# Force-Field Benchmarking by Alternatives: A Systematic Study of Ten Small α- and β-Proteins

**DOI:** 10.1101/2020.03.03.974477

**Authors:** Kalyanashis Jana, Kasper P. Kepp

## Abstract

Predicting protein structure from sequence is a central challenge of biochemistry, yet different force fields feature distinct structural biases that are hard to quantify, preventing clear assessment of results. Since structural transitions occur on milliseconds to seconds, sampling is out of reach in almost all routine studies, we inherently rely on local sampled structures, and benchmarks have emphasized the ability to reproduce these local structures. Here we approach the force field bias problem in a different way, via *alternatives*, by revisiting the old question: How unique is the sequence-structure relationship when studied computationally? To circumvent the sampling problem, the system-bias (specific structure choices affect apparent force field structural preference) and the complexity of tertiary structure, we studied ten small α- and β-proteins (20-35 amino acids) with one helix or sheet. For each of the ten sequences, we then designed alternative β- or α-structures and subjected all 20 proteins to molecular dynamics simulations. We apply this “alternative structure” benchmark to five of the best modern force fields: Amber ff99SB-ILDN, Amber ff99SB*-ILDN, CHARMM22*, CHARMM36, and GROMOS54A8. Surprisingly, we find that *all* sequences with reported β-structures also feature stable native-like α-structures with all five force fields. In contrast, only the alternative β-1T5Q and to some extent β-1CQ0 and β-1V1D resembled native β-proteins. With full phase space sampling being impossible in almost all cases, our benchmark by alternatives, which samples another local part of phase space in direct comparison, is a useful complement to millisecond benchmarks when these become more common.

## Introduction

Predicting protein structure from sequence is one of the main challenges of biochemistry and lends promise to major innovations in biotechnology.^1–5^ Computational techniques provide the necessary oversight and have guided the reported successes.^2,6–11^ By design, these previous successful case studies have assumed that the designed structure is unique. The studies do not reveal whether protein design is easy or difficult, since negative results rarely surface. In particular, they do not answer the question of *multiple minima*^12^: Can most amino acid sequences fold into different types of structures, depending on conditions? Karplus and co-workers originally explored this problem in 1984 by energy minimization of sea-worm hemerythrin and mouse immuno-globulin, finding that it is straightforward to produce alternative α- or β-structure, although the “designed” structures tend to be less ordered.^12^

Since this seminal work, the structural preferences of protein force fields have grown massively in relevance, as force fields are routinely used with molecular dynamics (MD) simulations to predict and rationalize protein structure and dynamics in thousands of studies each year.^5,6,13–17^ However, the timescales of folding-unfolding and structural transitions of small proteins are typically between milliseconds and seconds.^18,19^ Despite progress in computer power and dedicated software,^20^ and although many good force fields probably describe well the native structures when sampling is sufficient,^21^ sampling these timescales will not be routine for many years to come. This is particularly true for the 1000-fold shift from milliseconds to seconds that is relevant to most larger proteins. For all practical purposes, we must thus consider almost all MD studies non-ergodic, i.e. sampling a small local part of phase space close to the native structure, and absolute structure biases of force fields, which requires ergodic sampling, will be inaccessible until second-timescales become routine.

The problem is particularly relevant for small proteins that easily change structure. Some disordered proteins can exist with either α- or β-structure, a prominent example being β-amyloid involved in Alzheimer’s disease^22–26^. The sequence-structure relationship is evidently context dependent, and changes in the environment, e.g. heating or membrane interaction, commonly induce secondary structure changes^27–29^; it is also well-known that salt and pH can affect the structure of a sequence^30,31^, and even very similar sequences can in some cases feature quite distinct structures^32^. Adding to this complexity is the structure biases of modern protein force fields, which may predict another structure than the one seen experimentally^14,25,33,34^.

Because of the sampling problem, we propose to a new method of assessing relative structure preferences of force fields by “benchmarking of alternatives”. We systematically study the shortest possible protein sequences with well-defined secondary structure, since tertiary structure cannot be sampled well as the fold space contains thousands of alternative structures.^35^ Second, we systematically study both the native conformations, based on the experimental structures, and well-defined *alternative* α- or β-structures of the same sequences made by homology modeling. Comparison of the secondary structure obtained provides a way quantify relative structure biases of the force fields. The protocol amounts to sampling locally the two states of a structural transition between the native and alternative structure, comparing the ability of the force fields to maintain these structures nearly the minima.

We studied five small α-proteins and five small β-proteins with experimentally known structures, and from these ten sequences we designed alternative β- and α-structures using pair-wise structural homology (Figure 1). We then subjected all 20 protein structures to independent triplicate MD simulations using five modern force fields reported to work well for protein structure: Amber ff99SB-ILDN,^36^ Amber ff99SB*-ILDN,^37^ CHARMM22*,^13^ CHARMM36,^38,39^ and GROMOS 54A8.^37^ We specifically sought to answer the questions: i) How much does the ability to form the alternative structure depend on the force field? ii) How common is the multiple-minima problem at typical current force field and sampling quality? iii) Which differences in structural propensity is observed for the force fields across the ten native and ten alternative structures?

**Figure 1.**
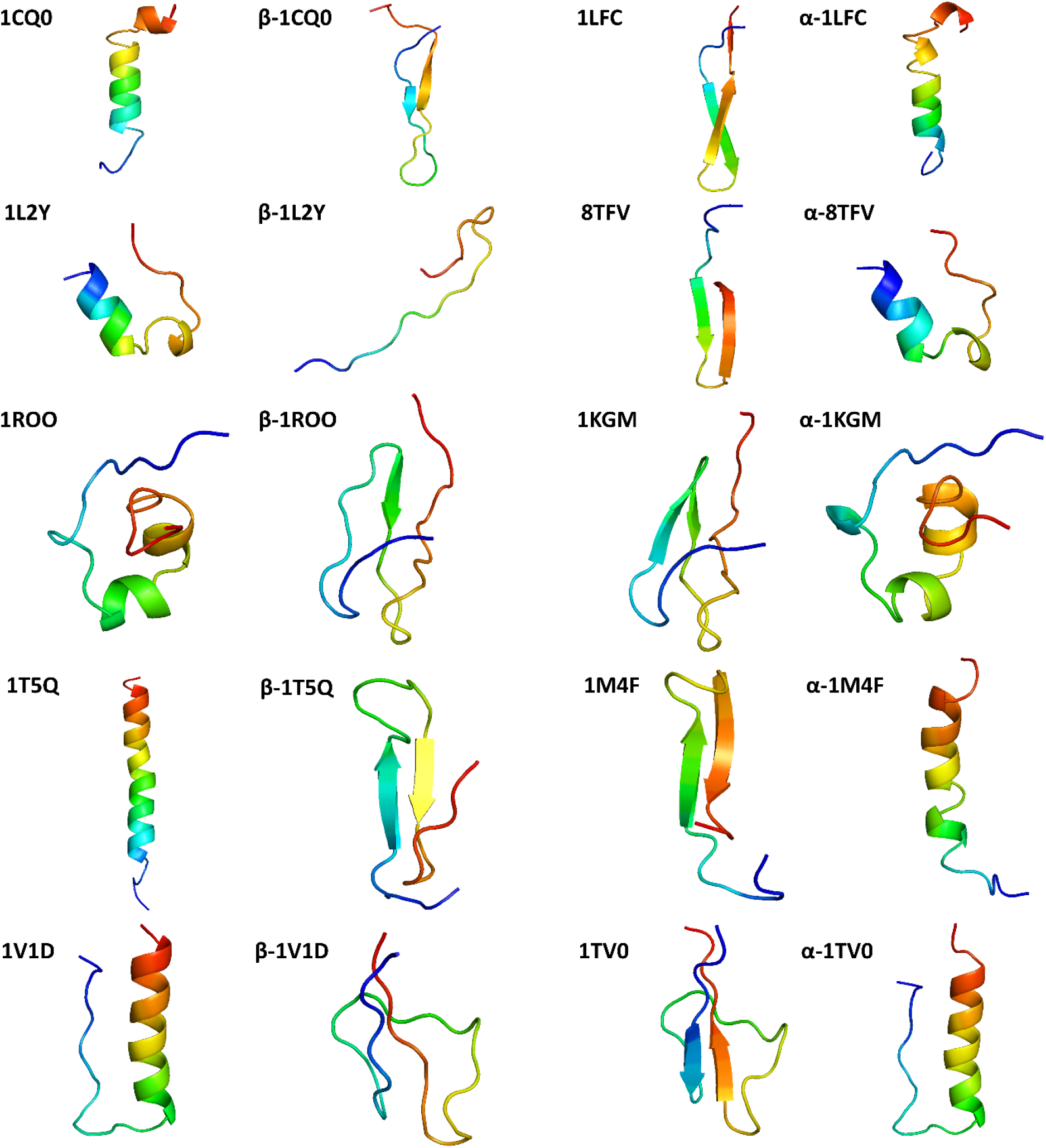
Studied small proteins and their alternative scaffolds. The α–proteins include human hypocretin-2/orexin-B (1CQ0), the TRP-cage miniprotein (1L2Y), ShK Toxin (1ROO), gastric inhibitory polypeptide (1T5Q), and pancreatic hormone (1V1D). The β–proteins include lactoferricin (1LFC), thanatin (8TFV), serine protease inhibitor I (1KGM), hepcidin (1M4F), and cryptdin-4 (1TV0). Proteins were designed from structure and sequence in the relation 1CQ0-1LFC, 1L2Y-8TFV, 1ROO-1KGM, 1T5Q-1M4F, 1V1D-1TV0.

## Computational methods

### Accuracy of molecular dynamic simulations

MD is a powerful tool for exploring protein structure and dynamics.^19,40–43^ MD is the only method that provides atomistic dynamic information at infinite resolution, whereas most experimental data are either from crystals or at lower resolution in solution. The three main approximations that reduce accuracy are the sampling extent, the realism of the chemical model, and the quality of the physical model (force field).

The required simulation time depends on the property of interest. Many modes have characteristic timescales of pico- and nanoseconds, including water-protein interactions, water-water interactions, and most protein-protein interactions^19^. However, conformational changes, loop motions, and partial unfolding events may occur on the millisecond to second time scales^19^. Thus, sub-second room temperature MD simulations cannot fully sample phase space, a fact known as the sampling or ergodicity problem^44,45^. Fortunately, MD starting from a folded stable structure typically probes the relevant fast dynamics of this subspace on the 10-100 nanosecond scale, unless major conformational changes are important^19,42^. Modern computers can routinely study this timescale without passing large diffusion barriers, explaining why MD has been successful when starting from experimental crystal or NMR structures. We utilize this fact to circumvent the need for millisecond sampling by constructing alternative starting structures of the β- and α-basins and test force field structural bias via the local stability of the alternative vs. native basins.

The realism of the chemical model, i.e. its chemical composition, is also an issue. Concentration and periodic artefacts are overcome by increasing the simulation cell size; recent studies disagree on the importance of the cell size beyond the standard size^46,47^. Another issue is salt: Many MD studies are performed in demineralized water despite experiments being typically performed near physiological ion strength, which affects protein conformation.^30,48^

The third approximation is the force field itself. Protein and water force fields represent different energy functions of structure and therefore estimate differently the thermal population of conformations. Specifically, water and protein force fields are not generally consistent^59^, and all commonly used force fields have fixed atomic charges that cannot polarize^40^. Accordingly, different force fields also favor certain types of secondary structure.^14,25,60–64^ Thus, the best force field depends on the properties of interest. If interested in free energies of hydration and interaction of ions and ligands, OPLS force fields^48,65,66^ commonly work well; if interested in secondary structure, improved versions of Charmm^13,67,68^ and Amber^36,69,70^ have been used widely. Amber ff99SB-ILDN and modified versions thereof have been recommended.^15,25,33,71^ Charmm22*^13^ and Charmm36^72^ are examples of structure-balanced Charmm force fields.

### Proteins and homology models studied

We considered five small α-proteins and five β-proteins with β-hairpins (Table 1). By some definitions (e.g. length less than 40 amino acids), these systems are called “peptides”, but as they have distinct, stable secondary structure others prefer to call them mini-proteins; we call them proteins below for consistency but could equally well have called them peptides. We collected the NMR structures from the protein data bank (PDB). All the proteins are complete and have 20-35 amino acids. Importantly, they either have one major helix or one main sheet composed of two β-strands, thereby in the cleanest way possible probing the secondary structure preferences of the sequences and force fields. Larger proteins, including those originally studied by Karplus and co-workers^12^, include tertiary structure and the potential to fold into many fold types, which makes any investigation subject to a large element of arbitrariness, even beyond the sampling issues associated with equilibrating such fold classes *de novo*.

**Table 1.** All the examined small α- and β-proteins details.

| α-proteins | | | | β-proteins | | | |
| --- | --- | --- | --- | --- | --- | --- | --- |
| Protein | PDB | Structure | Length | Protein | PDB | Structure | Length |
| Human hypocretin-2/orexin-B | 1CQ0<sup>49</sup> | α-helix | 28 | Bovine lactoferricin | 1LFC<sup>50</sup> | Tight β-hairpin | 25 |
| Designed Trp-Cage Miniprotein | 1L2Y<sup>51</sup> | α-helix, turn, coil | 20 | Protein (thanatin) | 8TFV<sup>52</sup> | Tight β-hairpin | 21 |
| ShK Toxin | 1ROO<sup>53</sup> | α-helix, major | 35 | Serine protease inhibitor I | 1KGM<sup>54</sup> | β-hairpin, turn, coil | 35 |
| Gastric inhibitory polypeptide | 1T5Q<sup>55</sup> | α-helix | 30 | Human hepcidin | 1M4F<sup>56</sup> | β-hairpin, coil | 25 |
| Designed esterase | 1V1D<sup>57</sup> | α-helix, turn, coil | 31 | Mouse cryptdin-4 | 1TV0<sup>58</sup> | β-hairpin, turn, coil | 32 |

The α-proteins are human hypocretin-2/orexin-B (PDB ID: 1CQ0)^49^, the designed TRP-case miniprotein (1L2Y)^51^, ShK toxin (1RRO)^53^, gastric inhibitory polypeptide (1T5Q)^55^, and the designed esterase (1V1D)^57^, whereas β-proteins are lactoferricin (1LFC)^50^, thanatin (8TFV)^52^, serine protease inhibitor I (1KGM)^54^, hepcidin (1M4F)^56^, and cryptdin-4 (1TV0)^58^. Obviously, some of the proteins may be more naturally inclined to display multiple structure via their sequence structural propensity: This real multiple-minimum tendency (or structural promiscuity) should emerge in the comparison of the systems when keeping the force field constant. In contrast, keeping the sequence constant while varying the force field will tell us about force field structural bias. These considerations are the main rationale behind the computational experiment suggested and carried out below.

We asked, along the line of Karplus and co-workers^12^, whether it is possible to define stable proteins with opposite structures from the same sequence. However to do this, we used modern force fields and MD simulation rather than energy minimization, and we studied ten native proteins each as templates for a new protein, by pair-wise structural homology based on similarity in sequence length only, using the MODELLER software^73^. For these small proteins with only one secondary structure element, the production of the alternative structure is the least arbitrary (larger proteins would create major issues in this regard, beyond the tertiary structure issue) and the approach is thus fully reproducible. To produce a consistent nomenclature, each designed protein is named according to the PDB identifier of its template, with α or β in front. All 20 native and template-modeled structures are shown in Figure 1, organized according to templates used. For example, β-1CQ0 is structurally similar to 1LFC whereas α-1LFC is structurally analogous to 1CQ0. The DSSP-calculated α-helix content is 57% for 1CQ0, 35% for 1L2Y, 31% for 1ROO, 76% for 1T5Q, and 55% for 1V1D. The β-sheet content is 24% for 1LFC, 38% for 8TFV, 11% for 1KGM, 32% for 1M4F, and 31% for 1TV0.

We found that the α-proteins were readily designable and reached significant α-helix content approaching that of the templates (Figure 1 right): 52% for α-1LFC, 24% for α-8TFV, 14% for α-1KGM, 52% for α-1M4F, and 47% for α-1TV0. We found that the design of the β-proteins was more difficult, both when optimizing and restricting the inter-strand hydrogen bonds, as the sheets put larger restrictions on the overall template-modeled structure, and thus the need for minimal RMSD and compact structure made the sheet tendency strongest in β-1T5Q and β-1V1D, with clear β-hairpin structures having 27% and 29% β-sheet content. β-1CQ0, β-1L2Y, and β-1ROO do not form full β-sheets but do feature significant β-bridge, except β-1L2Y, which was be modeled as a disordered folded protein.

### Molecular simulation details

All 20 proteins were subjected to MD simulation using five different force fields, i.e., Amber ff99SB-ILDN, Amber ff99SB*-ILDN, CHARMM22*, CHARMM36, and GROMOS 54A8 using GROMACS version 2018.5.^74^ All simulations included TIP3P water in neutral, cubic simulation boxes filled with Na^+^ or Cl^−^ ions to a final concentration of 0.15 M NaCl, with box sizes defined by a minimum of 10 Å from any box edge to the protein atoms. The total system was energy minimized using steepest-descent followed by full equilibration for 100 ps in the NVT ensemble and 100 ps in the NPT ensemble. The Verlet cut-off scheme was used for Coulomb and Lennard-Jones interactions with a cut-off of 10 Å. The Particle Mesh Ewald summation was employed to evaluate long-range electrostatic interactions, and the Linear Constraint Solver (LINCS) algorithm was used to constrain all covalent bonds during simulation.^75,76^ The 100 ps NVT simulations were performed using a V-rescale thermostat^77^ with a 0.1 ps temperature coupling constant at 300 K. The NPT simulations were carried out using the Parrinello−Rahman barostat^78^ at 1 atm with a 2 ps pressure coupling constant. For all 100 systems (20 proteins x 5 force fields), simulations were performed in triplicate by initiating them from three different random velocity seeds, and run for 100 ns each, for a total of 30 µs of simulation time; the simulation details are summarized in **Table S1**.

### Trajectory analysis

To understand the dynamical behavior caused by the different force fields, we monitored the evolution of protein secondary structural elements, the root mean square deviation (RMSD), root mean square fluctuation (RMSF), radius of gyration (R_g_), hydrogen bonding, solvent accessible surface area (SASA), and free energy landscape (FEL) of each simulation. A general summary of this analysis is given in **Table S2**. All the analyses were carried out using the full simulation trajectories since the proteins reached horizontal RMSD curves soon after MD simulation started, as expected from their small size (full curves in **Table S3**; averages in **Table S4**).

The FEL was plotted using the g_sham tool, based on the eigenvalues of principal components (PC) 1 and 2. The Gibb free energy was obtained as:

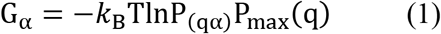

In this equation, G_α_ is the Gibbs free energy of a particular conformation state, *k*_B_ is the Boltzmann constant, T is the simulation temperature (300 K), P_(qα)_ is the probability density (number of counts along the trajectory), and P_max_(q) is the maximum of the probability density function.^79^

Contact analysis was carried out using the Arpeggio web server.^80^ The secondary structures (SS) were analyzed using the DSSP tool implemented in the GROMACS.^81^ In addition to the full trajectory analysis reported below, we also performed cluster analysis to provide a simple representation of the conformational ensemble and characterize the most representative structural states of each protein, using the cluster algorithm by Daura et al.^82^ implemented in GROMACS with a cut-off 0.2 nm.^82^

## Results and discussion

### Computed structures of native and alternative protein fold structures

Whereas Figure 1 presents the structures used in the study, Figure 2 summarizes the resulting simulated proteins as representative structures from cluster analysis (in green). The cyan structures are the experimental coordinates for the native proteins and the template coordinates for the alternative structures. Despite distinct differences between the five force fields, the native α- and β-proteins in their stable fold structures are well-modeled in all cases, with some variation in disordered parts as can be expected. In contrast, the designed β-proteins and α-proteins (Figure 2, right) differ substantially from the original templates showing the impact of all-atom MD equilibration of the alternative structures in solvent.

**Figure 2.**
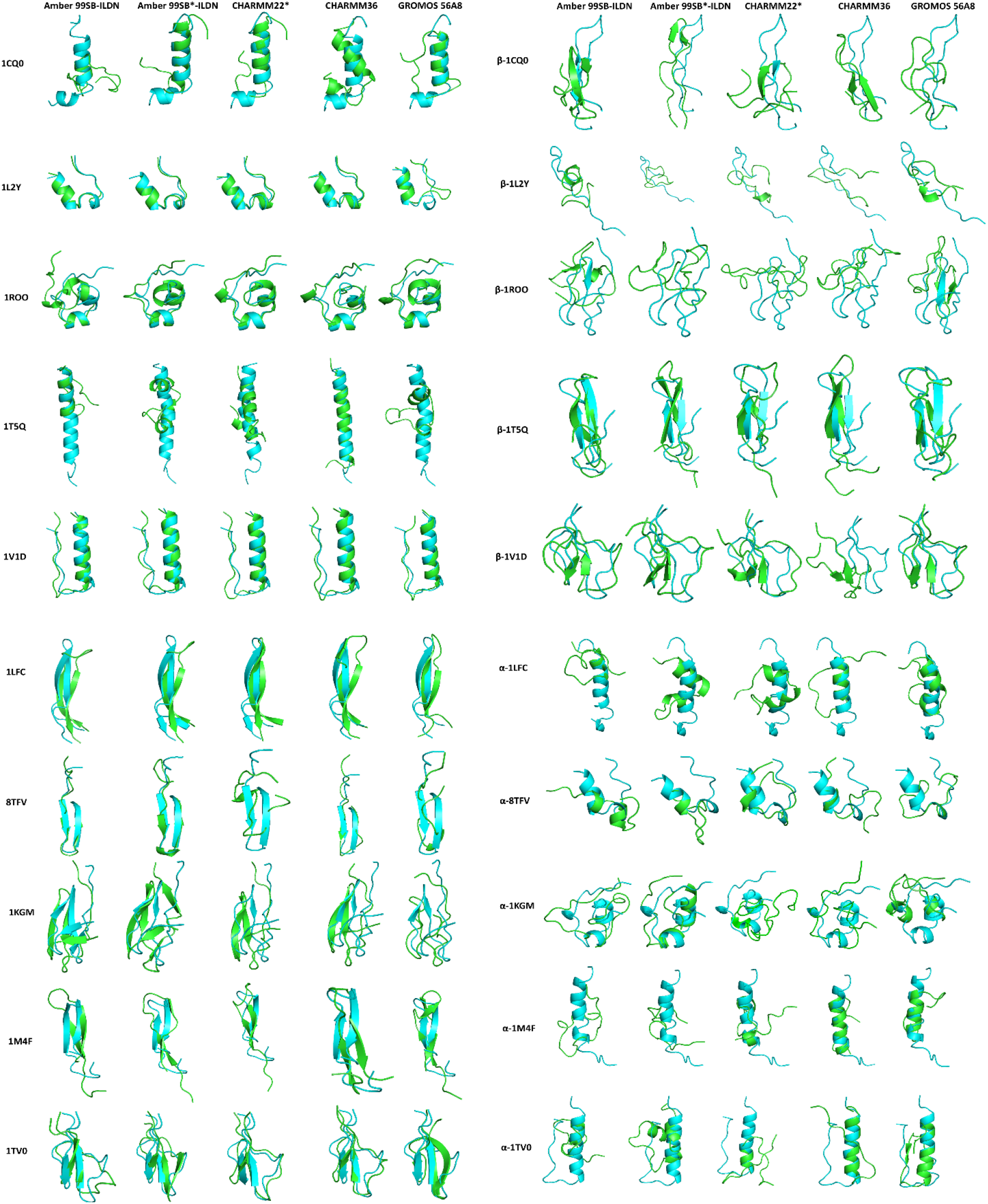
Comparison of experimental and simulated proteins using five force fields (green color). The cyan structures are experimental for the α-proteins (left) and modeled structures before simulation (right). The simulated structures are representative structures from cluster analysis.

Among the ten native proteins investigated in this work, the designed 20-residue Trp-cage miniprotein (1L2Y) has been extensively studied by many computational research groups as a test case of folding mechanisms and force field performance^83–87^. Recent computations suggest that Amber99SB*-ILDN produces excellent structures for this peptide^88^; several of the other force fields used here are also expected to produce structure-balanced ensembles, notably perhaps Charmm22*^13,25,33^. Previous studies focused on reproducing the experimentally observed helix-coil structure without account of false positives. We know today that false positives are common with many of the strand or helix-biased force fields available at the time of these studies, as shown by later benchmarks^14,25,33,34^. Yet the Trp-cage miniprotein has the clear advantage of being so small (20 residues, the smallest studied in the present work) that alternative β-structure becomes impossible, as seen from the five tested force fields all producing coil ensembles when starting closest possibly to a β-structure, which is very unstable due to the inability to form a stabilizing sheet in this short sequence (Figure 1).

The antimicrobial bovine milk protein lactoferricin (1LFC) has also been studied extensively^89,90^, and reversible α-β transitions have been suggested to explain the peptide’s *in vivo* function^50^. Previous MD-based conversion from β-hairpin to helix supports the possibility of dual structure tendency^90^. As seen from Figure 2, we can produce stable helix structure for this protein with all five tested force fields, but this was possible for *all* the five β-proteins and thus we conclude that local helix minima are not unique to lactoferricin but a common feature of small proteins.

We found that all five force fields are capable of producing stable α-proteins for all five sequences that are experimentally β-structures (Figure 2, right). In contrast, we only succeeded in designing one fully stable β-protein with a archetypical β-hairpin structure involving three strands (β-1T5Q) from a sequence with an experimental helix structure (1T5Q). The four other modeled β-proteins lost most strand during simulation, although all of them except β-1L2Y retained some β-structure. The ability of the force fields to stabilize the β-sheets differed substantially, as summarized below. The ten alternative structural ensembles contain more coil than the templates because much of the template-modeled secondary structure unfolds during simulation at 300 K. Despite this tendency, we observed some minor segments of the alternative secondary structure with all force fields.

We conclude that the force fields are well capable of reproducing the experimental structures in all ten cases. Most of the sequences have other minima in structure space of alternative structure type. Our results indicate that this is a rule rather than an exception, although the relative stability of the α- and β-structures will be very condition-dependent. We do not know if any of the alternative structures can be synthesized, but the α-structures may be stabilized specifically in nonpolar environments, as indicated by previous experimental structures of lactoferricin^50^. The alternative structures in all cases represent local minima that enable us to compare the force field secondary structure preferences in the same general ensembles.

### Properties of studied native and alternative structures

To characterize the structural ensembles produced by the five force fields we analyzed the secondary structure, RMSD, RMSF, R_g_, intra-protein hydrogen bonding, SASA, and FEL (Figures 3 and 4). The reported data are in all cases trajectory averages, subsequently averaged over all three independent simulations for each protein and force field.

**Figure 3.**
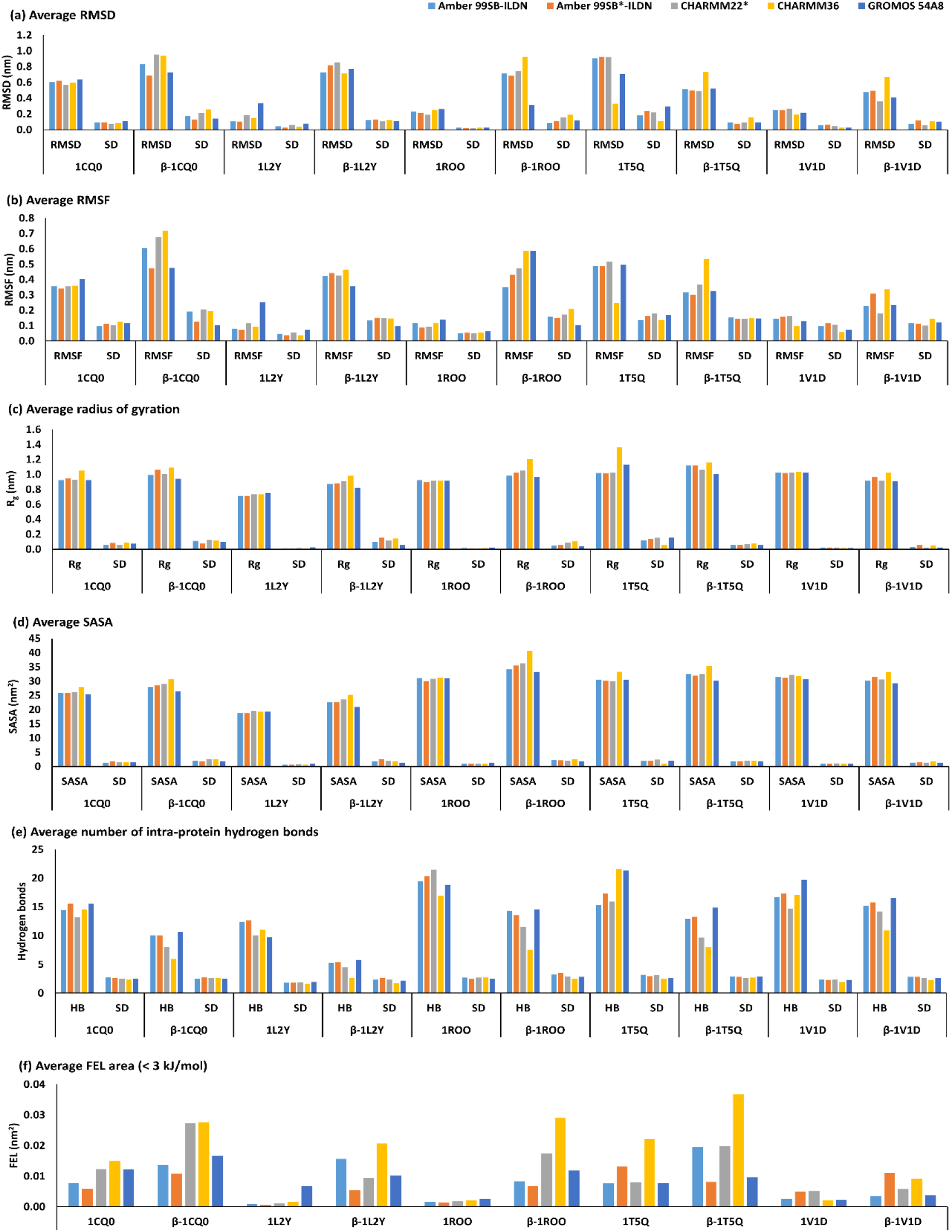
Simulated properties of α-proteins and artificial β-proteins averaged over three simulations, represented by the PDB code and force fields. (a) Average RMSD. (b) Average RMSF. (c) Average radius of gyration. (d) Average SASA. (e) Average intra-protein hydrogen bond. (f) Average area of free energy landscape within 3.0 kJ/mol free energy.

The five force fields generally produce very similar overall structural variations as quantified by the within-ensemble Cα RMSD (Figure 3a and 4a), although Charmm36 tended to produce significantly more structural variation than the other force fields (full data in **Tables S2-S4**). The alternative structures generally displayed higher structural variation consistent with their increased disorder. This was true both for the designed β-proteins (Figure 3a) and designed α-proteins (Figure 4a), the only exception being the gastric inhibitory peptide (1TQ5), which we succeeded in converting into a stable β-hairpin structure (Figure 2) with small disorder. The structural fluctuations (RMSF) strongly confirmed this picture (Figures 3b and 4b and **Tables S5-S6**).

**Figure 4.**
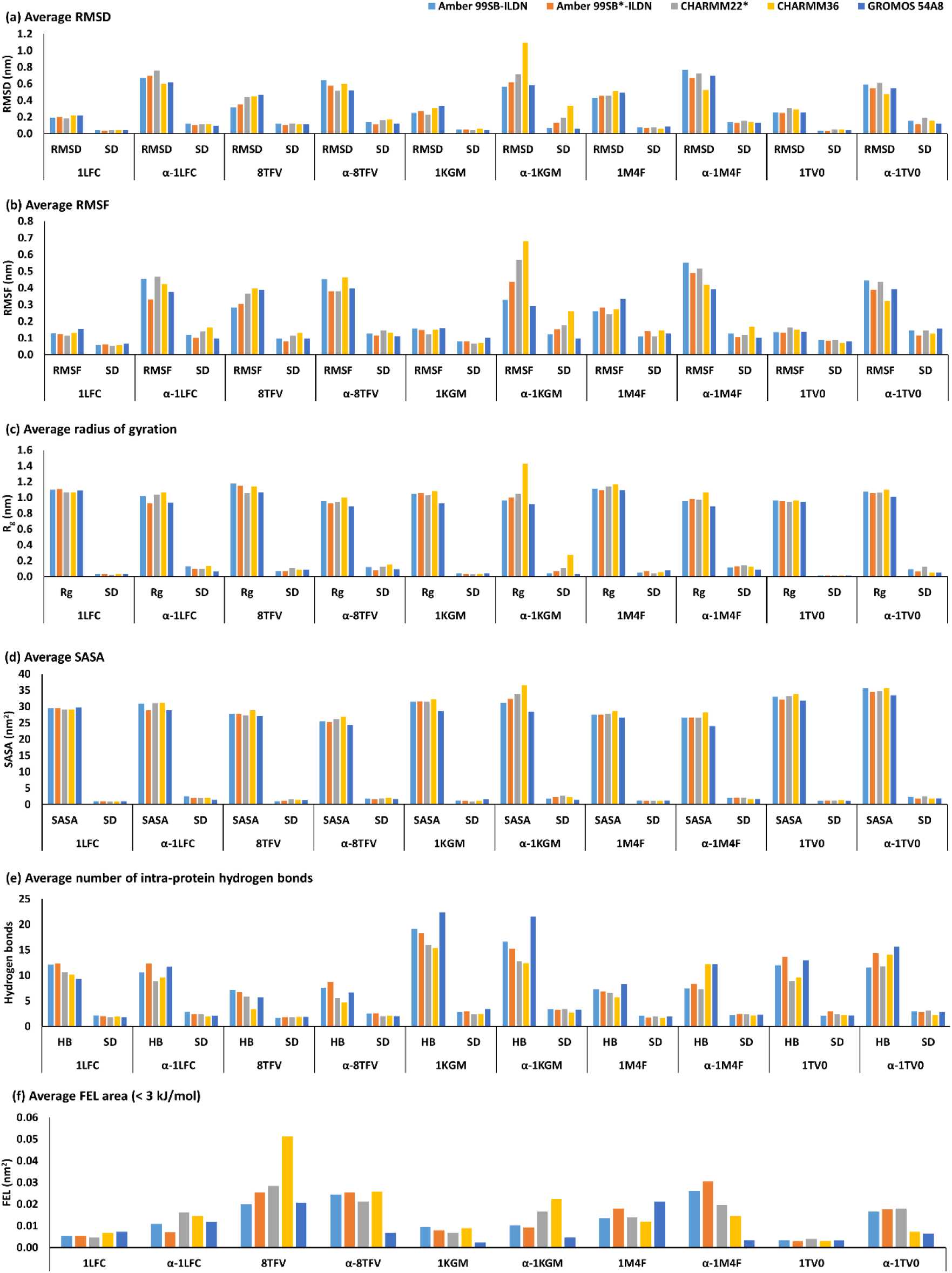
Simulated properties of β-proteins and artificial α-proteins averaged over three simulations, represented by the PDB code and force fields. (a) Average RMSD. (b) Average RMSF. (c) Average radius of gyration. (d) Average SASA. (e) Average intra-protein hydrogen bond. (f) Average area of free energy landscape within 3.0 kJ/mol free energy.

In terms of the protein sizes, the calculated average radius of gyration (R_g_) was very similar for the tested force fields, and the force field variations are clearly smaller than the peptide variations (Figures 3c and 4c and **Tables S7-S8**). The native proteins display average R_g_ of 7-11 Å. Again, the CHARMM36 force field produced consistently more expanded proteins R_g_ except for 1LFC and 8TFV. We note that the alternative and native structures have similar compactness in water when folded into secondary structure, but the structures that are disordered are slightly less compact. Thus, 1T5Q and β-1T5Q have similar compactness, and to some extent 1V1D and β-V1D, but the other designed β-proteins are larger (Figure 3d). In contrast, the designed α-proteins tended to be more compact than the native β-proteins, except for 1TV0/α-1TV0. Average solvent exposure (SASA) excellently confirmed these tendencies (Figures 3d and 4d and **Tables S9-S10**). In both cases, Charmm36 produced distinctly more expanded ensembles consistent with structural disorder (Figure 3 and 4).

An important property of the proteins is their ability to form and maintain intra-protein hydrogen bonds. The average number of intra-protein hydrogen bonds (Figures 3e and 4e and **Tables S11-S12**) show clearly the loss of intra-protein hydrogen bonding in the modeled β-proteins (Figure 3e). In contrast, the designed α-proteins did not display a loss of intra-protein hydrogen bonding averaged over the triplicate simulations (Figure 4e), quantifying the strong secondary structures formed in all cases (Figure 1). These findings were consistent for all studied force fields.

We constructed free energy landscapes (FEL) of the conformational ensembles of all three simulations of the 20 proteins with all five force fields, using a perimeter of 3.0 kJ/mol to define free energy basins (Figures 3f and 4f and **Tables S13-S14**). All the designed β-proteins displayed large free energy basins, consistent with their disordered ensembles. In contrast, the designed α-proteins displayed unique and diverse conformational states indicative of real folded protein ensembles. Consistent with the disorder documented above, CHARMM36 displayed larger basins, indicating a larger conformational entropy and weaker strength of the interactions. Amber ff99SB*-ILDN and GROMOS54A8 displayed the smallest entropy, consistent with their more compact ensembles. The details of the cluster analysis of the full trajectories, summarized in **Table S15**, show that the alternative structures typically feature more clusters than the native proteins, again driven by disordered ensembles.

### Secondary structure analysis

Finally, we analyzed the secondary structure in detail, by averaging the content of each DSSP-calculated structure type across the full trajectories of the triplicate simulations for each protein and force field (Figure 5). The native α-proteins are shown in Figure 5a, and the β-proteins designed for these sequences are shown in Figure 5b. Similarly, Figure 5c shows the native β-proteins, and Figure 5d shows the α-proteins designed from the sequences of the native β-proteins. The NMR-derived α-helix content of the native α-proteins is 57% for 1CQ0, 35% for 1L2Y, 31% for 1ROO, 76% for 1T5Q, and 55% for 1V1D; the experimentally estimated β-sheet content is 24% for 1LFC, 38% for 8TFV, 11% for 1KGM, 32% for 1M4F, and 31% for 1TV0 (Figure 5 and **Table S16**). We note that NMR-based secondary structure assignment is not very accurate but still provides an important comparison for the MD-simulated data, and previous authors have largely assumed that these structures are reasonable^83,84,90^. The good agreement between the NMR-derived and simulated secondary structure (Figure 5a and 5c) show that these comparisons are meaningful. Comparison of the corresponding ensemble-averaged residue-residue interactions (**Figure S1**) provides clear quantitative insight into the native-like properties of both classes of proteins, with a tendency towards less hydrophobic contacts in the alternative structures relative to the experimental structures mostly favored by the sequence; this findings substantially extends previous similar observations^12^ and supports the idea that the native protein structure is sequence-optimized to maximize hydrophobic packing in water.

**Figure 5.**
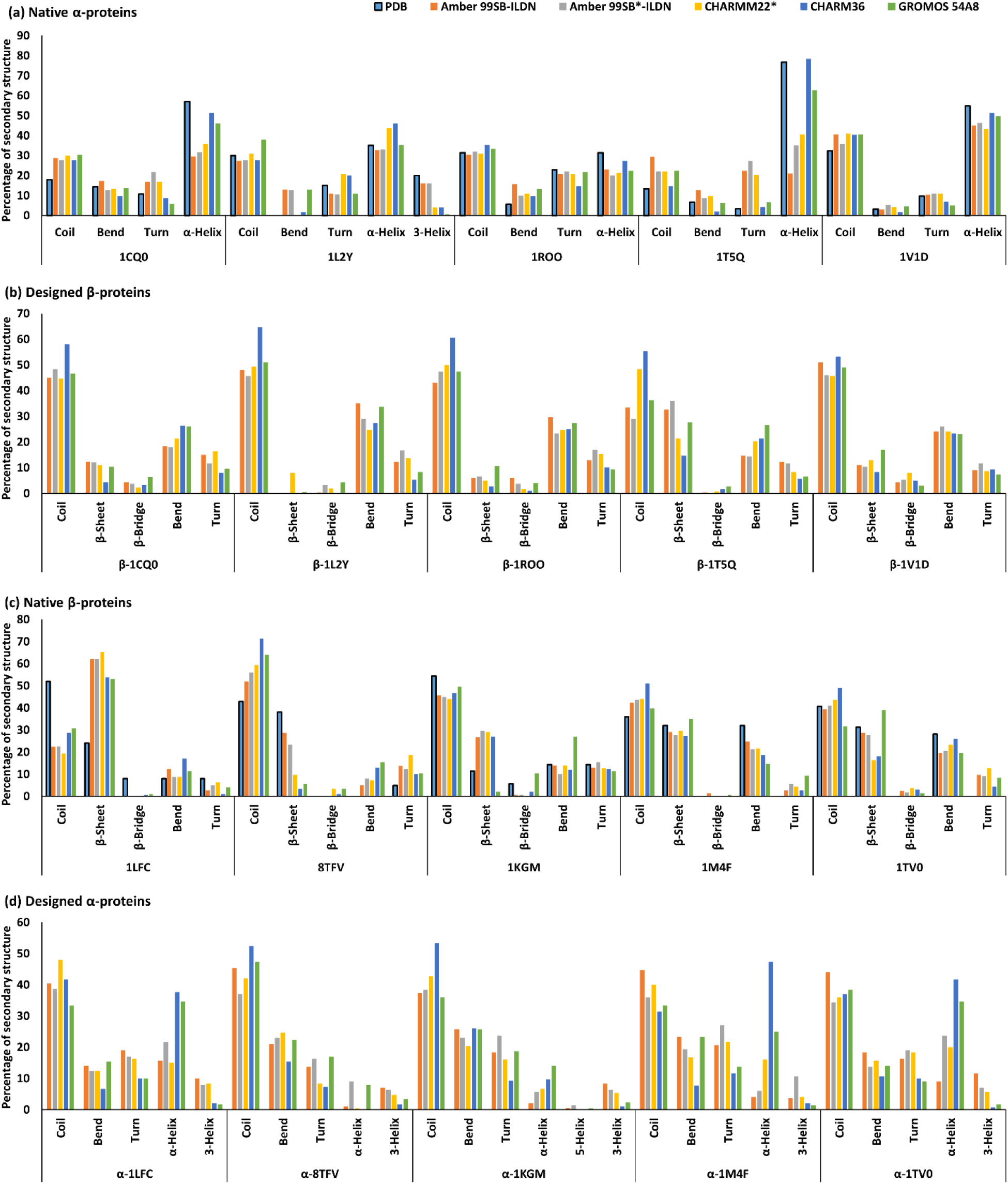
Percentage of coil, β-sheet, β-bridge, bend, turn, α-helix, and 3_10_-helix averaged over three simulations. (a) Native α-proteins. (b) Designed β-proteins. (c) Native β-proteins. (d) Designed α-proteins. The experimental secondary structure is labeled by PDB code, and the calculated secondary structure is labelled after force field.

For the native α-proteins (Figure 5a), three of the force fields, Amber99sb-ILDN, Amber99sb*-ILDN, and Charmm22*, tend to produce consistently less helix over all triplicate simulations than the Charmm36 and Gromos54A8 force fields, quantifying their different energy functions which stabilize helix torsion angles and hydrogen bonds differently. For 1CQ0 and 1T5Q, only the helix-stabilizing force fields Charmm36 and Gromos54A8 have similar amount of helix as the NMR models, whereas for the other native α-proteins, agreement is good for all force fields, considering the uncertainties in assigning helix structure from NMR experiments. Describing the helix content of 1T5Q is a major challenge for MD, as the results are very sensitive to force field.

In the case of the native β-proteins (Figure 5c), the β-sheet content and coil correlate inversely due to persistent β-coil transitions during the simulation time, indicating good sampling of these changes. The β-sheet propensity is generally similar for the five tested force fields. All force fields predict more sheet in 1LFC than suggested by the NMR model; for 8TFV, the opposite is the case, but otherwise the force fields span the experimental range well, and the relative amount of coil and sheet is quite well described in all other cases.

Of the five designed β-proteins (Figure 5b), β-1CQ0, β-1T5Q, and β-1V1D maintained a significant amount, more than 10%, of sheet structure and no helix structure in all triplicate simulations for most force fields, in particular our designed β-1T5Q, which has properties very similar to a native β-protein with up to 30% sheet structure. In contrast we did not observe significantly stable sheet structure in β-1ROO and β-1L2Y although these coil-rich ensembles did feature β-bridges and small strand segments. Also, the ability to preserve alternative structures is in some cases quite force field dependent; for example, Charmm22* is the only force field that, when averaged over three simulations, can produce some sheet in β-1L2Y. Except for β-1T5Q, which illustrates well a dual-structure peptide, the other four designed peptides in Figure 5b are disordered with mainly coil and little secondary structure. This may imply that these did not have energy minima strong enough to enable a β-structure basin in the free energy landscape.

Finally, all the five designed α-proteins (Figure 5d) contained significant α-helix content consistent with typical native α-proteins of the same size (Figure 5a). The triplicate ensemble-averaged α-helix content is 52% for α-1LFC, 24% for α-8TFV, 14% for α-1KGM, 52% for α-1M4F, and 47% for α-1TV0. The designed α-proteins excellently confirm the observation from the native proteins (Figure 5a) that Charmm36 and Gromos54A8 have substantially more helix propensity than the three other studied force fields. Previous versions of Gromos tended to favor sheet and disfavor helix^14^, so this could possibly be an overcorrection in the new version of Gromos.

## Conclusions

Delicate sequence-structure relationships and multiple-minima of real proteins,^32^ a problem of substantial biological interest, may be studied by from comparison of different proteins when keeping the force field constant. In contrast, keeping the sequence constant while varying the force field should tell us about force field structural bias, a problem of considerable technical interest. Karplus and co-workers originally explored this problem in 1984 by energy minimization of sea-worm hemerythrin and mouse immuno-globulin, showing that they could obtain minimized alternative α- or β-structures.^12^ While this study energy minimized rather than sampled the local minima, it indicated the weaknesses in force fields and spurred major interest in improving them.

Today, we have sophisticated computers and carefully developed all-atom protein force fields available, yet the problem remains very hard to assess: With relevant timescales approaching seconds, almost all MD simulations even today are non-ergodic, strictly preventing an assessment of these two problems. Also, force field biases for a few specific native structures may be system-dependent, i.e. coincidentally large or small for the studied systems. These considerations are the main rationale behind the computational experiment carried out in this work, which we refer to as “benchmarking by alternatives”, i.e. comparing structures sampled by the force fields locally near *both* the native structure and near a well-defined alternative α- or β-structure.

Since the fold space includes thousands of structural folds, which is also beyond any sampling, we studied five small α-proteins, 1CQ0, 1L2Y, 1ROO, 1T5Q, and 1V1D, and five small β-proteins, 1LFC, 8TFV, 1KGM, 1M4F, and 1TV0, with only one secondary structure element to avoid artefacts from arbitrary tertiary structures. We then designed alternative β- and α-structures by simple pairwise structural homology using the template of the alternative structure closest in sequence length. We tested five force fields completely in parallel: Amber ff99SB-ILDN, Amber ff99SB*-ILDN, CHARMM22*, CHARMM36, and GROMOS 54A8. We subjected all 100 systems (20 proteins studied with five force fields each) to triplicate 100-ns MD simulations, which is the practical time scale of most applied studies. Mots importantly, our protocol circumvents the need for millisecond-to-second sampling, which is virtually never met, by comparing directly the alternative structures of the β- and α-basins produces by the force fields.

We find that most of the studied force fields maintain stable alternative α- or β-structures. We were able to design all the alternative α-proteins from sequences reported to have experimental β-structures. Our alternative structures β-1T5Q and β-1V1D were generally stable for most examined force fields. While this says little about real structural preferences, which are typically very condition-dependent, it illustrates the local sampling problem already with secondary structure in small sequences of 20-35 amino acids. For larger tertiary structures, particular structural folds encounter many more local minima, which will be rarely accessible at the all-atom level for many years to come.^19,40^ Instead, our study provides detailed information on the structural preferences of the force fields via their ability to stabilize the native and alternative structures in strict comparison, as analyzed in the paper. The tendency to maintain alternative secondary structure was taken as indication of a preference towards that type of structure in the force field.

As all other studies (except perhaps those studying the fast-folding Trp-cage miniprotein, which folds on the microsecond time scale) our sampling is non-ergodic. However, considering the weaknesses in current protein force fields, chemical model realism, and structure sensitivity to temperature, salt, co-solvents, and pH, and since MD will not enjoy second timescales routinely in the coming years, we recommend benchmarking by alternatives for several force fields as a supplement to current simulations, even those that sample the native states for milliseconds, as our protocol may be the only way to quantify α-β preferences in practice.

## Supplementary material

The supporting information file contains all the time-evolution plots for all 300 MD simulations of the RMSD, RMSF, R_g_, SASA, and intra-protein hydrogen bonds, the full free energy landscapes for all simulations, and tables with the main averaged properties derived form each simulation, averaged over full trajectories performed in triplicate throughout.

## Author contributions

K. Y. performed all simulations and most of the analysis and wrote the first draft of the paper. K.P.K. planned the project, analyzed results, and co-wrote the paper.

## Funding

Authors are grateful to the Danish Council for Independent Research | Natural Sciences (DFF) for supporting this work (grant case 8022-00041B).

## Acknowledgments

We acknowledge the use of the Steno Cluster at DTU Chemistry, originally funded by the Danish Council for Independent Research, and the high-performance-computing cluster facility of DTU. The authors declare that they have no conflicts of interest associated with this work.

## References

1. Regan, L. & Degrado, W. F. Characterization of a helical protein designed from first principles. Science (80-.). 241, 976–978 (1988).

2. Koga, N. et al. Principles for designing ideal protein structures. Nature 491, 222–227 (2012).

3. Huang, P. S. et al. High thermodynamic stability of parametrically designed helical bundles. Science (80-.). 346, 481–485 (2014).

4. Huang, P.-S., Boyken, S. E. & Baker, D. The coming of age of de novo protein design. Nature 537, 320 (2016).

5. Skolnick, J., Kolinski, A., Brooks III, C. L., Godzik, A. & Rey, A. A method for predicting protein structure from sequence. Curr. Biol. 3, 414–423 (1993).

6. Lapidoth, G. D. et al. AbDesign: An algorithm for combinatorial backbone design guided by natural conformations and sequences. Proteins Struct. Funct. Bioinforma. 83, 1385–1406 (2015).

7. Brunette, T. J. et al. Exploring the repeat protein universe through computational protein design. Nature 528, 580–584 (2015).

8. Siegel, J. B. et al. Computational protein design enables a novel one-carbon assimilation pathway. Proc. Natl. Acad. Sci. 201500545 (2015). doi:10.1073/pnas.1500545112

9. Doyle, L. et al. Rational design of α-helical tandem repeat proteins with closed architectures. Nature 528, 585–588 (2015).

10. Boyken, S. E. et al. De novo design of protein homo-oligomers with modular hydrogen-bond network-mediated specificity. Science (80-.). 352, 680–687 (2016).

11. Woolfson, D. N. et al. De novo protein design: how do we expand into the universe of possible protein structures? Curr. Opin. Struct. Biol. 33, 16–26 (2015).

12. Novotný, J., Bruccoleri, R. & Karplus, M. An analysis of incorrectly folded protein models: implications for structure predictions. J. Mol. Biol. 177, 787–818 (1984).

13. Piana, S., Lindorff-Larsen, K. & Shaw, D. E. How robust are protein folding simulations with respect to force field parameterization? Biophys. J. 100, 47–49 (2011).

14. Best, R. B., Buchete, N.-V. & Hummer, G. Are current molecular dynamics force fields too helical? Biophys. J. 95, L07–L09 (2008).

15. Beauchamp, K. A., Lin, Y. S., Das, R. & Pande, V. S. Are protein force fields getting better? A systematic benchmark on 524 diverse NMR measurements. J. Chem. Theory Comput. 8, 1409–1414 (2012).

16. Allison, B. et al. Computational design of protein-small molecule interfaces. J. Struct. Biol. 185, 193–202 (2014).

17. Fiser, A., Feig, M., Brooks, C. L. & Sali, A. Evolution and physics in comparative protein structure modeling. Acc. Chem. Res. 35, 413–421 (2002).

18. Zwier, M. C. & Chong, L. T. Reaching biological timescales with all-atom molecular dynamics simulations. Curr. Opin. Pharmacol. 10, 745–752 (2010).

19. Karplus, M. & McCammon, J. A. Molecular dynamics simulations of biomolecules. Nat. Struct. Biol. 9, 646–652 (2002).

20. Klepeis, J. L., Lindorff-Larsen, K., Dror, R. O. & Shaw, D. E. Long-timescale molecular dynamics simulations of protein structure and function. Current Opinion in Structural Biology 19, 120–127 (2009).

21. Lindorff-Larsen, K., Piana, S., Dror, R. O. & Shaw, D. E. How fast-folding proteins fold. Science (80-.). 334, 517–520 (2011).

22. Xu, Y. et al. Conformational transition of amyloid beta-peptide. Proc. Natl. Acad. Sci. U. S. A. 102, 5403–5407 (2005).

23. Kotler, S. A., Walsh, P., Brender, J. R. & Ramamoorthy, A. Differences between amyloid-β aggregation in solution and on the membrane: insights into elucidation of the mechanistic details of Alzheimer’s disease. Chem. Soc. Rev. 43, 8–10 (2014).

24. Somavarapu, A. K. & Kepp, K. P. Direct Correlation of Cell Toxicity to Conformational Ensembles of Genetic Aβ Variants. ACS Chem. Neurosci. 6, 1990–1996 (2015).

25. Somavarapu, A. K. & Kepp, K. P. The Dependence of Amyloid-β Dynamics on Protein Force Fields and Water Models. ChemPhysChem 16, 3278–3289 (2015).

26. Vivekanandan, S., Brender, J. R., Lee, S. Y. & Ramamoorthy, A. A partially folded structure of amyloid-β(1-40) in an aqueous environment. Biochem. Biophys. Res. Commun. 411, 312–316 (2011).

27. Korshavn, K. J., Bhunia, A., Lim, M. H. & Ramamoorthy, A. Amyloid-β adopts a conserved, partially folded structure upon binding to zwitterionic lipid bilayers prior to amyloid formation. Chem. Commun. 52, 882–885 (2016).

28. Tomaselli, S. et al. The α-to-β conformational transition of Alzheimer’s Aβ-(1-42) peptide in aqueous media is reversible: A step by step conformational analysis suggests the location of β conformation seeding. ChemBioChem 7, 257–267 (2006).

29. Dasmeh, P. & Kepp, K. P. Unfolding Simulations of Holomyoglobin from Four Mammals: Identification of Intermediates and β-Sheet Formation from Partially Unfolded States. PLoS One 8, e80308 (2013).

30. Zhang, Y. & Cremer, P. S. Interactions between macromolecules and ions: the Hofmeister series. Curr. Opin. Chem. Biol. 10, 658–663 (2006).

31. Okur, H. I. et al. Beyond the Hofmeister Series: Ion-Specific Effects on Proteins and Their Biological Functions. J. Phys. Chem. B 121, 1997–2014 (2017).

32. He, Y., Chen, Y., Alexander, P., Bryan, P. N. & Orban, J. NMR structures of two designed proteins with high sequence identity but different fold and function. Proc. Natl. Acad. Sci. 105, 14412–14417 (2008).

33. Lindorff-Larsen, K. et al. Systematic validation of protein force fields against experimental data. PLoS One 7, 1–6 (2012).

34. Cino, E. A., Choy, W. & Karttunen, M. Comparison of Secondary Structure Formation Using 10 Different Force Fields in Microsecond Molecular Dynamics Simulations. J. Chem. Theory Comput. 8, 2725–2740 (2012).

35. Marks, D. S. et al. Protein 3D structure computed from evolutionary sequence variation. PLoS One 6, e28766 (2011).

36. Lindorff-Larsen, K. et al. Improved side-chain torsion potentials for the Amber ff99SB protein force field. Proteins Struct. Funct. Bioinforma. 78, 1950–1958 (2010).

37. Aliev, A. E. et al. Motional timescale predictions by molecular dynamics simulations: Case study using proline and hydroxyproline sidechain dynamics. Proteins Struct. Funct. Bioinforma. 82, 195–215 (2014).

38. Best, R. B. et al. Optimization of the additive CHARMM all-atom protein force field targeting improved sampling of the backbone φ, ψ and side-chain χ1 and χ2 Dihedral Angles. J. Chem. Theory Comput. 8, 3257–3273 (2012).

39. Huang, J. et al. CHARMM36m: An improved force field for folded and intrinsically disordered proteins. Nat. Methods 14, 71–73 (2016).

40. Adcock, S. A. & McCammon, J. A. Molecular dynamics: survey of methods for simulating the activity of proteins. Chem. Rev. 106, 1589–1615 (2006).

41. Voelz, V. A., Bowman, G. R., Beauchamp, K. & Pande, V. S. Molecular simulation of ab initio protein folding for a millisecond folder NTL9(1-39). J. Am. Chem. Soc. 132, 1526–1528 (2010).

42. van Gunsteren, W. F. & Berendsen, H. J. C. Computer simulation of molecular dynamics: Methodology, applications, and perspectives in chemistry. Angew. Chemie Int. Ed. English 29, 992–1023 (1990).

43. Beck, D. A. C. & Daggett, V. Methods for molecular dynamics simulations of protein folding/unfolding in solution. Methods 34, 112–120 (2004).

44. Clarage, J. B., Romo, T., Andrews, B. K., Pettitt, B. M. & Phillips, G. N. A sampling problem in molecular dynamics simulations of macromolecules. Proc. Natl. Acad. Sci. 92, 3288–3292 (1995).

45. Stella, L. & Melchionna, S. Equilibration and sampling in molecular dynamics simulations of biomolecules. J. Chem. Phys. 109, 10115–10117 (1998).

46. Mehra, R. & Kepp, K. P. Cell size effects in the molecular dynamics of the intrinsically disordered Aβ peptide. J. Chem. Phys. 151, 85101 (2019).

47. El Hage, K., Hedin, F., Gupta, P. K., Meuwly, M. & Karplus, M. Valid molecular dynamics simulations of human hemoglobin require a surprisingly large box size. Elife 7, e35560 (2018).

48. Jensen, K. P. & Jorgensen, W. L. Halide, ammonium, and alkali metal ion parameters for modeling aqueous solutions. J. Chem. Theory Comput. 2, 1499–1509 (2006).

49. Lee, J. H. et al. Solution structure of a new hypothalamic neuropeptide, human hypocretin-2/orexin-B. Eur. J. Biochem. 266, 831–839 (1999).

50. Hwang, P. M., Zhou, N., Shan, X., Arrowsmith, C. H. & Vogel, H. J. Three-Dimensional Solution Structure of Lactoferricin B, an Antimicrobial Peptide Derived from Bovine Lactoferrin. Biochemistry 37, 4288–4298 (1998).

51. Neidigh, J. W., Fesinmeyer, R. M. & Andersen, N. H. Designing a 20-residue protein. Nat. Struct. Biol. 9, 425–430 (2002).

52. Mandard, N. et al. Solution structure of thanatin, a potent bactericidal and fungicidal insect peptide, determined from proton two-dimensional nuclear magnetic resonance data. Eur. J. Biochem. 256, 404–410 (1998).

53. Tudor, J. E., Pallaghy, P. K., Pennington, M. W. & Norton, R. S. Solution structure of ShK toxin, a novel potassium channel inhibitor from a sea anemone. Nat. Struct. Biol. 3, 317–320 (1996).

54. Gáspári, Z., Patthy, A., Gráf, L. & Perczel, A. Comparative structure analysis of proteinase inhibitors from the desert locust, Schistocerca gregaria. Eur. J. Biochem. 269, 527–537 (2002).

55. Alaña, I. et al. NMR structure of the glucose-dependent insulinotropic polypeptide fragment, GIP(1-30)amide. Biochem. Biophys. Res. Commun. 325, 281–286 (2004).

56. Hunter, H. N., Fulton, D. B., Ganz, T. & Vogel, H. J. The solution structure of human hepcidin, a peptide hormone with antimicrobial activity that is involved in iron uptake and hereditary hemochromatosis. J. Biol. Chem. 277, 37597–603 (2002).

57. Nicoll, A. J. & Allemann, R. K. Nucleophilic and general acid catalysis at physiological pH by a designed miniature esterase. Org. Biomol. Chem. 2, 2175–80 (2004).

58. Jing, W., Hunter, H. N., Tanabe, H., Ouellette, A. J. & Vogel, H. J. Solution structure of cryptdin-4, a mouse paneth cell α-defensin. Biochemistry 43, 15759–15766 (2004).

59. Jensen, K. P. Improved interaction potentials for charged residues in proteins. J. Phys. Chem. B 112, 1820–7 (2008).

60. Freddolino, P. L., Park, S., Roux, B. & Schulten, K. Force Field Bias in Protein Folding Simulations. Biophys. J. 96, 3772–3780 (2009).

61. Mittal, J. & Best, R. B. Tackling force-field bias in protein folding simulations: Folding of Villin HP35 and Pin WW domains in explicit water. Biophys. J. 99, L26–L28 (2010).

62. Best, R. B. & Mittal, J. Free-energy landscape of the GB1 hairpin in all-atom explicit solvent simulations with different force fields: Similarities and differences. Proteins Struct. Funct. Bioinforma. 79, 1318–1328 (2011).

63. Ulmschneider, M. B. & Ulmsehneider, J. P. Folding peptides into lipid bilayer membranes. J. Chem. Theory Comput. 4, 1807–1809 (2008).

64. Shaw, D. E. et al. Atomic-level characterization of the structural dynamics of proteins. Science (80-.). 330, 341–346 (2010).

65. Harder, E. et al. OPLS3: A Force Field Providing Broad Coverage of Drug-like Small Molecules and Proteins. J. Chem. Theory Comput. 12, 281–296 (2016).

66. Jorgensen, W. L. & Tirado-Rives, J. The OPLS [optimized potentials for liquid simulations] potential functions for proteins, energy minimizations for crystals of cyclic peptides and crambin. J. Am. Chem. Soc. 110, 1657–1666 (1988).

67. MacKerell, A. D. J. et al. All-atom empirical potential for molecular modeling and dynamics studies of proteins. J. Phys. Chem. B 102, 3586–616 (1998).

68. Mackerell, A. D., Feig, M. & Brooks, C. L. Extending the treatment of backbone energetics in protein force fields: Limitations of gas-phase quantum mechanics in reproducing protein conformational distributions in molecular dynamics simulation. J. Comput. Chem. 25, 1400–1415 (2004).

69. Maier, J. A. et al. ff14SB: Improving the Accuracy of Protein Side Chain and Backbone Parameters from ff99SB. J. Chem. Theory Comput. 11, 3696–3713 (2015).

70. Hornak, V. et al. Comparison of multiple amber force fields and development of improved protein backbone parameters. Proteins: Structure, Function and Genetics 65, 712–725 (2006).

71. Lange, O. F., van der Spoel, D. & de Groot, B. L. Scrutinizing Molecular Mechanics Force Fields on the Submicrosecond Timescale with NMR Data. Biophys. J. 99, 647–655 (2010).

72. Huang, J. & MacKerell Jr, A. D. CHARMM36 all-atom additive protein force field: Validation based on comparison to NMR data. J. Comput. Chem. 34, 2135–2145 (2013).

73. Webb, B. & Sali, A. Protein structure modeling with MODELLER. in Methods in Molecular Biology 1654, 39–54 (Humana Press, New York, NY, 2017).

74. GROMACS 2018. 5 program package, freely available from the GROMACS Web site.

75. Darden, T., York, D. & Pedersen, L. Particle mesh Ewald: An N log(N) method for Ewald sums in large systems. J. Chem. Phys. 98, 10089 (1993).

76. Hess, B., Bekker, H., Berendsen, H. J. C. & Fraaije, J. G. E. M. LINCS: A Linear Constraint Solver for molecular simulations. J. Comput. Chem. 18, 1463–1472 (1997).

77. Bussi, G., Donadio, D. & Parrinello, M. Canonical sampling through velocity rescaling. J. Chem. Phys. 126, 014101 (2007).

78. Parrinello, M. & Rahman, A. Polymorphic Transitions in Single Crystals: a New Molecular Dynamics Method. J. Appl. Phys. 52, 7182–7190 (1981).

79. Singh, A., Somvanshi, P. & Grover, A. Drug repurposing against arabinosyl transferase (EmbC) of Mycobacterium tuberculosis: Essential dynamics and free energy minima based binding mechanics analysis. Gene 693, 114–126 (2019).

80. Jubb, H. C. et al. Arpeggio: A Web Server for Calculating and Visualising Interatomic Interactions in Protein Structures. J. Mol. Biol. 429, 365–371 (2017).

81. Kabsch, W. & Sander, C. Dictionary of protein secondary structure: pattern recognition of hydrogen-bonded and geometrical features. Biopolymers 22, 2577–2637 (1983).

82. Daura, X. et al. Peptide Folding: When Simulation Meets Experiment. Angew. Chemie Int. Ed. 38, 236–240 (1999).

83. Zhou, R. Trp-cage: Folding free energy landscape in explicit water. Proc. Natl. Acad. Sci. U. S. A. 100, 13280–13285 (2003).

84. Paschek, D., Nymeyer, H. & García, A. E. Replica exchange simulation of reversible folding/unfolding of the Trp-cage miniprotein in explicit solvent: On the structure and possible role of internal water. J. Struct. Biol. 157, 524–533 (2007).

85. Snow, C. D., Zagrovic, B. & Pande, V. S. The Trp cage: folding kinetics and unfolded state topology via molecular dynamics simulations. J. Am. Chem. Soc. 124, 14548–14549 (2002).

86. Chowdhury, S., Lee, M. C., Xiong, G. & Duan, Y. Ab initio folding simulation of the Trp-cage mini-protein approaches NMR resolution. J. Mol. Biol. 327, 711–717 (2003).

87. Pitera, J. W. & Swope, W. Understanding folding and design: Replica-exchange simulations of``Trp-cage’’miniproteins. Proc. Natl. Acad. Sci. 100, 7587–7592 (2003).

88. Robustelli, P., Piana, S. & Shaw, D. E. Developing a molecular dynamics force field for both folded and disordered protein states. Proc. Natl. Acad. Sci. U. S. A. 115, E4758–E4766 (2018).

89. Daidone, I. et al. Conformational study of bovine lactoferricin in membrane-micking conditions by molecular dynamics simulation and circular dichroism. Biometals 24, 259–268 (2011).

90. Zhou, N., Tieleman, D. P. & Vogel, H. J. Molecular dynamics simulations of bovine lactoferricin: turning a helix into a sheet. Biometals 17, 217–223 (2004).

