## Supporting information for "Force-Field Benchmarking by Alternatives: A Systematic Study of Ten Small α- and β-Proteins"

**Table S1**. Simulation details for native α-proteins, modeled β-proteins, β-proteins, and modeled α-proteins.

| | Amber ff99SB-ILDN | Amber ff99SB*-ILDN | CHARMM22* | CHARMM36 | GROMOS 54A8 | Total simulation time (μs) |
|---|---|---|---|---|---|---|
| | Seeds | Seeds | Seeds | Seeds | Seeds | |
| 1CQ0 | 3 | 3 | 3 | 3 | 3 | 1.5 |
| 1L2Y | 3 | 3 | 3 | 3 | 3 | 1.5 |
| 1ROO | 3 | 3 | 3 | 3 | 3 | 1.5 |
| 1T5Q | 3 | 3 | 3 | 3 | 3 | 1.5 |
| 1V1D | 3 | 3 | 3 | 3 | 3 | 1.5 |
| β-1CQ0 | 3 | 3 | 3 | 3 | 3 | 1.5 |
| β-1L2Y | 3 | 3 | 3 | 3 | 3 | 1.5 |
| β-1ROO | 3 | 3 | 3 | 3 | 3 | 1.5 |
| β-1T5Q | 3 | 3 | 3 | 3 | 3 | 1.5 |
| β-1V1D | 3 | 3 | 3 | 3 | 3 | 1.5 |
| 1LFC | 3 | 3 | 3 | 3 | 3 | 1.5 |
| 8TFV | 3 | 3 | 3 | 3 | 3 | 1.5 |
| 1KGM | 3 | 3 | 3 | 3 | 3 | 1.5 |
| 1M4F | 3 | 3 | 3 | 3 | 3 | 1.5 |
| 1TV0 | 3 | 3 | 3 | 3 | 3 | 1.5 |
| α-1LFC | 3 | 3 | 3 | 3 | 3 | 1.5 |
| α-8TFV | 3 | 3 | 3 | 3 | 3 | 1.5 |
| α-1KGM | 3 | 3 | 3 | 3 | 3 | 1.5 |
| α-1M4F | 3 | 3 | 3 | 3 | 3 | 1.5 |
| α-1TV0 | 3 | 3 | 3 | 3 | 3 | 1.5 |

**Table S2.** Summary of highest and lowest average values of simulated properties and the force field used when observing the value. Amber ff99SB-ILDN: SB; Amber ff99SB*-ILDN: SB*; CHARMM22*: C22*; CHARMM36: C36; GROMOS 54A8: 54A8.

| Protein | RMSD | | RMSF | | $R_g$ | |
|---|---|---|---|---|---|---|
| | High | Low | High | Low | High | Low |
| 1CQ0 | 54A8 (0.64) | C22* (0.57) | 54A8 (0.40) | SB* (0.34) | C36 (1.05) | SB/54A8 (0.92) |
| 1L2Y | 54A8 (0.34) | SB* (0.11) | 54A8 (0.25) | SB* (0.07) | 54A8 (0.75) | SB/SB* (0.71) |
| 1ROO | 54A8 (0.26) | C22* (0.19) | 54A8 (0.14) | SB* (0.09) | SB/C22*/C36/ 54A8 (0.92) | SB* (0.90) |
| 1T5Q | SB* (0.93) | C36 (0.33) | C22* (0.52) | C36 (0.25) | C36 (1.36) | SB* (1.01) |
| 1V1D | C22* (0.26) | C36 (0.20) | C22*/SB* (0.16) | C36 (0.10) | C36 (1.03) | SB* (1.02) |
| β-1CQ0 | C22* (0.96) | SB* (0.69) | C36 (0.72) | SB* (0.47) | C36 (1.09) | 54A8 (0.94) |
| β-1L2Y | C22* (0.85) | C36 (0.71) | C36 (0.46) | 54A8 (0.35) | C36 (0.98) | 54A8 (0.82) |
| β-1ROO | C36 (0.92) | 54A8 (0.32) | C36/54A8 (0.59) | SB* (0.43) | C36 (1.20) | 54A8 (0.97) |
| β-1T5Q | C36 (0.73) | C22* (0.49) | C36 (0.53) | SB* (0.30) | C36 (1.16) | 54A8 (1.01) |
| β-1V1D | C36 (0.67) | C22* (0.36) | C36 (0.15) | C22* (0.10) | C36 (1.02) | 54A8 (0.90) |
| 1LFC | C36/54A8 (0.22) | C22* (0.18) | 54A8 (0.15) | C22* (0.11) | SB* (1.11) | C22*/C36 (1.07) |
| 8TFV | 54A8 (0.47) | SB (0.32) | C36 (0.40) | SB (0.28) | SB (1.18) | SB* (1.06) |
| 1KGM | 54A8 (0.33) | C22* (0.23) | SB/54A8 (0.16) | C22* (0.12) | C36 (1.08) | 54A8 (0.93) |
| 1M4F | C36 (0.51) | SB (0.43) | 54A8 (0.24) | C22* (0.34) | C36 (1.17) | SB* (1.09) |
| 1TV0 | C22* (0.31) | SB* (0.25) | C22* (0.16) | SB* (0.13) | SB/ C36 (0.96) | 54A8 (0.94) |
| α-1LFC | C22* (0.76) | C36 (0.60) | C22* (0.47) | SB* (0.33) | C36 (1.07) | SB* (0.93) |
| α-8TFV | SB (0.64) | C22* (0.51) | C36 (0.46) | C22*/SB* (0.38) | C36 (1.00) | 54A8 (0.89) |
| α-1KGM | C36 (1.09) | SB (0.56) | C36 (0.68) | 54A8 (0.29) | C36 (1.43) | 54A8 (0.92) |
| α-1M4F | SB (0.77) | C36 (0.52) | SB (0.55) | 54A8 (0.39) | C36 (1.06) | 54A8 (0.89) |
| α-1TV0 | C22* (0.61) | C36 (0.48) | SB/ C22* (0.44) | C36 (0.32) | C36 (1.10) | 54A8 (1.01) |

**Table S2, continued.**

| Protein | SASA | | HB | | FEL | |
|---|---|---|---|---|---|---|
| | High | Low | Low | High | Low | Low |
| 1CQ0 | C36 (28) | 54A8/SB* (16) | C22* (13) | C36 (0.015) | SB* (0.006) | 54A8 (25) |
| 1L2Y | C22* (20) | SB* (13) | C22*/54A8 (10) | 54A8 (0.007) | SB* (0.001) | SB/SB*/C36/ 54A8 (19) |
| 1ROO | SB/C22*/C36 /54A8 (31) | C22* (21) | C36 (17) | 54A8 (0.003) | SB* (0.001) | SB* (30) |
| 1T5Q | C36 (33) | C36 (22) | SB (15) | C36 (0.022) | SB/54A8 (0.008) | SB*/C22* (30) |
| 1V1D | C36/SB/C22* (32) | 54A8 (20) | C22* (15) | SB*/C22* (0.005) | C36/54A8 (0.002) | SB*/54A8 (31) |
| β-1CQ0 | C36 (31) | 54A8 (11) | C36 (6) | C36 (0.028) | SB* (0.011) | 54A8 (26) |
| β-1L2Y | C36 (25) | 54A8 (6) | C36 (3) | C36 (0.021) | SB* (0.005) | 54A8 (21) |
| β-1ROO | C36 (41) | 54A8 (15) | C36 (8) | C36 (0.029) | SB* (0.007) | 54A8 (33) |
| β-1T5Q | C36 (33) | 54A8 (15) | C36 (8) | C36 (0.037) | SB* (0.008) | 54A8 (30) |
| β-1V1D | C36 (33) | 54A8 (17) | C36 (11) | SB* (0.011) | SB (0.003) | 54A8 (29) |
| 1LFC | 54A8 (30) | SB/SB* (12) | 54A8 (9) | C36/54A8 (0.007) | SB/SB*/ C22* (0.005) | SB/SB*/C22* /C36 (29) |
| 8TFV | C36 (29) | SB/SB* (7) | C36 (3) | C36 (0.051) | SB (0.020) | 54A8 (27) |
| 1KGM | C36 (32) | 54A8 (22) | C36 (15) | SB/ C36 (0.009) | 54A8 (0.002) | 54A8 (29) |
| 1M4F | C36 (29) | 54A8 (8) | C36 (6) | 54A8 (0.021) | C36 (0.012) | 54A8 (27) |
| 1TV0 | C36 (34) | SB* (14) | C22* (9) | C22* (0.004) | SB* (0.003) | 54A8 (32) |
| α-1LFC | C36/C22* (31) | 54A8/SB* (12) | C22* (9) | C22* (0.016) | SB* (0.007) | SB/SB*/54A8 (29) |
| α-8TFV | C36 (27) | SB* (9) | C36 (5) | C36 (0.026) | 54A8 (0.007) | 54A8 (24) |
| α-1KGM | C36 (36) | 54A8 (22) | C36 (12) | C36 (0.022) | 54A8 (0.005) | 54A8 (28) |
| α-1M4F | C36 (28) | 54A8/C36 (12) | SB/C22* (7) | SB* (0.030) | 54A8 (0.003) | 54A8 (24) |
| α-1TV0 | SB/C36 (36) | 54A8 (16) | SB/C22* (12) | SB*/C22* (0.018) | 54A8 (0.006) | 54A8 (33) |

**Table S3**. RMSD plots of native α-proteins, modeled β-proteins, β-proteins, and modeled α-proteins.

<table>
<tr><th colspan="3">RMSD</th></tr>
<tr><th></th><th>1CQ0</th><th>β-1CQ0</th></tr>
<tr><td>Amber 99SB-ILDN</td><td>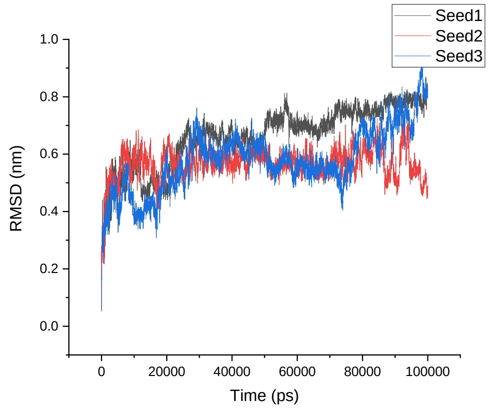
</td><td>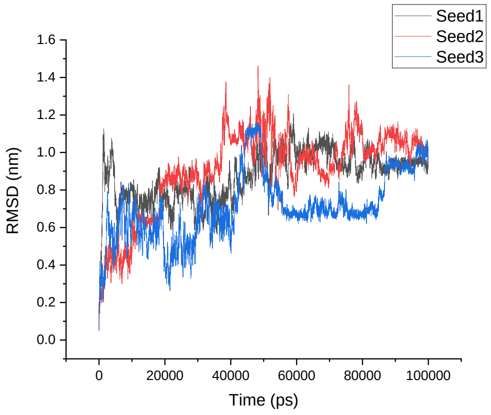
</td></tr>
<tr><td>Amber 99SB*-ILDN</td><td>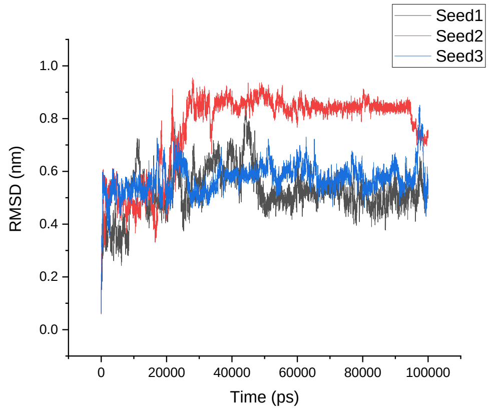
</td><td>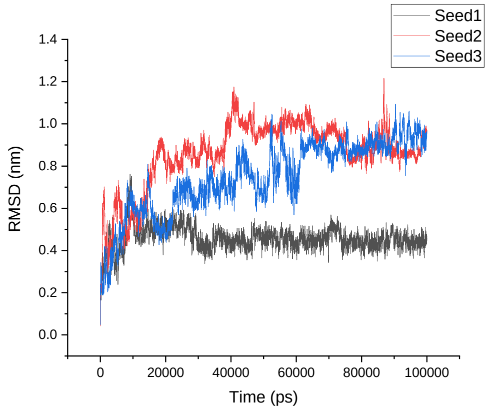
</td></tr>
<tr><td>CHARMM22*</td><td>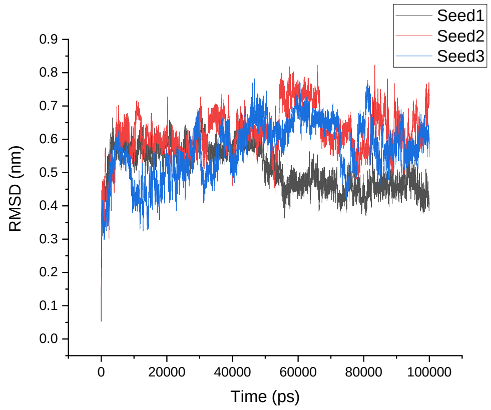
</td><td>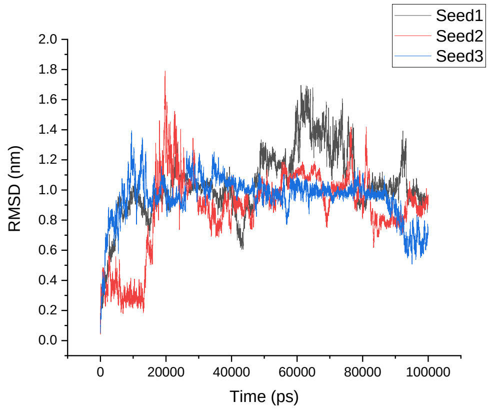
</td></tr>
</table>

<table>
<tr><td>CHARMM36</td><td>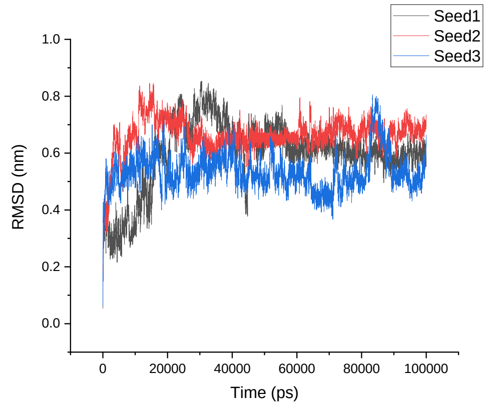
</td><td>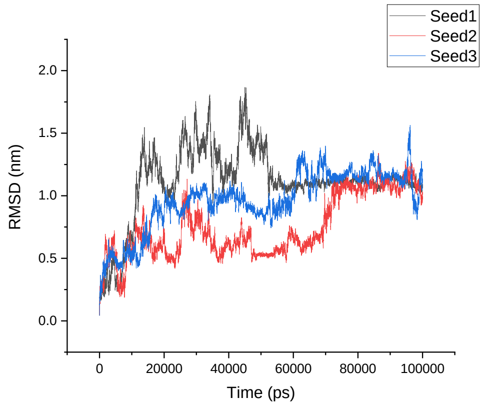
</td></tr>
<tr><td>GROMOS 54A8</td><td>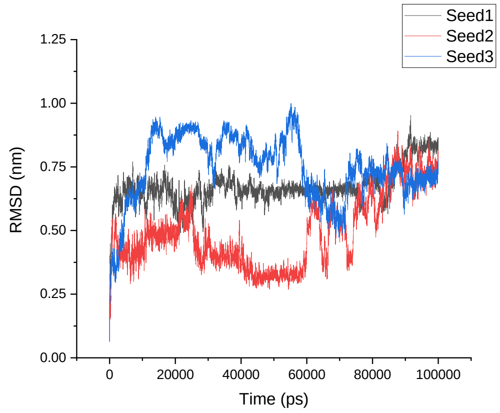
</td><td>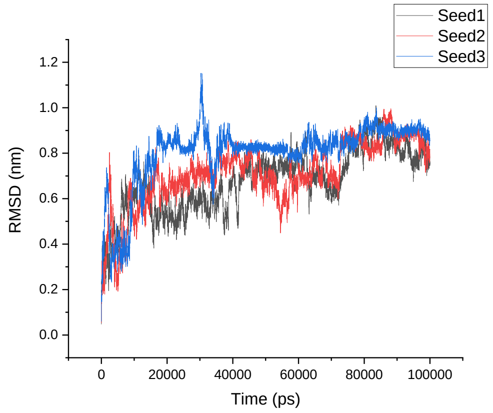
</td></tr>
<tr><td></td><td>1L2Y</td><td>β-1L2Y</td></tr>
<tr><td>Amber 99SB-ILDN</td><td>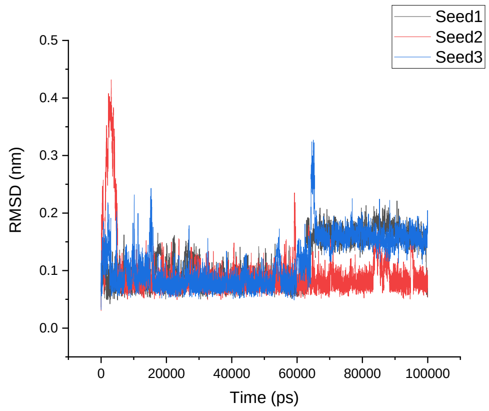
</td><td>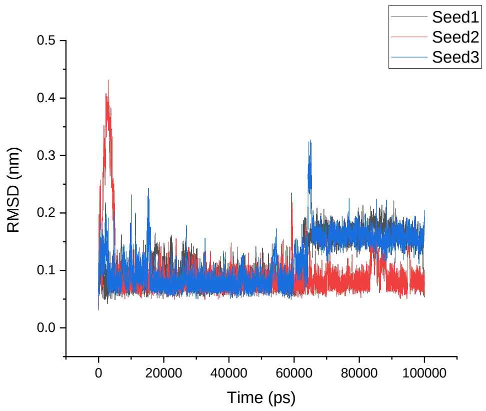
</td></tr>
</table>

| | | |
|---|---|---|
| Amber 99SB*-ILDN | 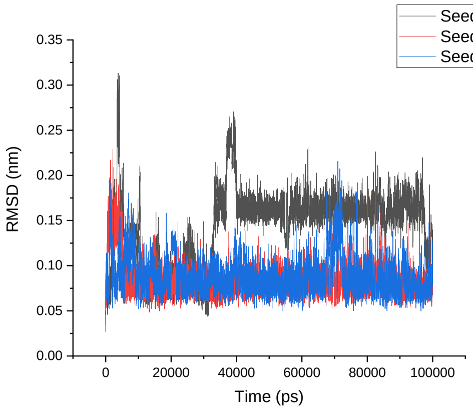 | 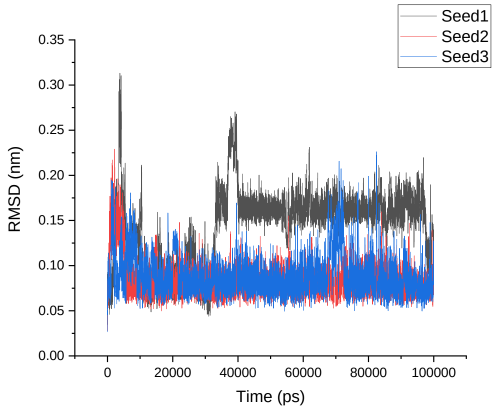 |
| CHARMM22* | 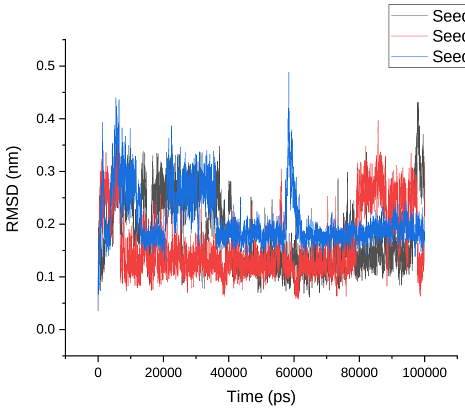 | 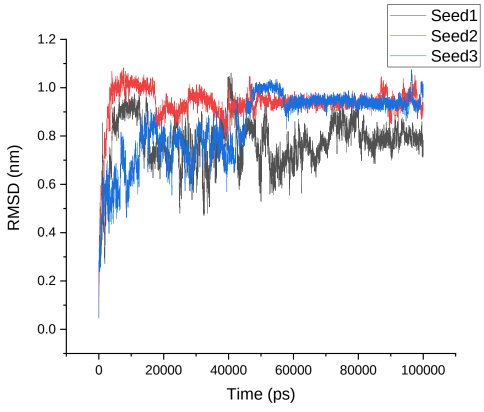 |
| CHARMM36 | 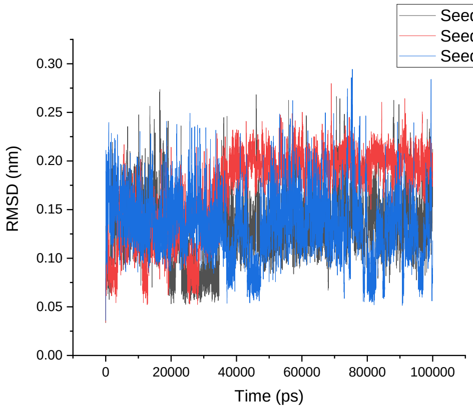 | 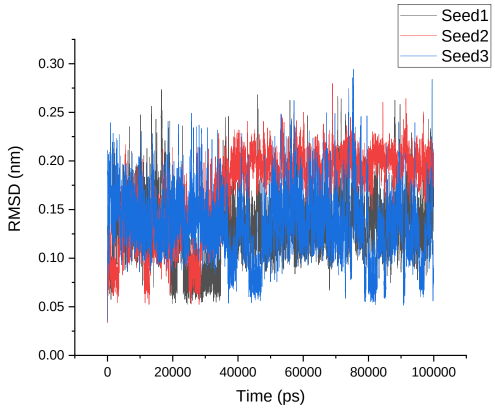 |
| GROMOS 54A8 | 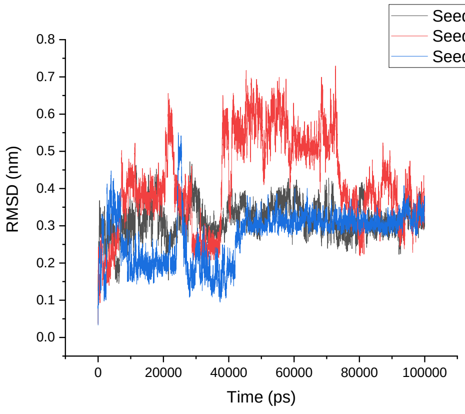 | 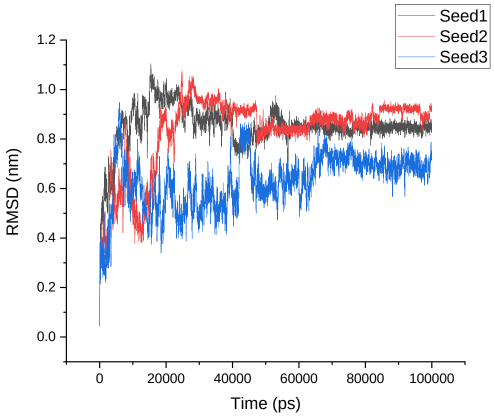 |

| | 1ROO | β-1ROO |
|---|---|---|
| Amber 99SB-ILDN | 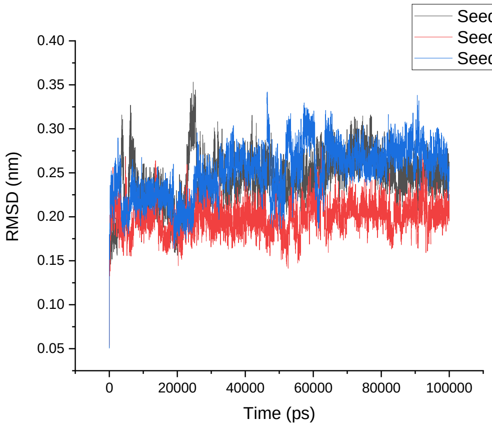 | 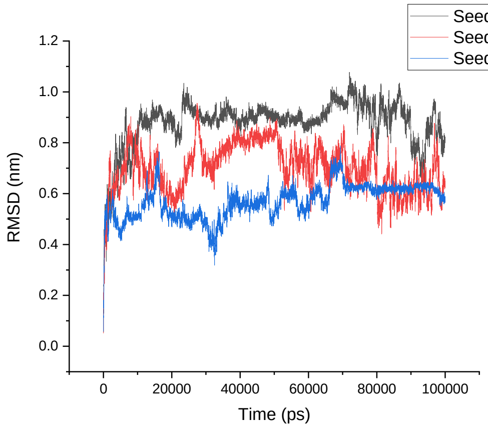 |
| Amber 99SB*-ILDN | 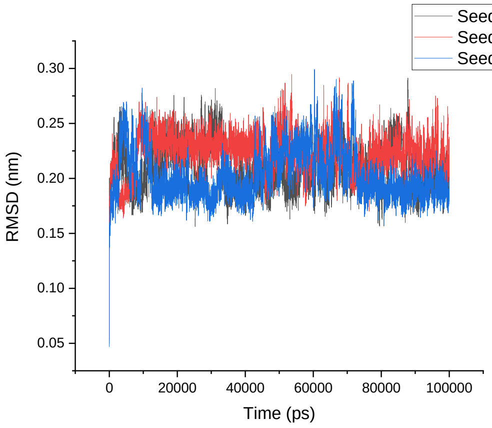 | 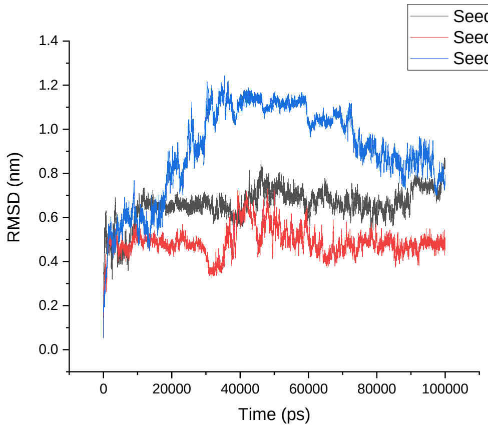 |
| CHARMM22* | 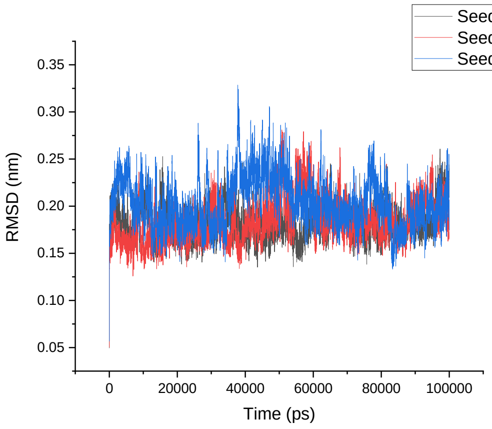 | 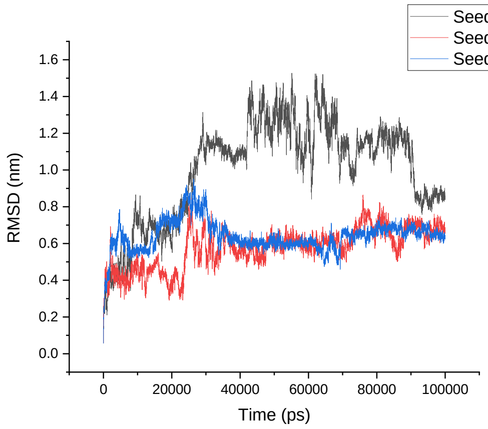 |

| | | |
|---|---|---|
| CHARMM36 | 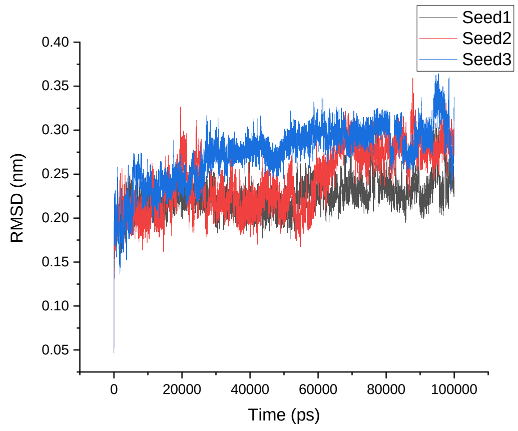 | 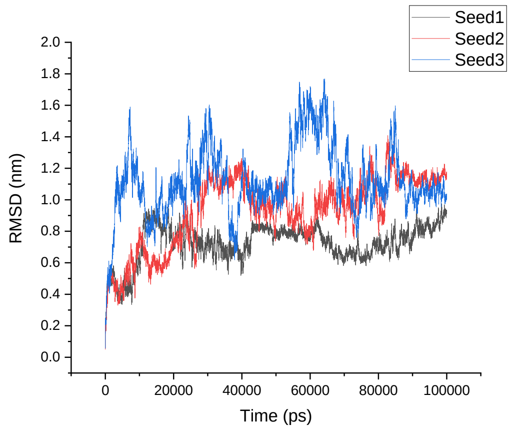 |
| GROMOS 54A8 | 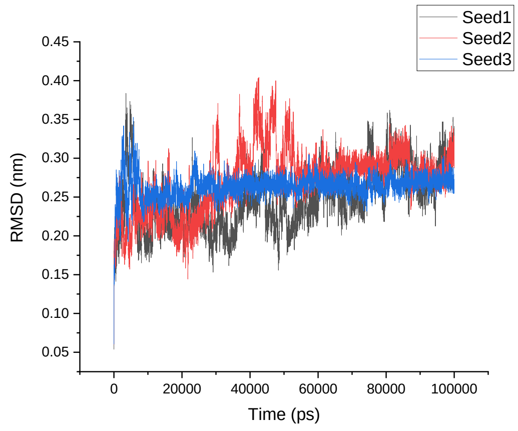 | 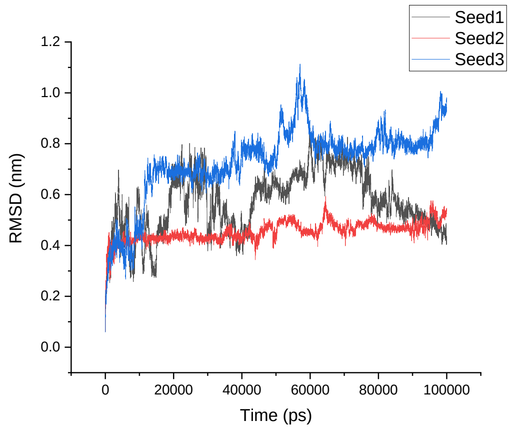 |
| | **1T5Q** | **β-1T5Q** |
| Amber 99SB-ILDN | 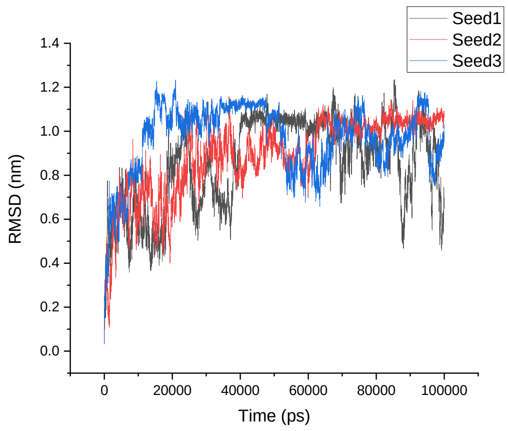 | 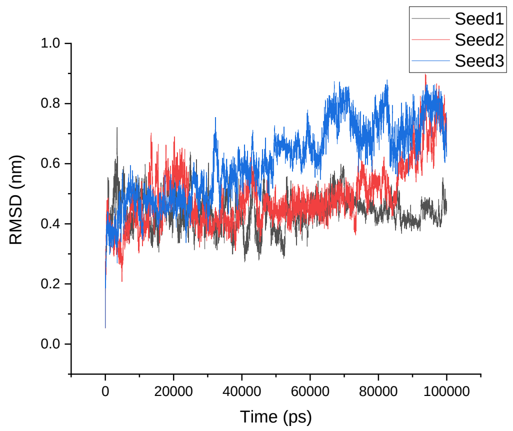 |

| | | |
|---|---|---|
| Amber 99SB*-ILDN | 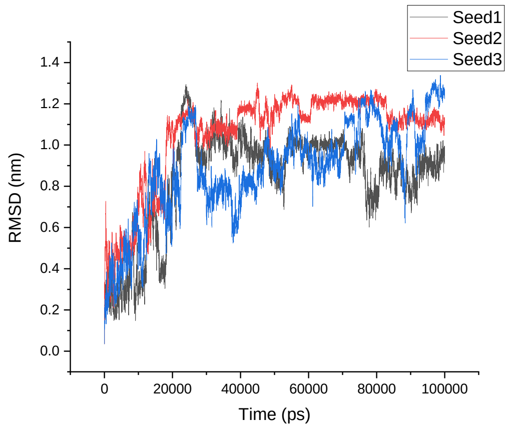  | 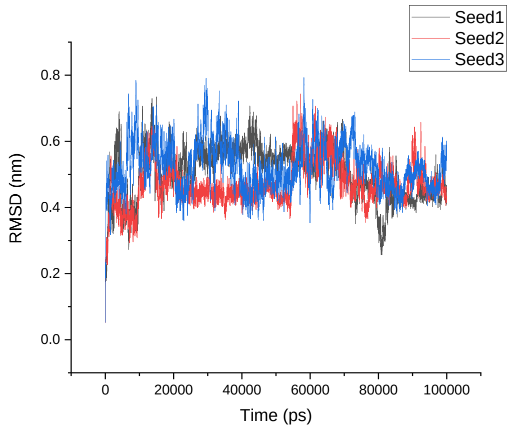  |
| CHARMM22* | 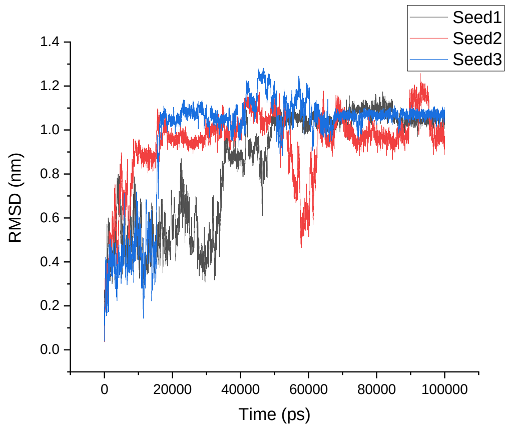  | 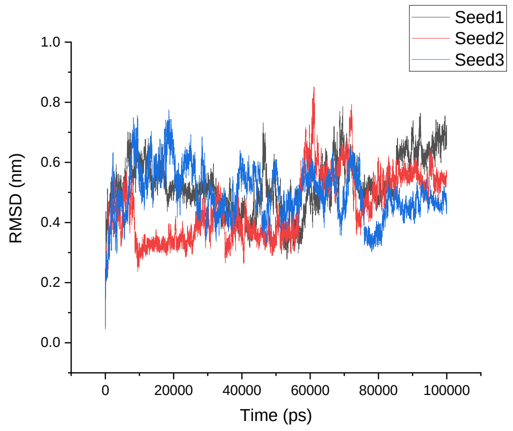  |
| CHARMM36 | 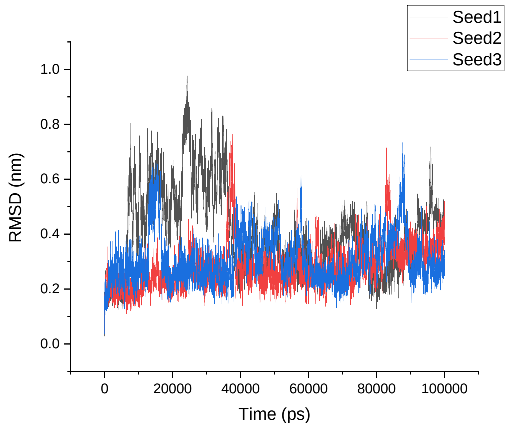  | 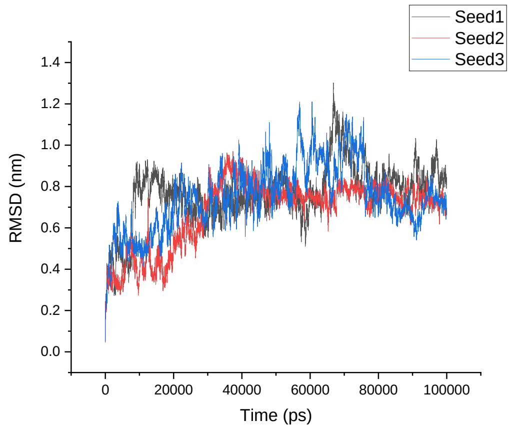  |

<table>
<tr><td>GROMOS 54A8</td><td>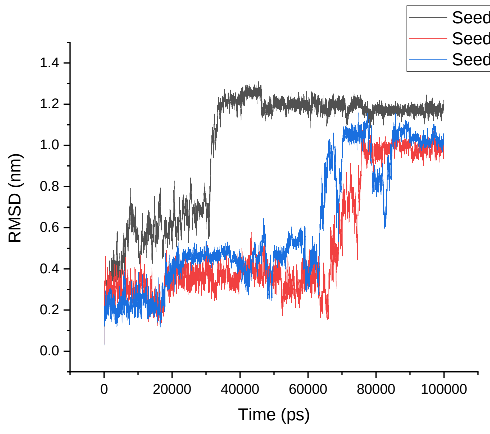
</td><td>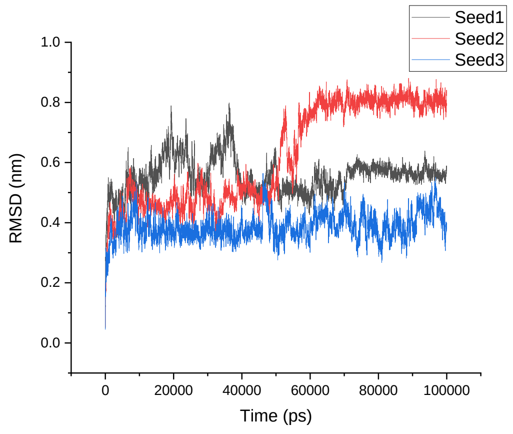
</td></tr>
<tr><td></td><td>1V1D</td><td>β-1V1D</td></tr>
<tr><td>Amber 99SB-ILDN</td><td>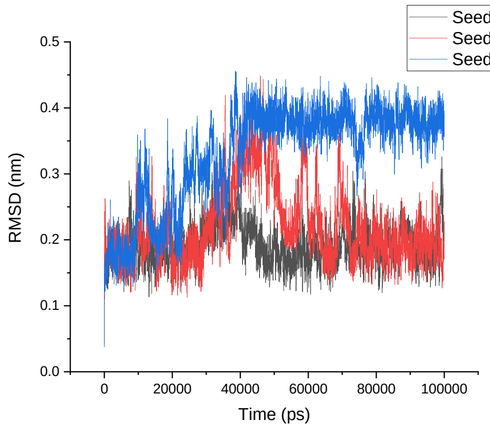
</td><td>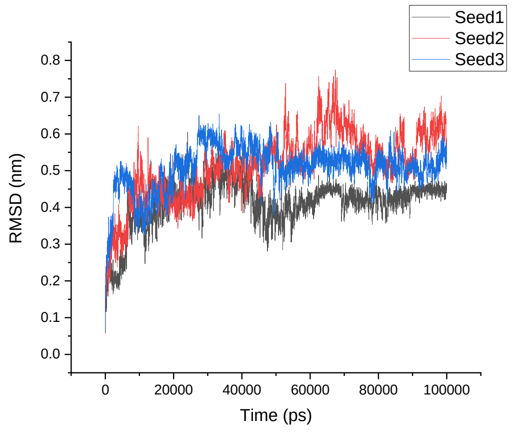
</td></tr>
<tr><td>Amber 99SB*-ILDN</td><td>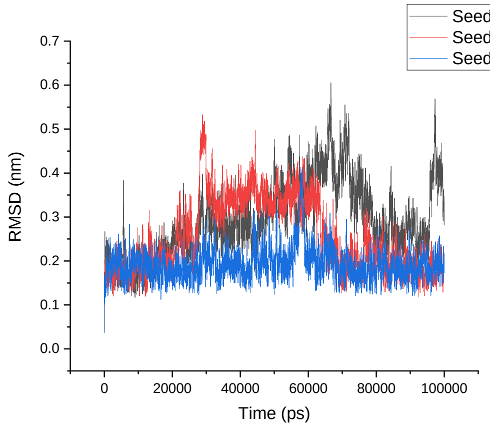
</td><td>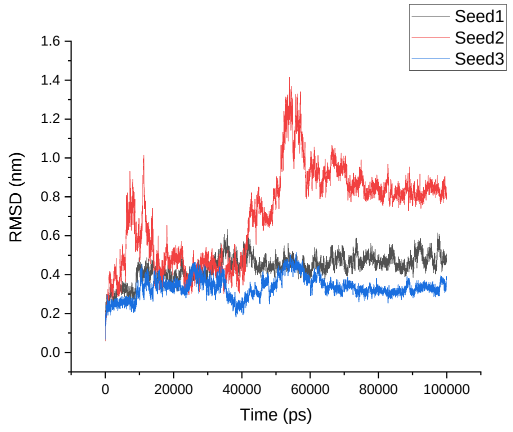
</td></tr>
</table>

<table>
<tr><td>CHARMM22*</td><td>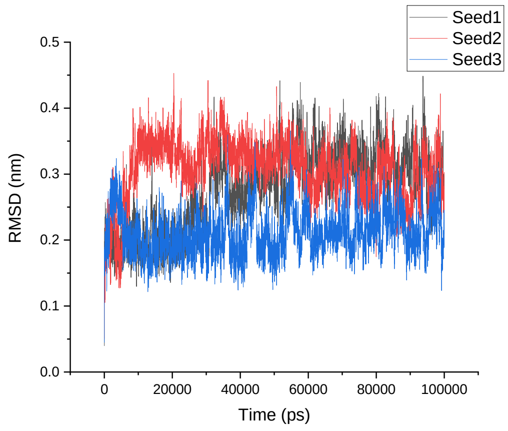
</td><td>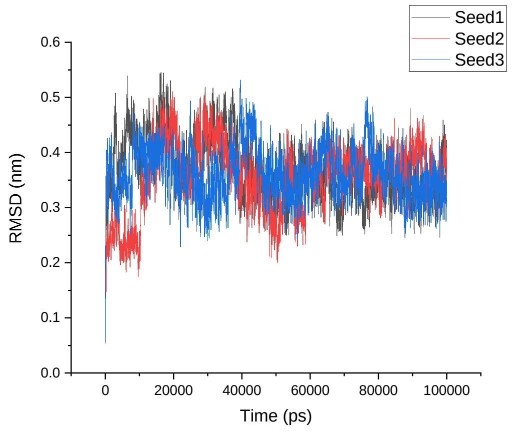
</td></tr>
<tr><td>CHARMM36</td><td>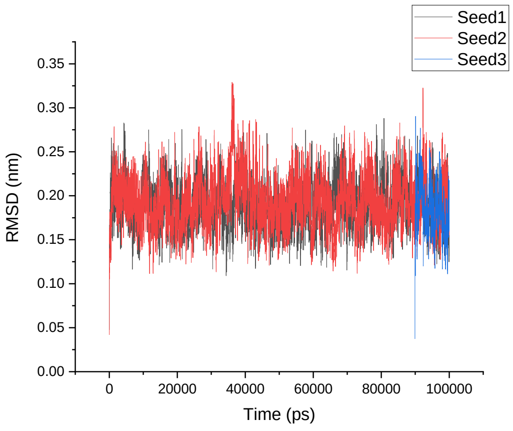
</td><td>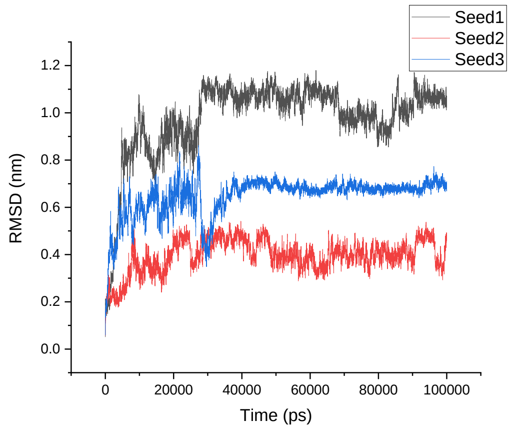
</td></tr>
<tr><td>GROMOS 54A8</td><td>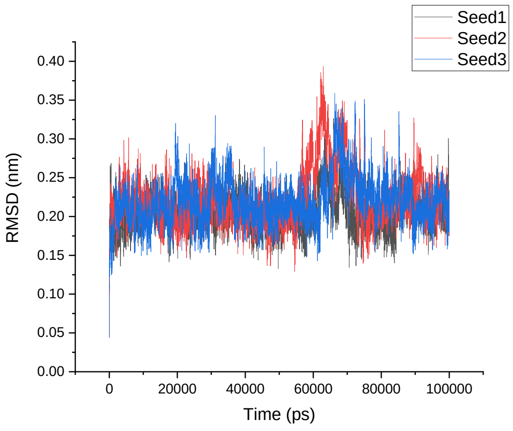
</td><td>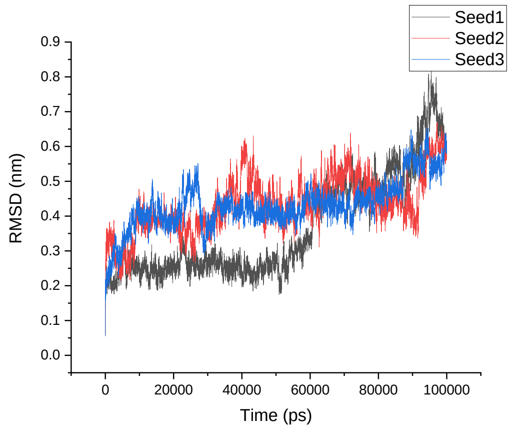
</td></tr>
<tr><td></td><td>1LFC</td><td>α-1LFC</td></tr>
</table>

Amber 99SB-ILDN

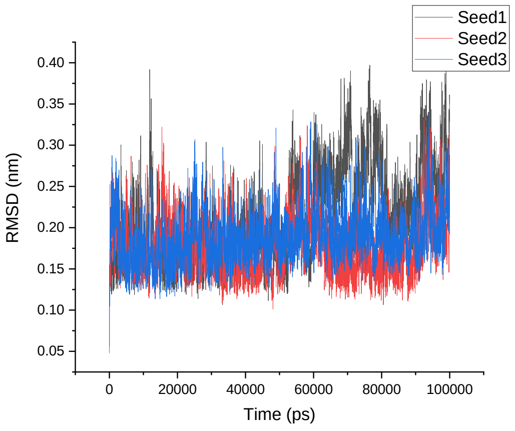


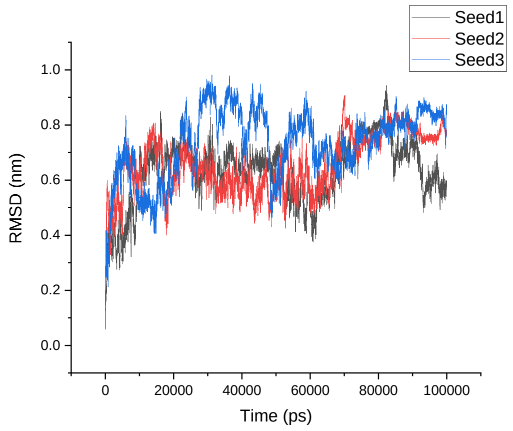


Amber 99SB*-ILDN

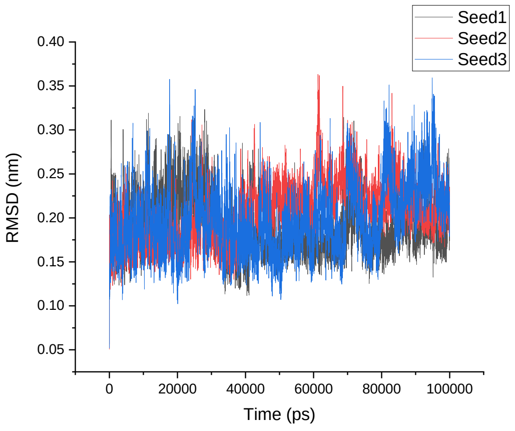


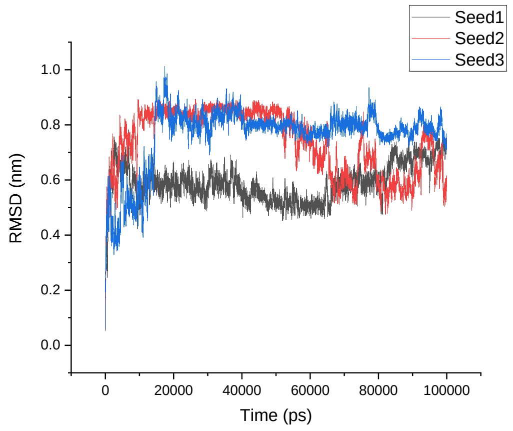


CHARMM22*

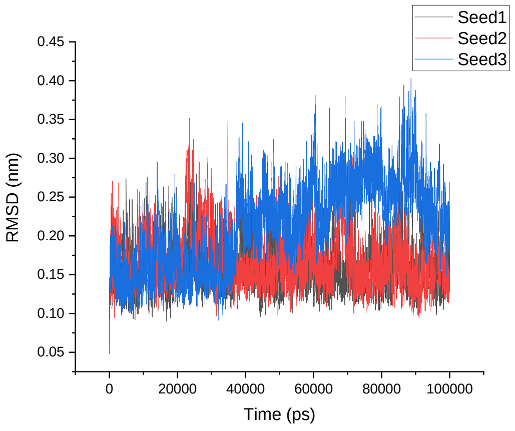


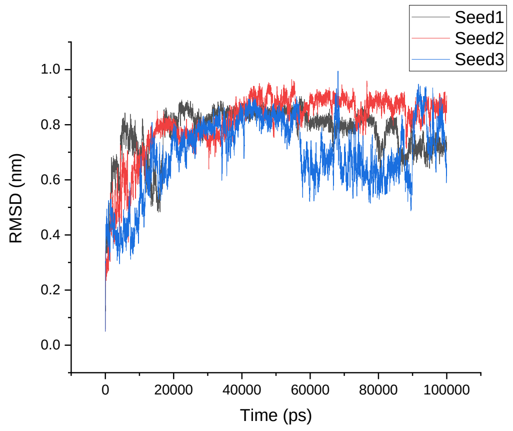

| | | |
|---|---|---|
| CHARMM36 | 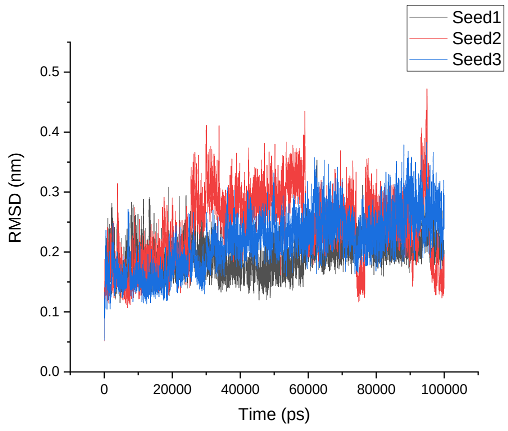  | 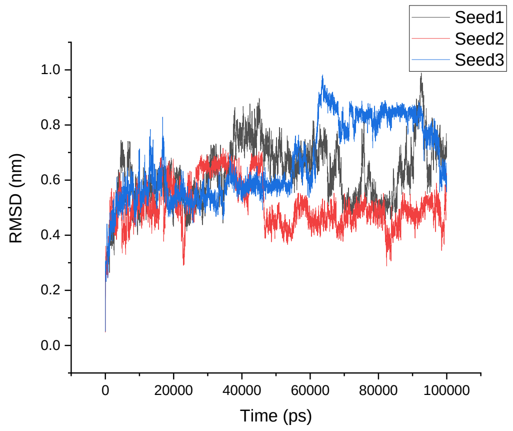  |
| GROMOS 54A8 | 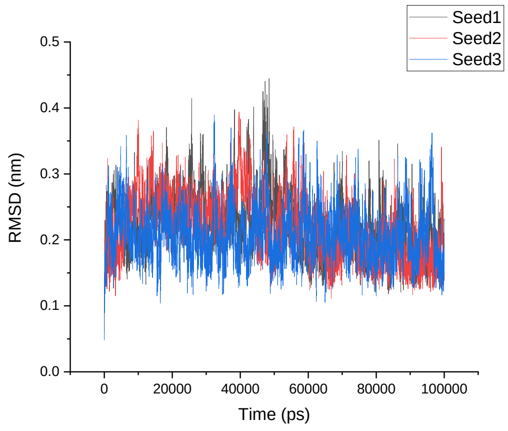  | 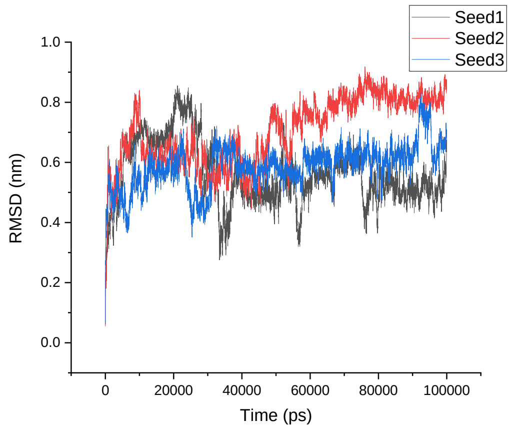  |
| | **8TFV** | **α-8TFV** |
| Amber 99SB-ILDN | 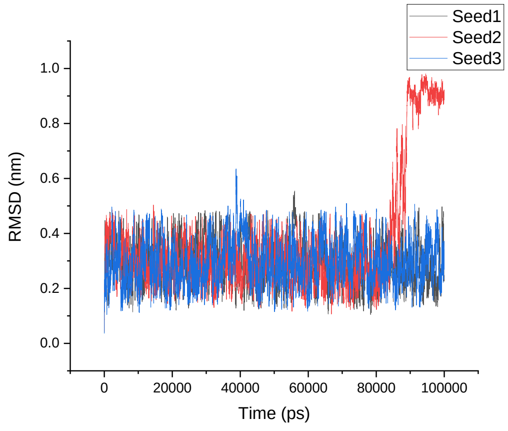  | 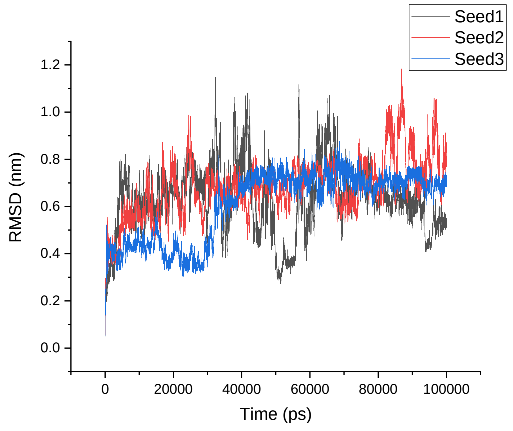  |

| | | |
|---|---|---|
| Amber 99SB*-ILDN | 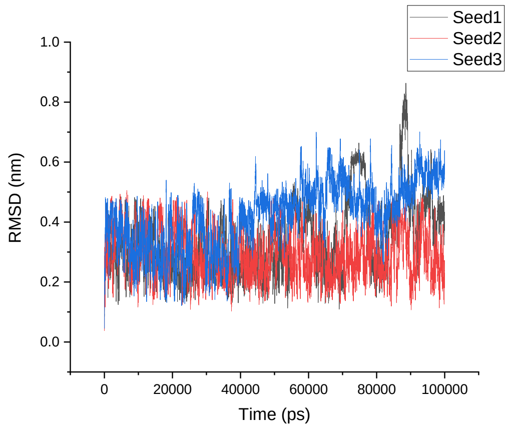 | 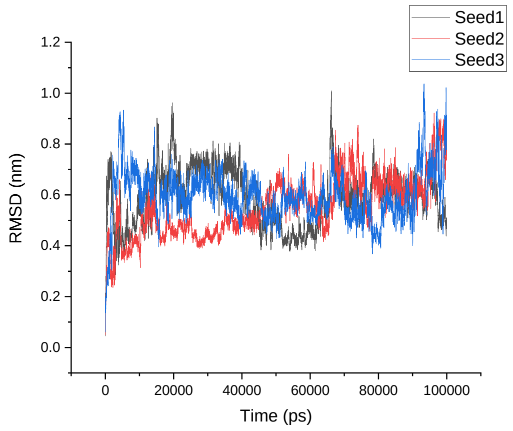 |
| CHARMM22* | 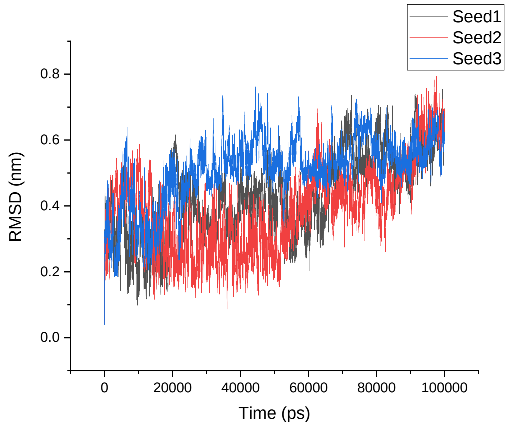 | 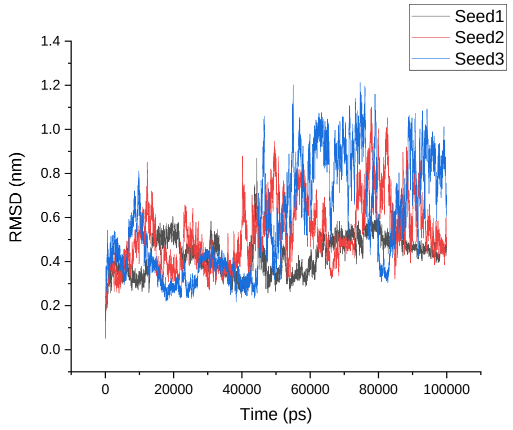 |
| CHARMM36 | 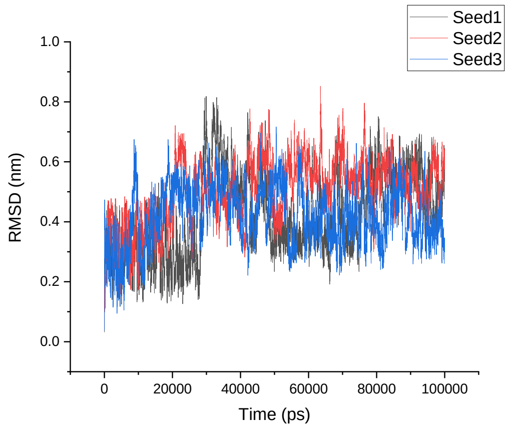 | 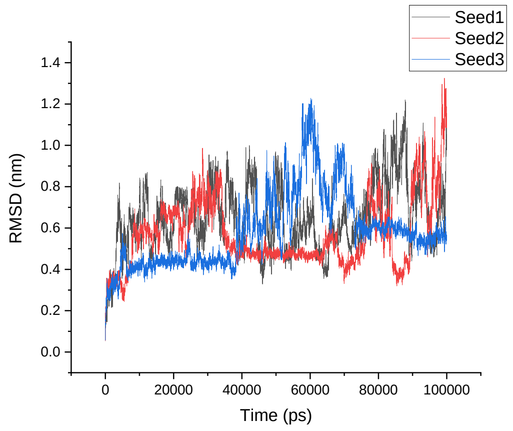 |

| | | |
|---|---|---|
| GROMOS 54A8 | 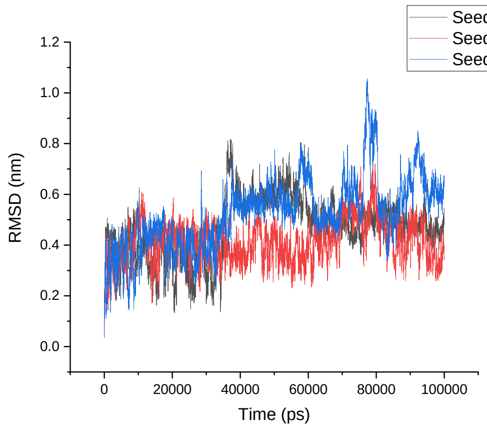 | 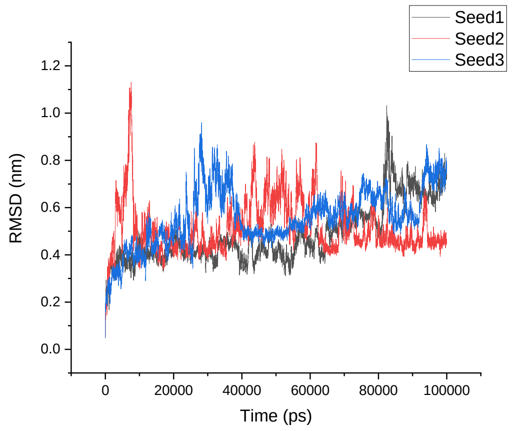 |
| | **1KGM** | **α-1KGM** |
| Amber 99SB-ILDN | 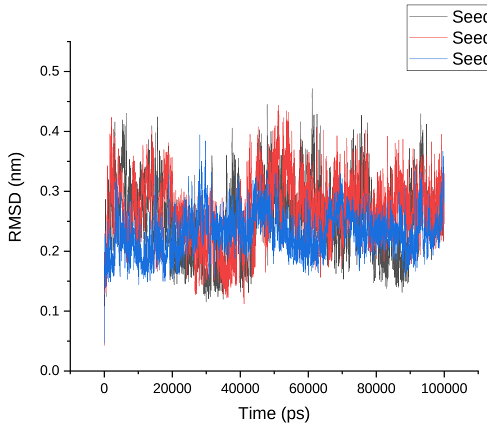 | 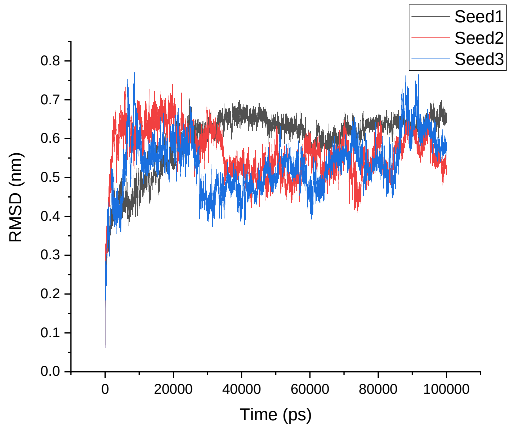 |
| Amber 99SB*-ILDN | 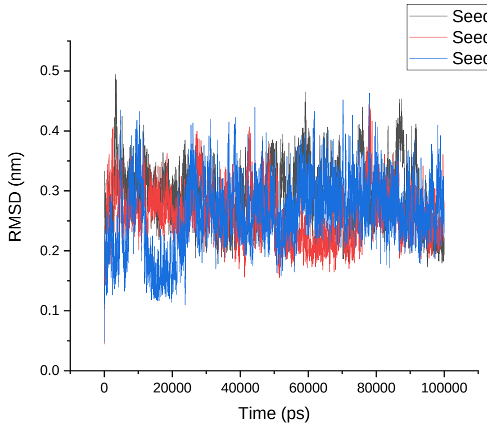 | 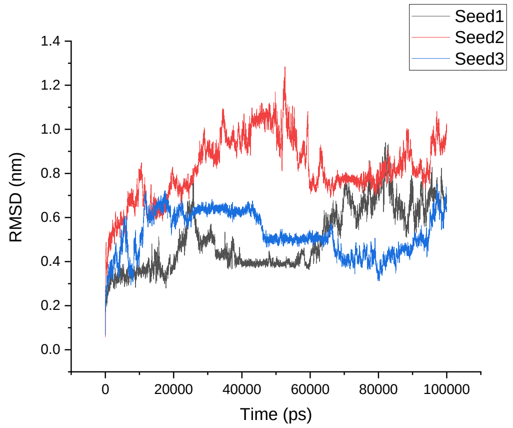 |

<table>
<tr><td>CHARMM22*</td><td>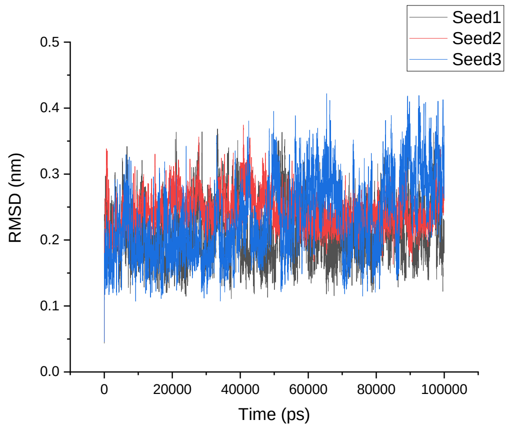
</td><td>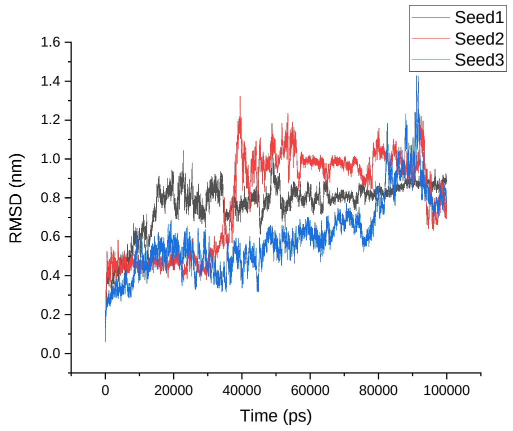
</td></tr>
<tr><td>CHARMM36</td><td>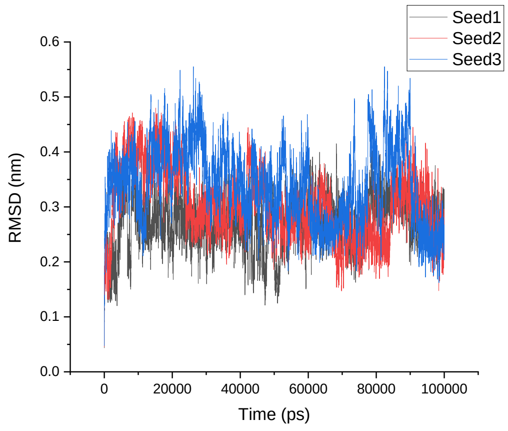
</td><td>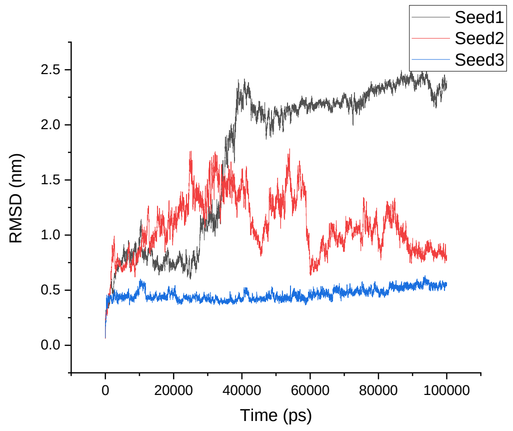
</td></tr>
<tr><td>GROMOS<br>54A8</td><td>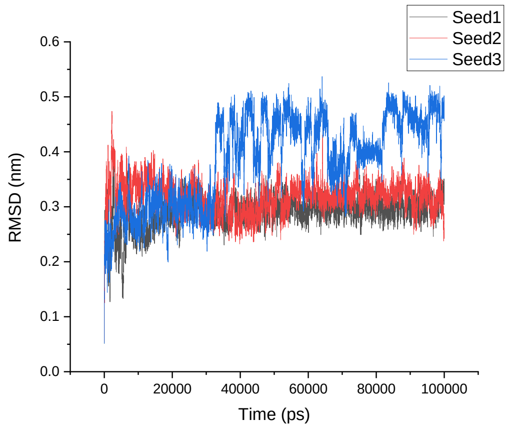
</td><td>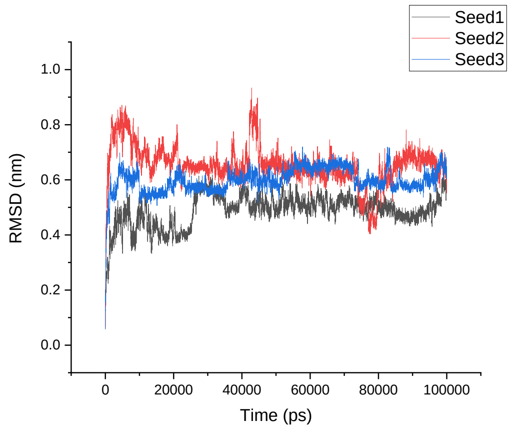
</td></tr>
<tr><td></td><td>1M4F</td><td>α-1M4F</td></tr>
</table>

Amber 99SB-ILDN

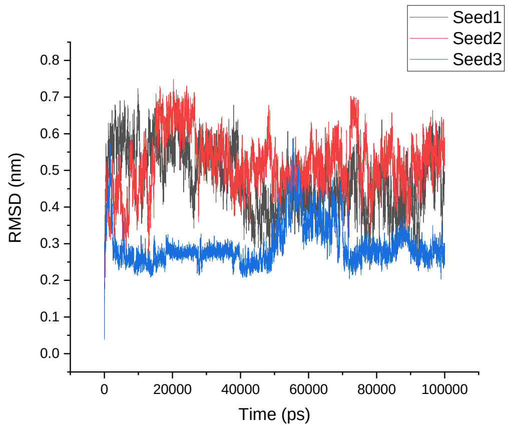


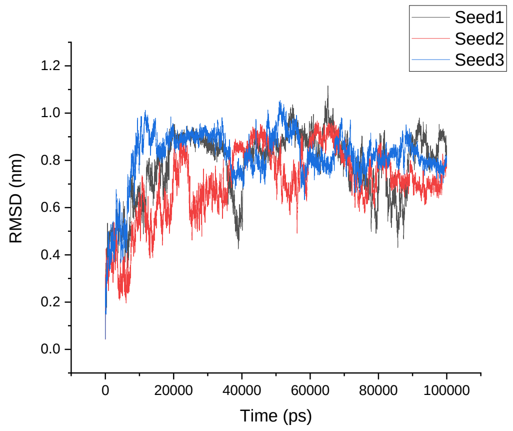


Amber 99SB*-ILDN

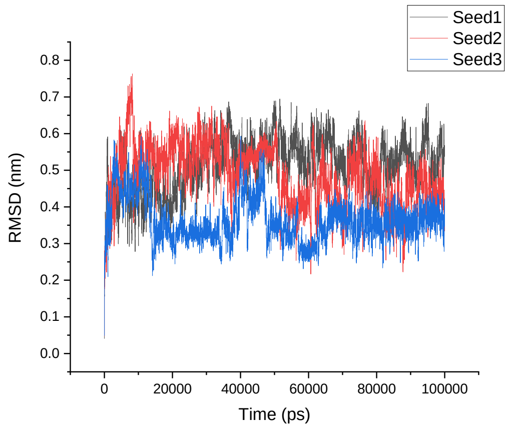


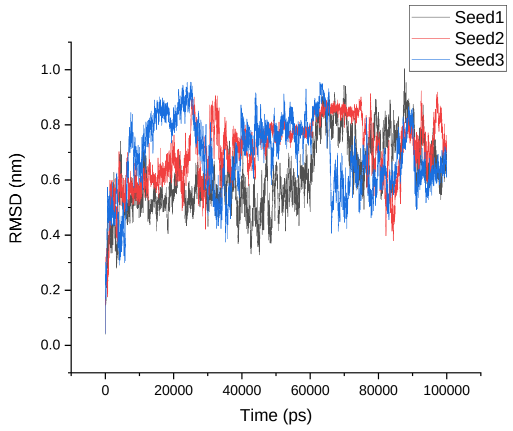


CHARMM22*

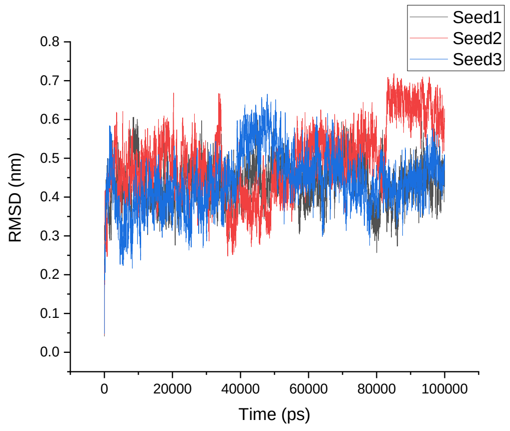


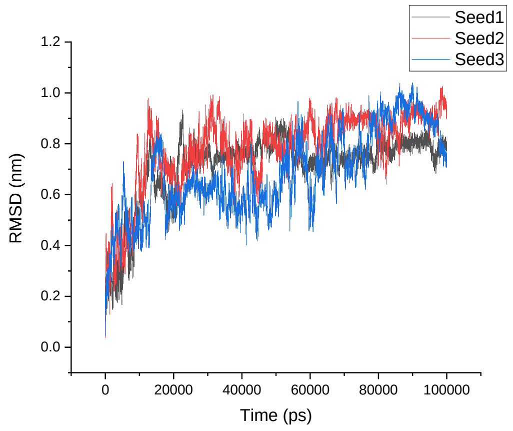

<table>
<tr><td>CHARMM36</td><td>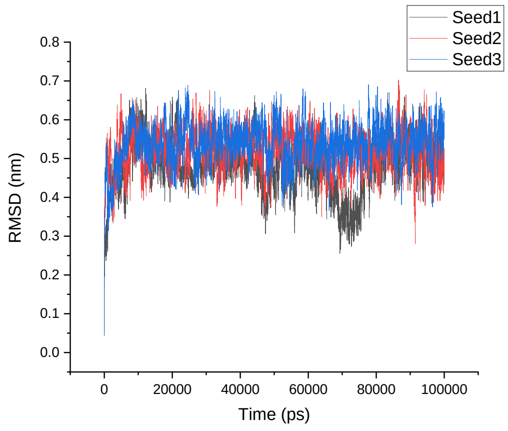
</td><td>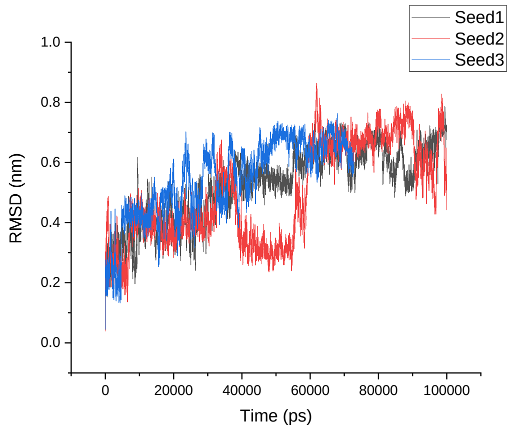
</td></tr>
<tr><td>GROMOS 54A8</td><td>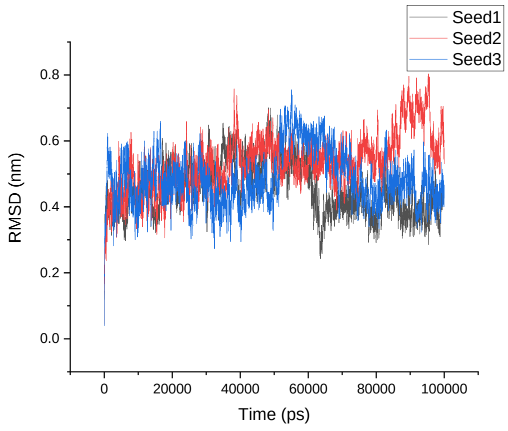
</td><td>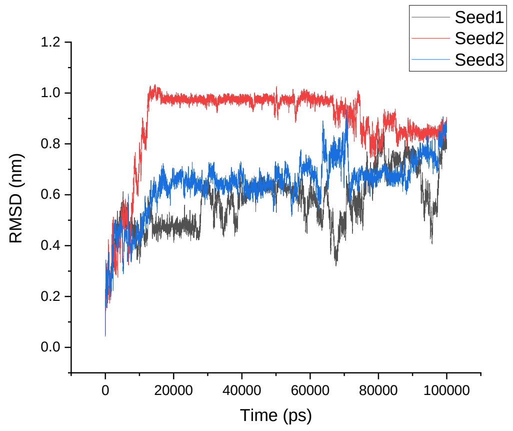
</td></tr>
<tr><td></td><td>1TV0</td><td>α-1TV0</td></tr>
<tr><td>Amber 99SB-ILDN</td><td>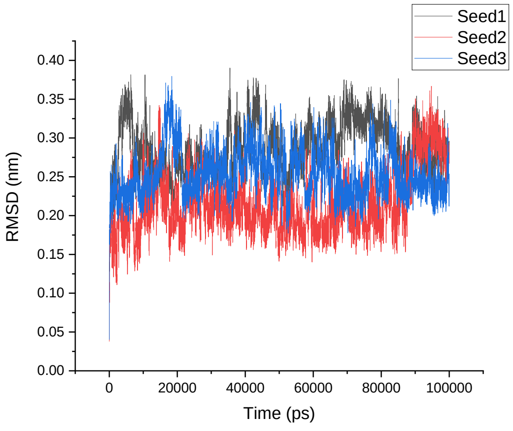
</td><td>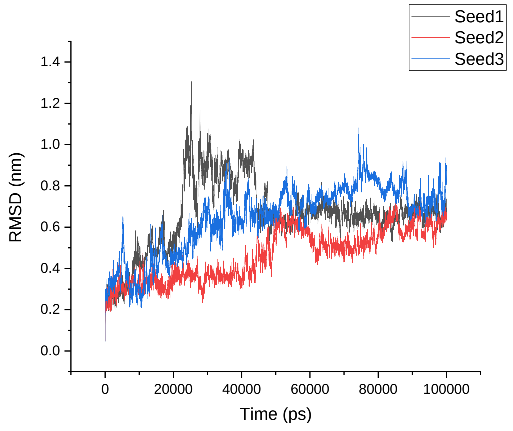
</td></tr>
</table>

Amber 99SB*-ILDN

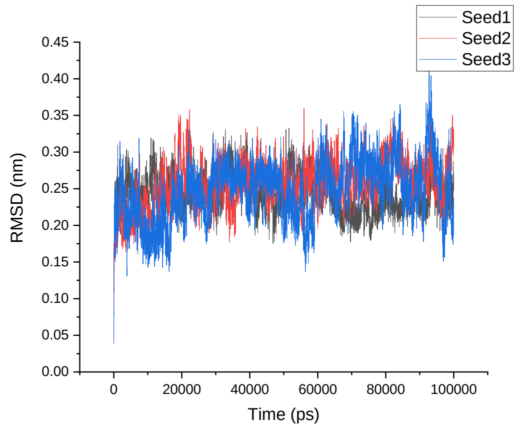


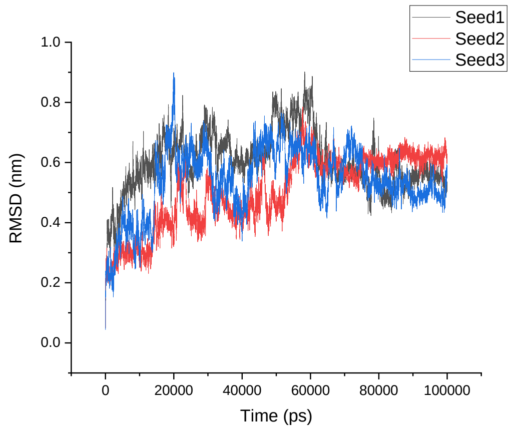


CHARMM22*

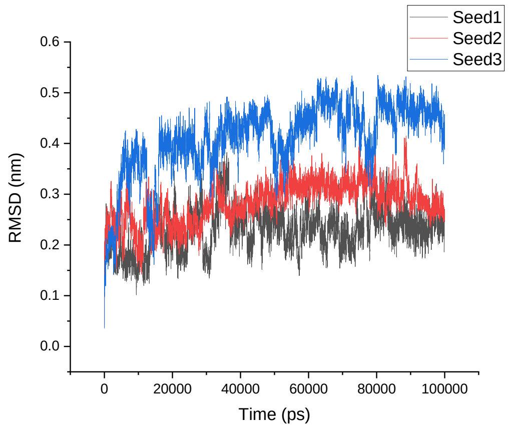


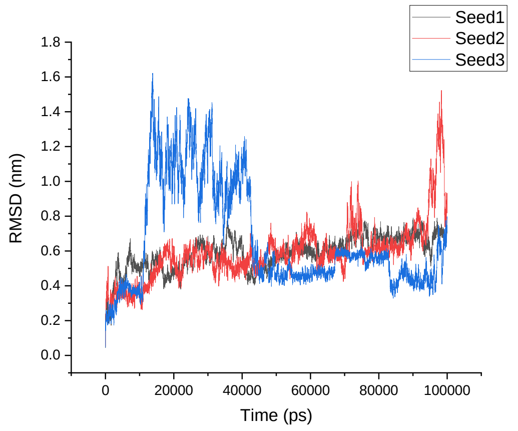


CHARMM36

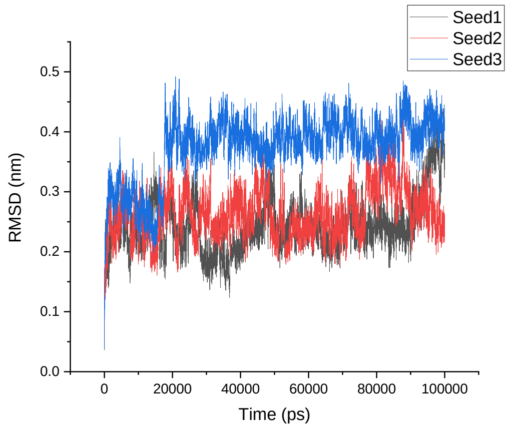


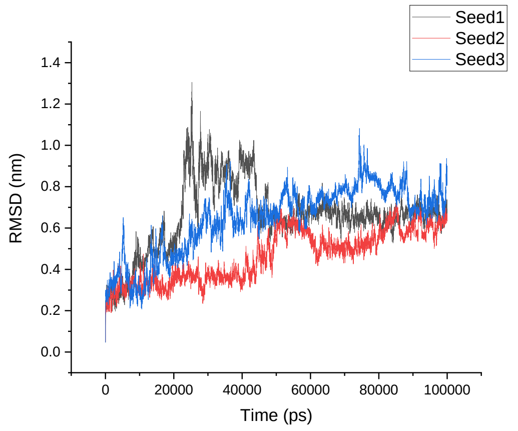

| | | |
|---|---|---|
| GROMOS 54A8 | 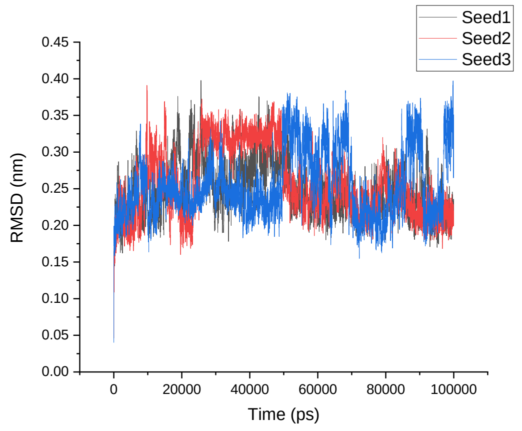 Seed1 Seed2 Seed3 RMSD (nm) Time (ps) | 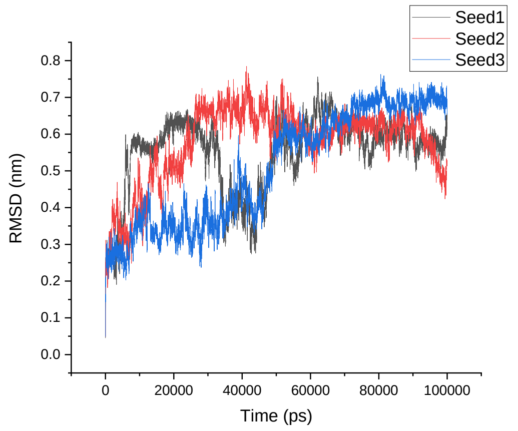 Seed1 Seed2 Seed3 RMSD (nm) Time (ps) |

**Table S4**. Average RMSD of the native α-proteins, modeled β-proteins, β-proteins, and modeled α-proteins.

| | RMSD | SD | RMSD | SD | RMSD | SD | RMSD | SD | RMSD | SD |
|---|---|---|---|---|---|---|---|---|---|---|
| α- proteins | | | | | | | | | | |
| | **1CQ0** | | **1L2Y** | | **1ROO** | | **1T5Q** | | **1V1D** | |
| Amber 99SB-ILDN | 0.60 | 0.09 | 0.11 | 0.04 | 0.23 | 0.02 | 0.90 | 0.18 | 0.25 | 0.05 |
| Amber 99SB*-ILDN | 0.62 | 0.10 | 0.11 | 0.03 | 0.21 | 0.02 | 0.93 | 0.24 | 0.25 | 0.07 |
| CHARMM22* | 0.57 | 0.08 | 0.18 | 0.06 | 0.19 | 0.02 | 0.92 | 0.22 | 0.26 | 0.05 |
| CHARMM36 | 0.60 | 0.08 | 0.15 | 0.04 | 0.25 | 0.03 | 0.33 | 0.11 | 0.20 | 0.03 |
| GROMOS 54A8 | 0.64 | 0.11 | 0.34 | 0.08 | 0.26 | 0.03 | 0.70 | 0.30 | 0.22 | 0.03 |
| Modelled β-proteins | | | | | | | | | | |
| | **β-1CQ0** | | **β-1L2Y** | | **β-1ROO** | | **β-1T5Q** | | **β-1V1D** | |
| Amber 99SB-ILDN | 0.83 | 0.18 | 0.73 | 0.12 | 0.72 | 0.09 | 0.51 | 0.10 | 0.48 | 0.07 |
| Amber 99SB*-ILDN | 0.69 | 0.13 | 0.81 | 0.13 | 0.68 | 0.11 | 0.50 | 0.07 | 0.49 | 0.12 |
| CHARMM22* | 0.96 | 0.21 | 0.85 | 0.11 | 0.74 | 0.15 | 0.49 | 0.09 | 0.36 | 0.05 |
| CHARMM36 | 0.94 | 0.26 | 0.71 | 0.12 | 0.92 | 0.19 | 0.73 | 0.15 | 0.67 | 0.11 |
| GROMOS 54A8 | 0.73 | 0.14 | 0.77 | 0.11 | 0.32 | 0.12 | 0.52 | 0.09 | 0.41 | 0.10 |
| β- proteins | | | | | | | | | | |
| | **1LFC** | | **8TFV** | | **1KGM** | | **1M4F** | | **1TV0** | |
| Amber 99SB-ILDN | 0.19 | 0.19 | 0.32 | 0.12 | 0.25 | 0.05 | 0.43 | 0.07 | 0.25 | 0.03 |
| Amber 99SB*-ILDN | 0.20 | 0.20 | 0.35 | 0.10 | 0.27 | 0.05 | 0.45 | 0.07 | 0.25 | 0.03 |
| CHARMM22* | 0.18 | 0.18 | 0.44 | 0.12 | 0.23 | 0.04 | 0.46 | 0.07 | 0.31 | 0.05 |
| CHARMM36 | 0.22 | 0.22 | 0.45 | 0.12 | 0.30 | 0.06 | 0.51 | 0.06 | 0.29 | 0.05 |
| GROMOS 54A8 | 0.22 | 0.22 | 0.47 | 0.11 | 0.33 | 0.05 | 0.49 | 0.08 | 0.25 | 0.04 |
| Modelled α- proteins | | | | | | | | | | |
| | **α-1LFC** | | **α-8TFV** | | **α-1KGM** | | **α-1M4F** | | **α-1TV0** | |
| Amber 99SB-ILDN | 0.67 | 0.12 | 0.64 | 0.14 | 0.56 | 0.07 | 0.77 | 0.14 | 0.59 | 0.15 |
| Amber 99SB*-ILDN | 0.70 | 0.10 | 0.58 | 0.11 | 0.62 | 0.13 | 0.67 | 0.13 | 0.55 | 0.11 |
| CHARMM22* | 0.76 | 0.11 | 0.51 | 0.16 | 0.72 | 0.19 | 0.73 | 0.15 | 0.61 | 0.19 |
| CHARMM36 | 0.60 | 0.11 | 0.60 | 0.17 | 1.09 | 0.33 | 0.52 | 0.14 | 0.48 | 0.15 |
| GROMOS 54A8 | 0.62 | 0.10 | 0.52 | 0.12 | 0.58 | 0.06 | 0.70 | 0.13 | 0.54 | 0.12 |

**Table S5**. RMSF plots of the native α-proteins, modeled β-proteins, β-proteins, and modeled α-proteins.

| RMSF | | |
|---|---|---|
| | **1CQ0** | **β-1CQ0** |
| Amber 99SB-ILDN | 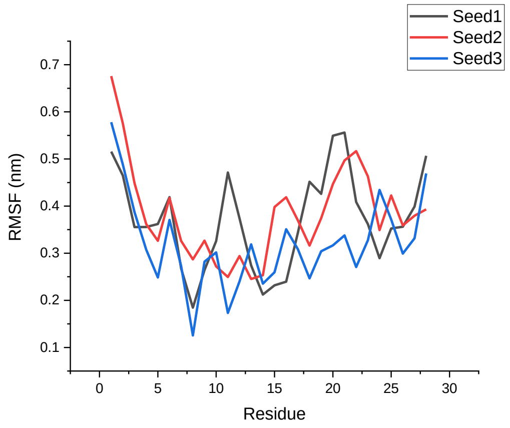 | 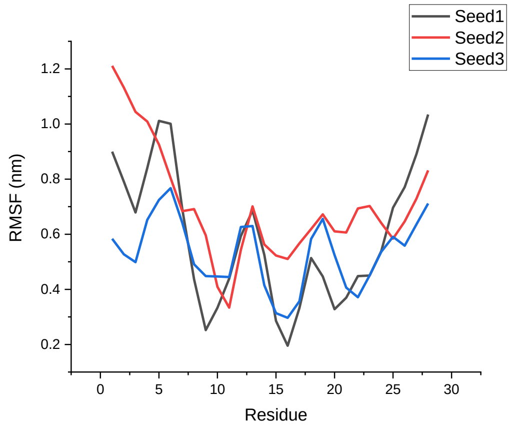 |
| Amber 99SB*-ILDN | 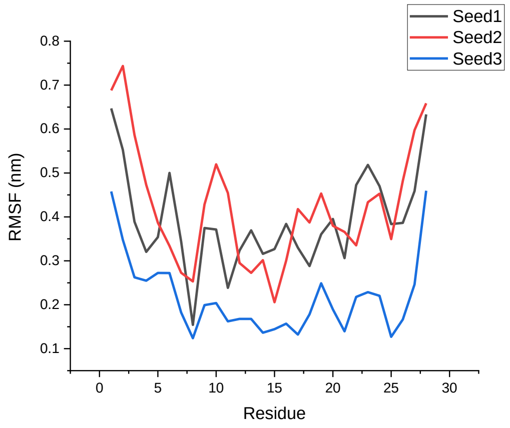 | 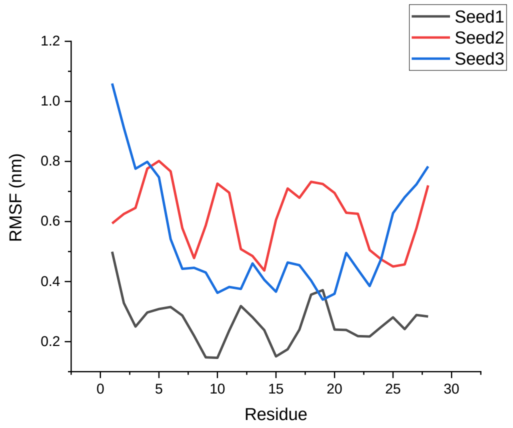 |
| CHARMM22* | 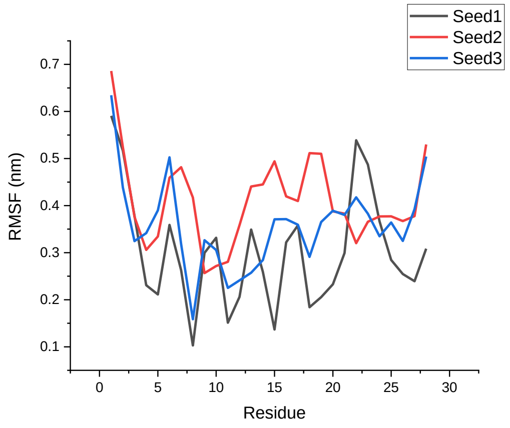 | 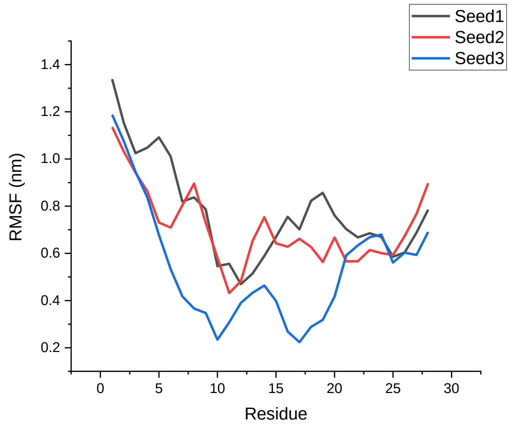 |

| | | |
|---|---|---|
| CHARMM36 | 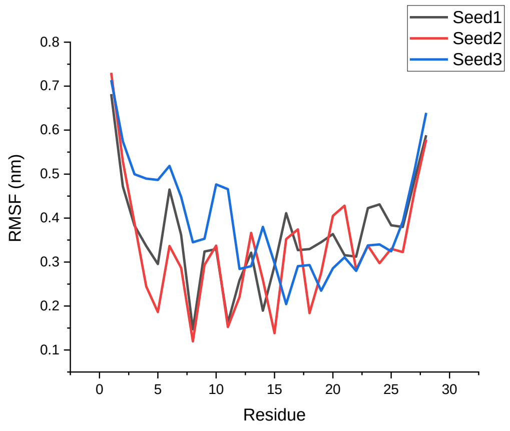 | 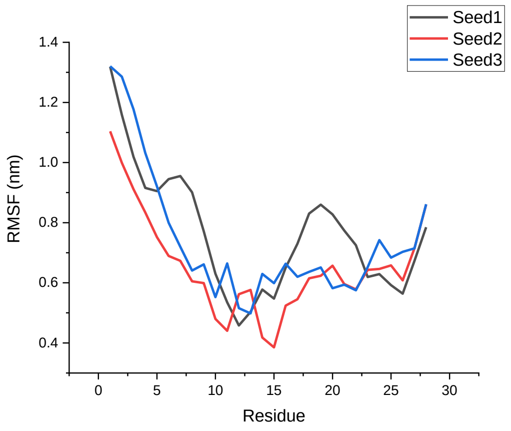 |
| GROMOS 54A8 | 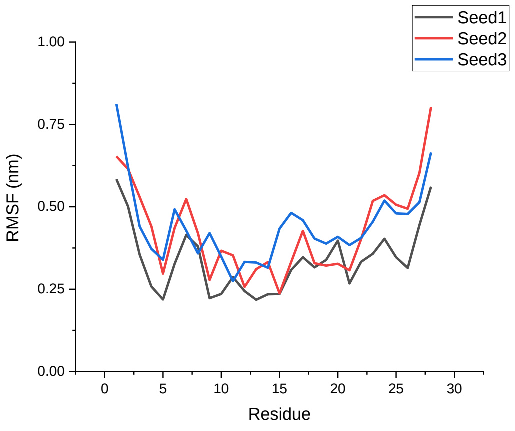 | 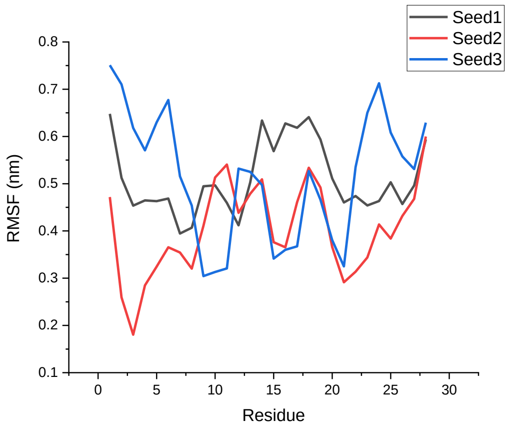 |
| | **1L2Y** | **β-1L2Y** |
| Amber 99SB-ILDN | 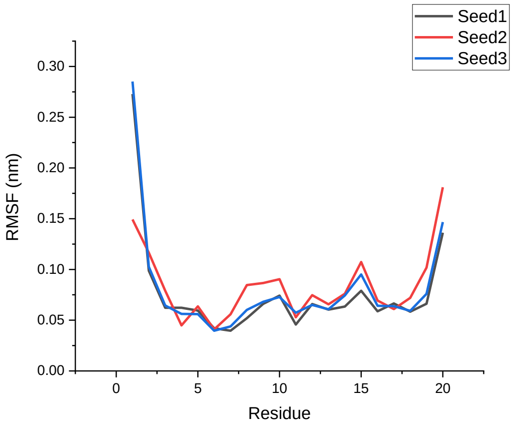 | 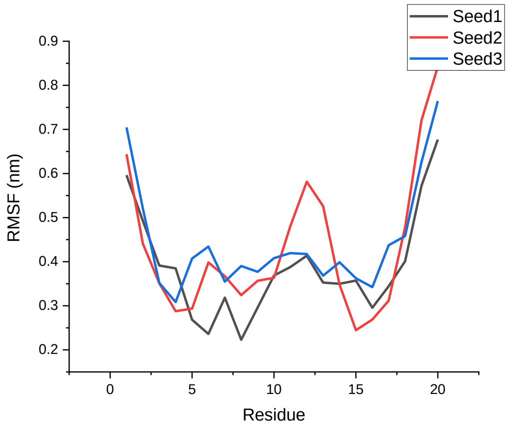 |

<table>
<tr><td>Amber 99SB*-ILDN</td><td>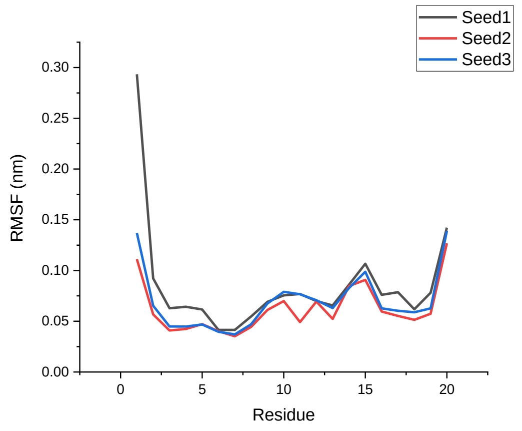
</td><td>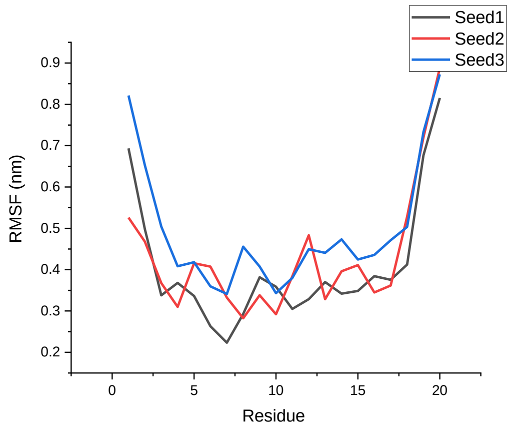
</td></tr>
<tr><td>CHARMM22*</td><td>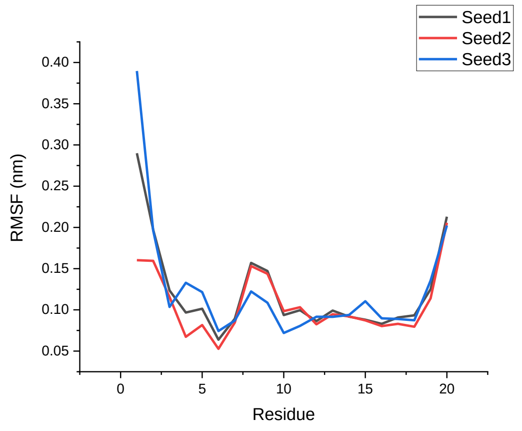
</td><td>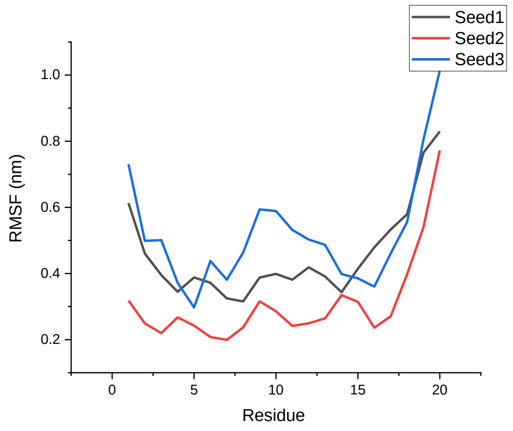
</td></tr>
<tr><td>CHARMM36</td><td>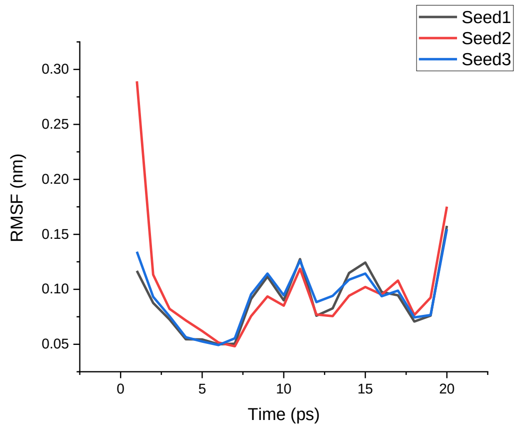
</td><td>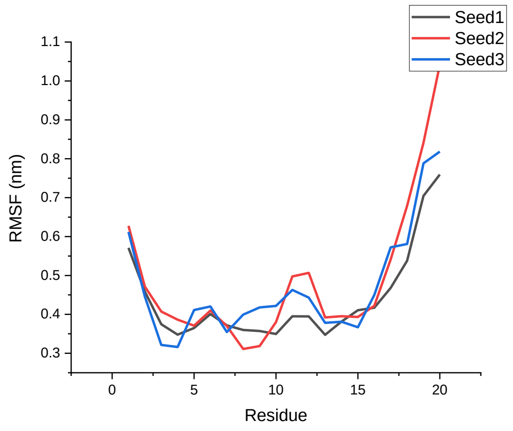
</td></tr>
</table>

<table>
<tr><td>GROMOS 54A8</td><td>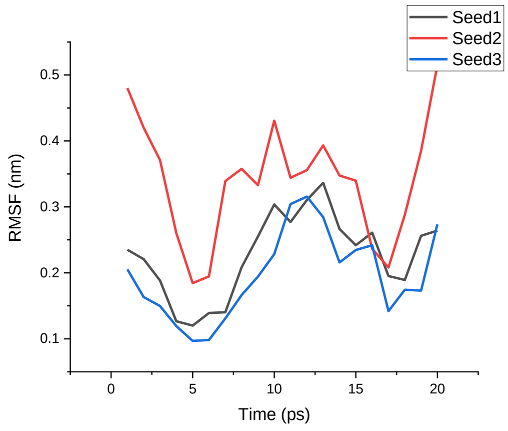</td><td>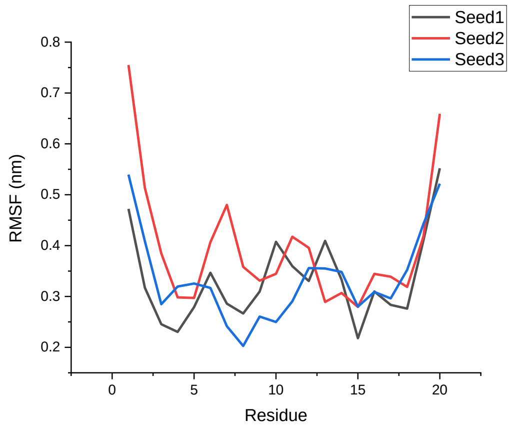</td></tr>
<tr><td></td><td>1ROO</td><td>β-1ROO</td></tr>
<tr><td>Amber 99SB-ILDN</td><td>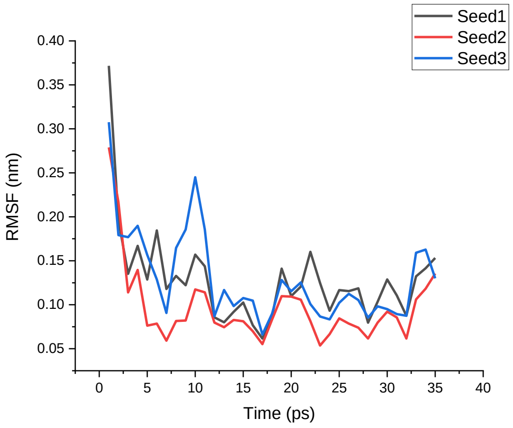</td><td>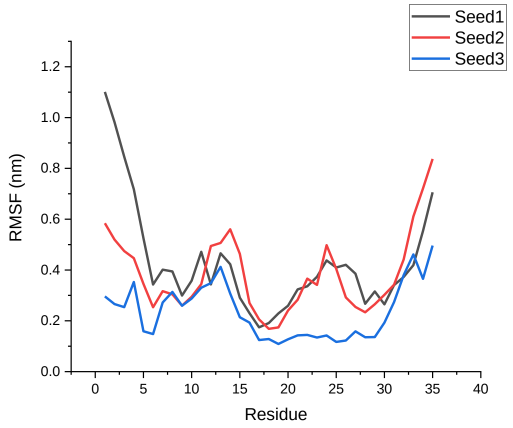</td></tr>
<tr><td>Amber 99SB*-ILDN</td><td>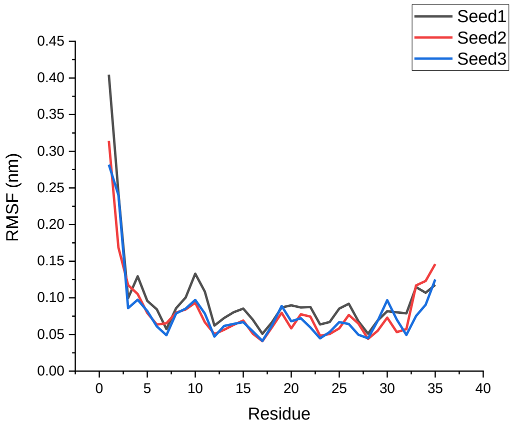</td><td>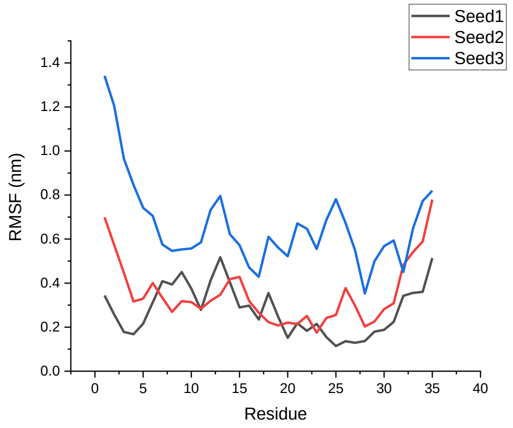</td></tr>
</table>

<table>
<tr><td>CHARMM22*</td><td>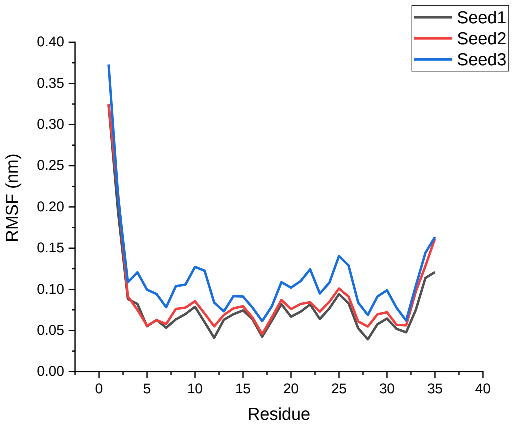</td><td>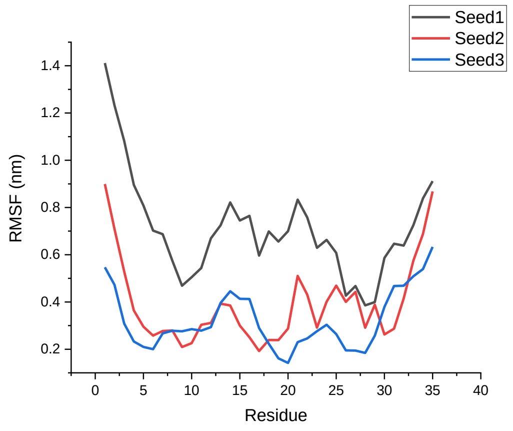</td></tr>
<tr><td>CHARMM36</td><td>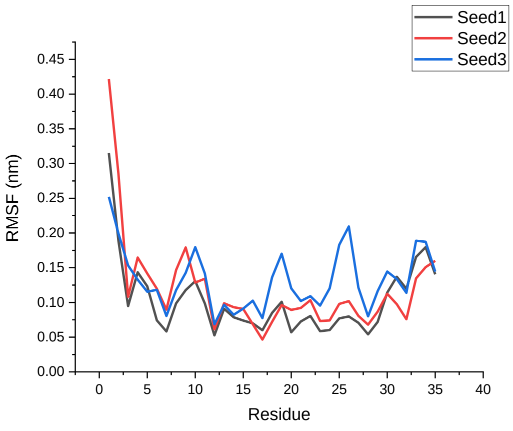</td><td>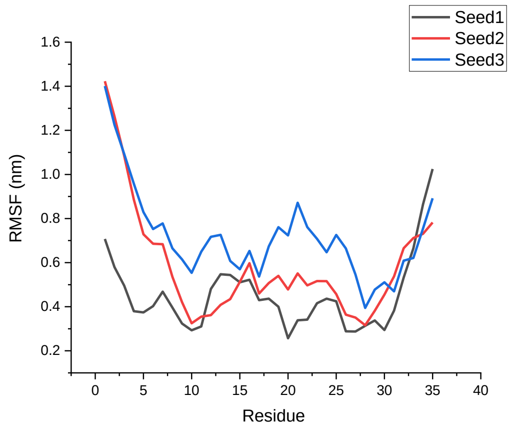</td></tr>
<tr><td>GROMOS 54A8</td><td>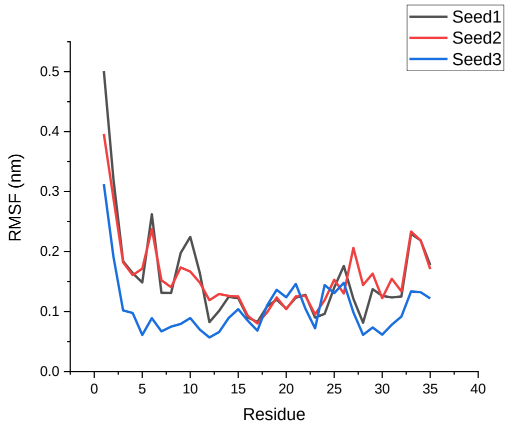</td><td>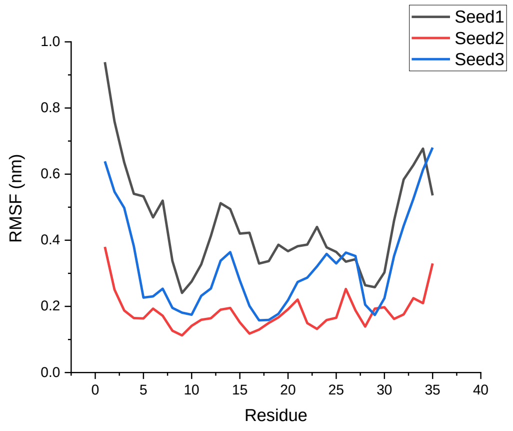</td></tr>
<tr><td></td><td>1T5Q</td><td>β-1T5Q</td></tr>
</table>

<table>
<tr>
<td>Amber 99SB-ILDN</td>
<td>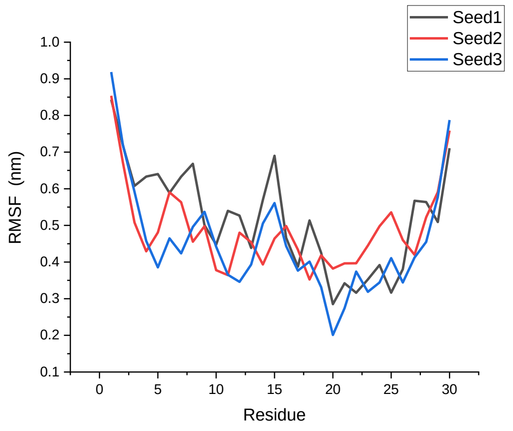
</td>
<td>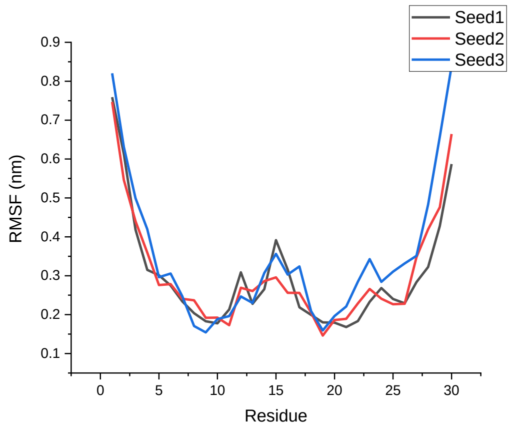
</td>
</tr>
<tr>
<td>Amber 99SB*-ILDN</td>
<td>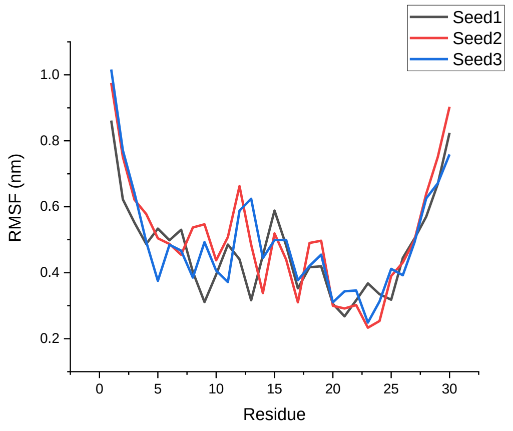
</td>
<td>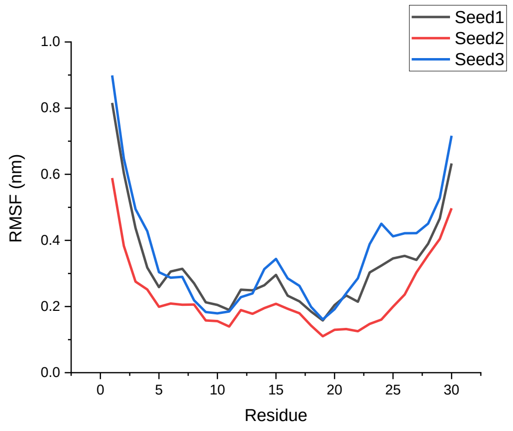
</td>
</tr>
<tr>
<td>CHARMM22*</td>
<td>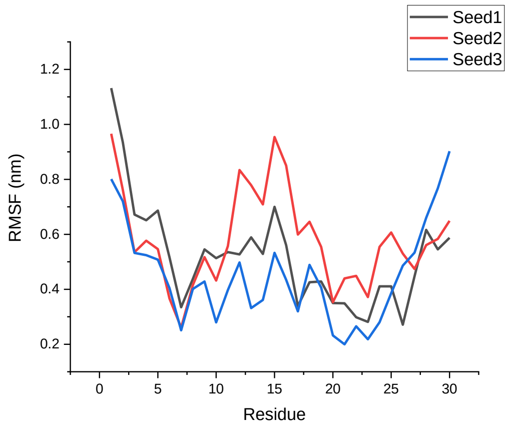
</td>
<td>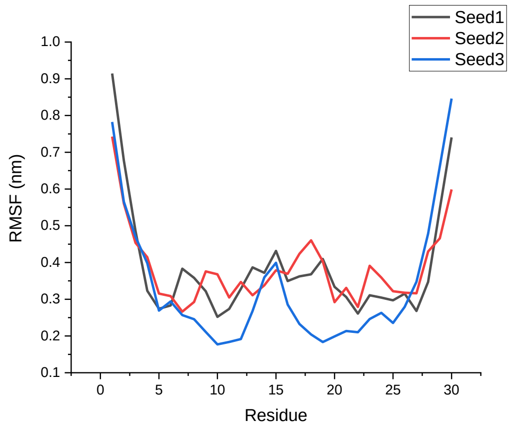
</td>
</tr>
</table>

<table>
<tr><td>CHARMM36</td><td>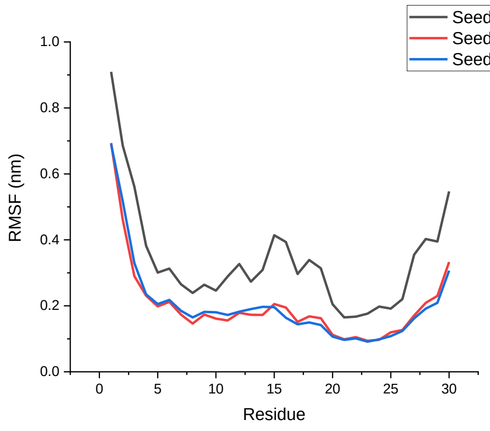
</td><td>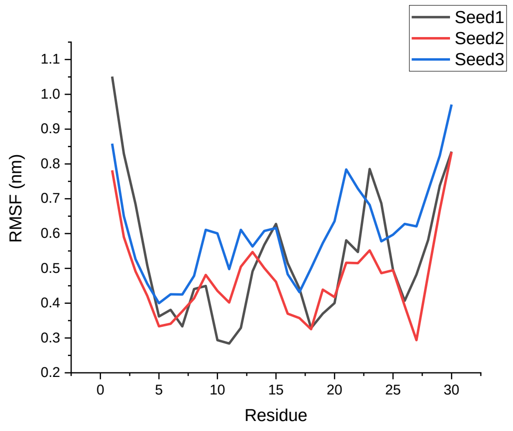
</td></tr>
<tr><td>GROMOS 54A8</td><td>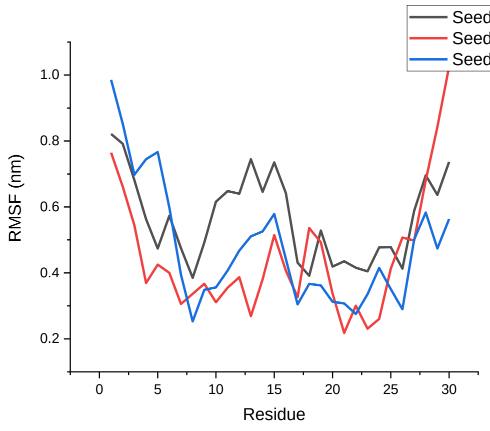
</td><td>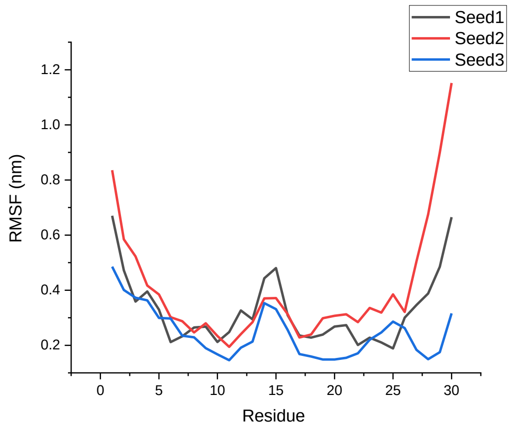
</td></tr>
<tr><td></td><td>1V1D</td><td>β-1V1D</td></tr>
<tr><td>Amber 99SB-ILDN</td><td>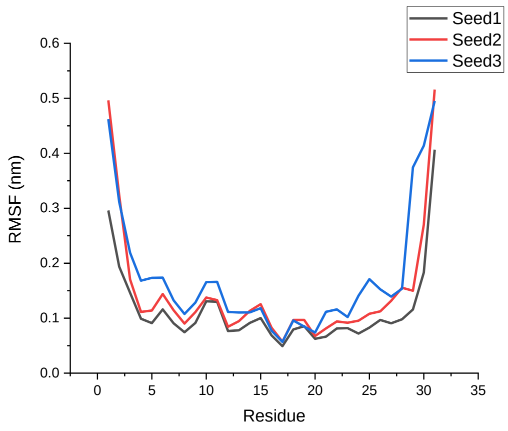
</td><td>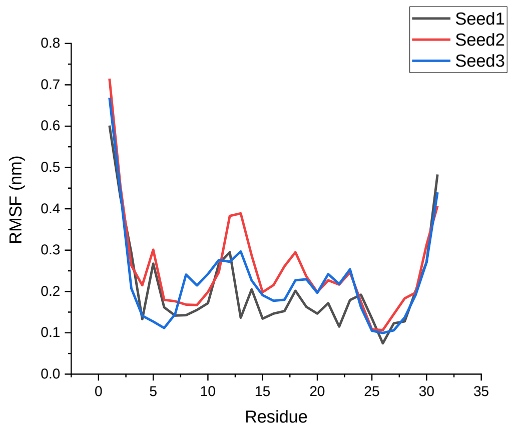
</td></tr>
</table>

| | | |
|---|---|---|
| Amber 99SB*-ILDN | 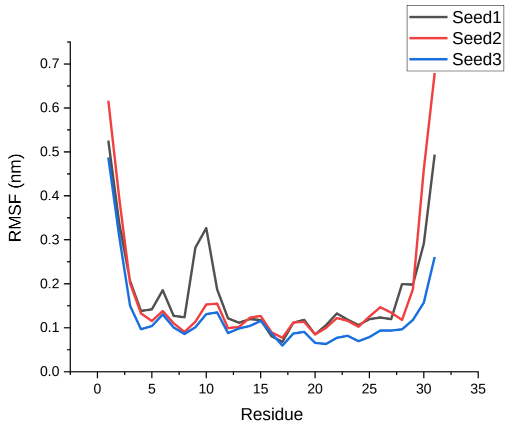 | 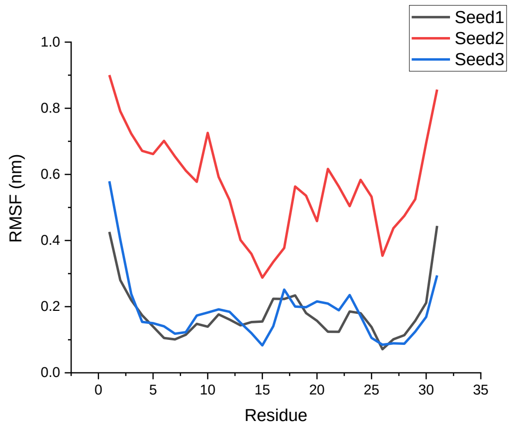 |
| CHARMM22* | 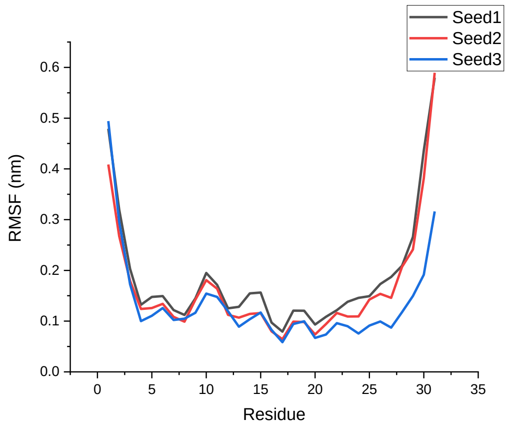 | 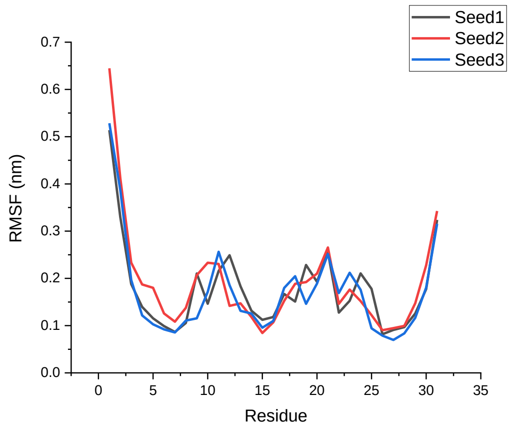 |
| CHARMM36 | 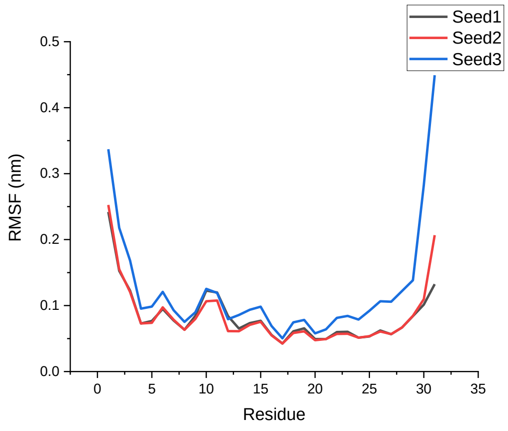 | 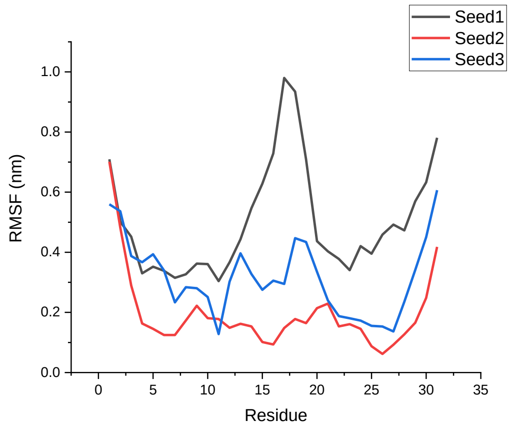 |

<table>
<tr><td>GROMOS 54A8</td><td>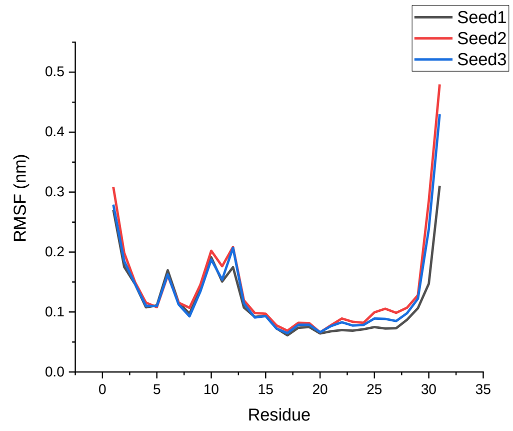
</td><td>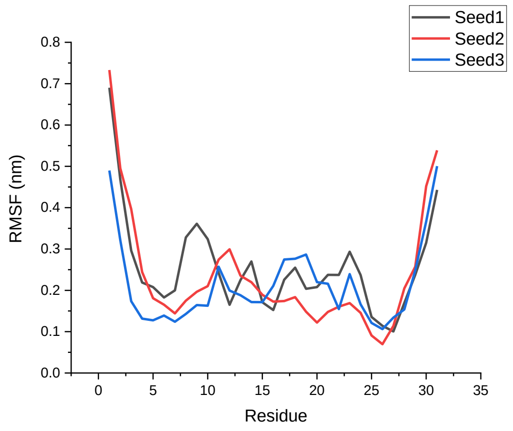
</td></tr>
<tr><td></td><td>1LFC</td><td>α-1LFC</td></tr>
<tr><td>Amber 99SB-ILDN</td><td>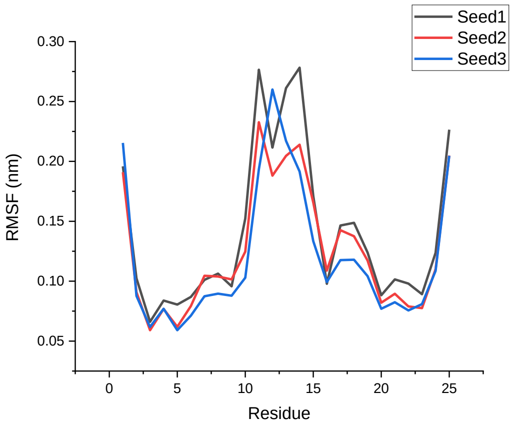
</td><td>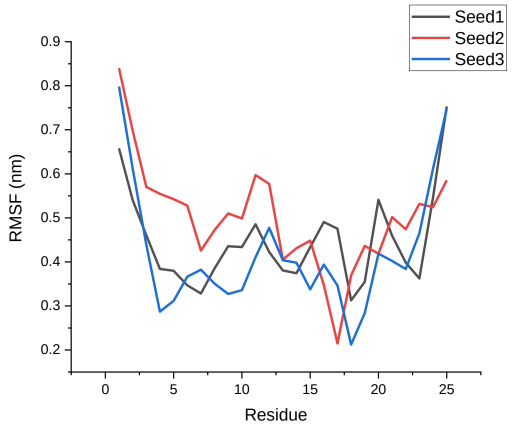
</td></tr>
<tr><td>Amber 99SB*-ILDN</td><td>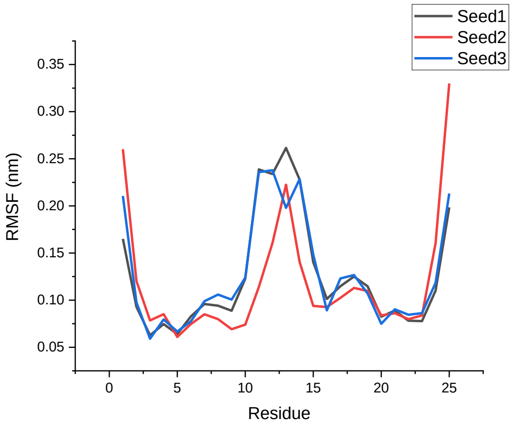
</td><td>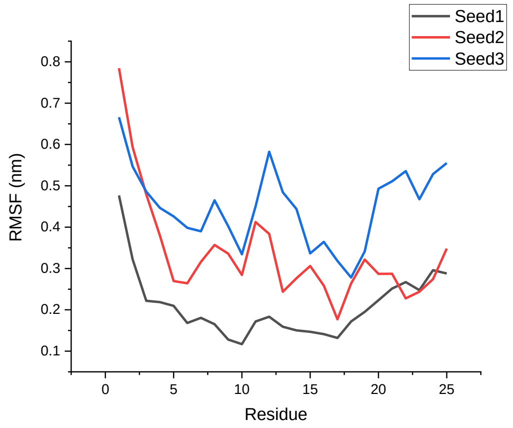
</td></tr>
</table>

| | | |
|---|---|---|
| CHARMM22* | 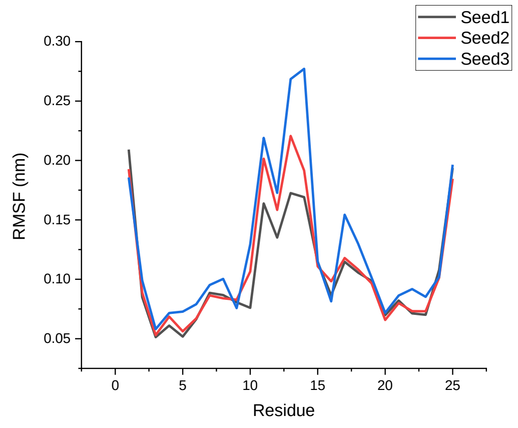 | 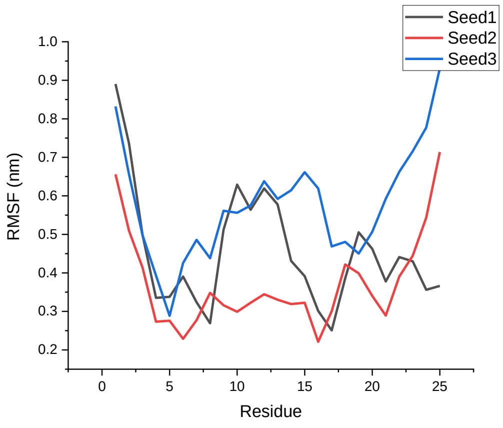 |
| CHARMM36 | 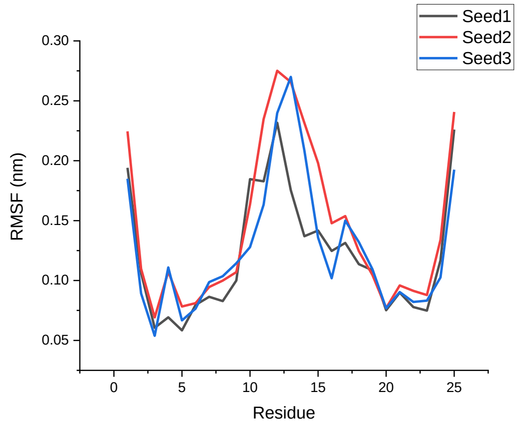 | 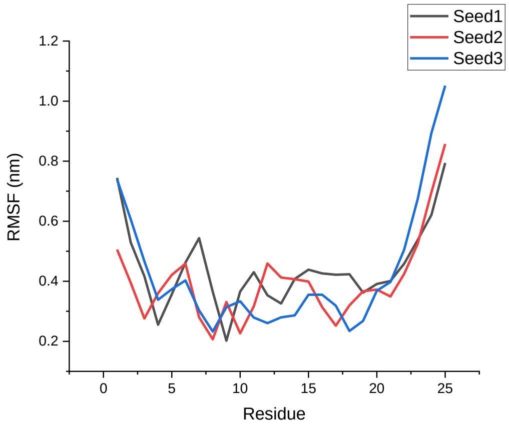 |
| GROMOS 54A8 | 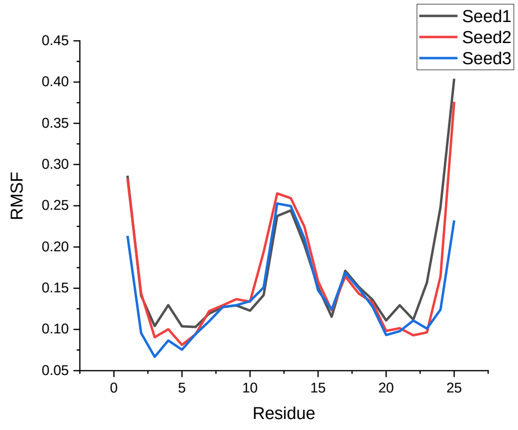 | 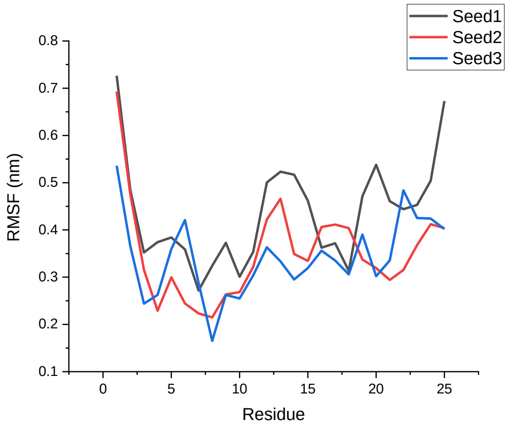 |
| | **8TFV** | **α-8TFV** |

| | | |
|---|---|---|
| Amber 99SB-ILDN | 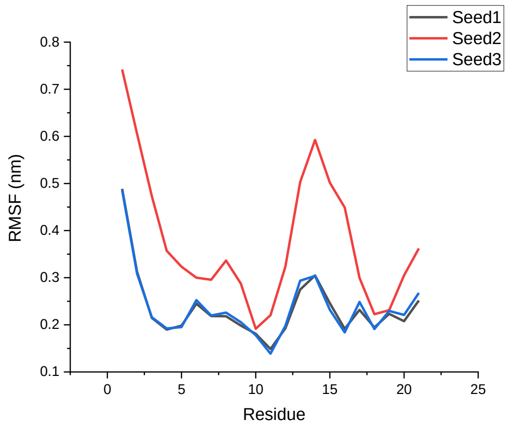 | 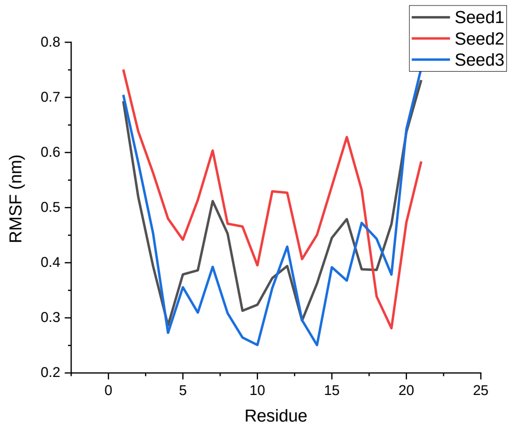 |
| Amber 99SB*-ILDN | | |
| CHARMM22* | 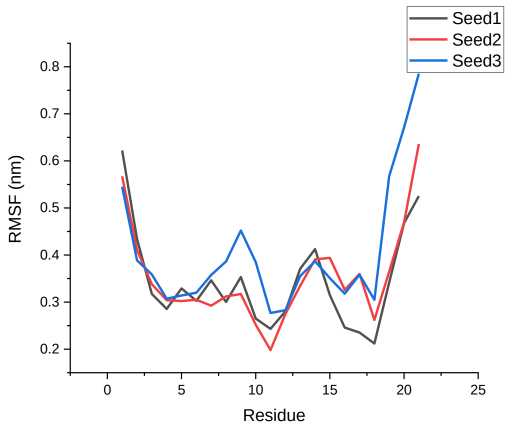 | 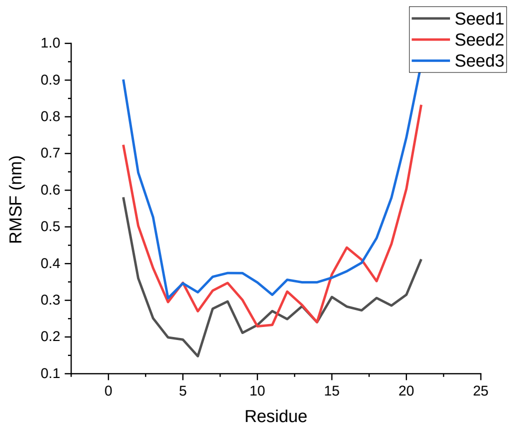 |
| CHARMM36 | 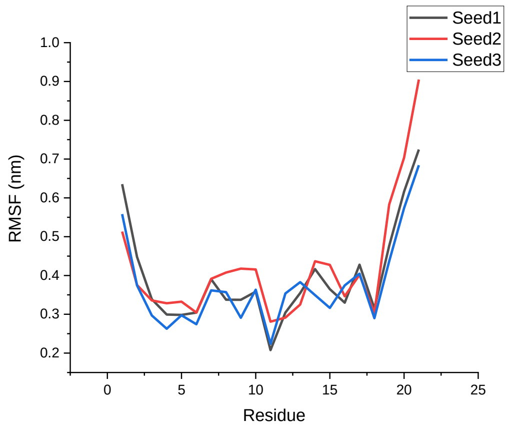 | 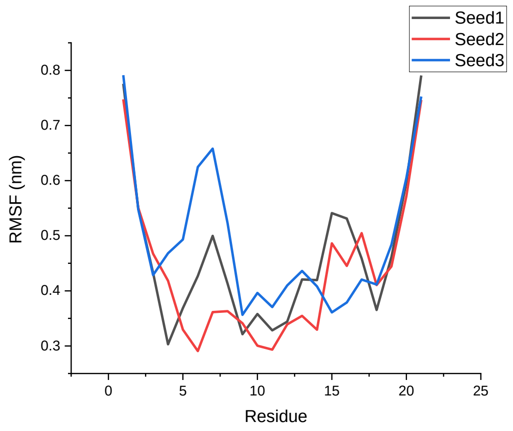 |

| | | |
|---|---|---|
| GROMOS 54A8 | 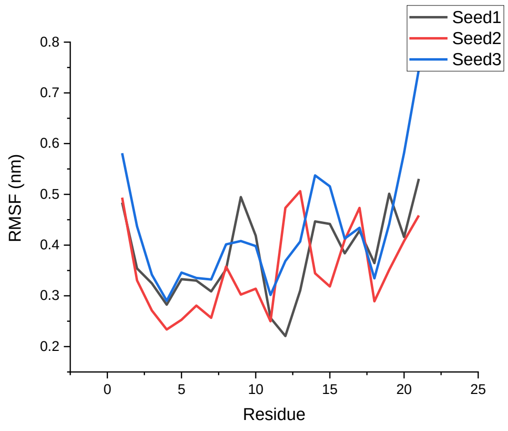  | 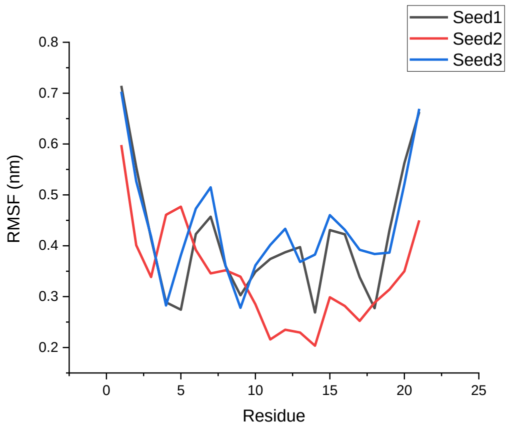  |
| | **1KGM** | **α-1KGM** |
| Amber 99SB-ILDN | 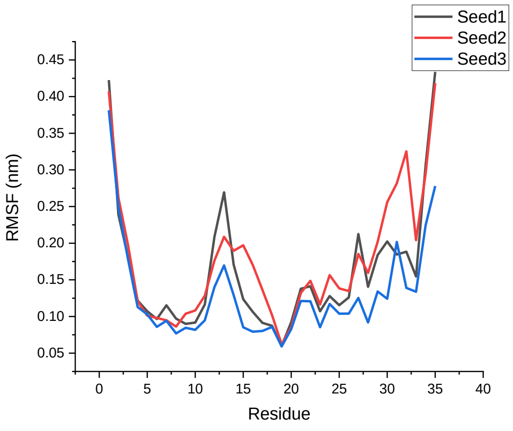  | 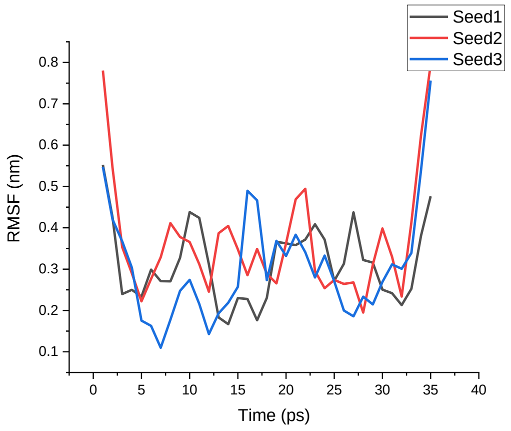  |
| Amber 99SB*-ILDN | 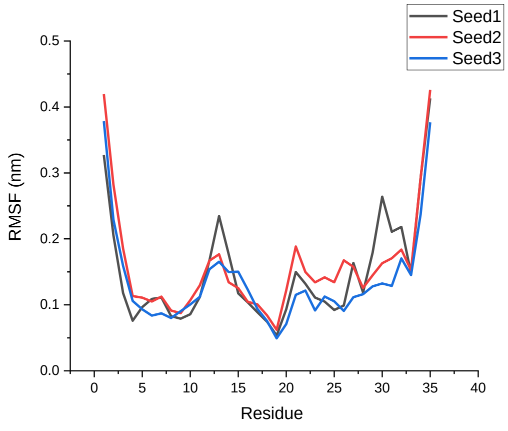  | 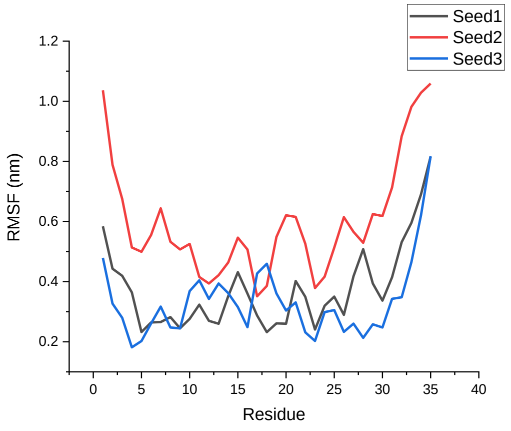  |

<table>
<tr>
<td>CHARMM22*</td>
<td>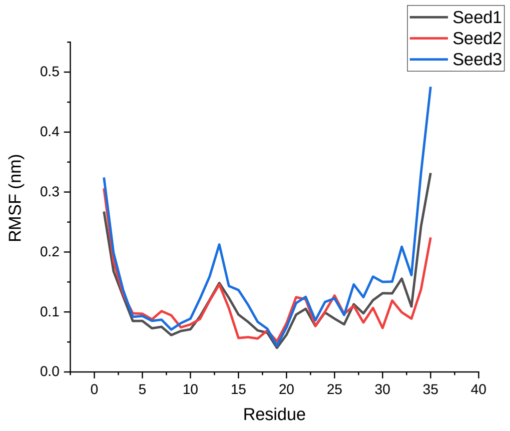
</td>
<td>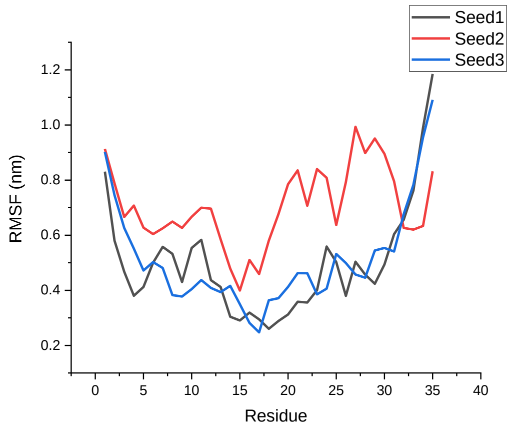
</td>
</tr>
<tr>
<td>CHARMM36</td>
<td>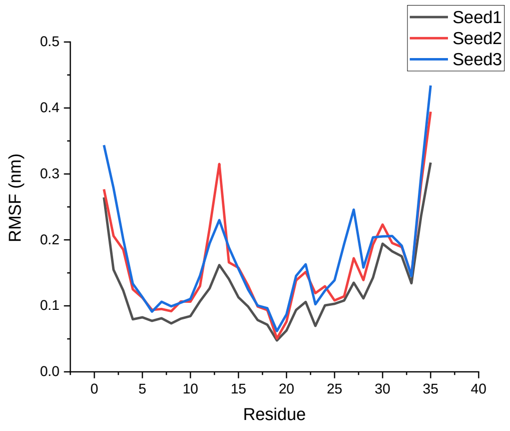
</td>
<td>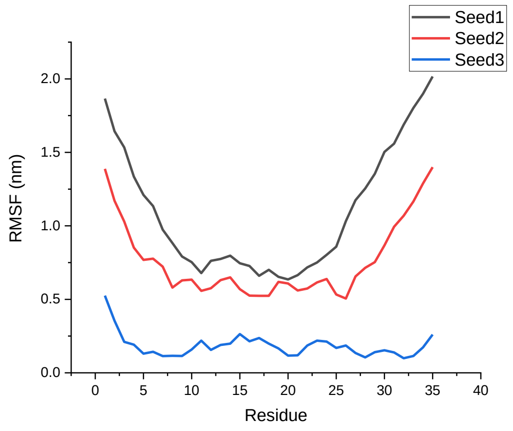
</td>
</tr>
<tr>
<td>GROMOS 54A8</td>
<td>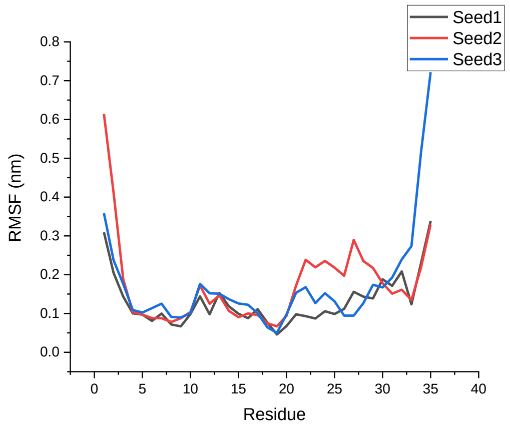
</td>
<td>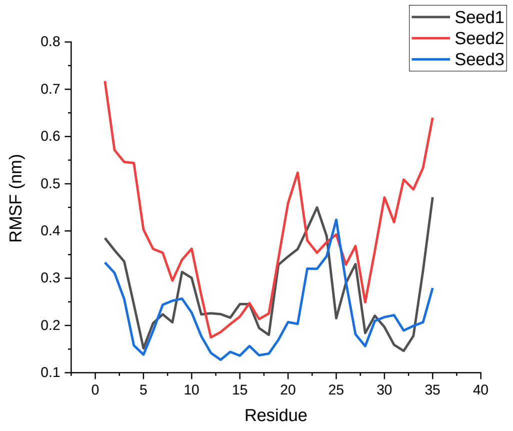
</td>
</tr>
<tr>
<td></td>
<td>1M4F</td>
<td>α-1M4F</td>
</tr>
</table>

| | | |
|---|---|---|
| Amber 99SB-ILDN | 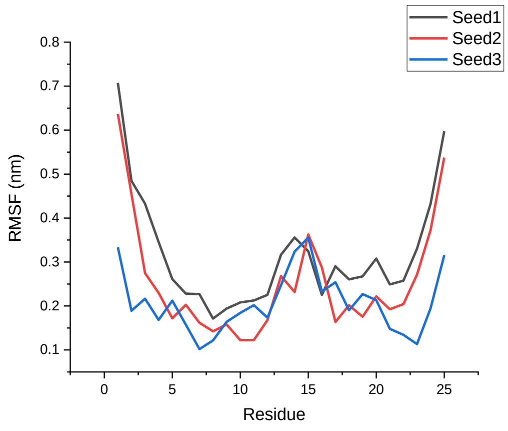 | 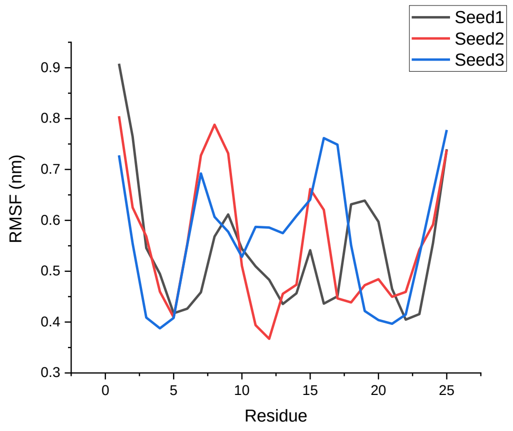 |
| Amber 99SB*-ILDN | 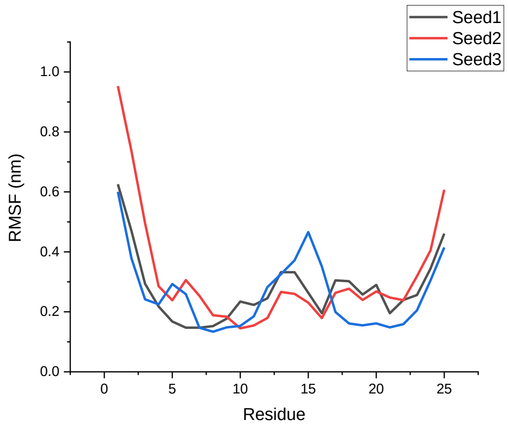 | 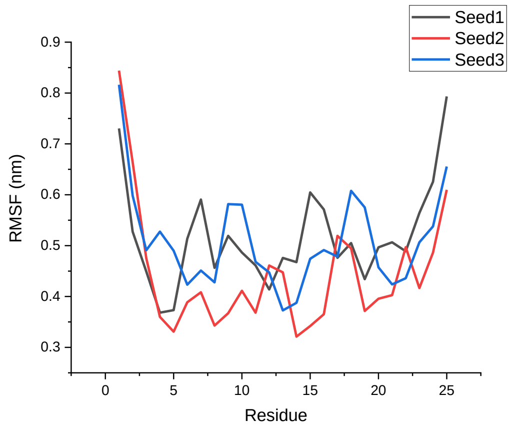 |
| CHARMM22* | 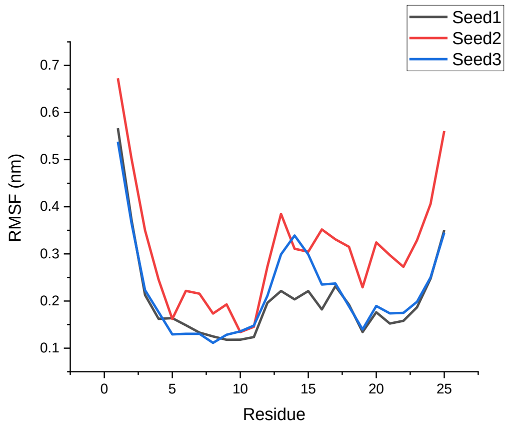 | 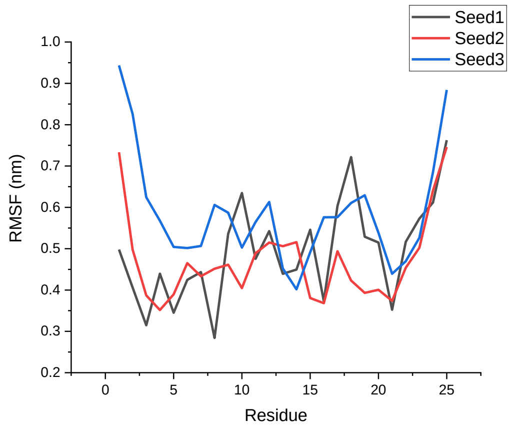 |

| | | |
|---|---|---|
| CHARMM36 | 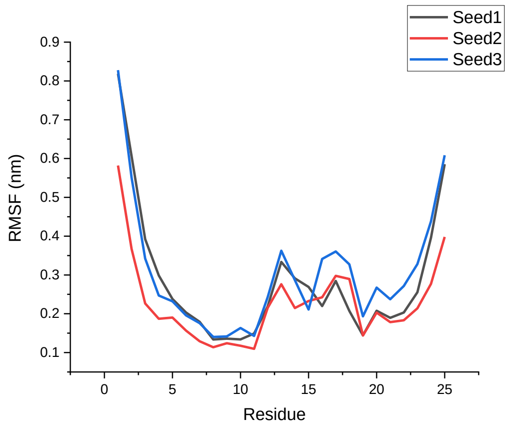 | 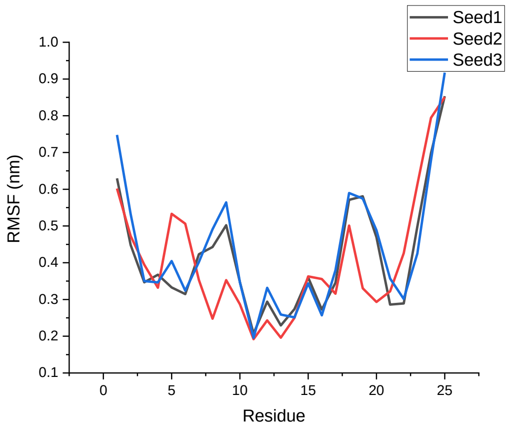 |
| GROMOS 54A8 | 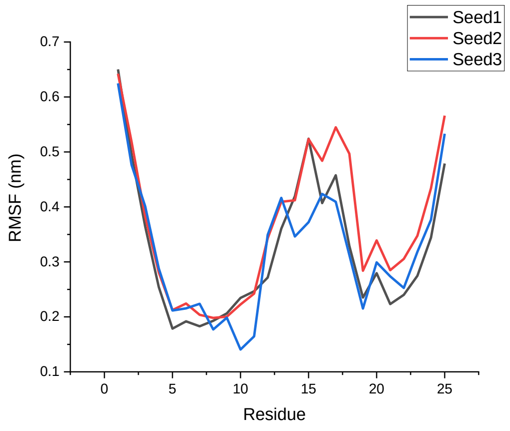 | 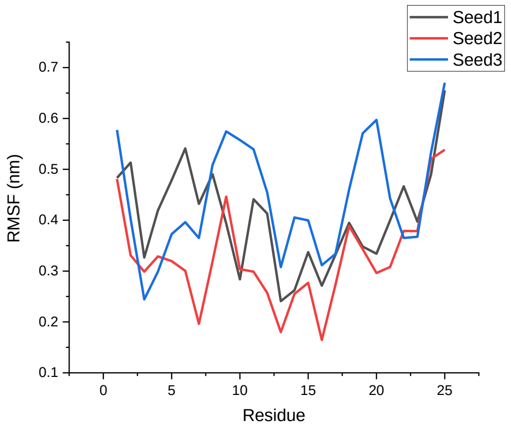 |
| | **1TV0** | **α-1TV0** |
| Amber 99SB-ILDN | 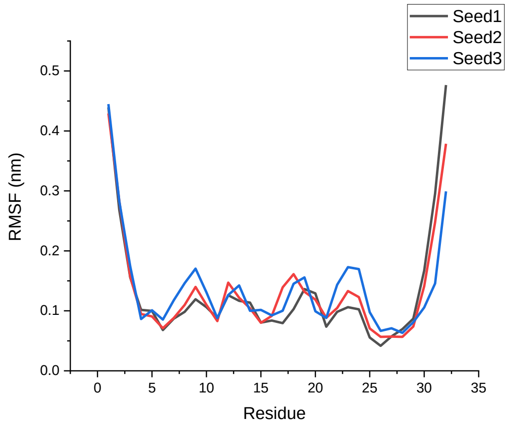 | 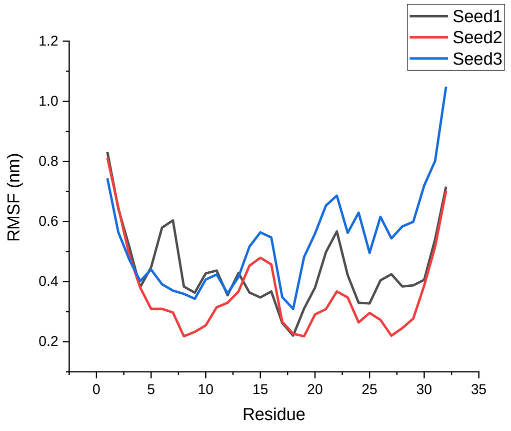 |

| | | |
|---|---|---|
| Amber 99SB*-ILDN | 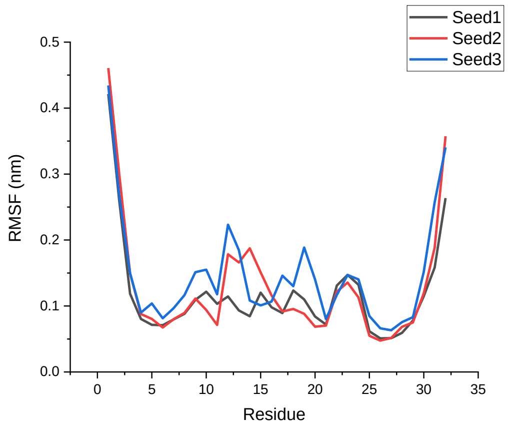 | 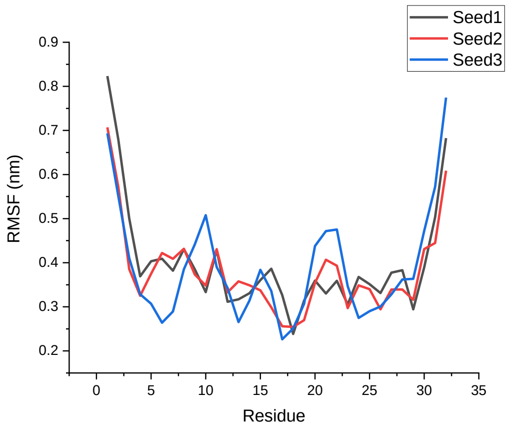 |
| CHARMM22* | 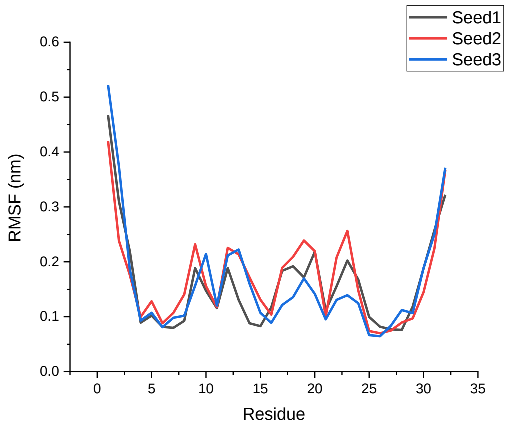 | 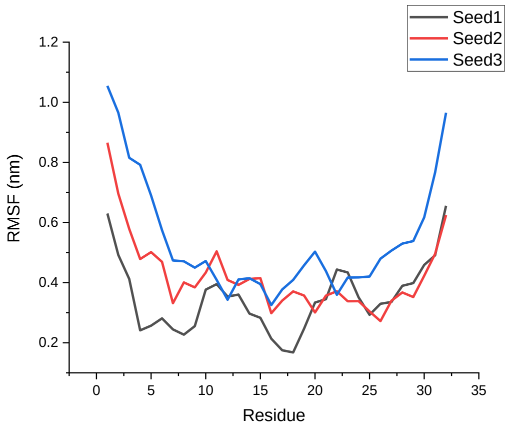 |
| CHARMM36 | 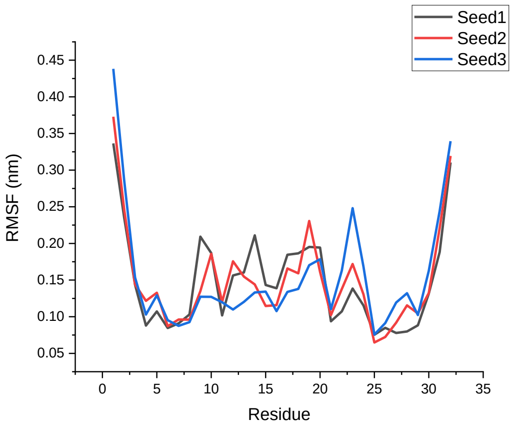 | 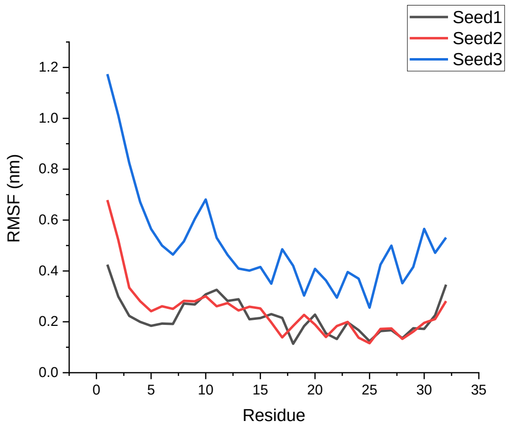 |

| | | |
|---|---|---|
| GROMOS 54A8 | 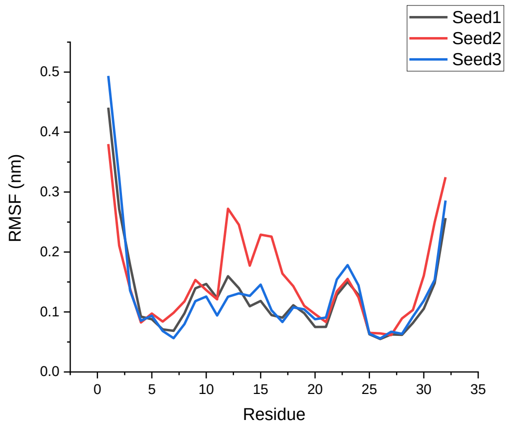 Seed1<br>Seed2<br>Seed3<br>RMSF (nm)<br>0.0<br>0.1<br>0.2<br>0.3<br>0.4<br>0.5<br>0<br>5<br>10<br>15<br>20<br>25<br>30<br>35<br>Residue | 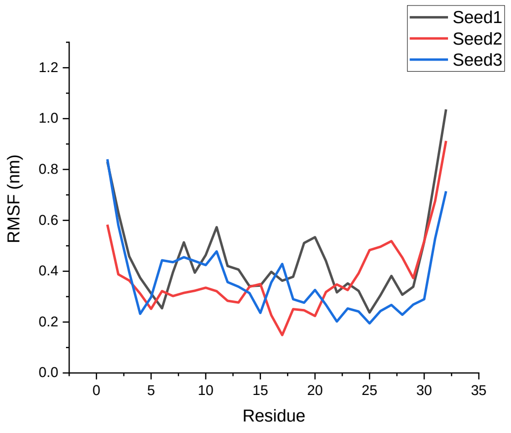 Seed1<br>Seed2<br>Seed3<br>RMSF (nm)<br>0.0<br>0.2<br>0.4<br>0.6<br>0.8<br>1.0<br>1.2<br>0<br>5<br>10<br>15<br>20<br>25<br>30<br>35<br>Residue |

**Table S6**. Average RMSF of the native α-proteins, modeled β-proteins, β-proteins, and modeled α-proteins.

| | RMSF | SD | RMSF | SD | RMSF | SD | RMSF | SD | RMSF | SD |
|---|---|---|---|---|---|---|---|---|---|---|
| α- proteins | | | | | | | | | | |
| | **1CQ0** | | **1L2Y** | | **1ROO** | | **1T5Q** | | **1V1D** | |
| Amber 99SB-ILDN | 0.36 | 0.10 | 0.08 | 0.05 | 0.12 | 0.05 | 0.49 | 0.13 | 0.15 | 0.10 |
| Amber 99SB*-ILDN | 0.34 | 0.11 | 0.07 | 0.04 | 0.09 | 0.05 | 0.49 | 0.16 | 0.16 | 0.12 |
| CHARMM22* | 0.36 | 0.10 | 0.12 | 0.06 | 0.09 | 0.05 | 0.52 | 0.18 | 0.16 | 0.11 |
| CHARMM36 | 0.36 | 0.12 | 0.09 | 0.04 | 0.12 | 0.05 | 0.25 | 0.14 | 0.10 | 0.06 |
| GROMOS 54A8 | 0.40 | 0.12 | 0.25 | 0.07 | 0.14 | 0.06 | 0.50 | 0.17 | 0.13 | 0.07 |
| Modelled β-proteins | | | | | | | | | | |
| | **β-1CQ0** | | **β-1L2Y** | | **β-1ROO** | | **β-1T5Q** | | **β-1V1D** | |
| Amber 99SB-ILDN | 0.61 | 0.19 | 0.42 | 0.13 | 0.35 | 0.16 | 0.32 | 0.16 | 0.23 | 0.12 |
| Amber 99SB*-ILDN | 0.47 | 0.13 | 0.44 | 0.15 | 0.43 | 0.15 | 0.30 | 0.14 | 0.31 | 0.11 |
| CHARMM22* | 0.68 | 0.21 | 0.43 | 0.15 | 0.47 | 0.17 | 0.37 | 0.14 | 0.18 | 0.10 |
| CHARMM36 | 0.72 | 0.20 | 0.46 | 0.15 | 0.59 | 0.21 | 0.53 | 0.15 | 0.34 | 0.15 |
| GROMOS 54A8 | 0.48 | 0.10 | 0.35 | 0.10 | 0.59 | 0.10 | 0.32 | 0.15 | 0.24 | 0.12 |
| β- proteins | | | | | | | | | | |
| | **1LFC** | | **8TFV** | | **1KGM** | | **1M4F** | | **1TV0** | |
| Amber 99SB-ILDN | 0.13 | 0.06 | 0.28 | 0.10 | 0.16 | 0.08 | 0.26 | 0.11 | 0.13 | 0.09 |
| Amber 99SB*-ILDN | 0.12 | 0.06 | 0.30 | 0.08 | 0.15 | 0.08 | 0.28 | 0.14 | 0.13 | 0.08 |
| CHARMM22* | 0.11 | 0.05 | 0.37 | 0.11 | 0.12 | 0.07 | 0.24 | 0.11 | 0.16 | 0.09 |
| CHARMM36 | 0.13 | 0.06 | 0.40 | 0.13 | 0.15 | 0.07 | 0.27 | 0.14 | 0.15 | 0.07 |
| GROMOS 54A8 | 0.15 | 0.07 | 0.39 | 0.09 | 0.16 | 0.10 | 0.34 | 0.13 | 0.14 | 0.08 |
| Modelled α- proteins | | | | | | | | | | |
| | **α-1LFC** | | **α-8TFV** | | **α-1KGM** | | **α-1M4F** | | **α-1TV0** | |
| Amber 99SB-ILDN | 0.46 | 0.12 | 0.45 | 0.13 | 0.33 | 0.12 | 0.55 | 0.13 | 0.44 | 0.14 |
| Amber 99SB*-ILDN | 0.33 | 0.10 | 0.38 | 0.11 | 0.44 | 0.15 | 0.49 | 0.10 | 0.39 | 0.12 |
| CHARMM22* | 0.47 | 0.14 | 0.38 | 0.15 | 0.57 | 0.18 | 0.52 | 0.12 | 0.44 | 0.15 |
| CHARMM36 | 0.42 | 0.16 | 0.46 | 0.13 | 0.68 | 0.26 | 0.42 | 0.17 | 0.32 | 0.13 |
| GROMOS 54A8 | 0.38 | 0.10 | 0.40 | 0.11 | 0.29 | 0.10 | 0.39 | 0.10 | 0.39 | 0.16 |

**Table S7**. $R_g$ plots of the native α-proteins, modeled β-proteins, β-proteins, and modeled α-proteins.

<table>
<tr><th colspan="3">$R_g$</th></tr>
<tr><th></th><th>1CQ0</th><th>β-1CQ0</th></tr>
<tr><td>Amber 99SB-ILDN</td><td>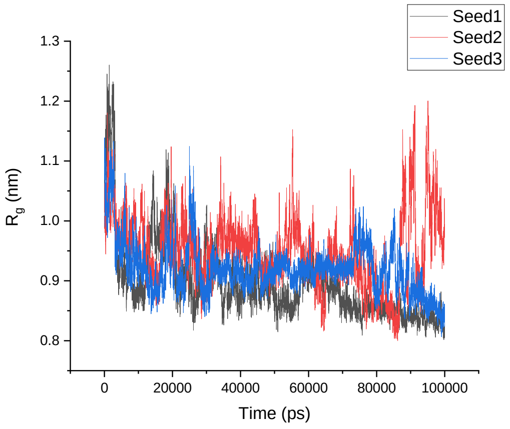
</td><td>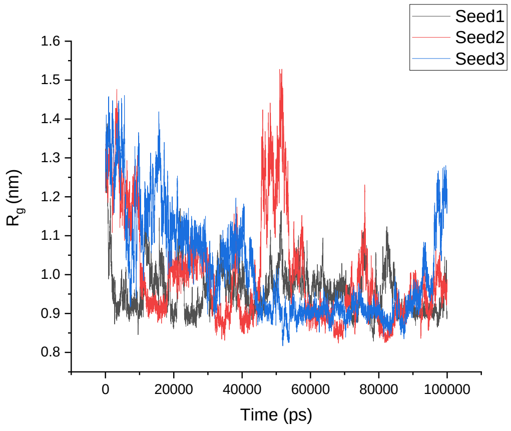
</td></tr>
<tr><td>Amber 99SB*-ILDN</td><td>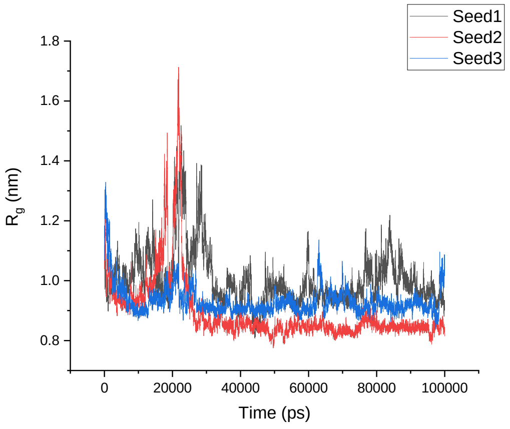
</td><td>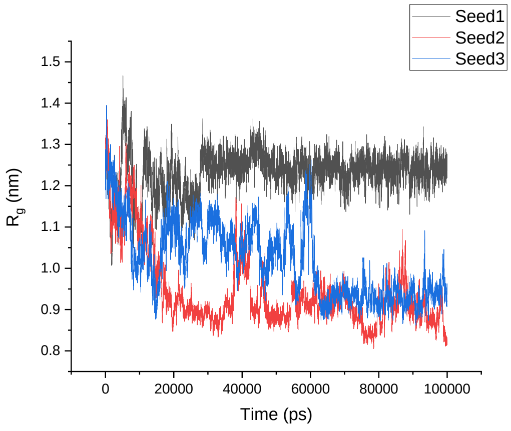
</td></tr>
<tr><td>CHARMM22*</td><td>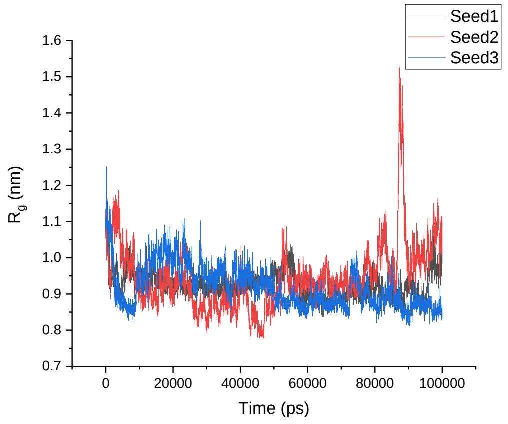
</td><td>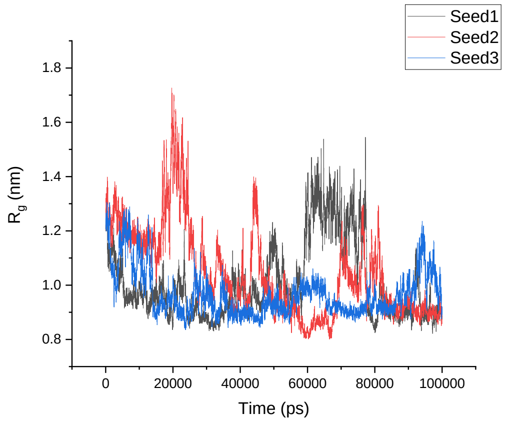
</td></tr>
</table>

<table>
<tr><td>CHARMM36</td><td>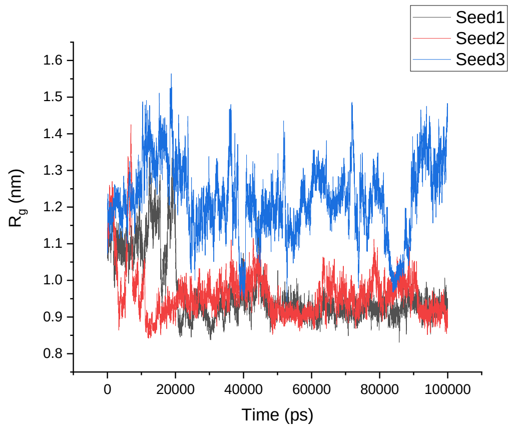
</td><td>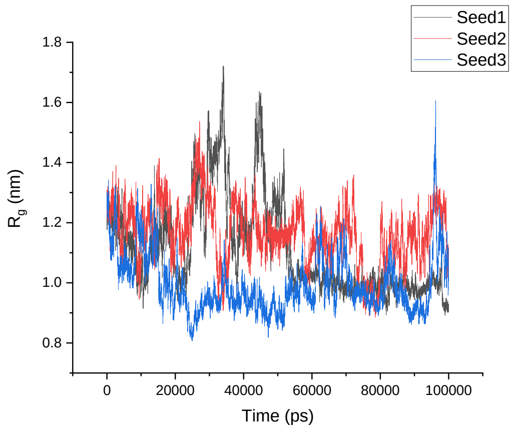
</td></tr>
<tr><td>GROMOS 54A8</td><td>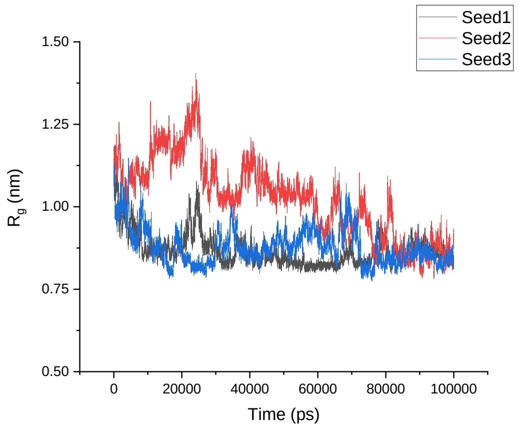
</td><td>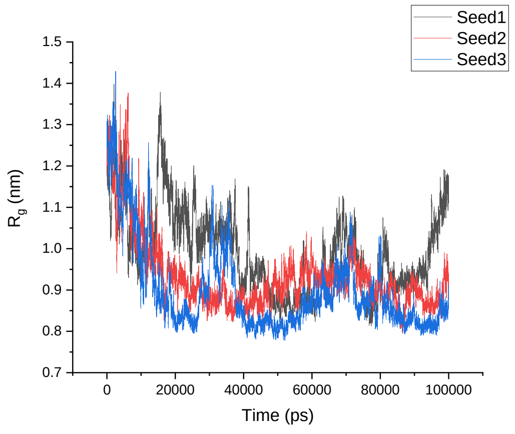
</td></tr>
<tr><td></td><td>1L2Y</td><td>β-1L2Y</td></tr>
<tr><td>Amber 99SB-ILDN</td><td>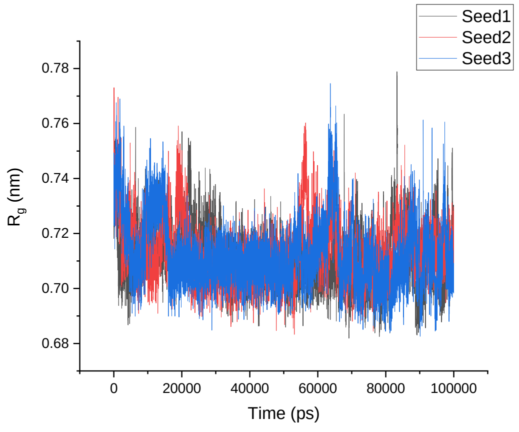
</td><td>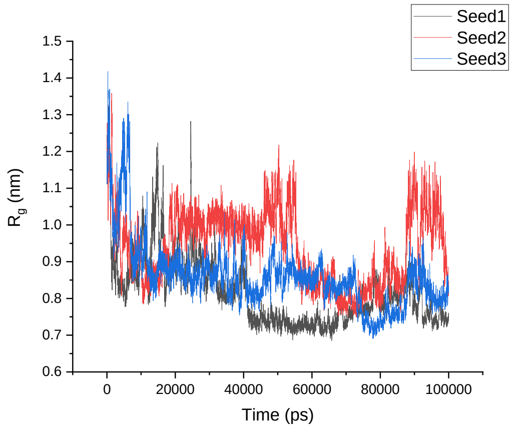
</td></tr>
</table>

Amber 99SB*-ILDN

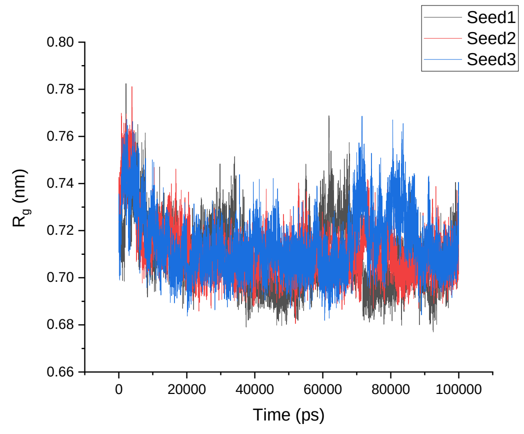


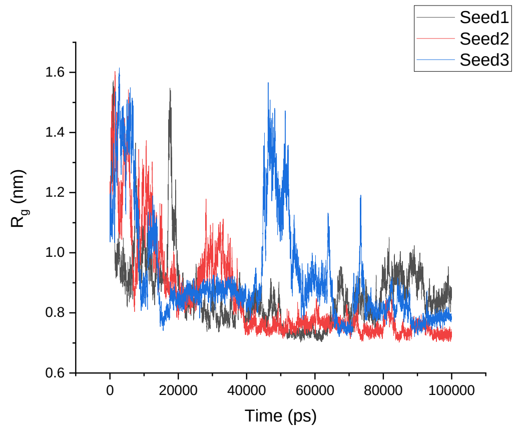


CHARMM22*

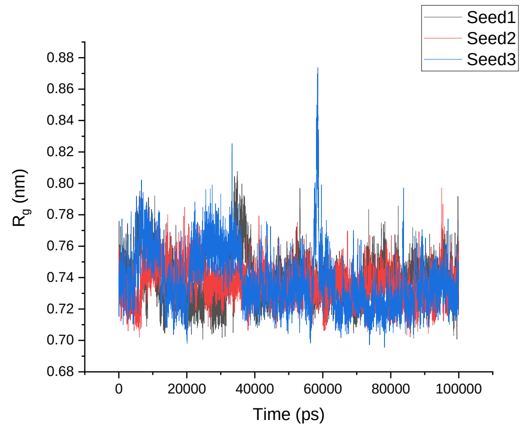


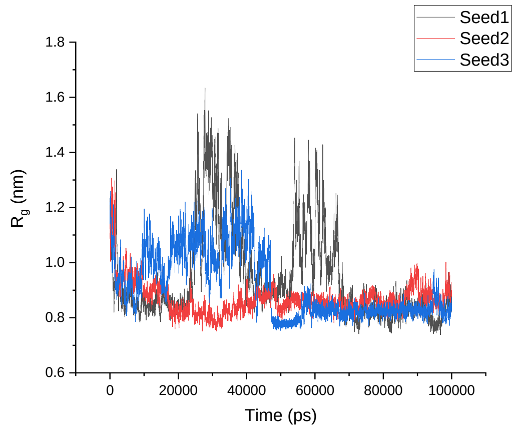


CHARMM36

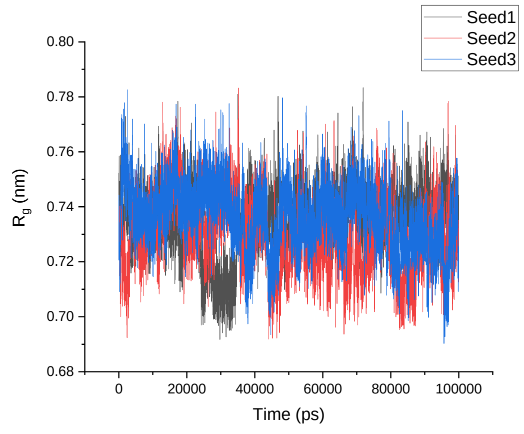


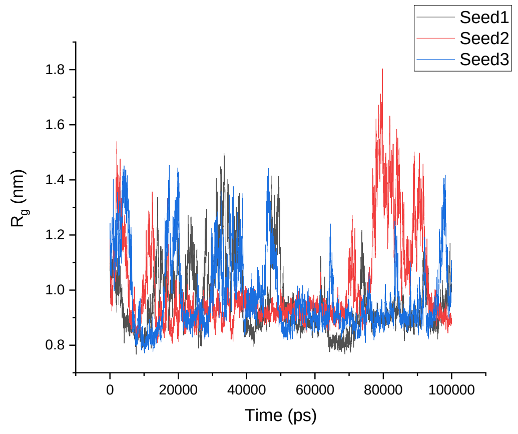

<table>
<tr><td>GROMOS 54A8</td><td>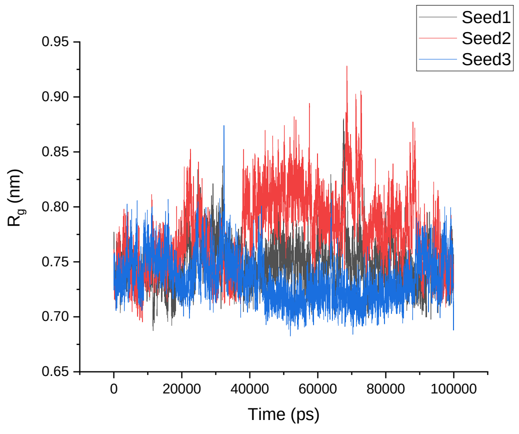
</td><td>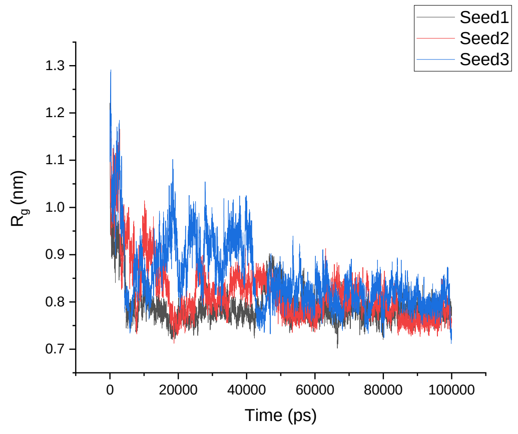
</td></tr>
<tr><td></td><td>1ROO</td><td>β-1ROO</td></tr>
<tr><td>Amber 99SB-ILDN</td><td>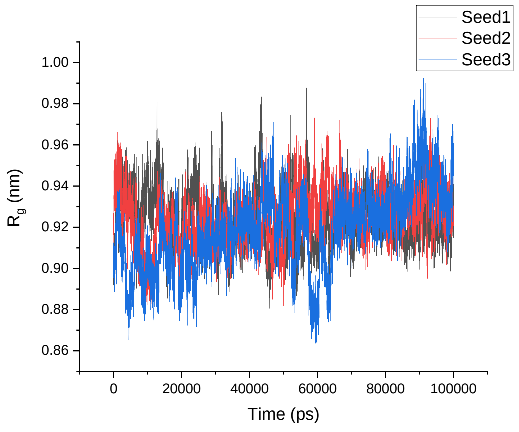
</td><td>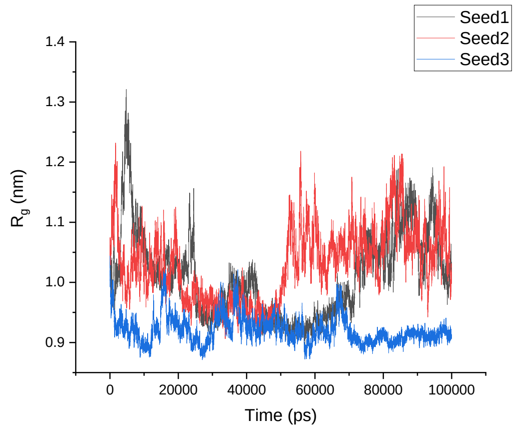
</td></tr>
<tr><td>Amber 99SB*-ILDN</td><td>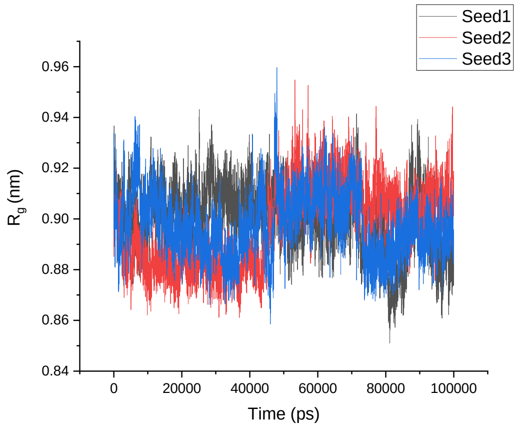
</td><td>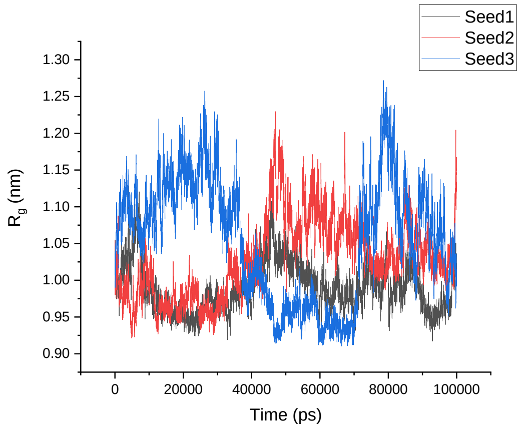
</td></tr>
</table>

<table>
<tr><td>CHARMM22*</td><td>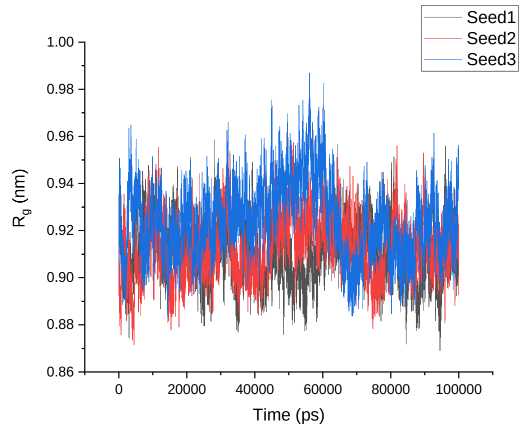
</td><td>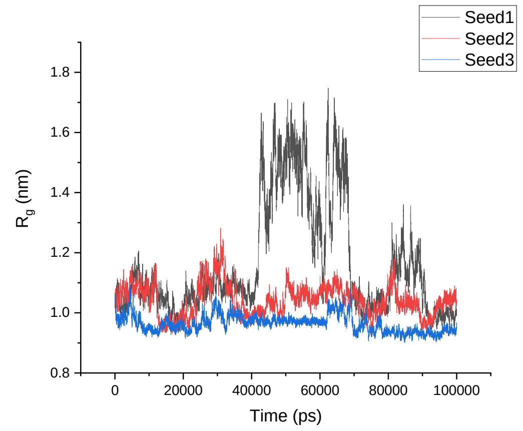
</td></tr>
<tr><td>CHARMM36</td><td>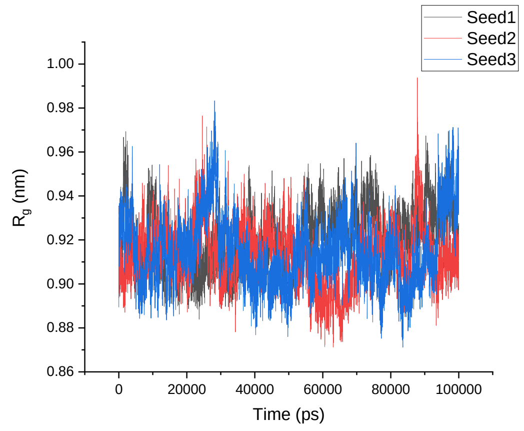
</td><td>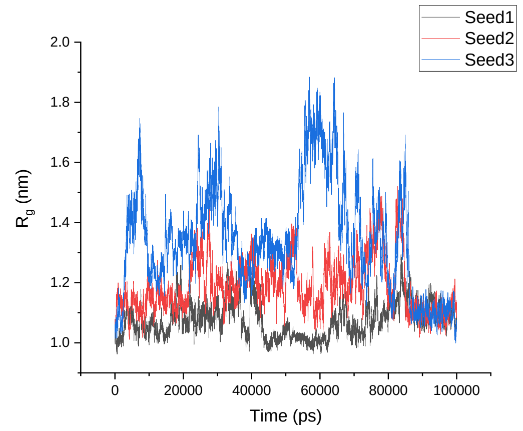
</td></tr>
<tr><td>GROMOS<br>54A8</td><td>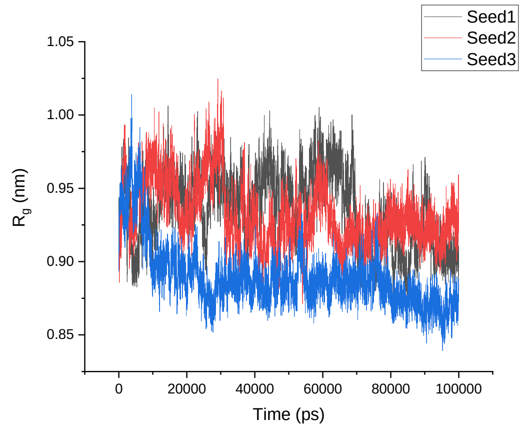
</td><td>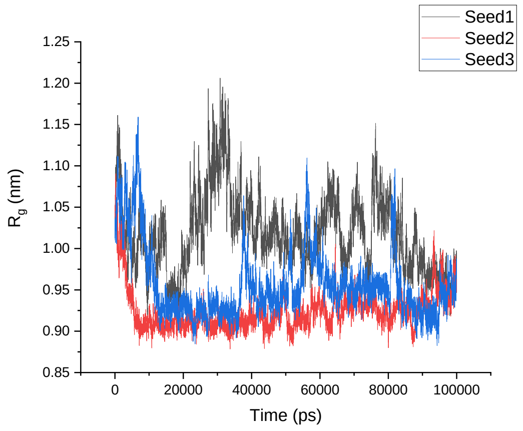
</td></tr>
<tr><td></td><td>1T5Q</td><td>β-1T5Q</td></tr>
</table>

| | | |
|---|---|---|
| Amber 99SB-ILDN | 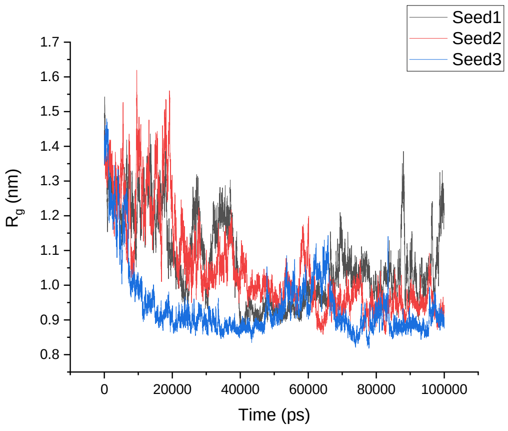 | 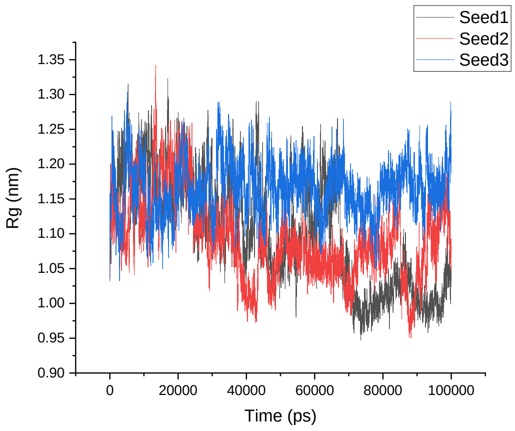 |
| Amber 99SB*-ILDN | 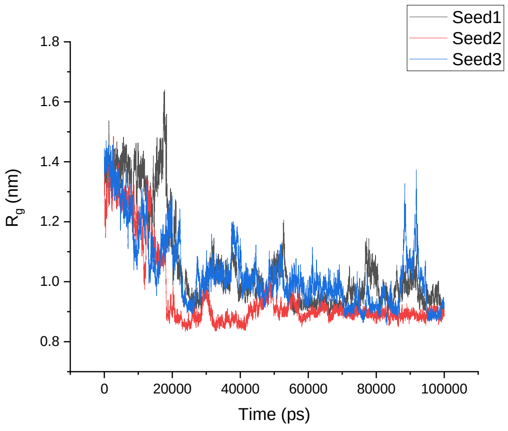 | 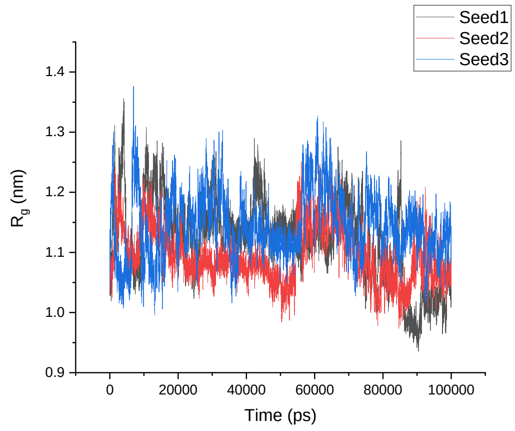 |
| CHARMM22* | 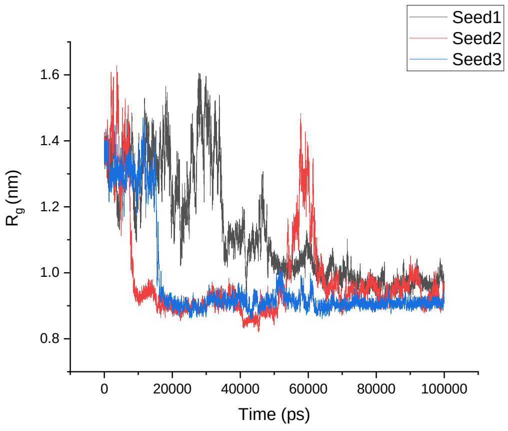 | 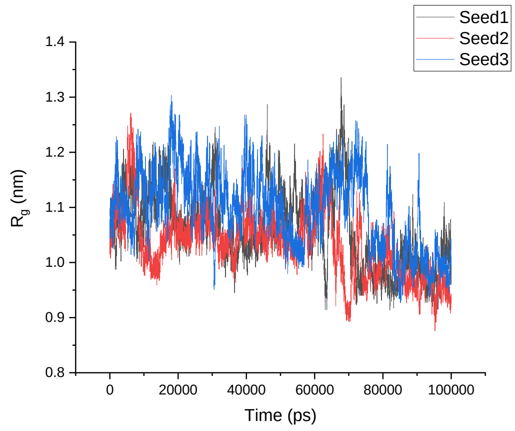 |

<table>
<tr><td>CHARMM36</td><td>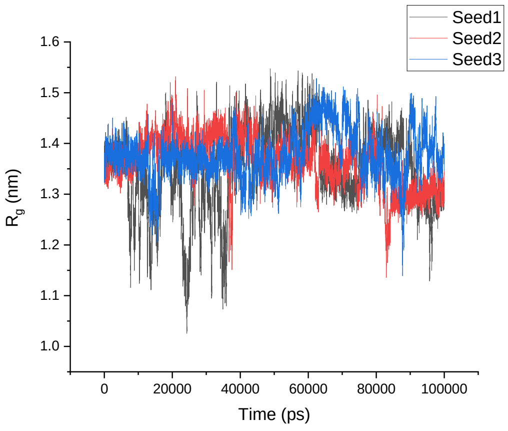
</td><td>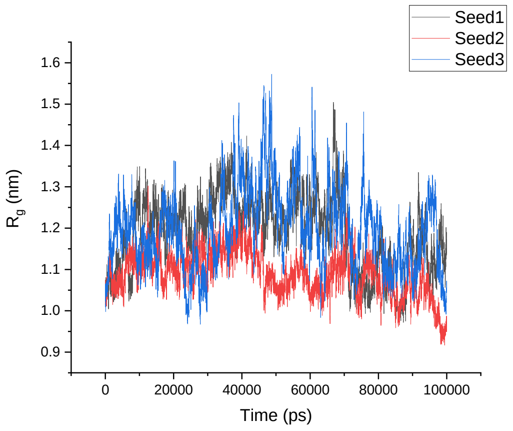
</td></tr>
<tr><td>GROMOS 54A8</td><td>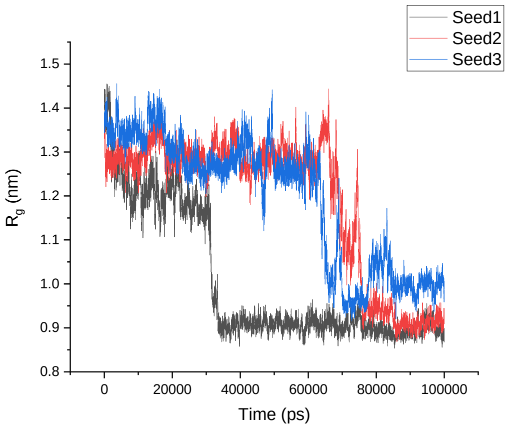
</td><td>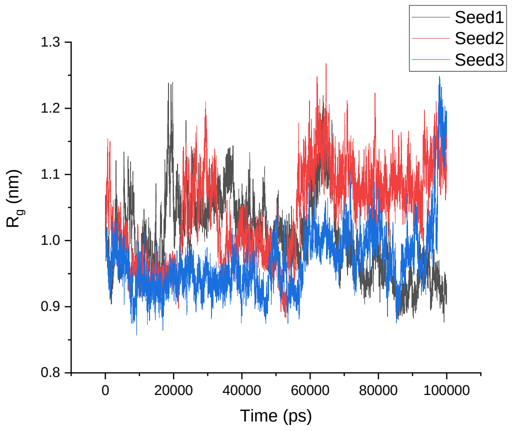
</td></tr>
<tr><td></td><td>1V1D</td><td>β-1V1D</td></tr>
<tr><td>Amber 99SB-ILDN</td><td>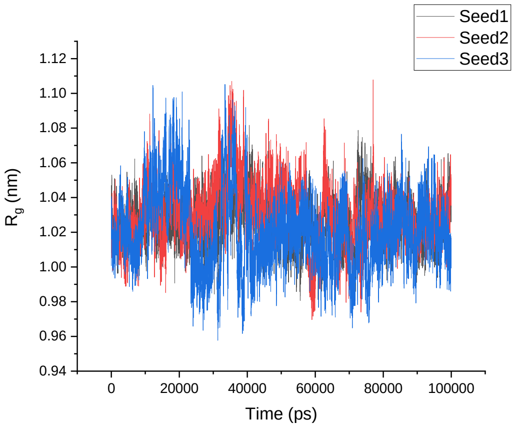
</td><td>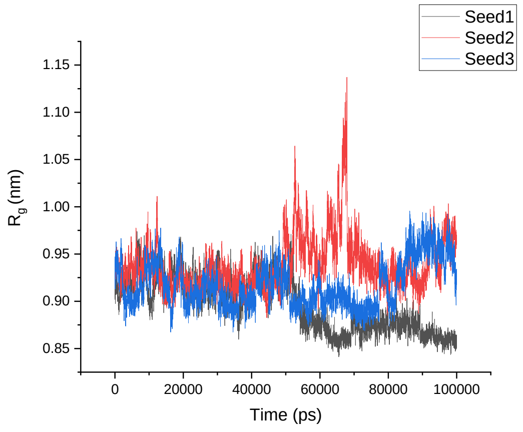
</td></tr>
</table>

Amber 99SB*-ILDN

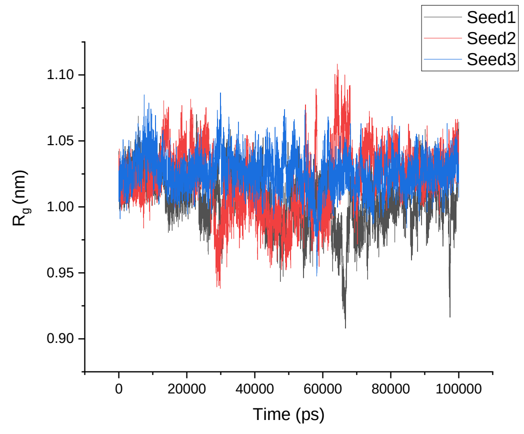


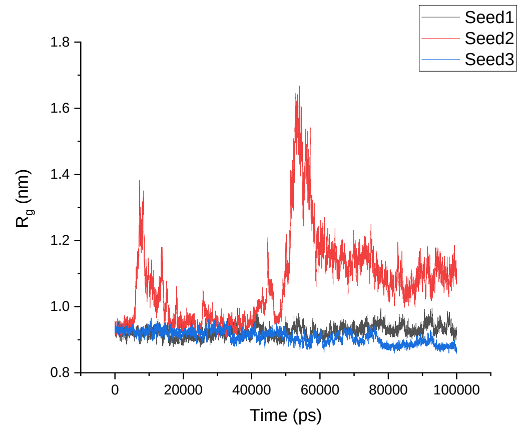


CHARMM22*

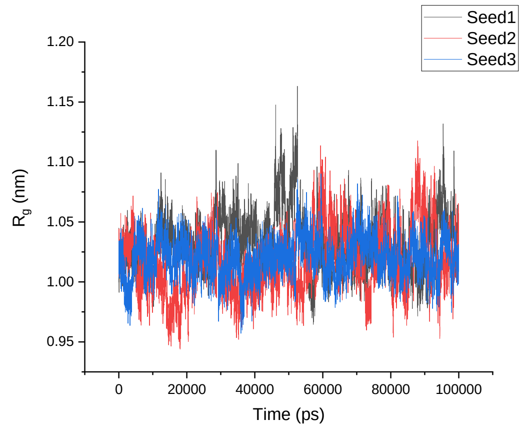


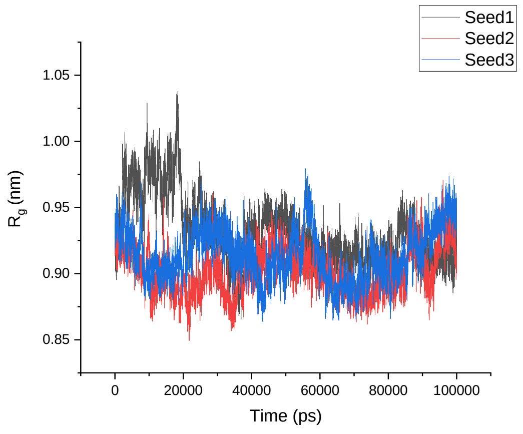


CHARMM36

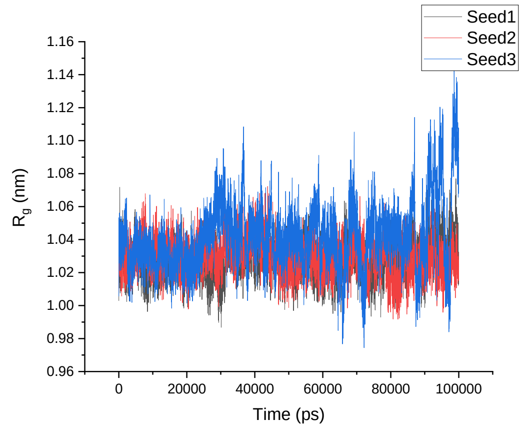


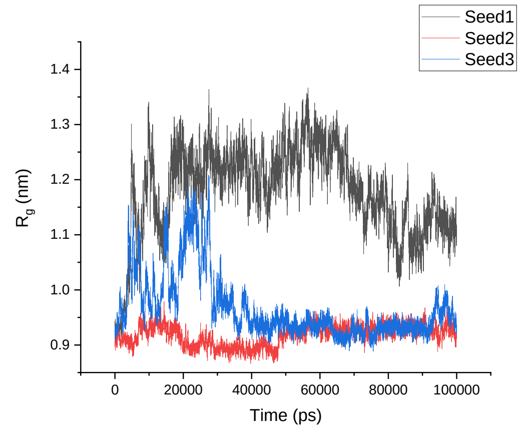

<table>
<tr><td>GROMOS 54A8</td><td>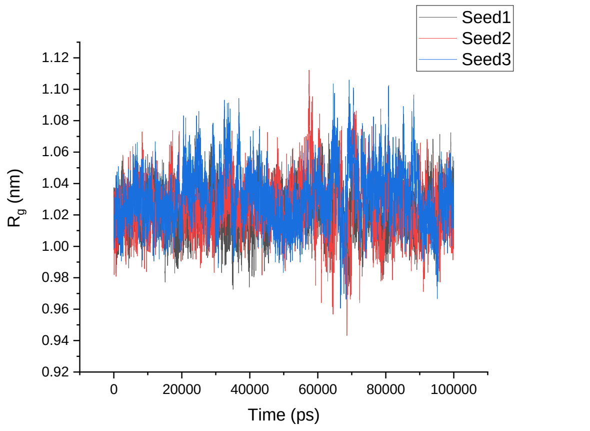
</td><td>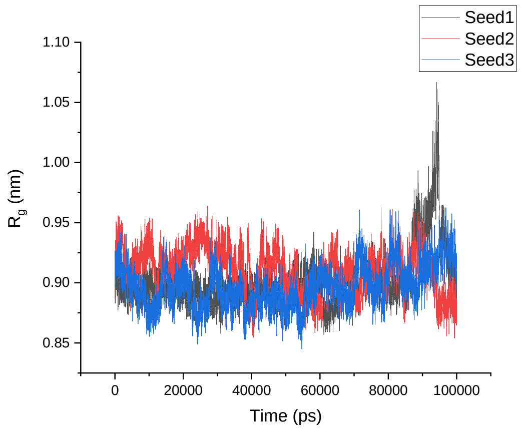
</td></tr>
<tr><td></td><td>1LFC</td><td>α-1LFC</td></tr>
<tr><td>Amber 99SB-ILDN</td><td>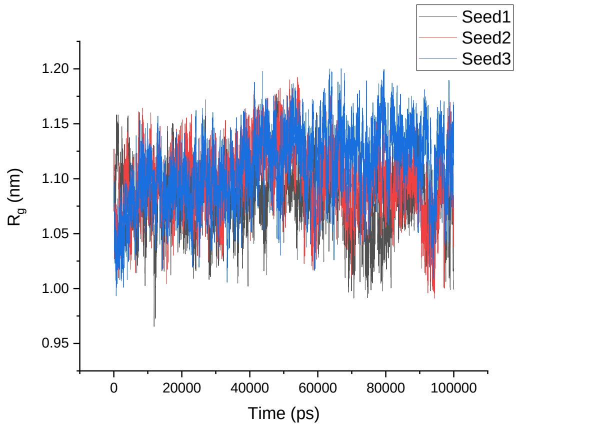
</td><td>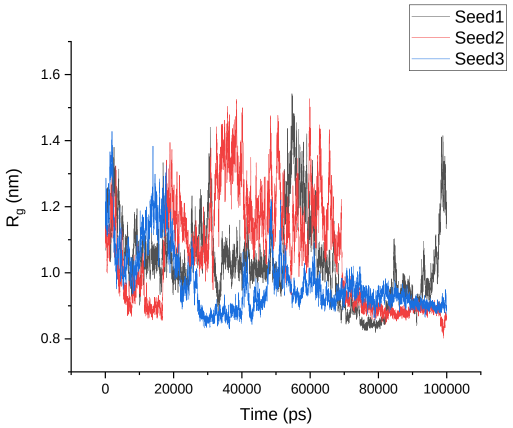
</td></tr>
<tr><td>Amber 99SB*-ILDN</td><td>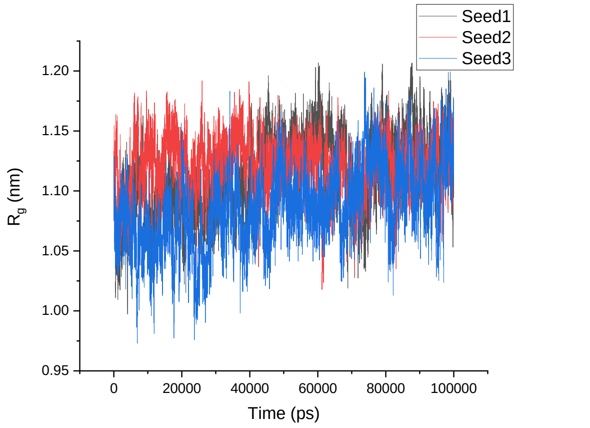
</td><td>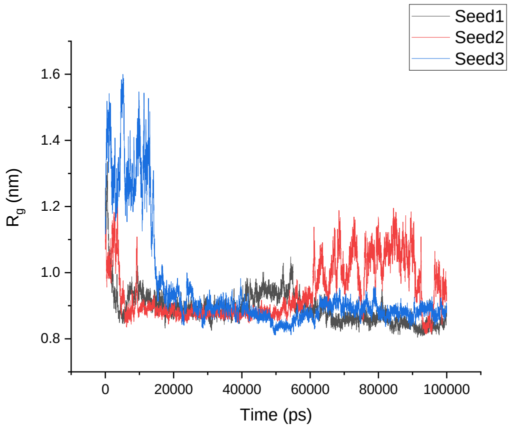
</td></tr>
</table>

| | | |
|---|---|---|
| CHARMM22* | 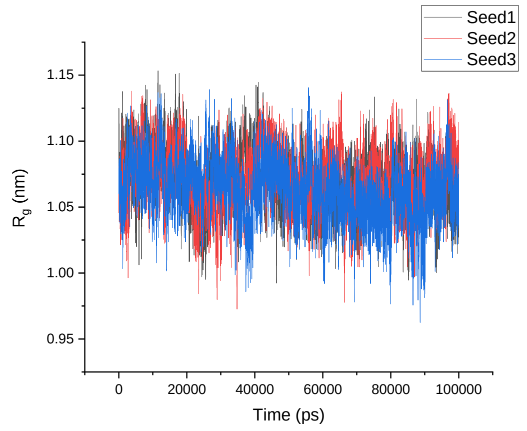 | 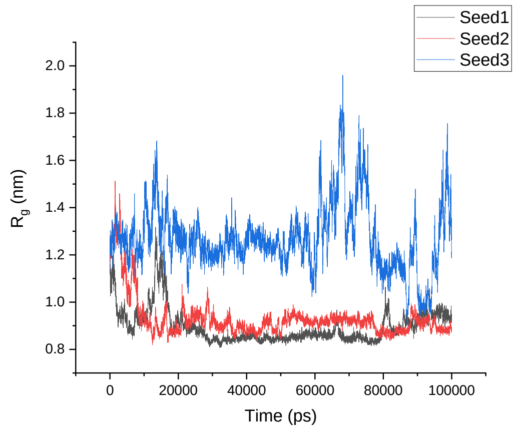 |
| CHARMM36 | 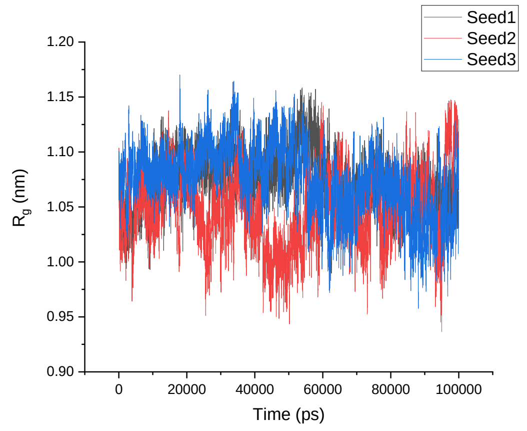 | 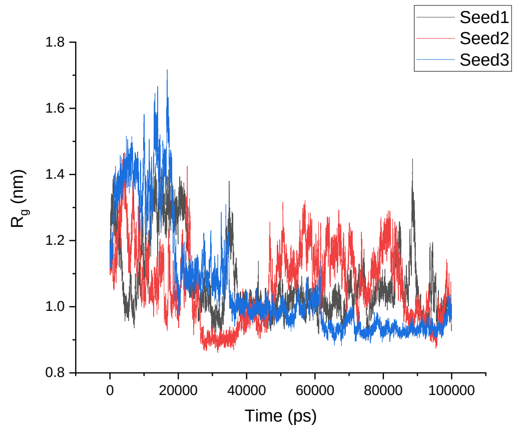 |
| GROMOS 54A8 | 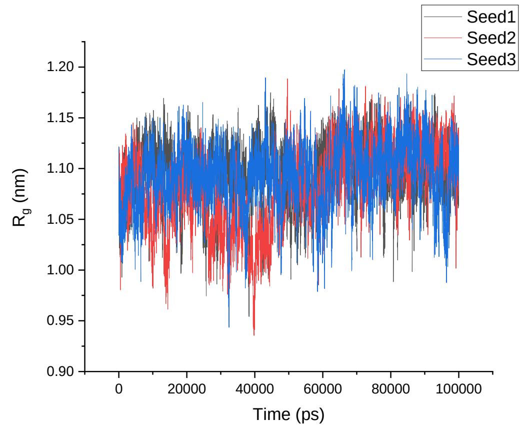 | 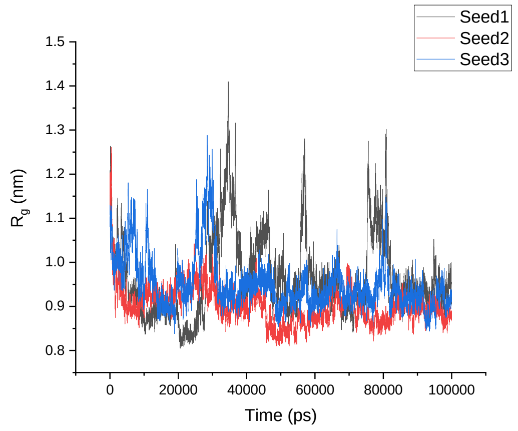 |
| | **8TFV** | **α-8TFV** |

<table>
<tr><td>Amber 99SB-ILDN</td><td>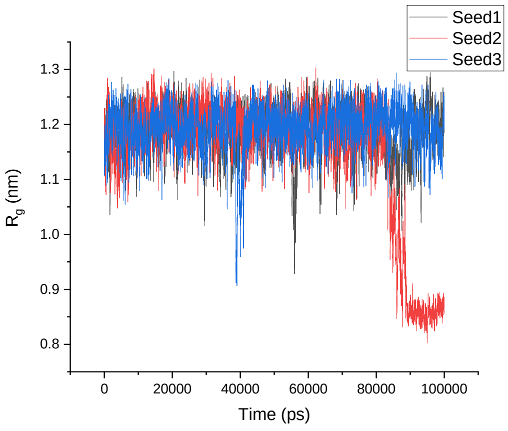
</td><td>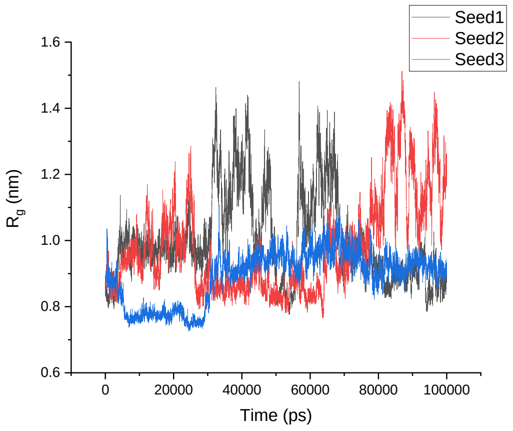
</td></tr>
<tr><td>Amber 99SB*-ILDN</td><td>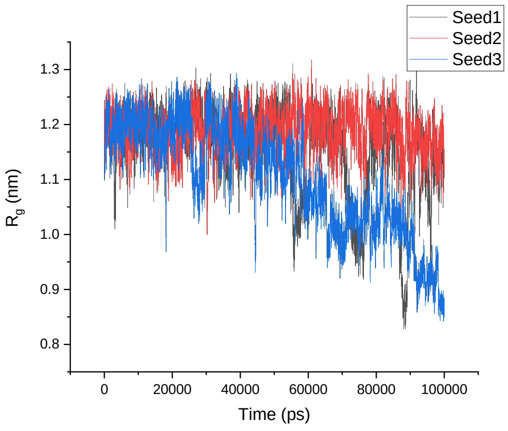
</td><td>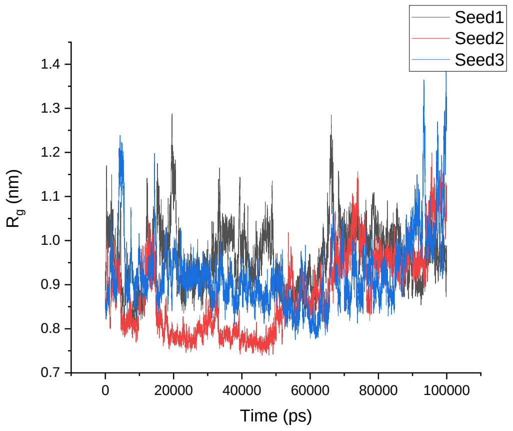
</td></tr>
<tr><td>CHARMM22*</td><td>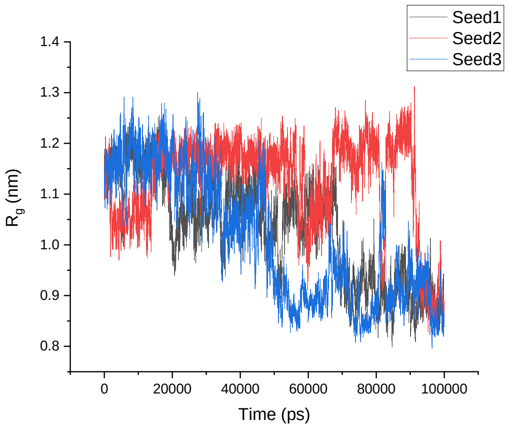
</td><td>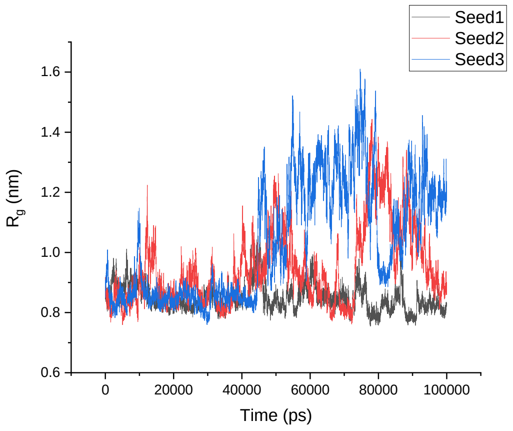
</td></tr>
</table>

<table>
<tr><td>CHARMM36</td><td>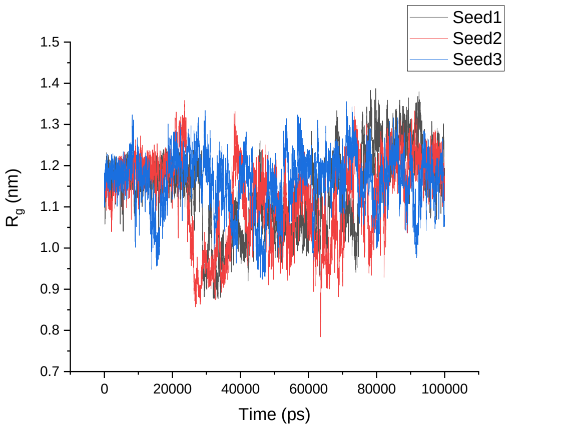
</td><td>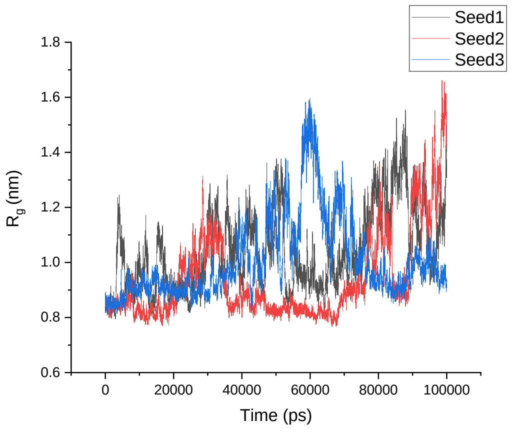
</td></tr>
<tr><td>GROMOS 54A8</td><td>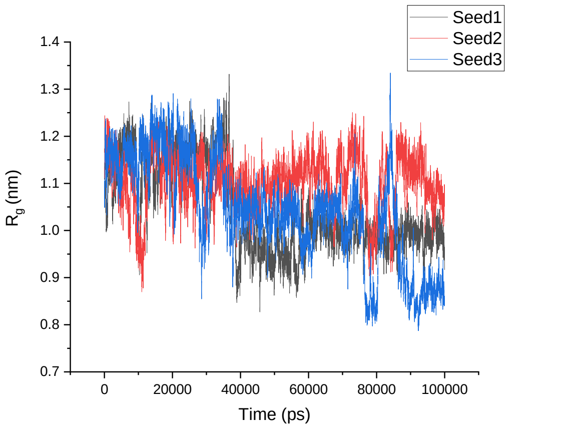
</td><td>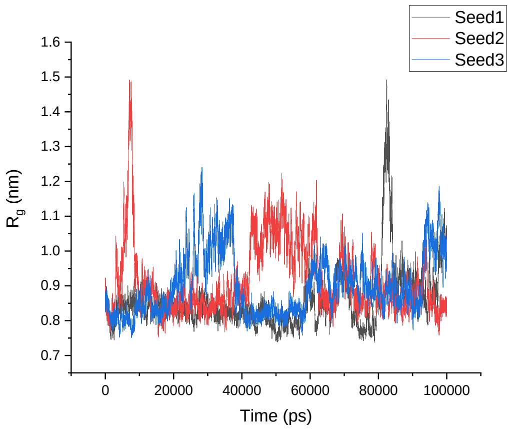
</td></tr>
<tr><td></td><td>1KGM</td><td>α-1KGM</td></tr>
<tr><td>Amber 99SB-ILDN</td><td>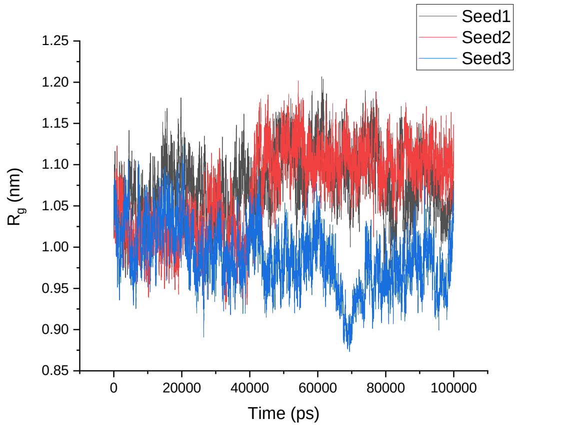
</td><td>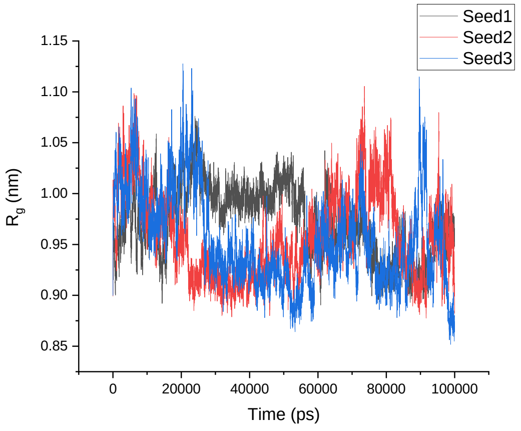
</td></tr>
</table>

<table>
<tr><td>Amber 99SB*-ILDN</td><td>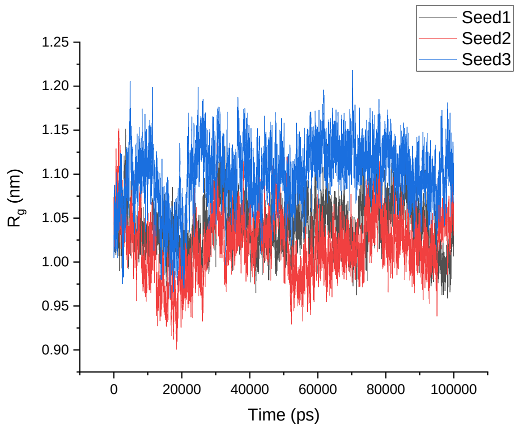
</td><td>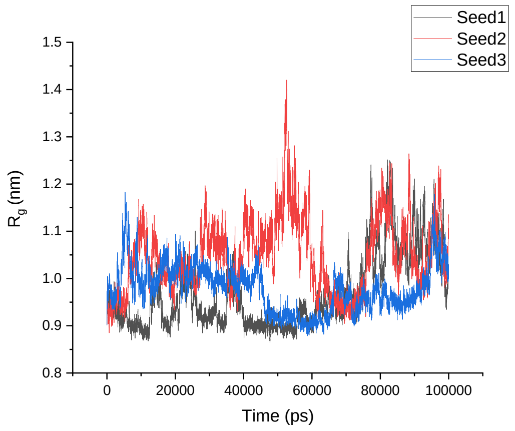
</td></tr>
<tr><td>CHARMM22*</td><td>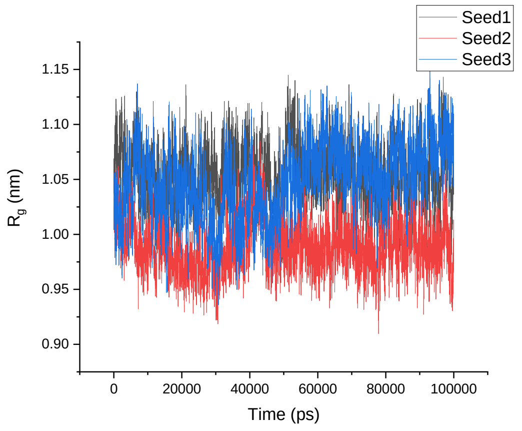
</td><td>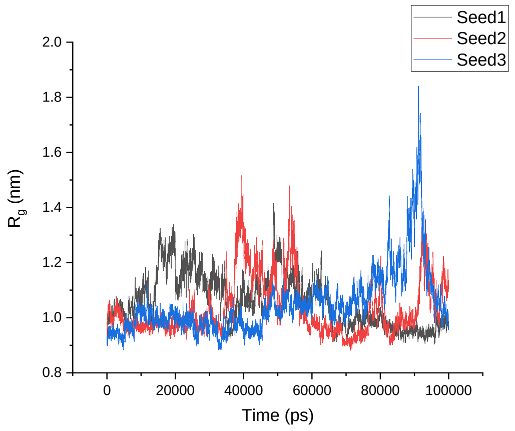
</td></tr>
<tr><td>CHARMM36</td><td>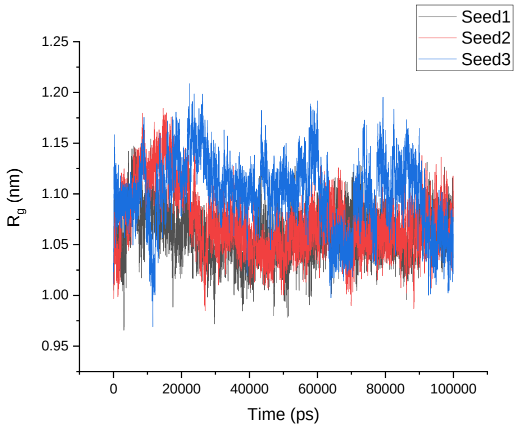
</td><td>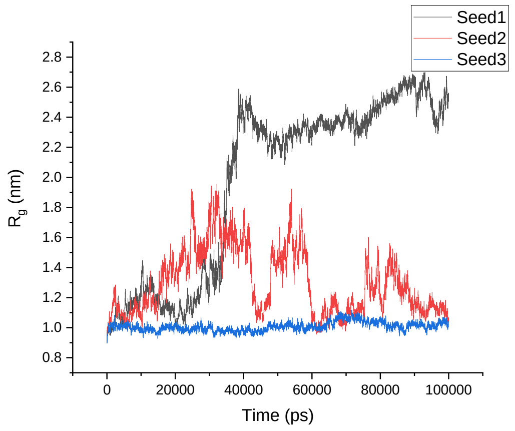
</td></tr>
</table>

<table>
<tr><td>GROMOS 54A8</td><td>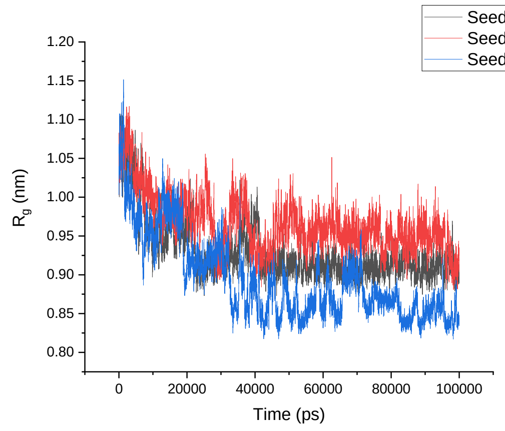
</td><td>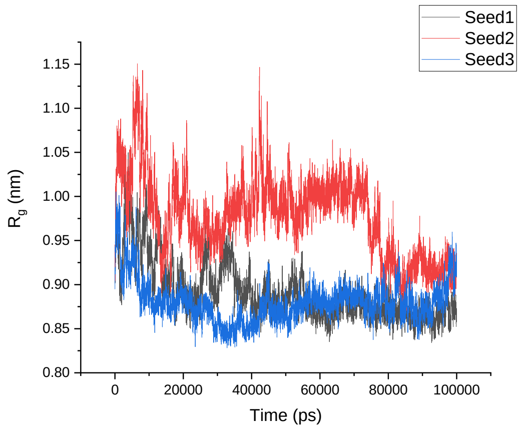
</td></tr>
<tr><td></td><td>1M4F</td><td>α-1M4F</td></tr>
<tr><td>Amber 99SB-ILDN</td><td>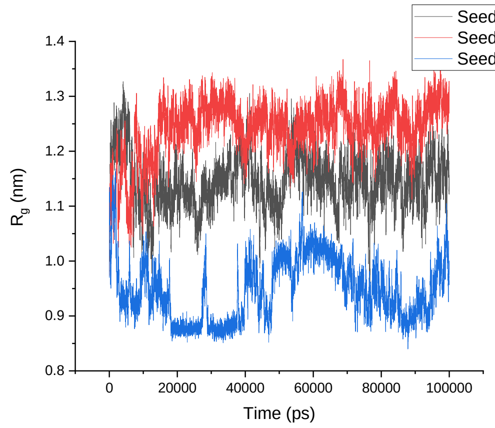
</td><td>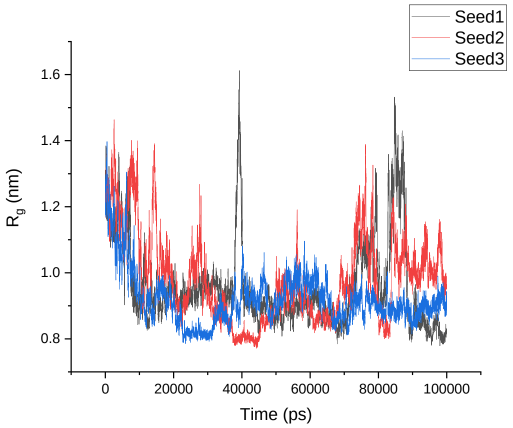
</td></tr>
<tr><td>Amber 99SB*-ILDN</td><td>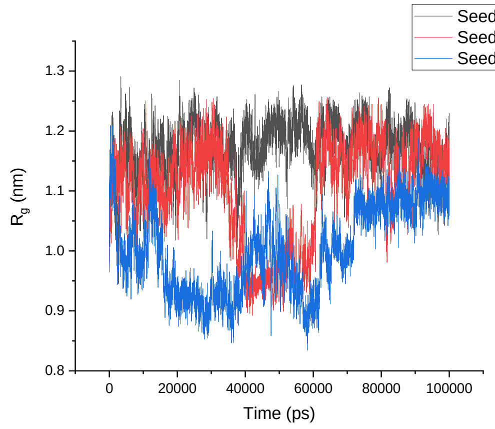
</td><td>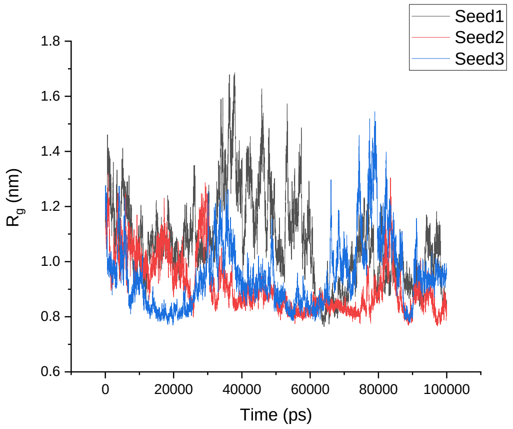
</td></tr>
</table>

<table>
<tr><td>CHARMM22*</td><td>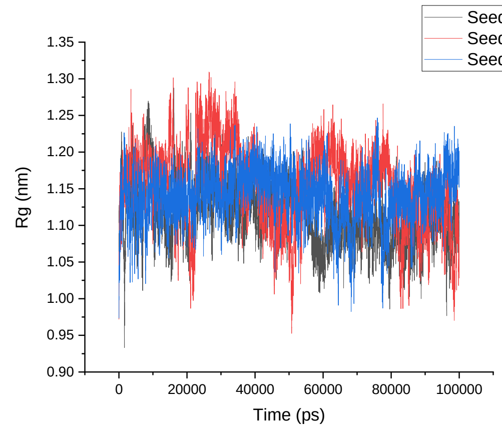
</td><td>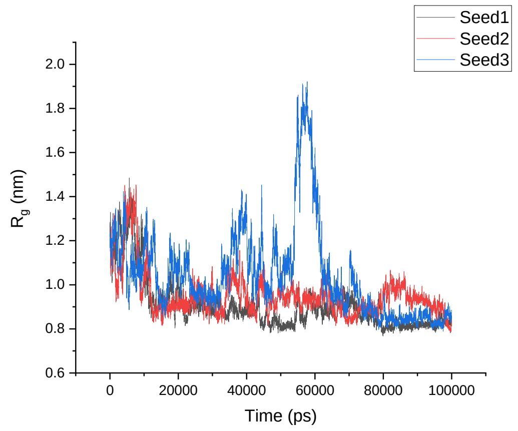
</td></tr>
<tr><td>CHARMM36</td><td>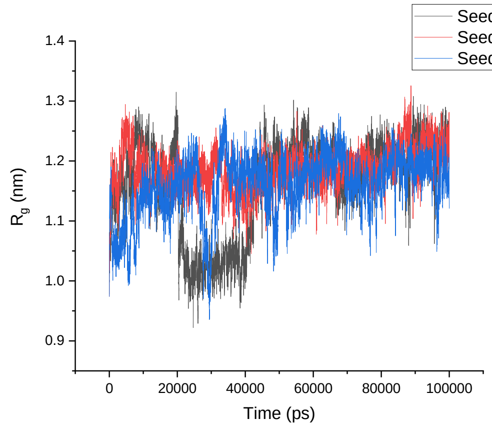
</td><td>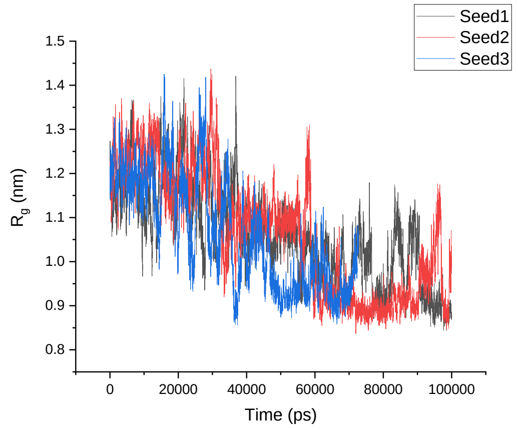
</td></tr>
<tr><td>GROMOS 54A8</td><td>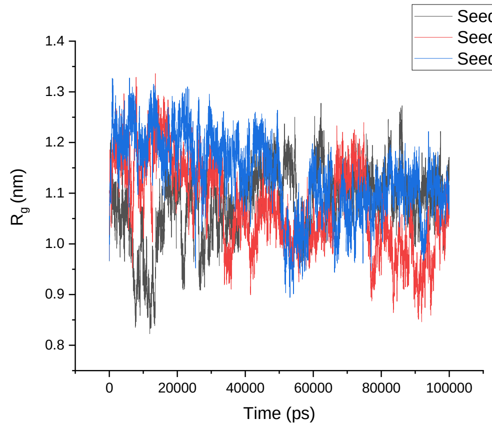
</td><td>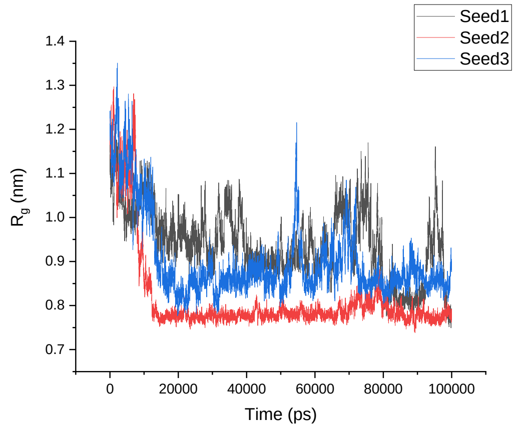
</td></tr>
<tr><td></td><td>1TV0</td><td>α-1TV0</td></tr>
</table>

| | | |
|---|---|---|
| Amber 99SB-ILDN | 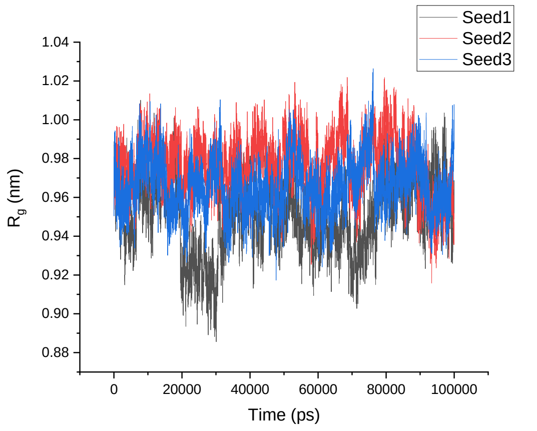 | 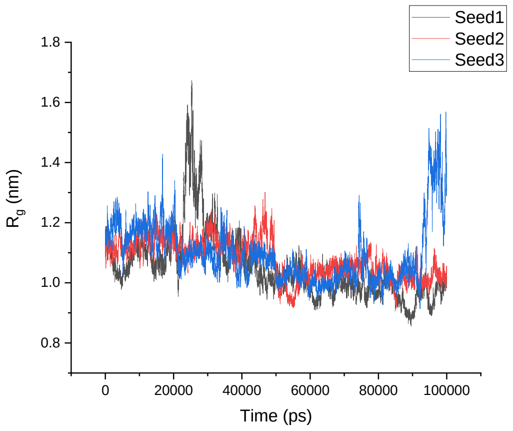 |
| Amber 99SB*-ILDN | 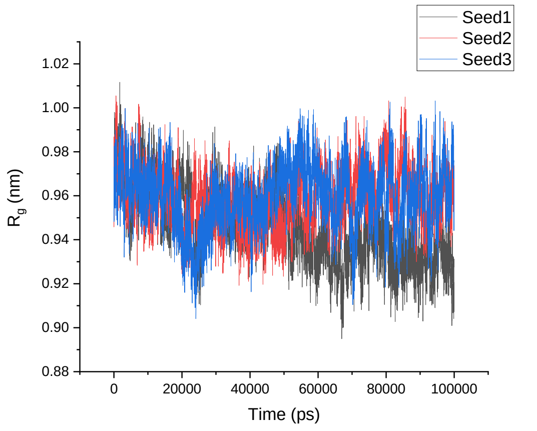 | 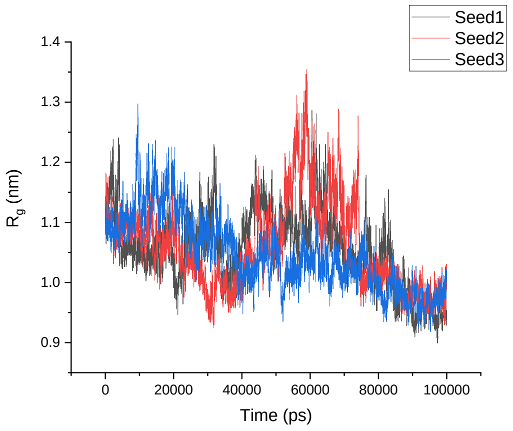 |
| CHARMM22* | 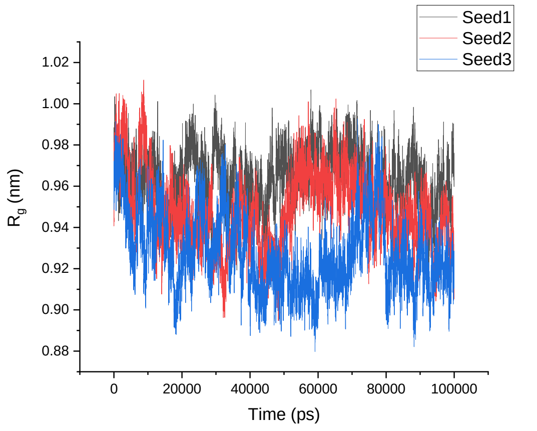 | 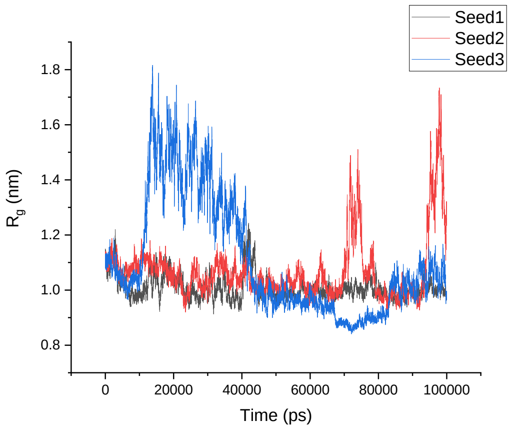 |

| | | |
|---|---|---|
| CHARMM36 | 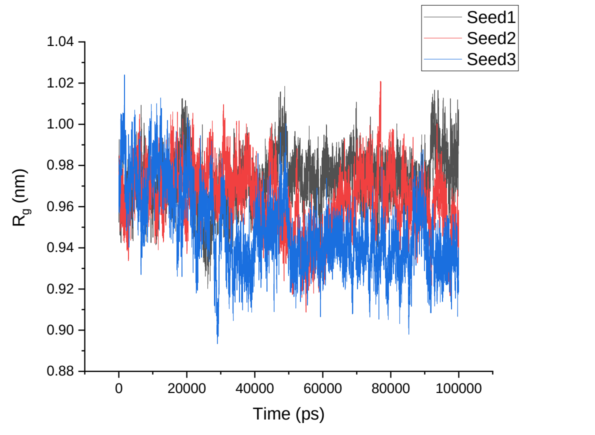 | 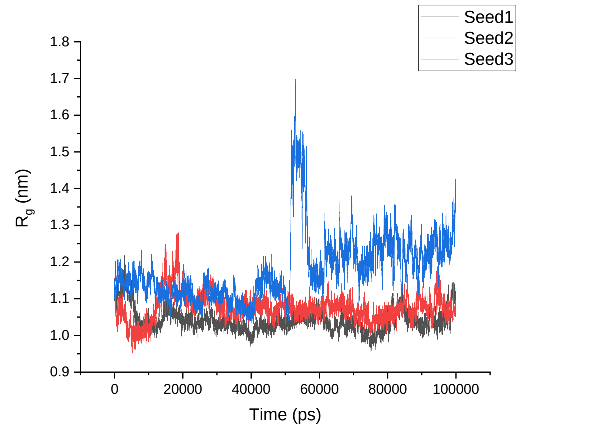 |
| GROMOS 54A8 | 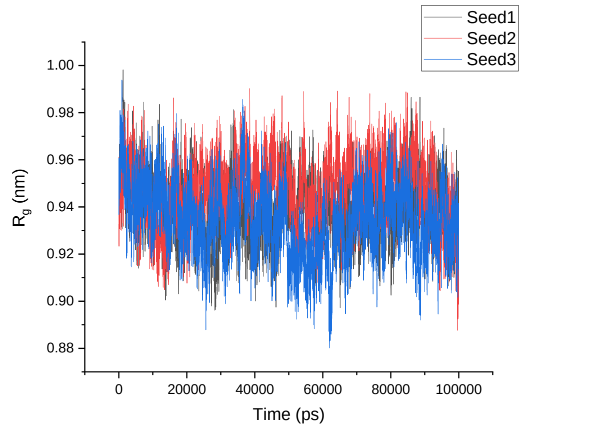 | 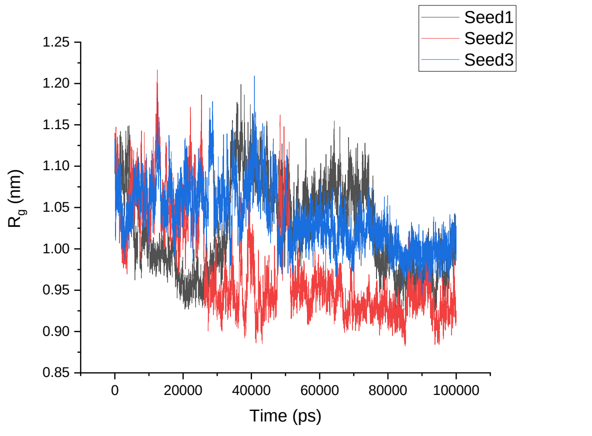 |

**Table S8**. Average $R_g$ over three simulations of the native α-proteins, modeled β-proteins, β-proteins, and modeled α-proteins.

| | $R_g$ | SD | $R_g$ | SD | $R_g$ | SD | $R_g$ | SD | $R_g$ | SD |
|---|---|---|---|---|---|---|---|---|---|---|
| α- proteins | | | | | | | | | | |
| | **1CQ0** | | **1L2Y** | | **1ROO** | | **1T5Q** | | **1V1D** | |
| Amber 99SB-ILDN | 0.92 | 0.05 | 0.71 | 0.01 | 0.92 | 0.02 | 1.02 | 0.12 | 1.03 | 0.02 |
| Amber 99SB*-ILDN | 0.94 | 0.08 | 0.71 | 0.01 | 0.90 | 0.01 | 1.01 | 0.14 | 1.02 | 0.02 |
| CHARMM22* | 0.93 | 0.06 | 0.74 | 0.01 | 0.92 | 0.01 | 1.03 | 0.15 | 1.02 | 0.02 |
| CHARMM36 | 1.05 | 0.09 | 0.73 | 0.01 | 0.92 | 0.01 | 1.36 | 0.06 | 1.03 | 0.01 |
| GROMOS 54A8 | 0.92 | 0.07 | 0.75 | 0.03 | 0.92 | 0.02 | 1.13 | 0.15 | 1.03 | 0.02 |
| Modelled β-proteins | | | | | | | | | | |
| | **β-1CQ0** | | **β-1L2Y** | | **β-1ROO** | | **β-1T5Q** | | **β-1V1D** | |
| Amber 99SB-ILDN | 0.99 | 0.11 | 0.87 | 0.10 | 0.99 | 0.05 | 1.12 | 0.06 | 0.92 | 0.03 |
| Amber 99SB*-ILDN | 1.06 | 0.08 | 0.88 | 0.15 | 1.02 | 0.06 | 1.12 | 0.06 | 0.97 | 0.06 |
| CHARMM22* | 1.01 | 0.13 | 0.91 | 0.12 | 1.06 | 0.09 | 1.06 | 0.07 | 0.92 | 0.02 |
| CHARMM36 | 1.09 | 0.12 | 0.98 | 0.14 | 1.20 | 0.11 | 1.16 | 0.08 | 1.02 | 0.05 |
| GROMOS 54A8 | 0.94 | 0.10 | 0.82 | 0.06 | 0.97 | 0.04 | 1.01 | 0.06 | 0.90 | 0.02 |
| β- proteins | | | | | | | | | | |
| | **1LFC** | | **8TFV** | | **1KGM** | | **1M4F** | | **1TV0** | |
| Amber 99SB-ILDN | 1.10 | 0.03 | 1.18 | 0.07 | 1.04 | 0.04 | 1.11 | 0.05 | 0.96 | 0.02 |
| Amber 99SB*-ILDN | 1.11 | 0.03 | 1.15 | 0.07 | 1.05 | 0.03 | 1.09 | 0.07 | 0.95 | 0.01 |
| CHARMM22* | 1.07 | 0.02 | 1.06 | 0.11 | 1.03 | 0.03 | 1.14 | 0.04 | 0.95 | 0.02 |
| CHARMM36 | 1.07 | 0.03 | 1.14 | 0.09 | 1.08 | 0.03 | 1.17 | 0.05 | 0.96 | 0.02 |
| GROMOS 54A8 | 1.09 | 0.03 | 1.06 | 0.09 | 0.93 | 0.04 | 1.10 | 0.08 | 0.94 | 0.01 |
| Modelled α- proteins | | | | | | | | | | |
| | **α-1LFC** | | **α-8TFV** | | **α-1KGM** | | **α-1M4F** | | **α-1TV0** | |
| Amber 99SB-ILDN | 1.02 | 0.13 | 0.96 | 0.12 | 0.96 | 0.04 | 0.95 | 0.11 | 1.07 | 0.09 |
| Amber 99SB*-ILDN | 0.93 | 0.10 | 0.92 | 0.08 | 1.00 | 0.07 | 0.98 | 0.13 | 1.05 | 0.07 |
| CHARMM22* | 1.03 | 0.10 | 0.94 | 0.12 | 1.05 | 0.10 | 0.97 | 0.14 | 1.06 | 0.12 |
| CHARMM36 | 1.07 | 0.13 | 1.00 | 0.15 | 1.43 | 0.28 | 1.06 | 0.12 | 1.10 | 0.05 |
| GROMOS 54A8 | 0.94 | 0.06 | 0.89 | 0.09 | 0.92 | 0.03 | 0.89 | 0.09 | 1.01 | 0.05 |

**Table S9**. SASA plots of the native α-proteins, modeled β-proteins, β-proteins, and modeled α-proteins.

<table>
<tr><th colspan="3">SASA</th></tr>
<tr><th></th><th>1CQ0</th><th>β-1CQ0</th></tr>
<tr><td>Amber 99SB-ILDN</td><td>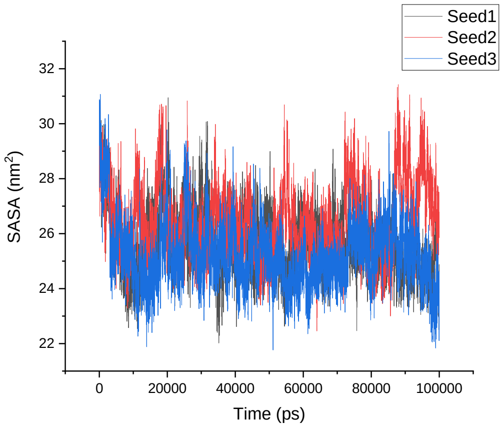
</td><td>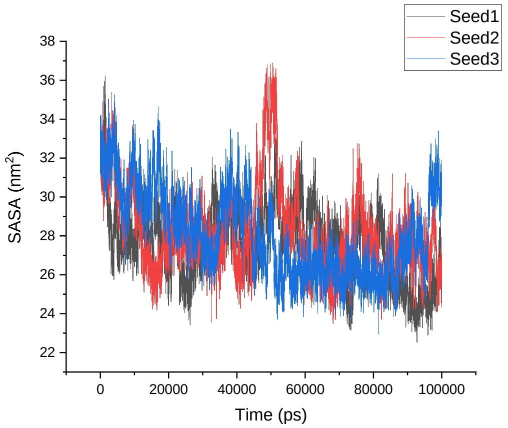
</td></tr>
<tr><td>Amber 99SB*-ILDN</td><td>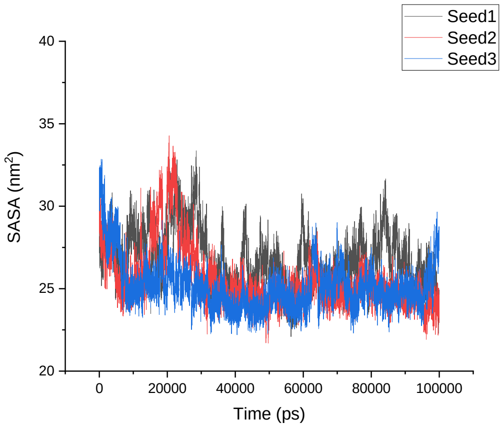
</td><td>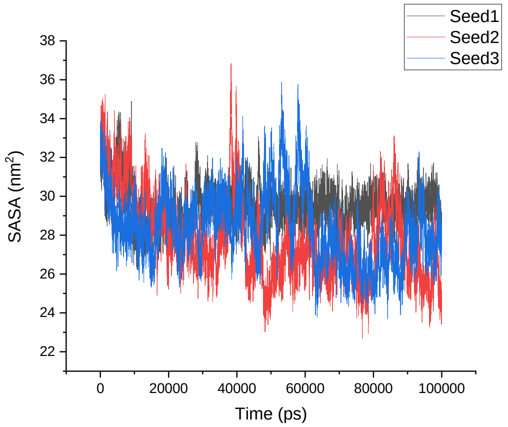
</td></tr>
<tr><td>CHARMM22*</td><td>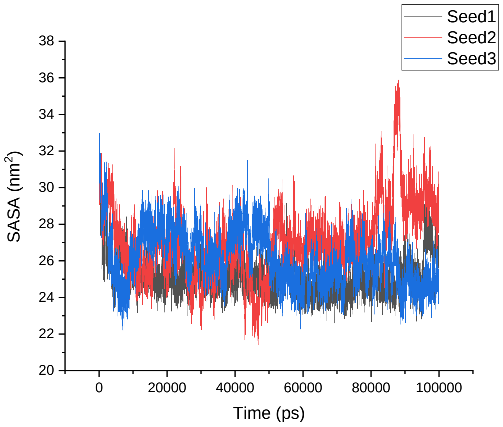
</td><td>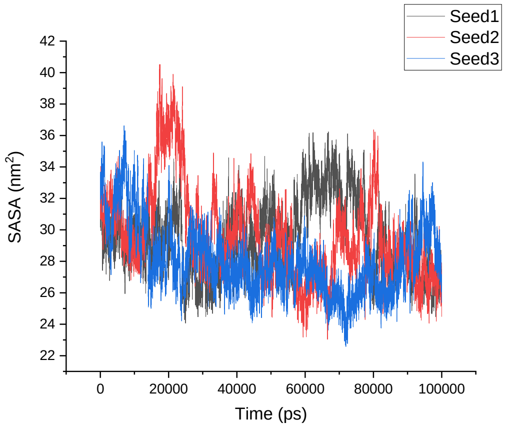
</td></tr>
</table>

| | | |
|---|---|---|
| CHARMM36 | 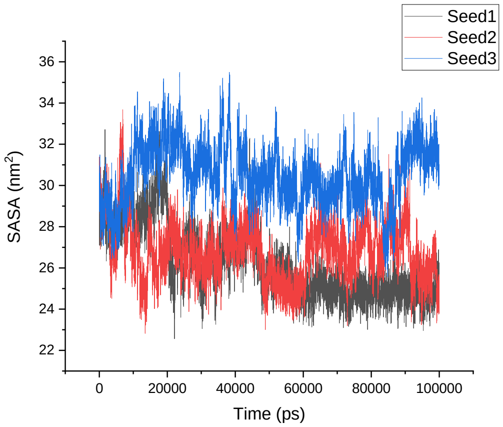 | 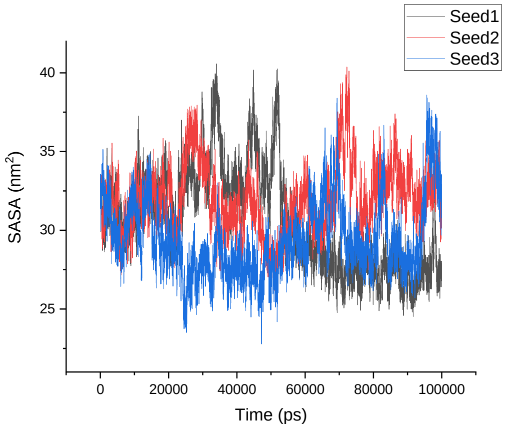 |
| GROMOS 54A8 | 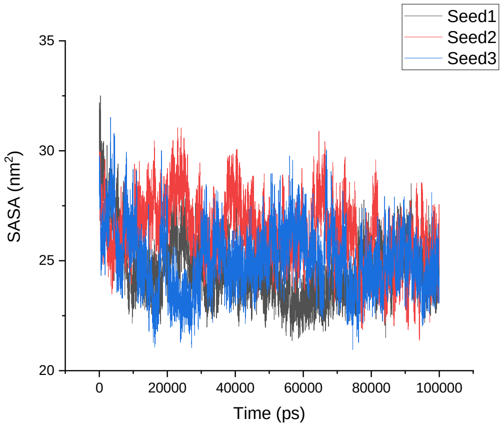 | 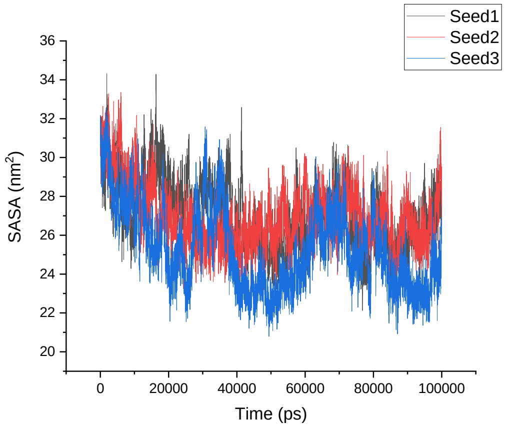 |
| | **1L2Y** | **β-1L2Y** |
| Amber 99SB-ILDN | 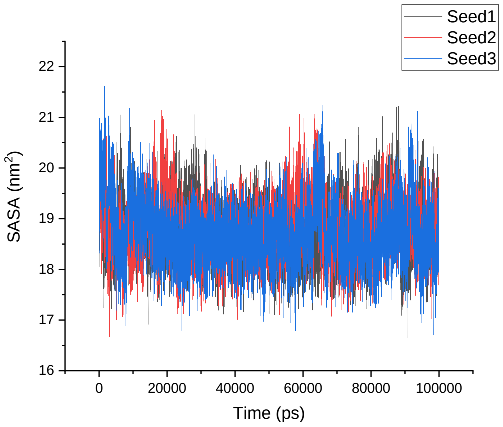 | 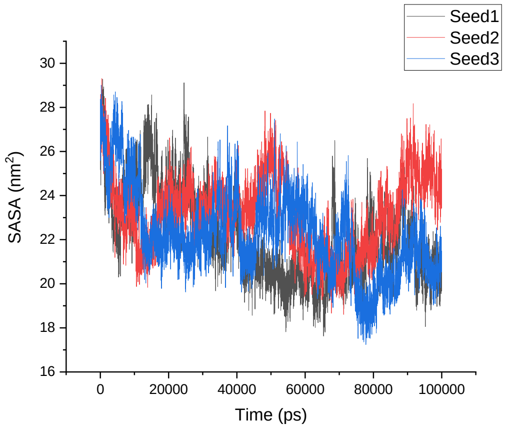 |

| | | |
|---|---|---|
| Amber 99SB*-ILDN | 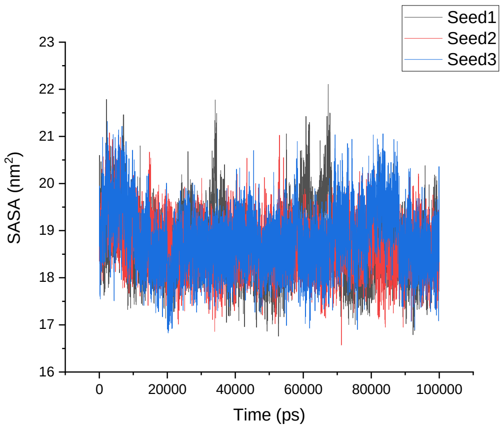 | 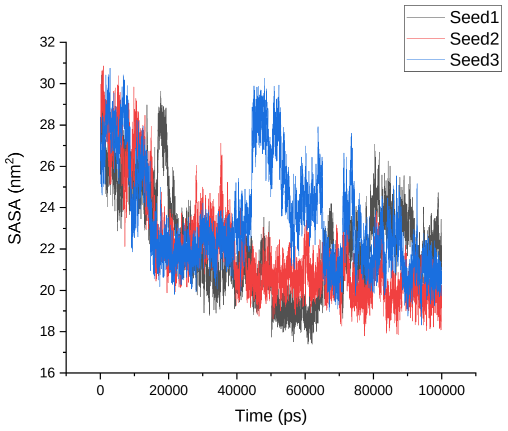 |
| CHARMM22* | 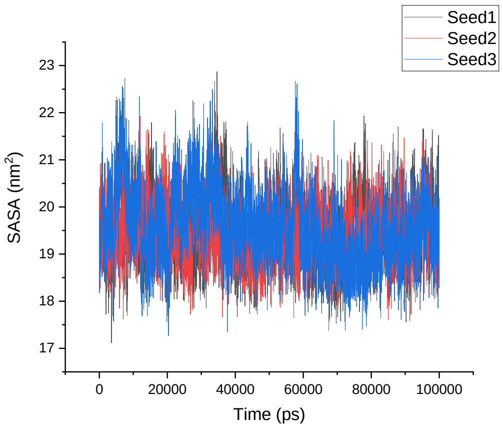 | 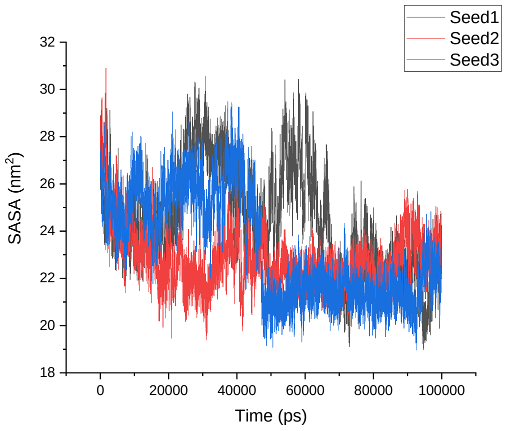 |
| CHARMM36 | 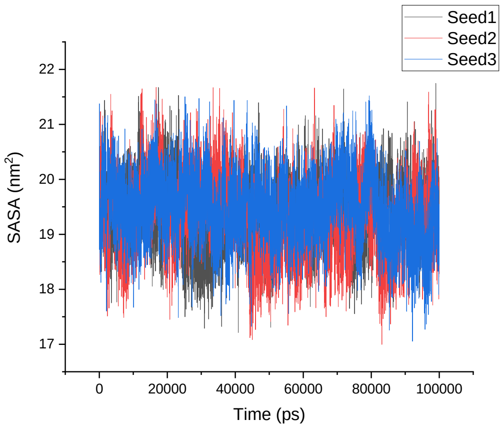 | 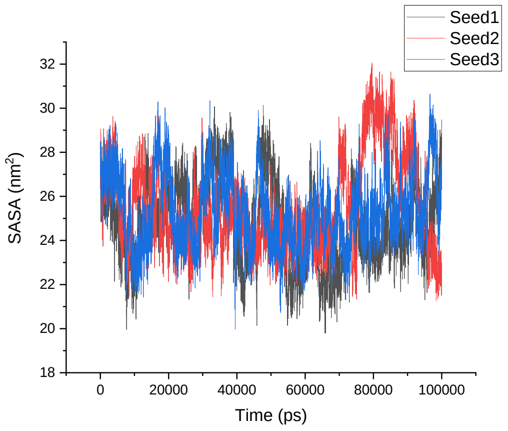 |

<table>
<tr><td>GROMOS 54A8</td><td>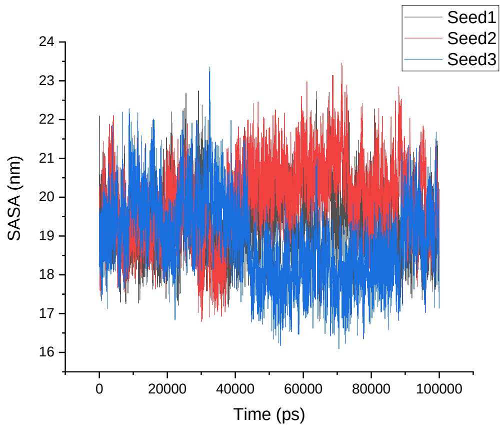
</td><td>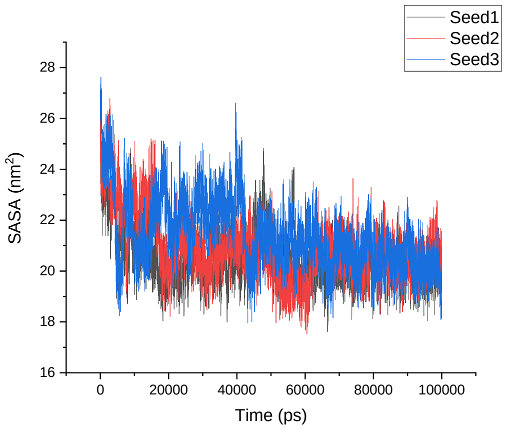
</td></tr>
<tr><td></td><td>1ROO</td><td>β-1ROO</td></tr>
<tr><td>Amber 99SB-ILDN</td><td>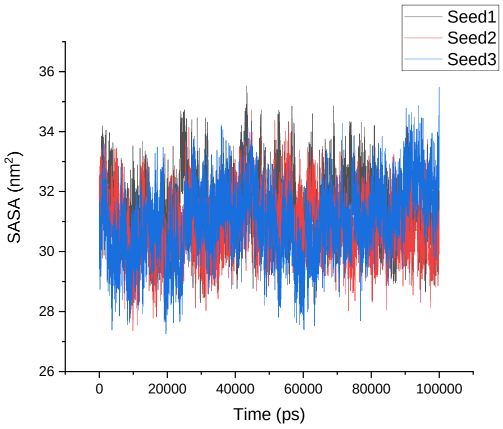
</td><td>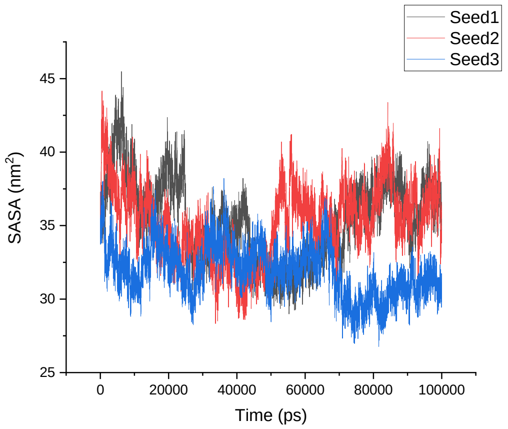
</td></tr>
<tr><td>Amber 99SB*-ILDN</td><td>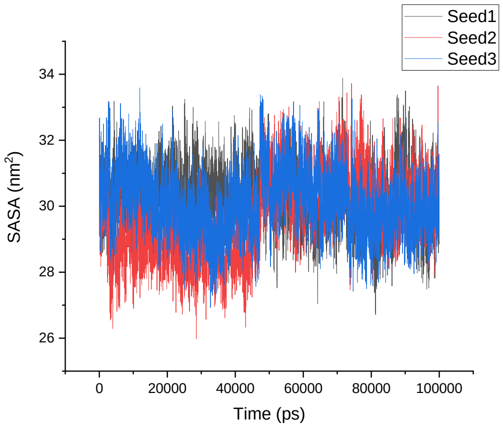
</td><td>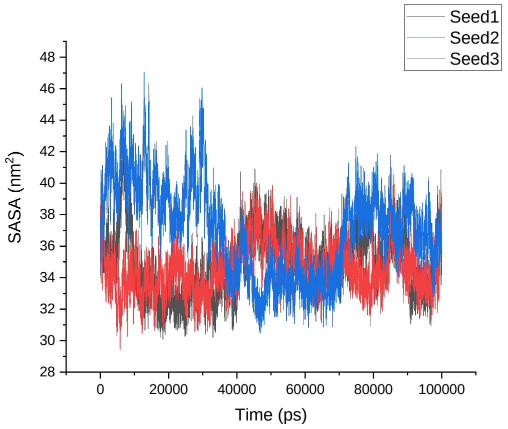
</td></tr>
</table>

<table>
<tr><td>CHARMM22*</td><td>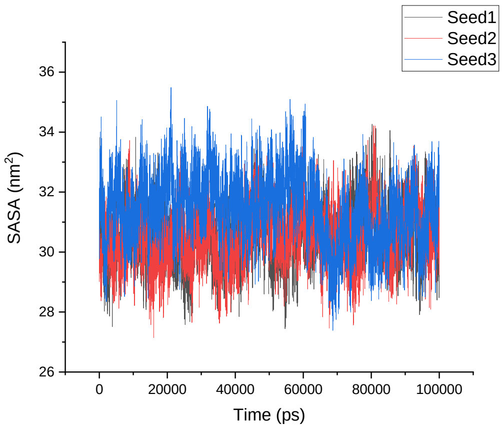
</td><td>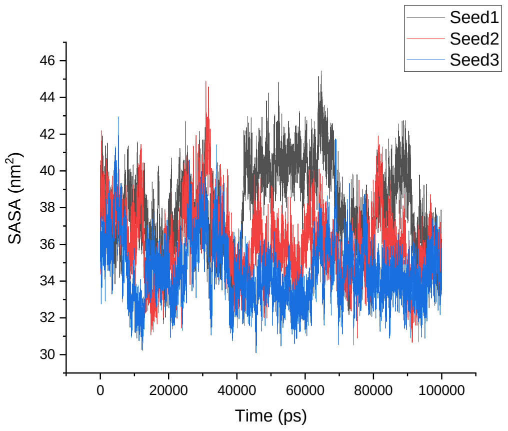
</td></tr>
<tr><td>CHARMM36</td><td>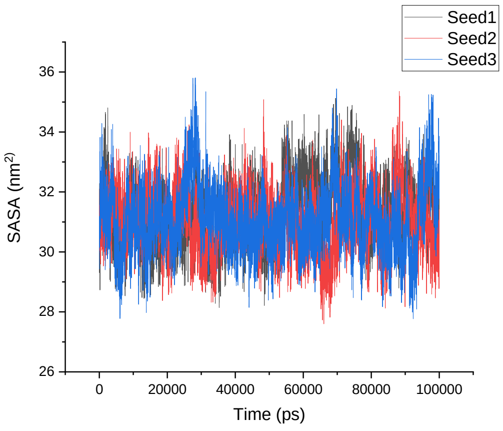
</td><td>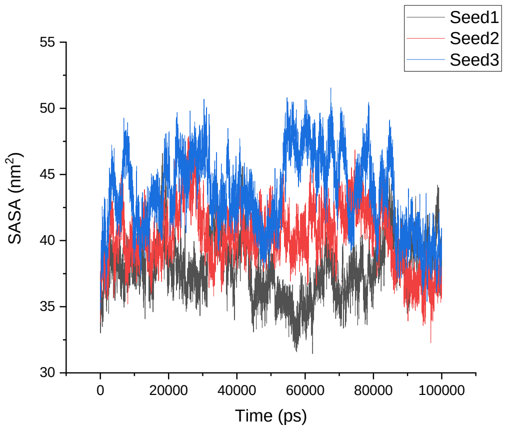
</td></tr>
<tr><td>GROMOS 54A8</td><td>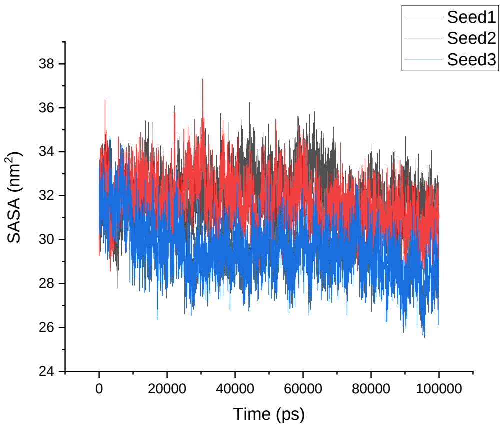
</td><td>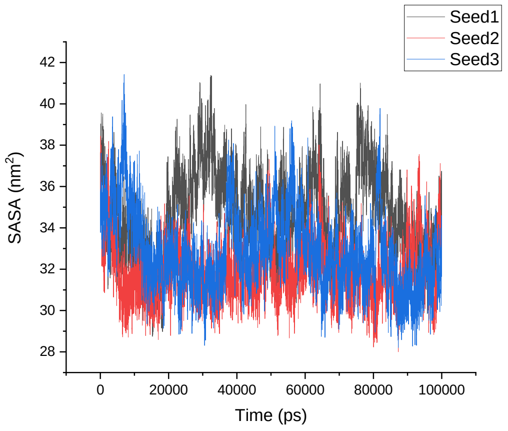
</td></tr>
<tr><td></td><td>1T5Q</td><td>β-1T5Q</td></tr>
</table>

<table>
<tr>
<td>Amber 99SB-ILDN</td>
<td>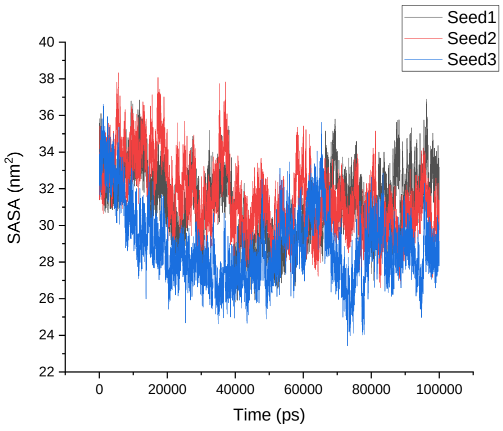
</td>
<td>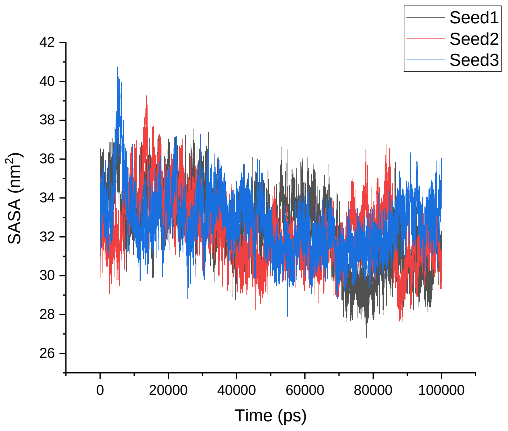
</td>
</tr>
<tr>
<td>Amber 99SB*-ILDN</td>
<td>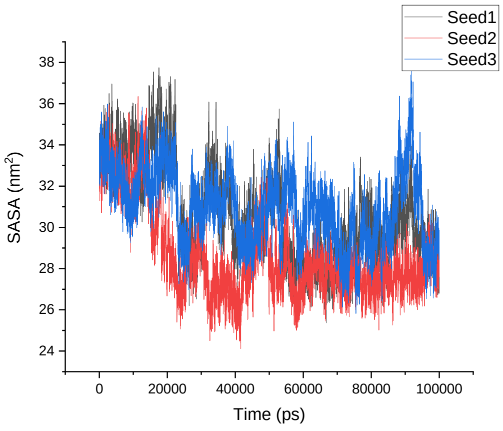
</td>
<td>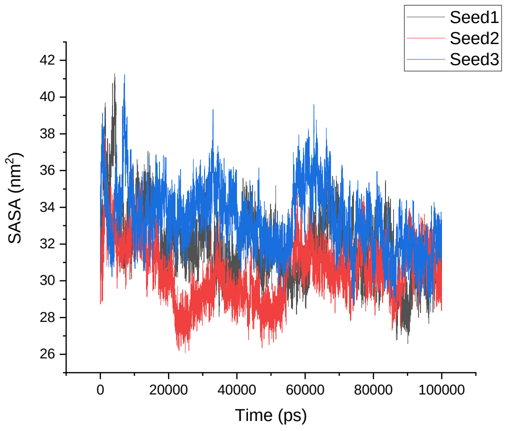
</td>
</tr>
<tr>
<td>CHARMM22*</td>
<td>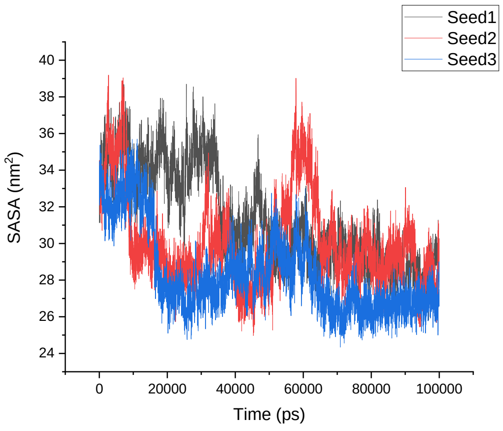
</td>
<td>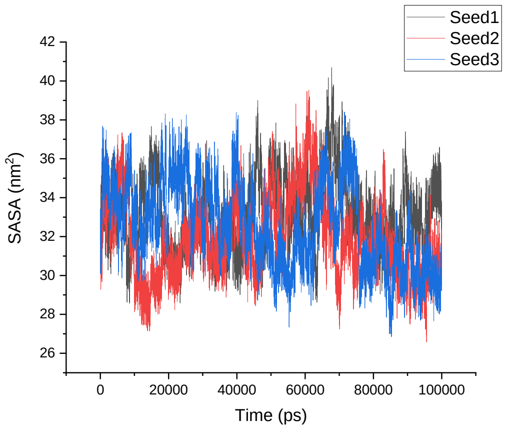
</td>
</tr>
</table>

<table>
<tr><td>CHARMM36</td><td>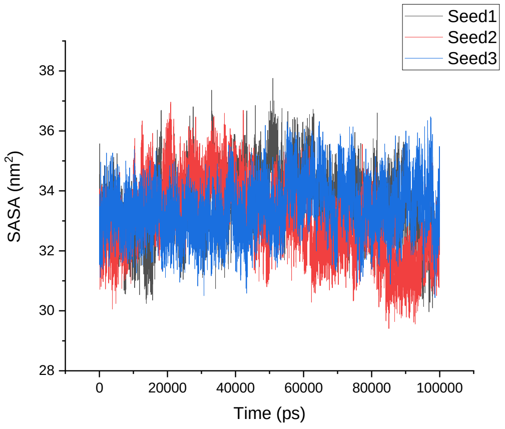</td><td>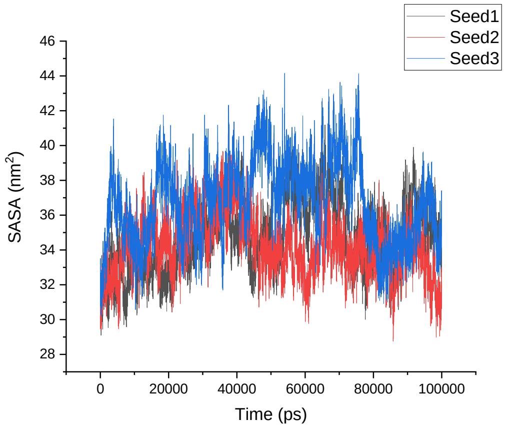</td></tr>
<tr><td>GROMOS 54A8</td><td>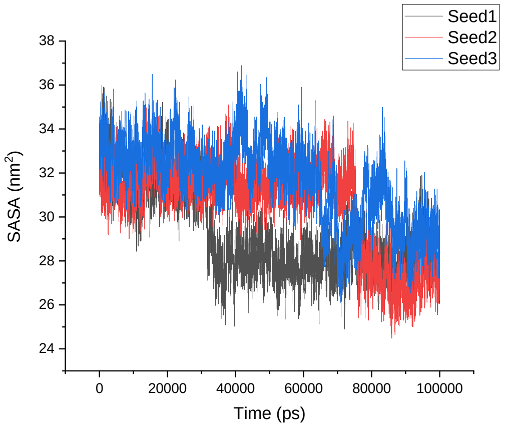</td><td>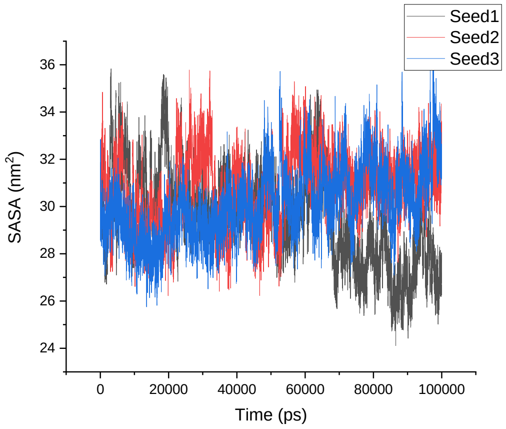</td></tr>
<tr><td></td><td>1V1D</td><td>β-1V1D</td></tr>
<tr><td>Amber 99SB-ILDN</td><td>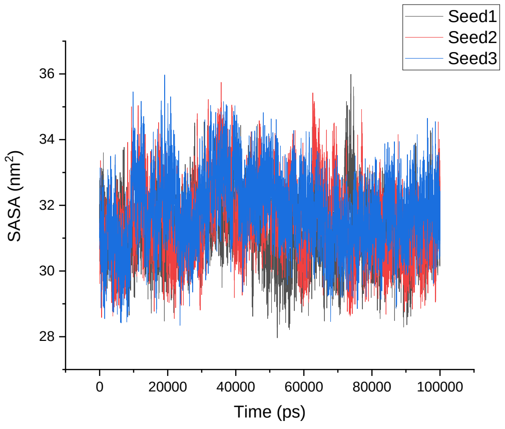</td><td>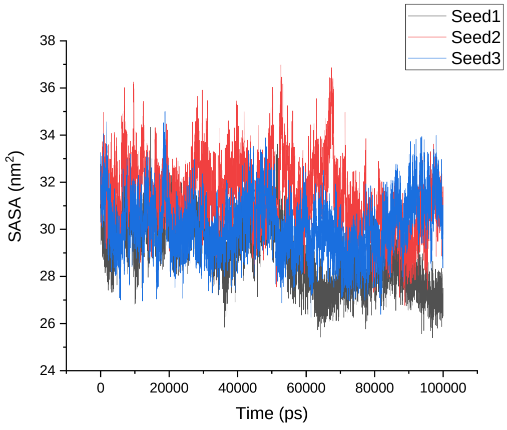</td></tr>
</table>

<table>
<tr>
<td>Amber 99SB*-ILDN</td>
<td>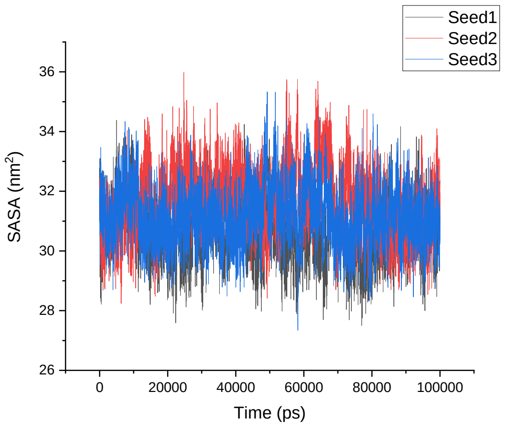
</td>
<td>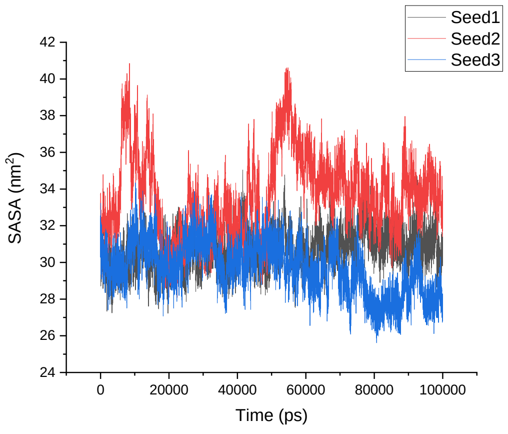
</td>
</tr>
<tr>
<td>CHARMM22*</td>
<td>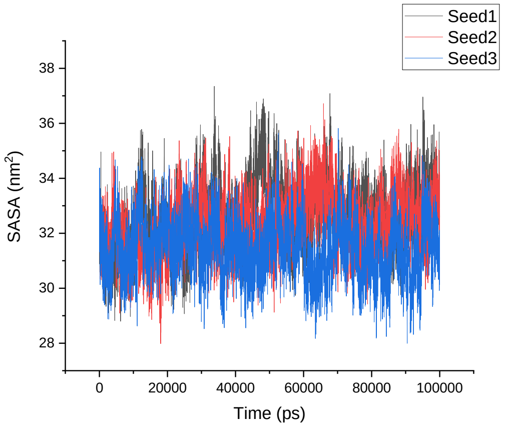
</td>
<td>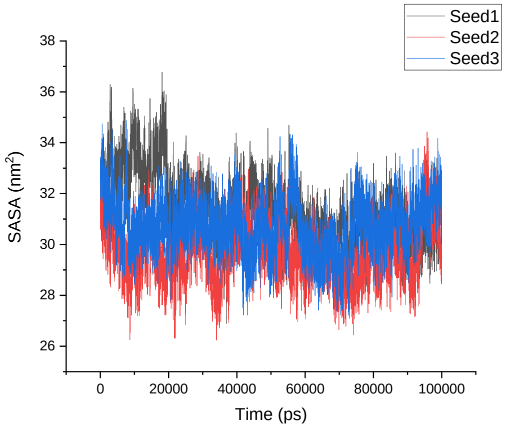
</td>
</tr>
<tr>
<td>CHARMM36</td>
<td>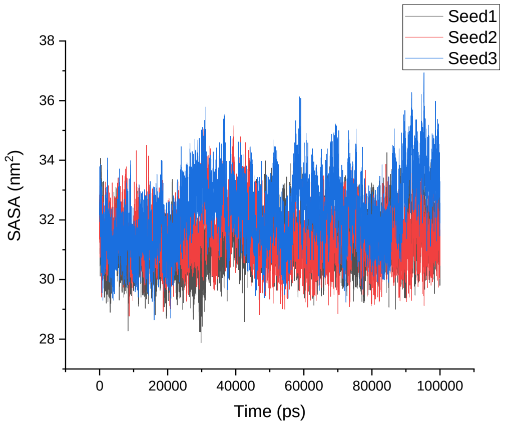
</td>
<td>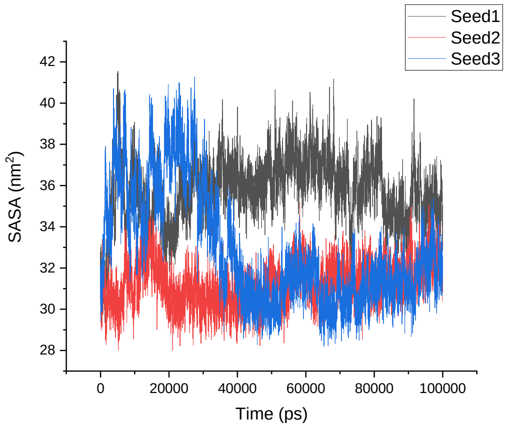
</td>
</tr>
</table>

| | | |
|---|---|---|
| GROMOS 54A8 | 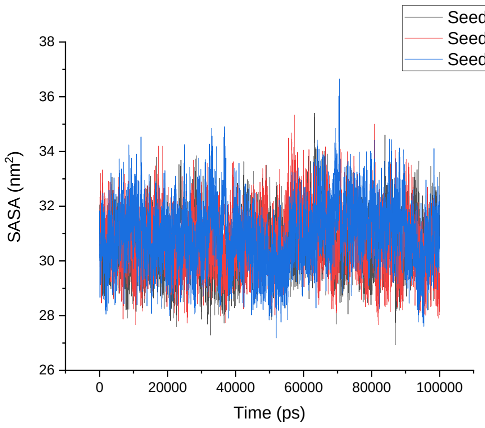 | 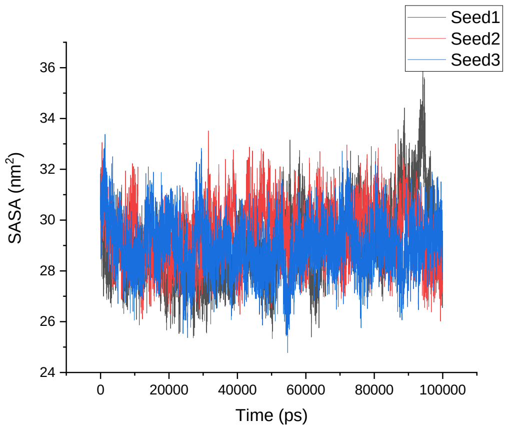 |
| | **1LFC** | **α-1LFC** |
| Amber 99SB-ILDN | 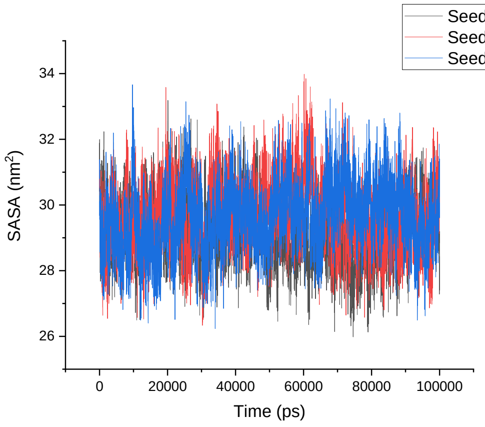 | 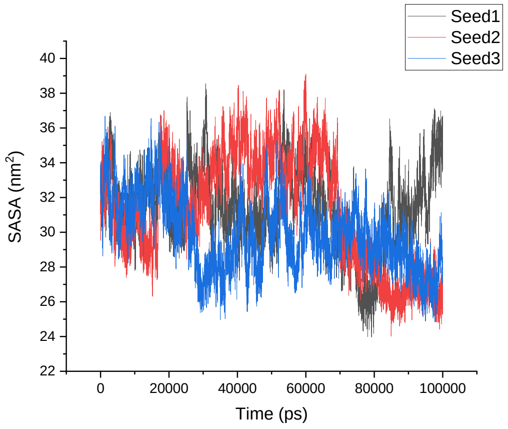 |
| Amber 99SB*-ILDN | 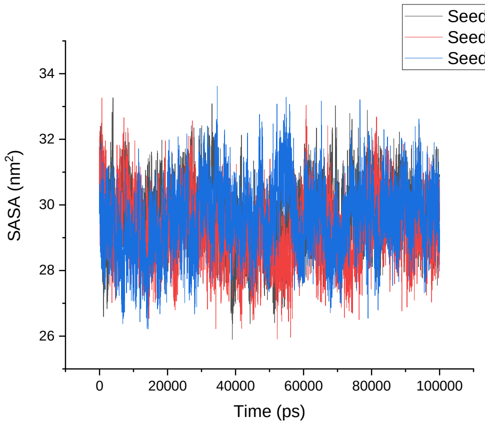 | 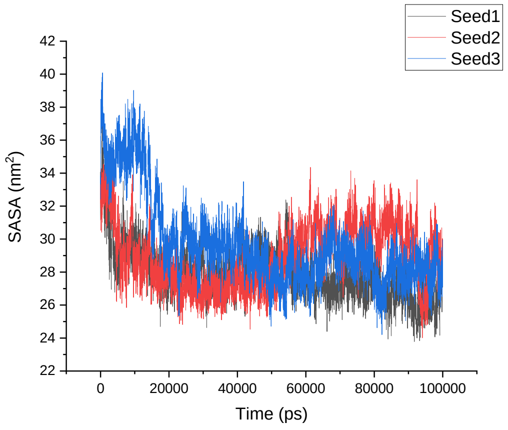 |

| | | |
|---|---|---|
| CHARMM22* | 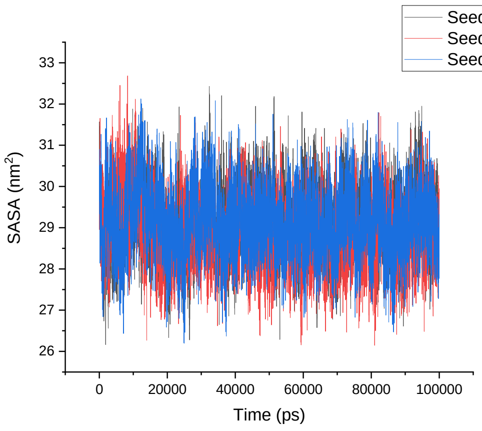  | 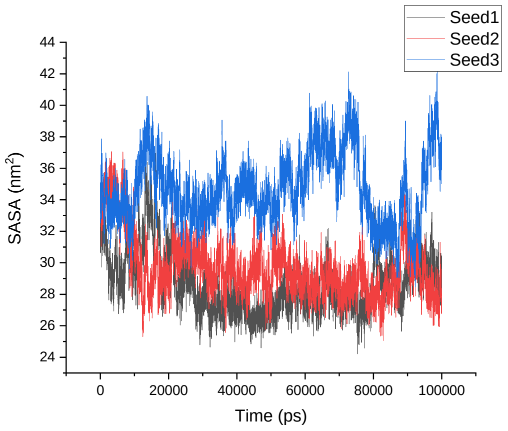  |
| CHARMM36 | 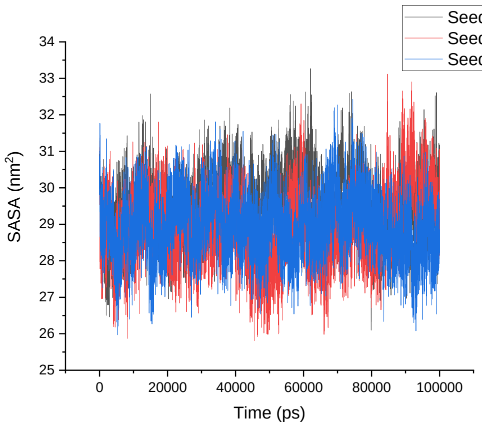  | 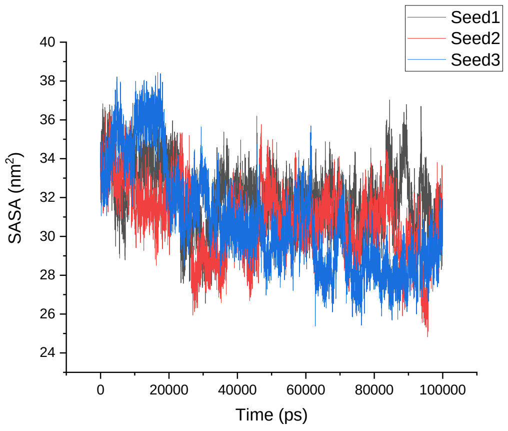  |
| GROMOS 54A8 | 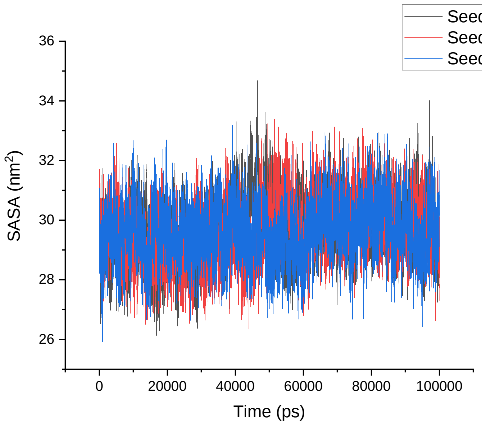  | 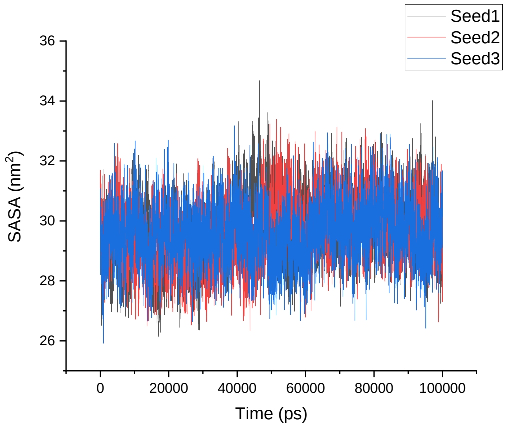  |
| | **8TFV** | **α-8TFV** |

| | | |
|---|---|---|
| Amber 99SB-ILDN | 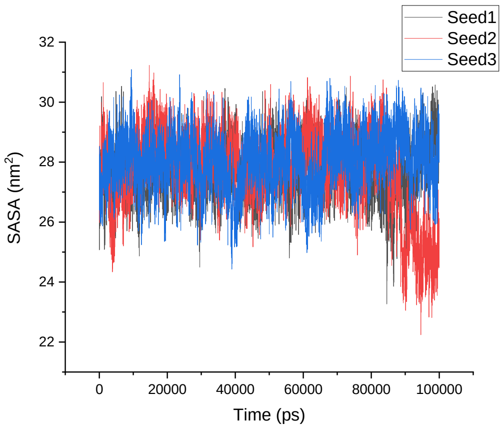 | 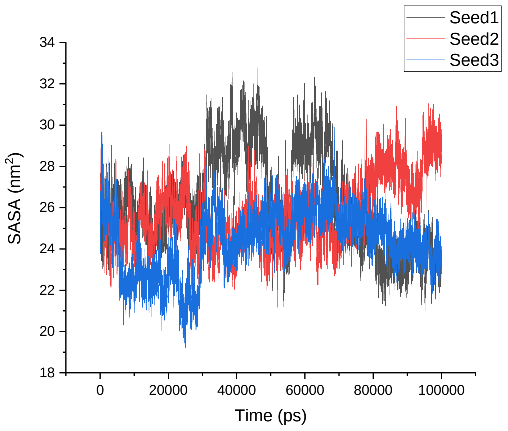 |
| Amber 99SB*-ILDN | 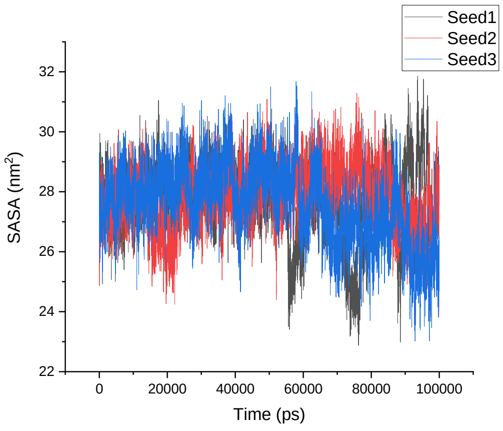 | 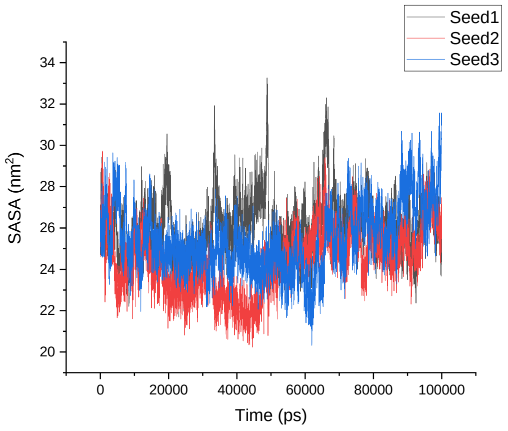 |
| CHARMM22* | 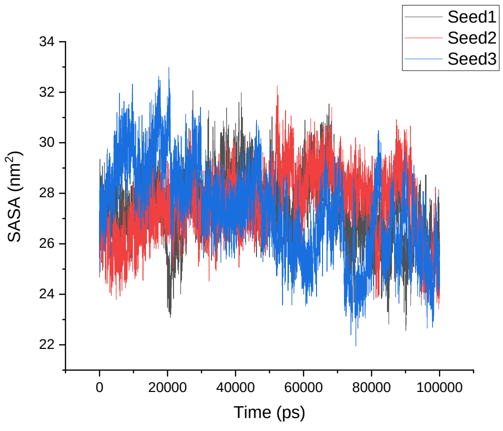 | 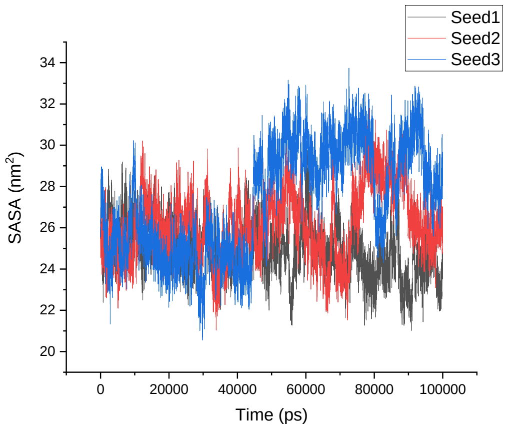 |

<table>
<tr><td>CHARMM36</td><td>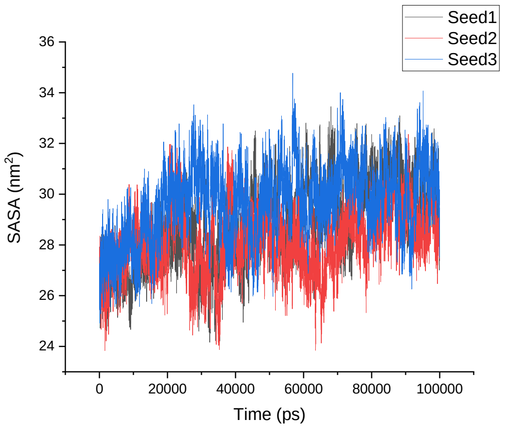
</td><td>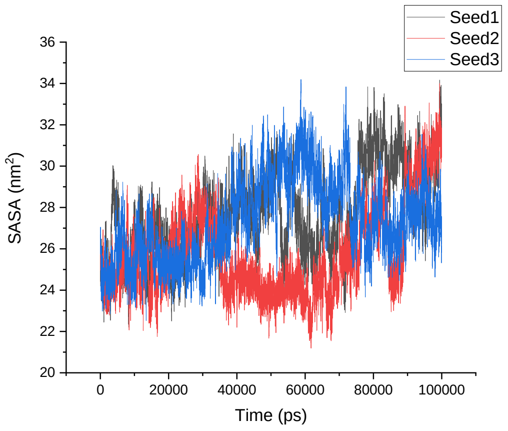
</td></tr>
<tr><td>GROMOS 54A8</td><td>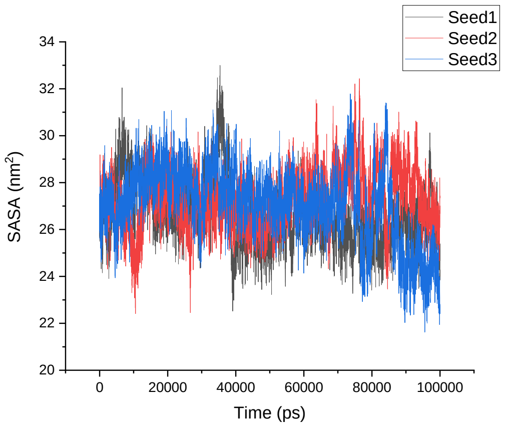
</td><td>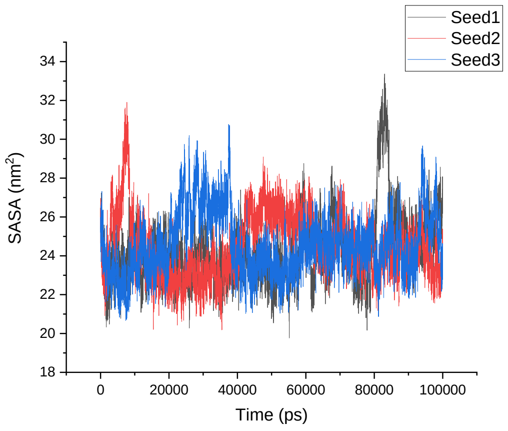
</td></tr>
<tr><td></td><td>1KGM</td><td>α-1KGM</td></tr>
<tr><td>Amber 99SB-ILDN</td><td>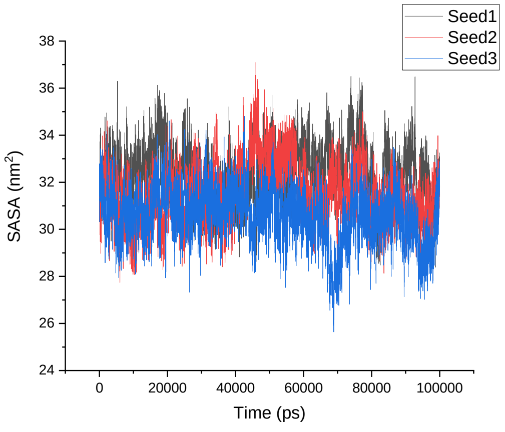
</td><td>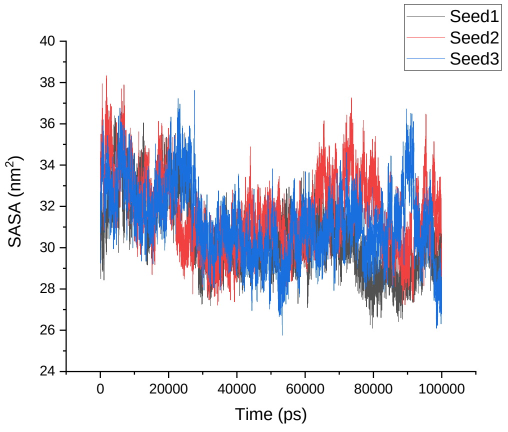
</td></tr>
</table>

| | | |
|---|---|---|
| Amber 99SB*-ILDN | 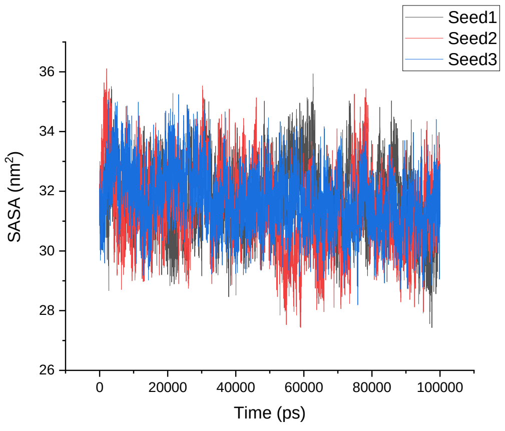  | 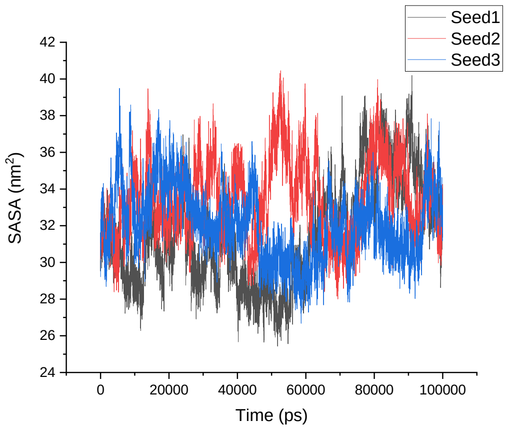  |
| CHARMM22* | 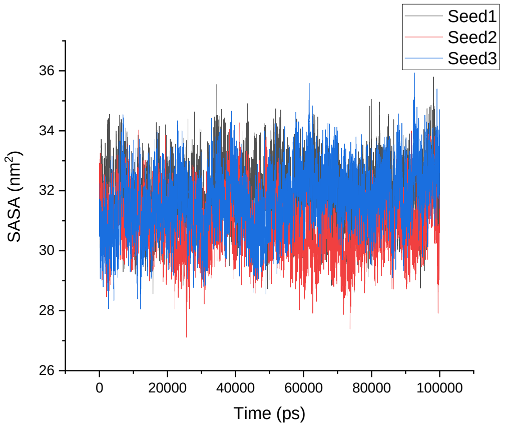  | 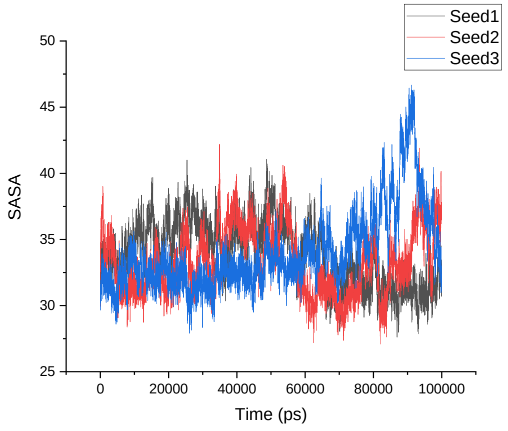  |
| CHARMM36 | 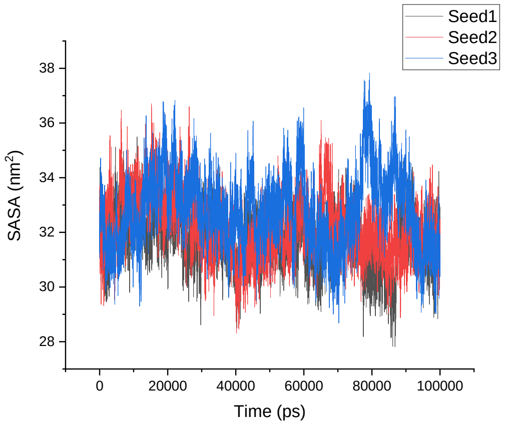  | 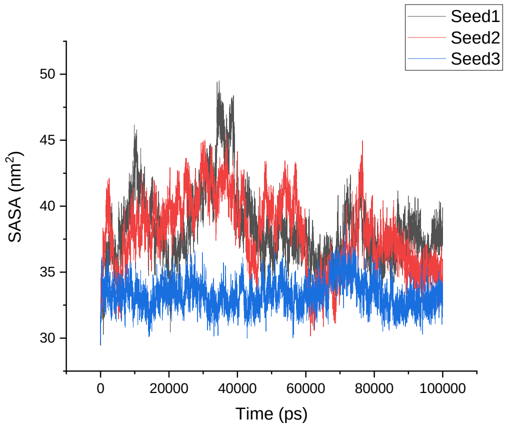  |

<table>
<tr><td>GROMOS 54A8</td><td>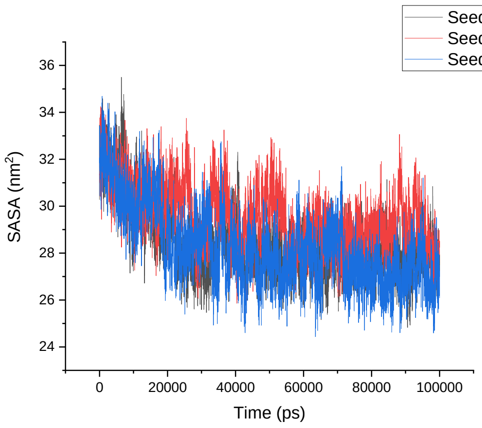
</td><td>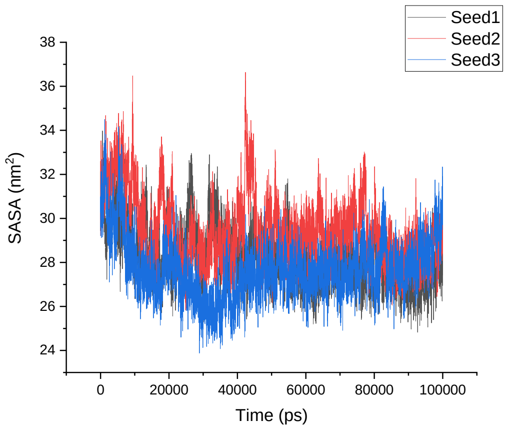
</td></tr>
<tr><td></td><td>1M4F</td><td>α-1M4F</td></tr>
<tr><td>Amber 99SB-ILDN</td><td>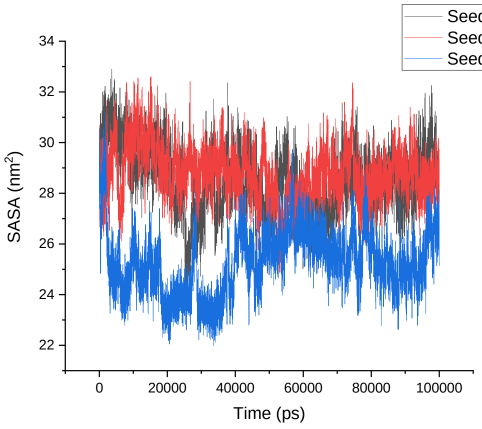
</td><td>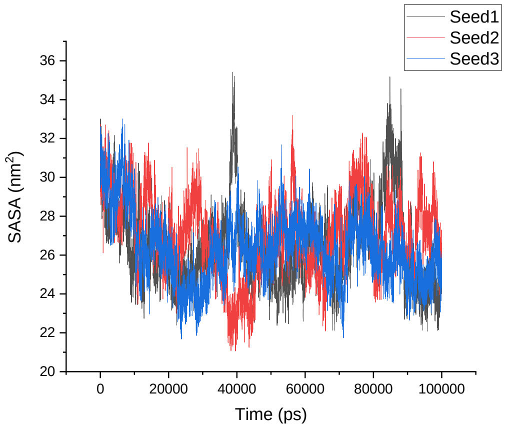
</td></tr>
<tr><td>Amber 99SB*-ILDN</td><td>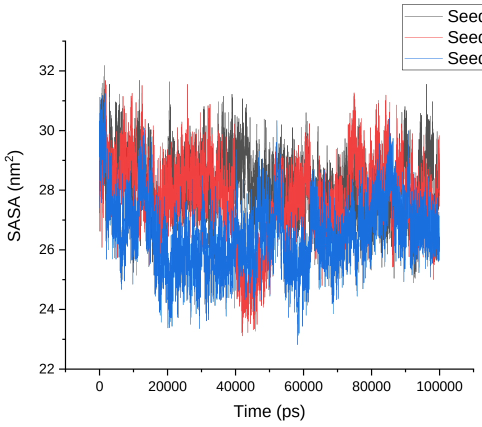
</td><td>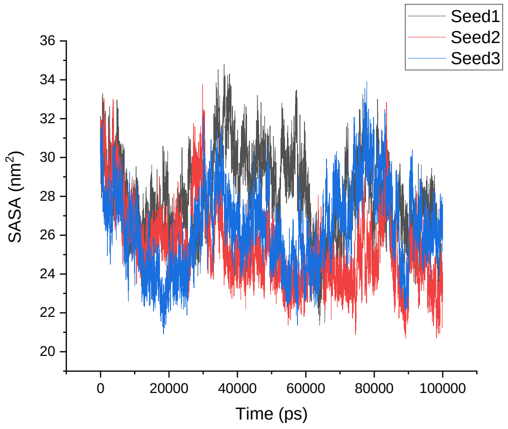
</td></tr>
</table>

| | | |
|---|---|---|
| CHARMM22* | 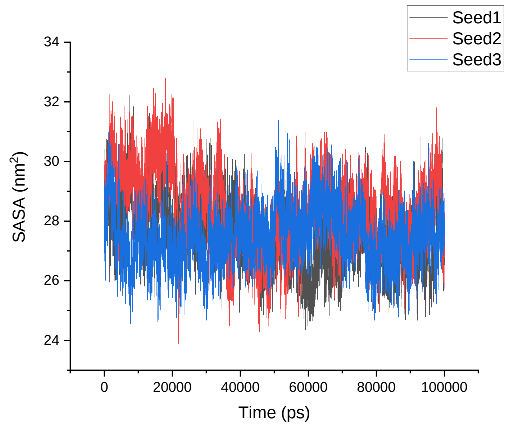 | 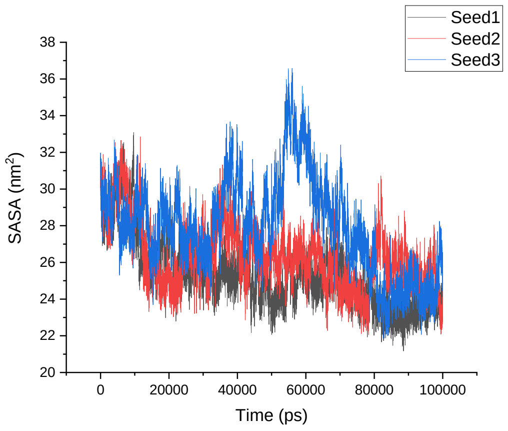 |
| CHARMM36 | 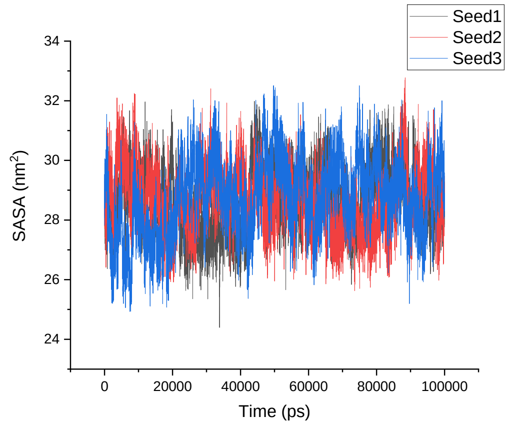 | 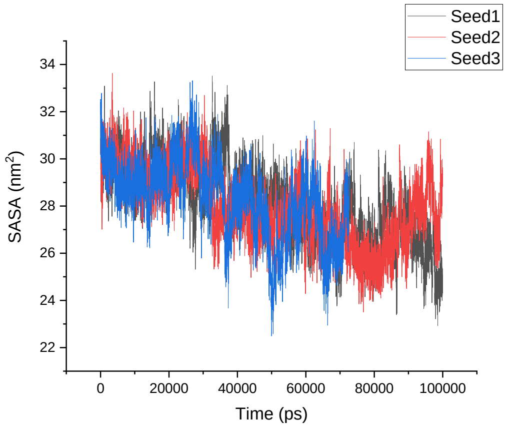 |
| GROMOS 54A8 | 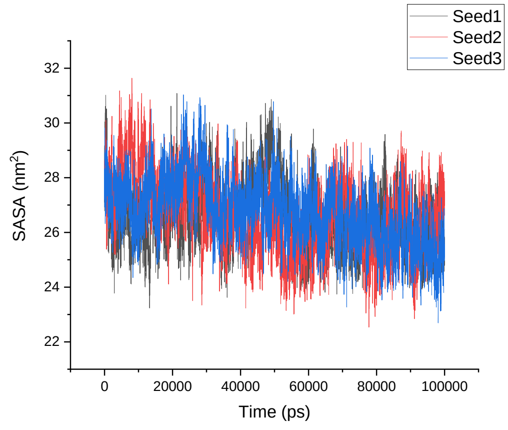 | 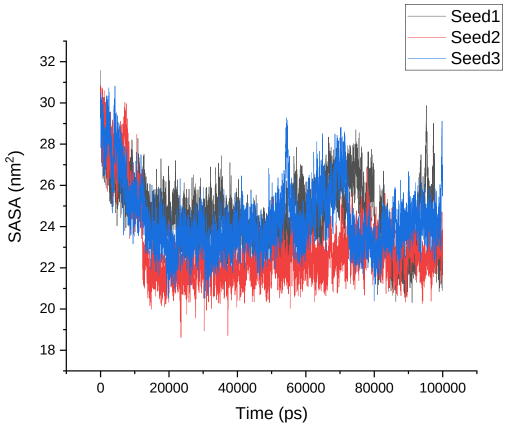 |
| | **1TV0** | **α-1TV0** |

<table>
<tr>
<td>Amber 99SB-ILDN</td>
<td>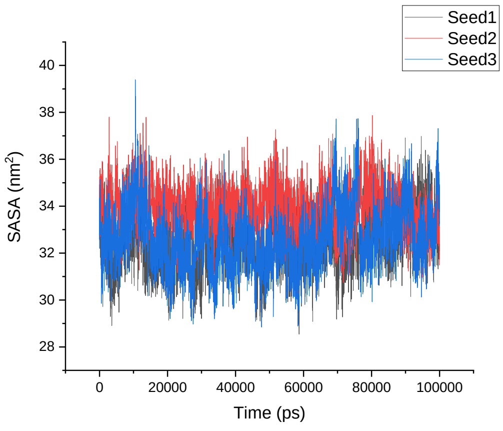
</td>
<td>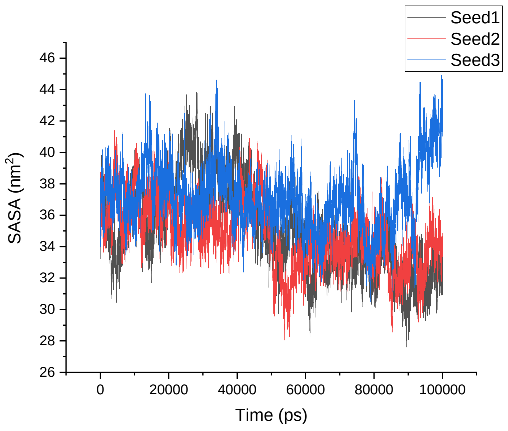
</td>
</tr>
<tr>
<td>Amber 99SB*-ILDN</td>
<td>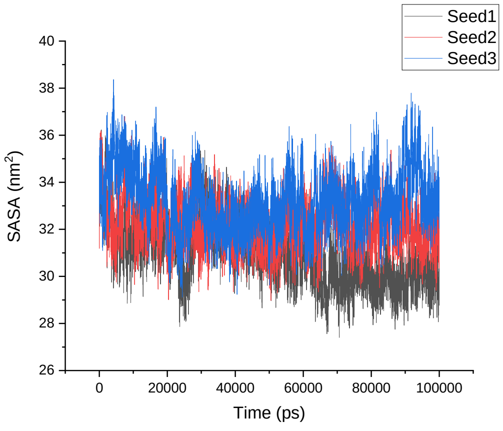
</td>
<td>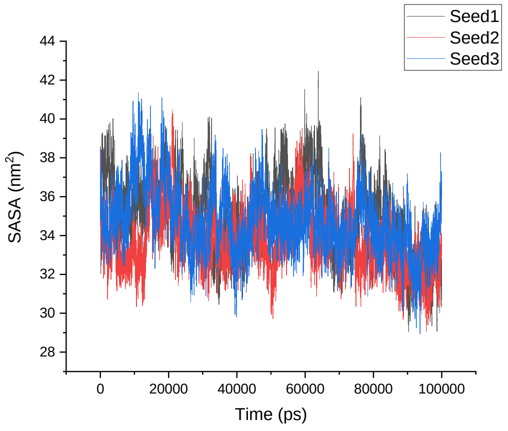
</td>
</tr>
<tr>
<td>CHARMM22*</td>
<td>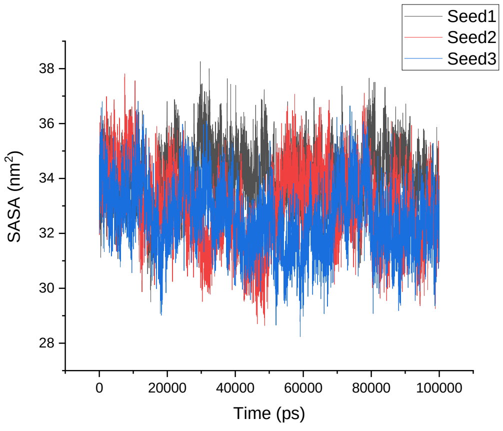
</td>
<td>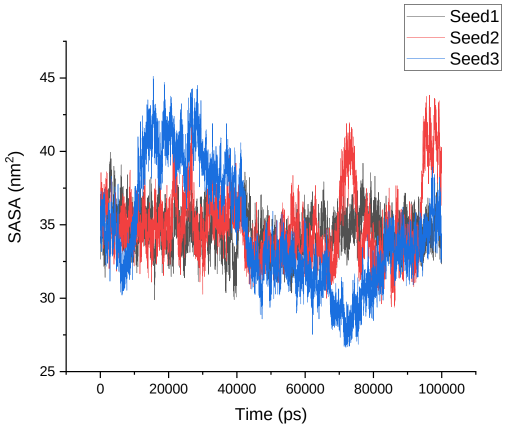
</td>
</tr>
</table>

| | | |
|---|---|---|
| CHARMM36 | 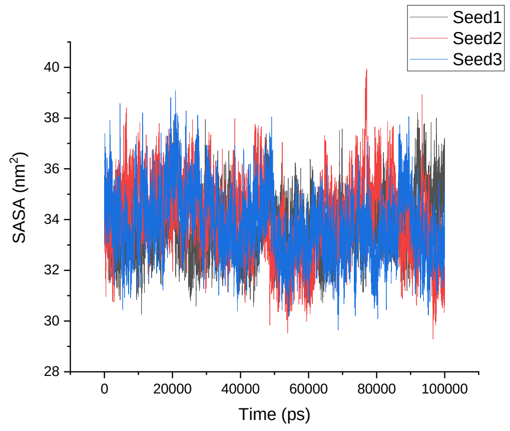  | 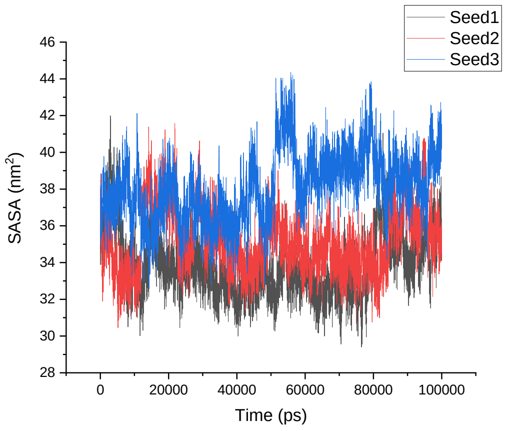  |
| GROMOS 54A8 | 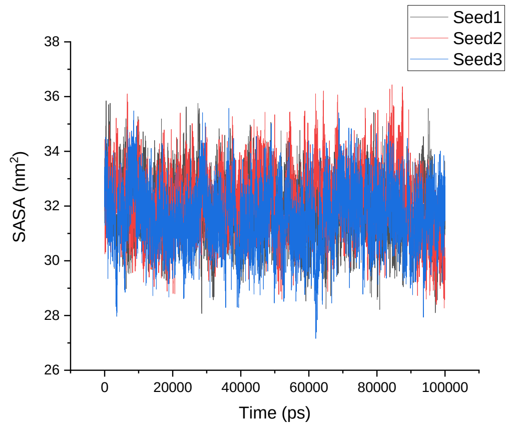  | 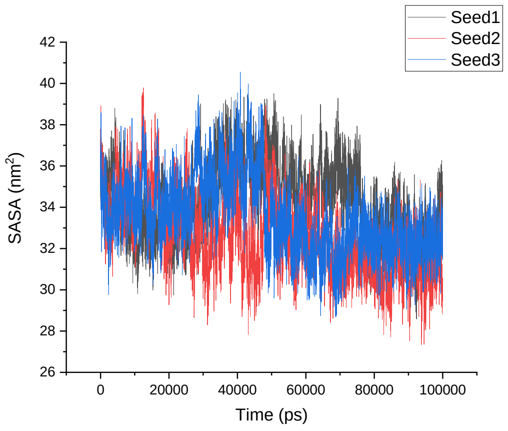  |

**Table S10**. Average SASA over three simulations of the native α-proteins, modeled β-proteins, β-proteins, and modeled α-proteins.

| | SASA | SD | SASA | SD | SASA | SD | SASA | SD | SASA | SD |
|---|---|---|---|---|---|---|---|---|---|---|
| α- proteins | | | | | | | | | | |
| | **1CQ0** | | **1L2Y** | | **1ROO** | | **1T5Q** | | **1V1D** | |
| Amber 99SB-ILDN | 26 | 1 | 19 | 1 | 31 | 1 | 31 | 2 | 32 | 1 |
| Amber 99SB*-ILDN | 26 | 2 | 19 | 1 | 30 | 1 | 30 | 2 | 31 | 1 |
| CHARMM22* | 26 | 2 | 20 | 1 | 31 | 1 | 30 | 2 | 32 | 1 |
| CHARMM36 | 28 | 2 | 19 | 1 | 31 | 1 | 33 | 1 | 32 | 1 |
| GROMOS 54A8 | 25 | 1 | 19 | 1 | 31 | 1 | 31 | 2 | 31 | 1 |
| Modelled β-proteins | | | | | | | | | | |
| | **β-1CQ0** | | **β-1L2Y** | | **β-1ROO** | | **β-1T5Q** | | **β-1V1D** | |
| Amber 99SB-ILDN | 28 | 2 | 23 | 2 | 34 | 2 | 33 | 2 | 30 | 1 |
| Amber 99SB*-ILDN | 29 | 2 | 23 | 2 | 36 | 2 | 32 | 2 | 31 | 2 |
| CHARMM22* | 29 | 2 | 24 | 2 | 36 | 2 | 33 | 2 | 31 | 1 |
| CHARMM36 | 31 | 3 | 25 | 2 | 41 | 2 | 35 | 2 | 33 | 2 |
| GROMOS 54A8 | 26 | 2 | 21 | 1 | 33 | 2 | 30 | 2 | 29 | 1 |
| β- proteins | | | | | | | | | | |
| | **1LFC** | | **8TFV** | | **1KGM** | | **1M4F** | | **1TV0** | |
| Amber 99SB-ILDN | 29 | 1 | 28 | 1 | 31 | 1 | 28 | 1 | 33 | 1 |
| Amber 99SB*-ILDN | 29 | 1 | 28 | 1 | 32 | 1 | 27 | 1 | 32 | 1 |
| CHARMM22* | 29 | 1 | 27 | 1 | 31 | 1 | 28 | 1 | 33 | 1 |
| CHARMM36 | 29 | 1 | 29 | 1 | 32 | 1 | 29 | 1 | 34 | 1 |
| GROMOS 54A8 | 30 | 1 | 27 | 1 | 29 | 2 | 27 | 1 | 32 | 1 |
| Modelled α- proteins | | | | | | | | | | |
| | **α-1LFC** | | **α-8TFV** | | **α-1KGM** | | **α-1M4F** | | **α-1TV0** | |
| Amber 99SB-ILDN | 31 | 3 | 25 | 2 | 31 | 2 | 27 | 2 | 36 | 2 |
| Amber 99SB*-ILDN | 29 | 2 | 25 | 2 | 32 | 2 | 27 | 2 | 35 | 2 |
| CHARMM22* | 31 | 2 | 26 | 2 | 34 | 3 | 27 | 2 | 35 | 3 |
| CHARMM36 | 31 | 2 | 27 | 2 | 36 | 2 | 28 | 2 | 36 | 2 |
| GROMOS 54A8 | 29 | 1 | 24 | 2 | 28 | 1 | 24 | 2 | 33 | 2 |

**Table S11**. Number of intra-protein hydrogen bonds during the simulation time of three seeded simulations of the native α-proteins, modeled β-proteins, β-proteins, and modeled α-proteins.

| Hydrogen bonds | | |
|---|---|---|
| | **1CQ0** | **β-1CQ0** |
| Amber 99SB-ILDN | 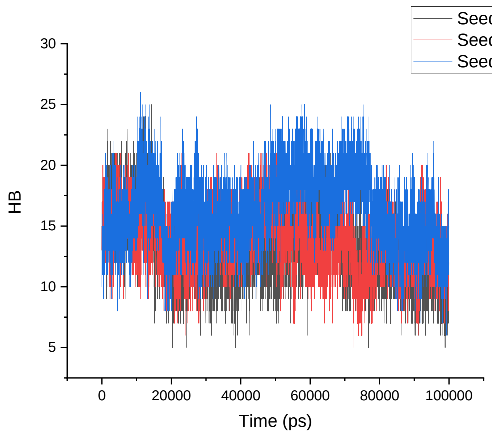 | 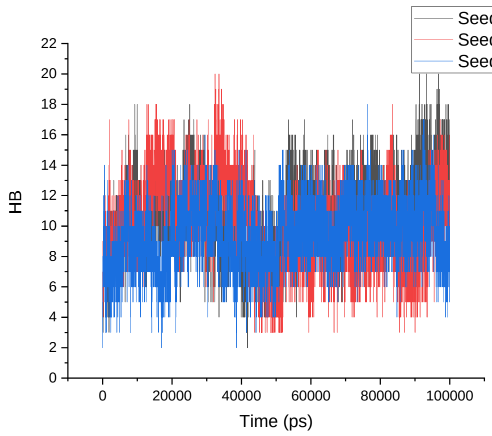 |
| Amber 99SB*-ILDN | 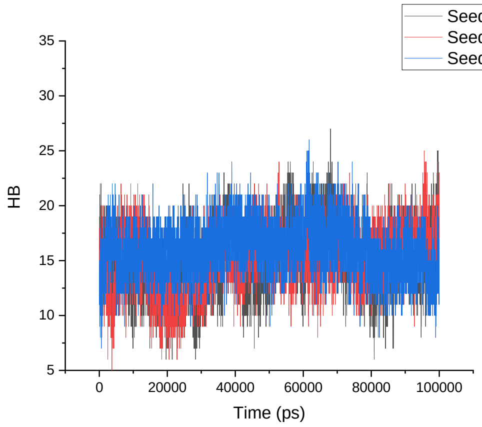 | 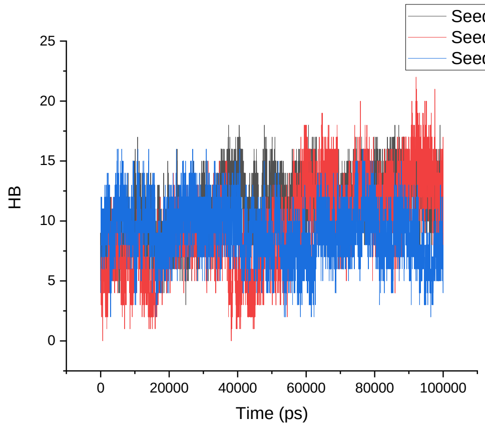 |
| CHARMM22* | 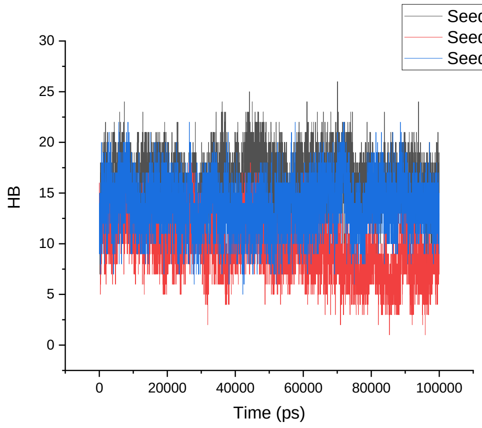 | 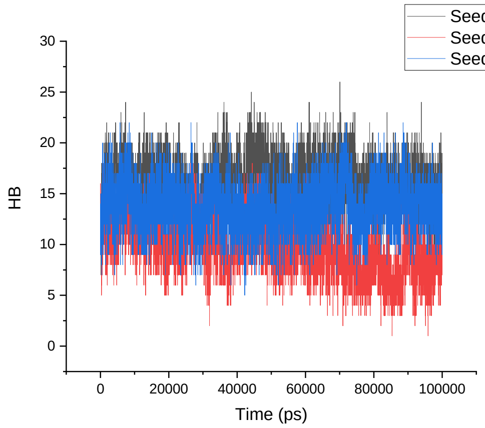 |

| | | |
|---|---|---|
| CHARMM36 | 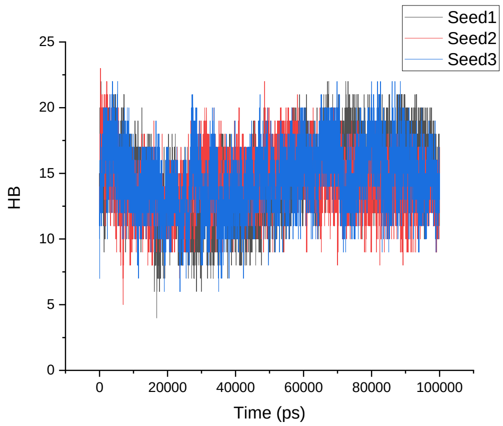 | 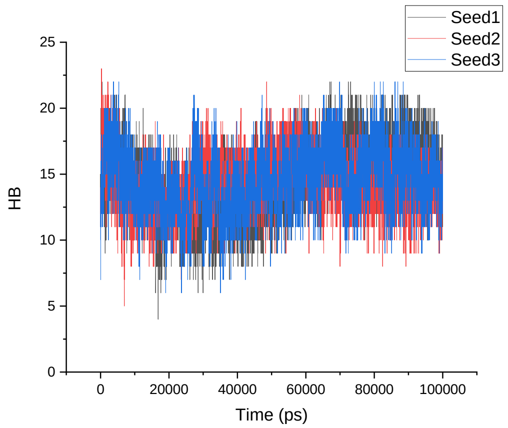 |
| GROMOS 54A8 | 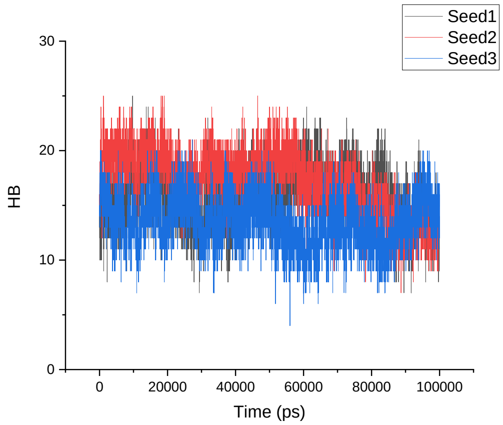 | 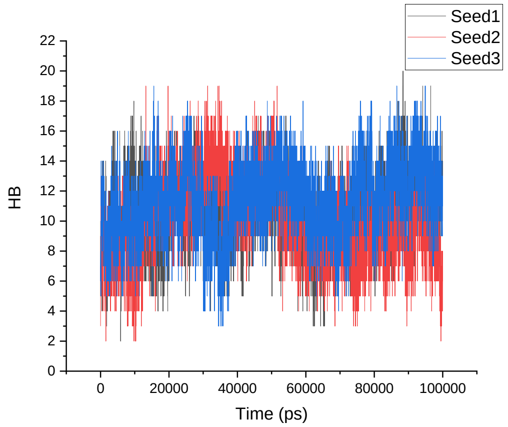 |
| | **1L2Y** | **β-1L2Y** |
| Amber 99SB-ILDN | 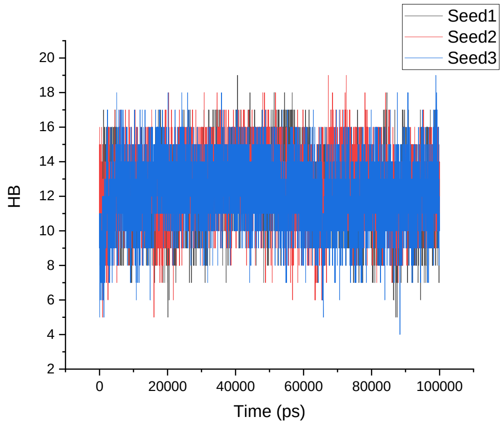 | 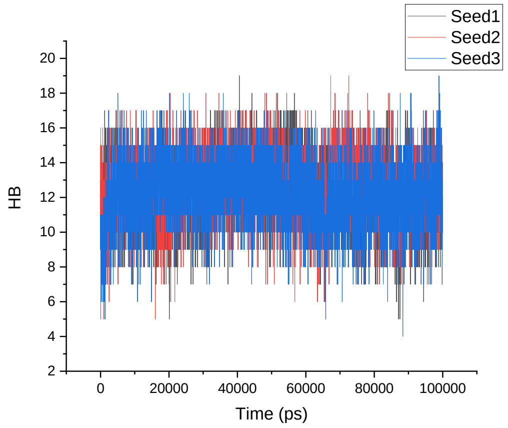 |

<table>
<tr>
<td>Amber 99SB*-ILDN</td>
<td>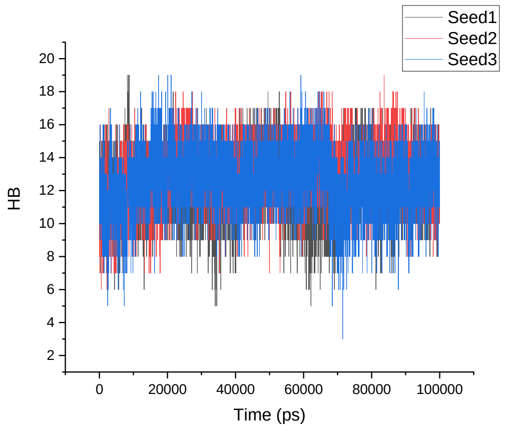
</td>
<td>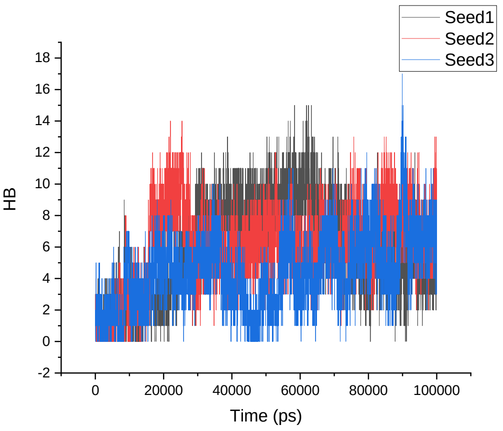
</td>
</tr>
<tr>
<td>CHARMM22*</td>
<td>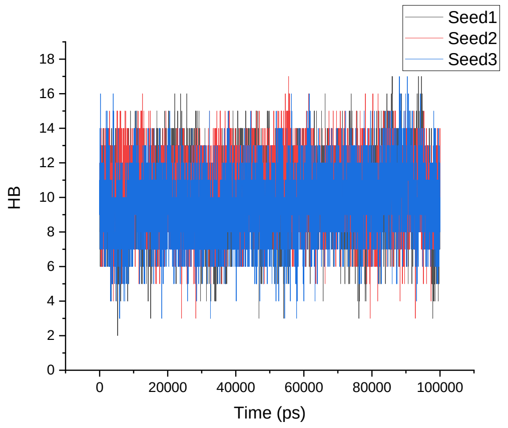
</td>
<td>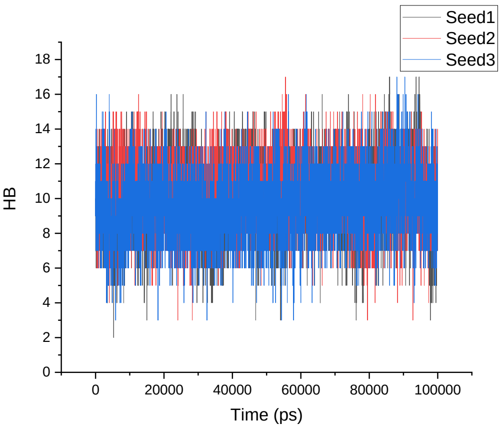
</td>
</tr>
<tr>
<td>CHARMM36</td>
<td>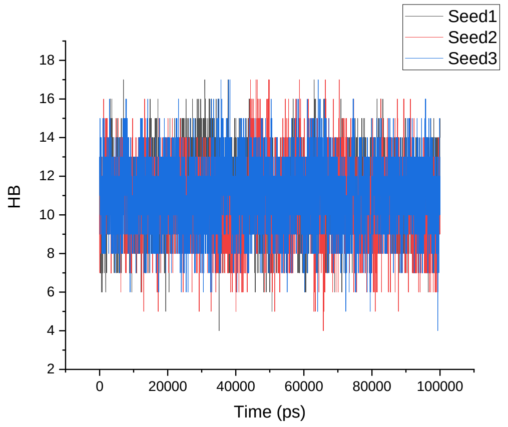
</td>
<td>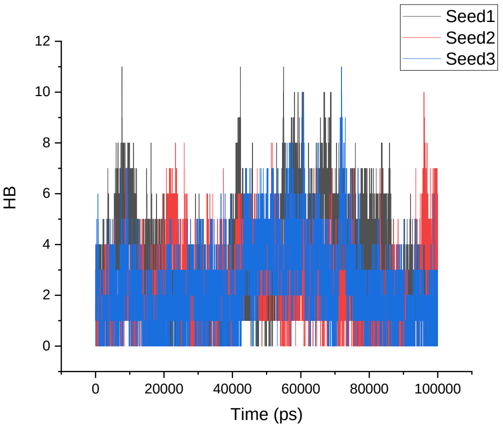
</td>
</tr>
</table>

<table>
<tr><td>GROMOS 54A8</td><td>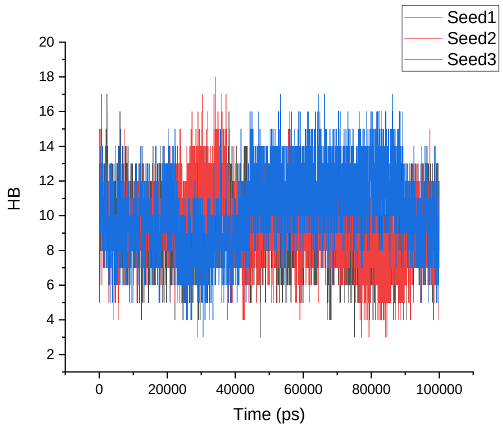
</td><td>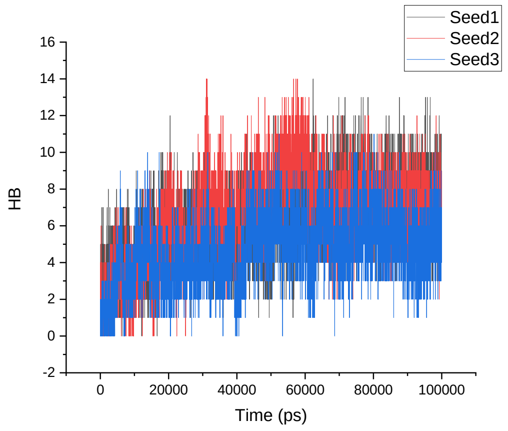
</td></tr>
<tr><td></td><td>1ROO</td><td>β-1ROO</td></tr>
<tr><td>Amber 99SB-ILDN</td><td>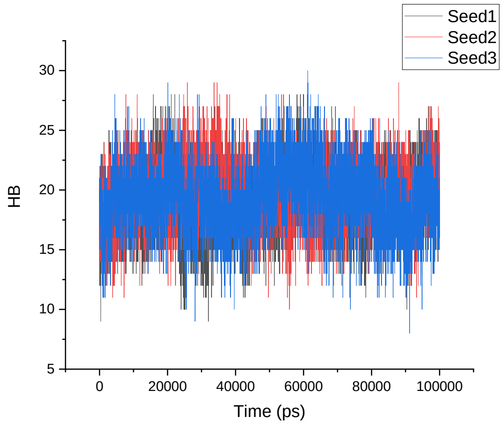
</td><td>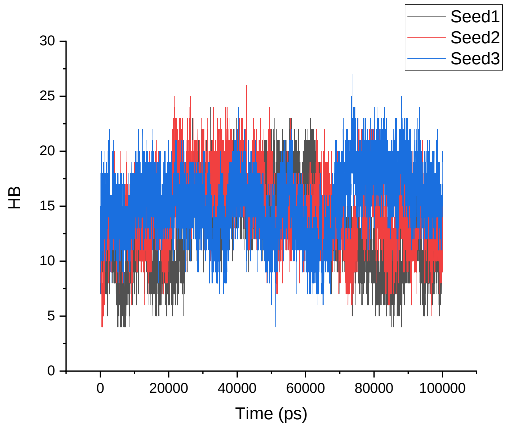
</td></tr>
<tr><td>Amber 99SB*-ILDN</td><td>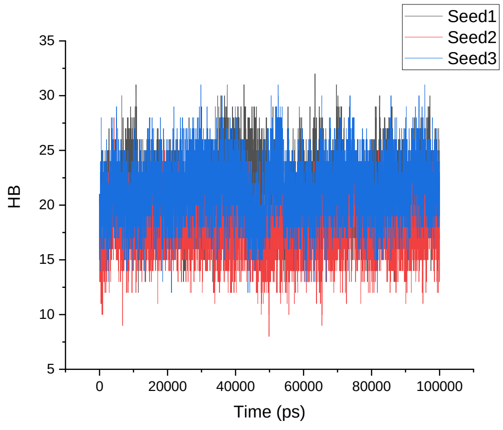
</td><td>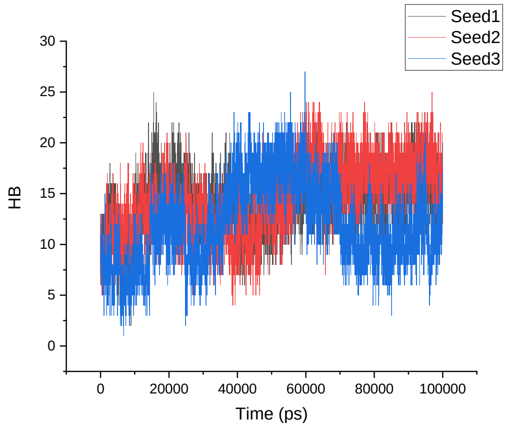
</td></tr>
</table>

| | | |
|---|---|---|
| CHARMM22* | 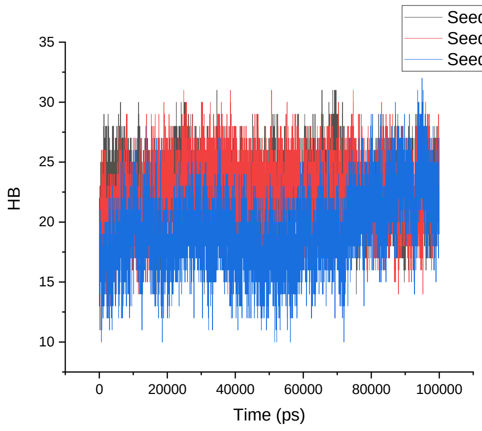 | 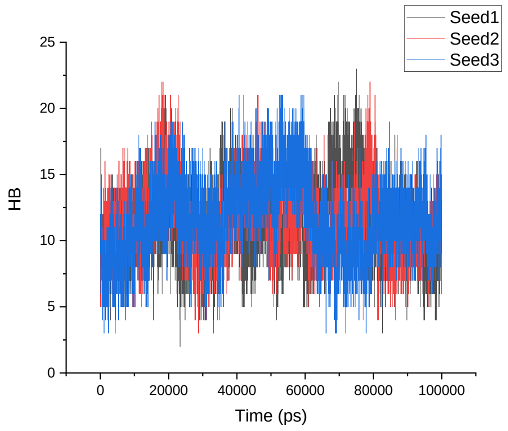 |
| CHARMM36 | 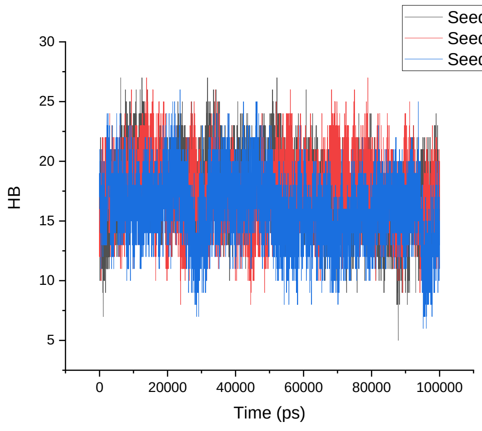 | 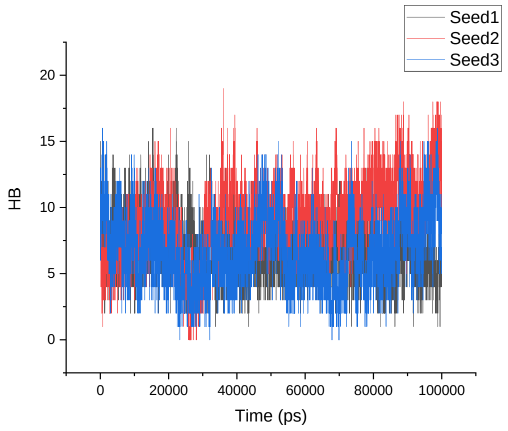 |
| GROMOS 54A8 | 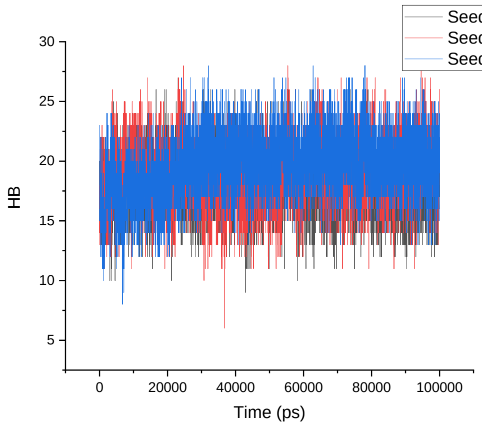 | 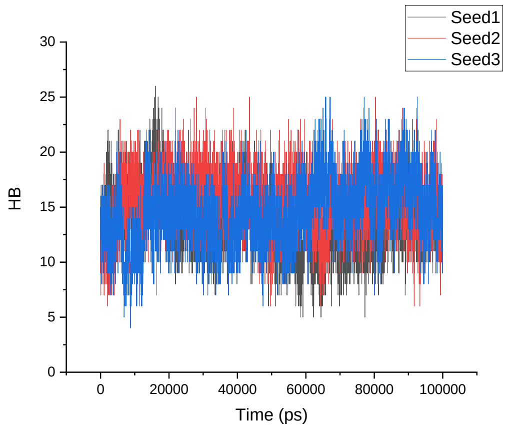 |
| | **1T5Q** | **β-1T5Q** |

| | | |
|---|---|---|
| Amber 99SB-ILDN | 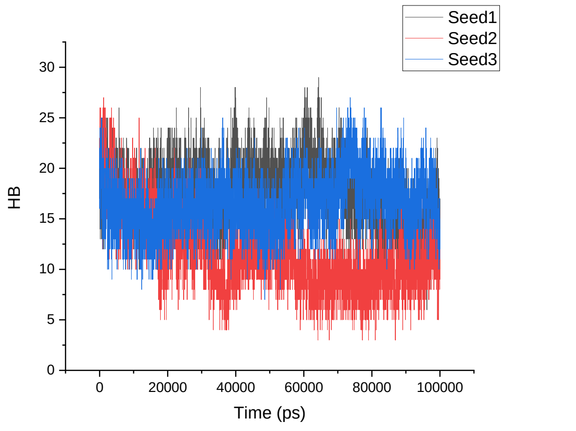 | 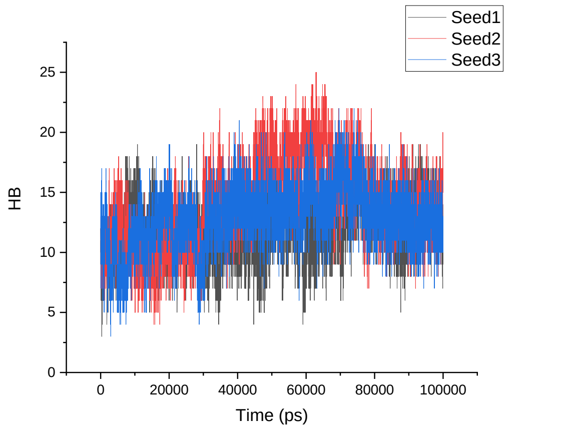 |
| Amber 99SB*-ILDN | 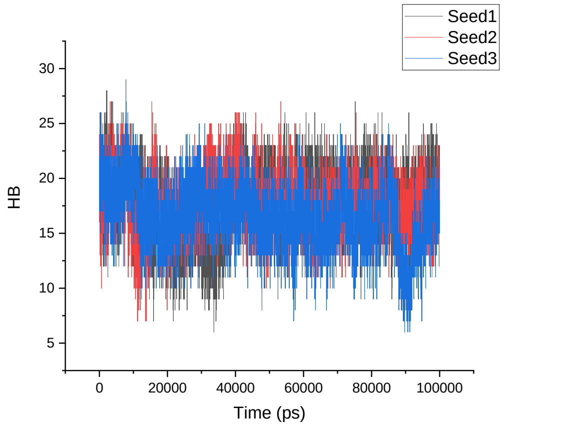 | 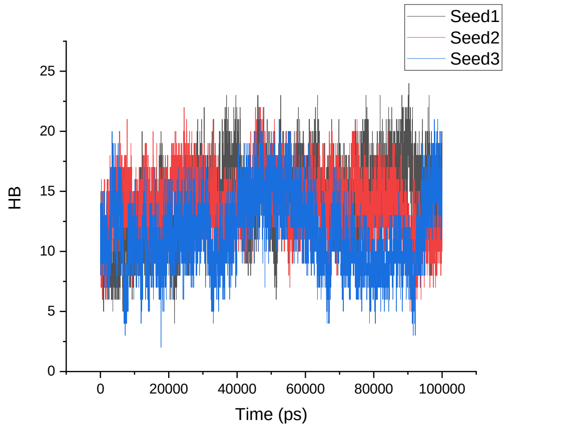 |
| CHARMM22* | 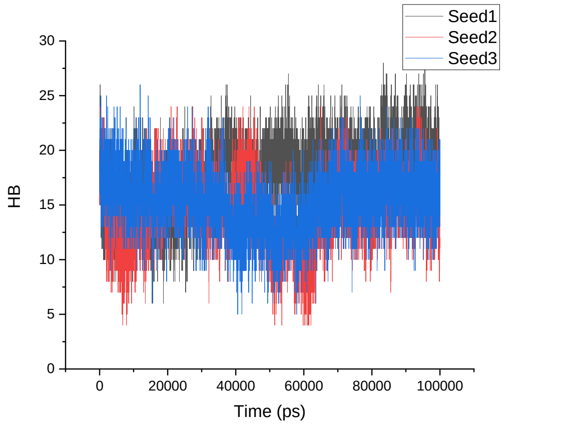 | 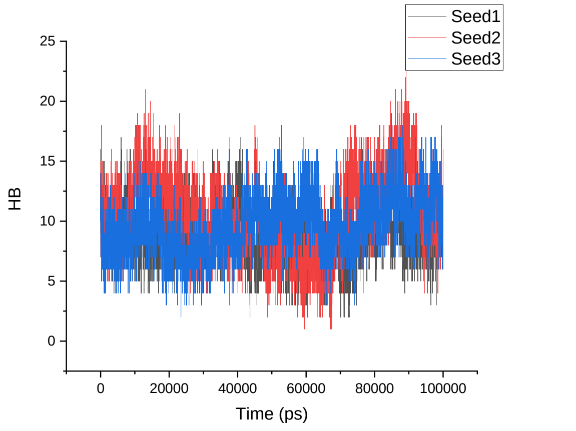 |

<table>
<tr><td>CHARMM36</td><td>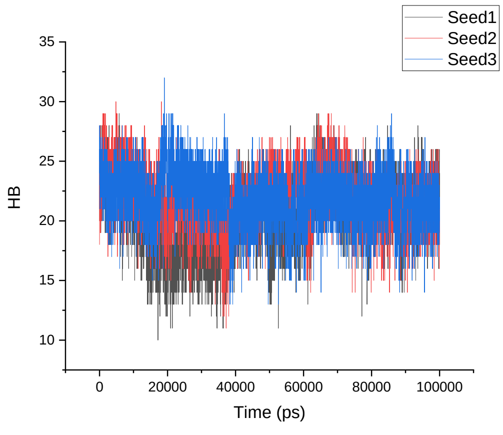
</td><td>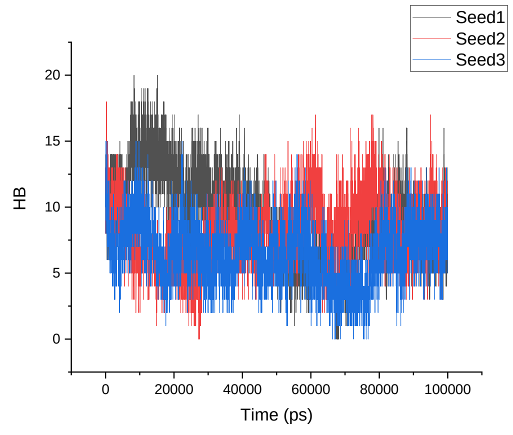
</td></tr>
<tr><td>GROMOS 54A8</td><td>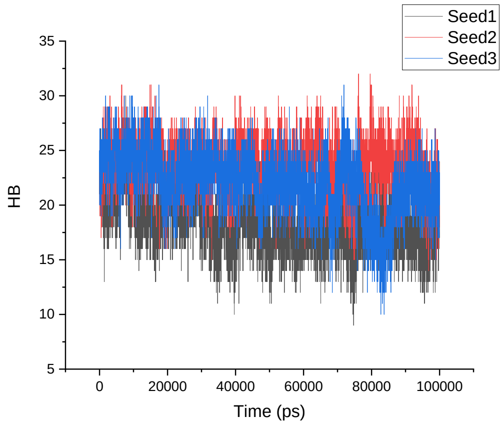
</td><td>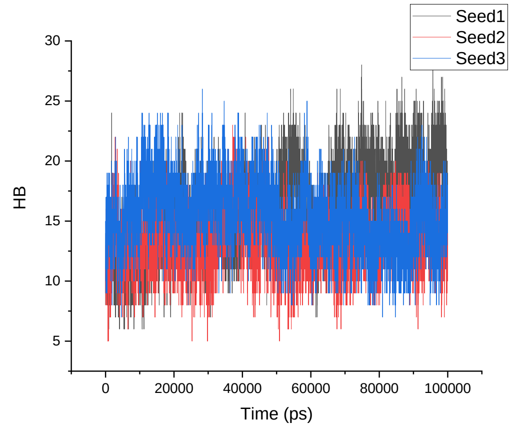
</td></tr>
<tr><td></td><td>1V1D</td><td>β-1V1D</td></tr>
<tr><td>Amber 99SB-ILDN</td><td>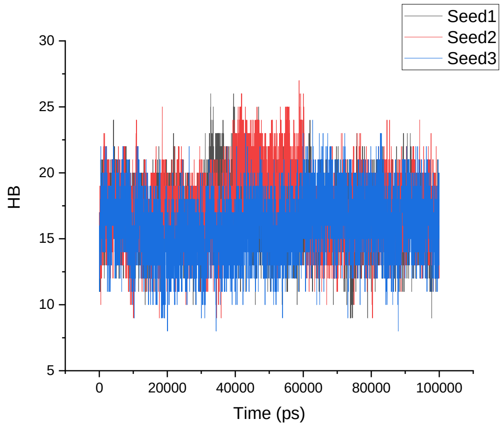
</td><td>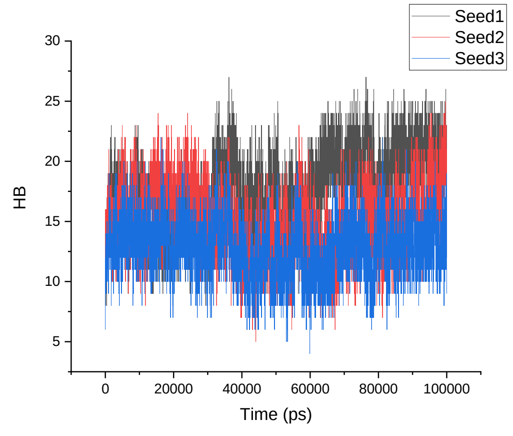
</td></tr>
</table>

| | | |
|---|---|---|
| Amber 99SB*-ILDN | 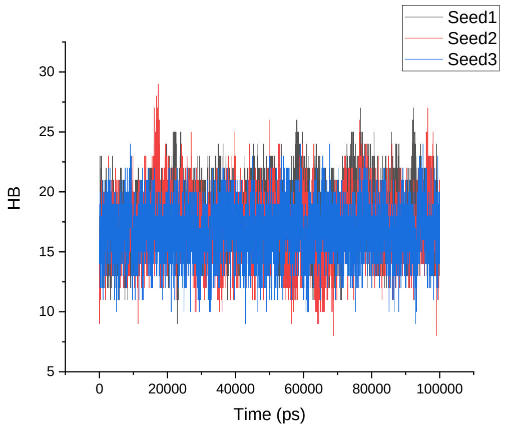 | 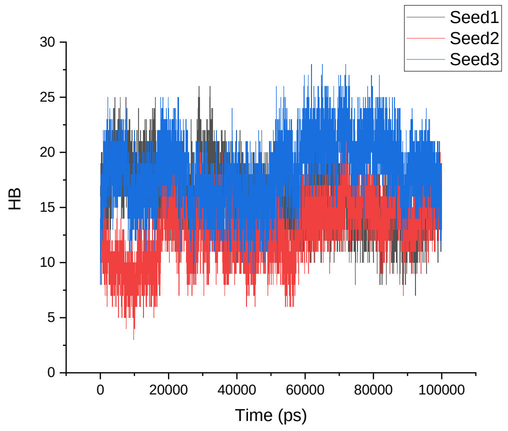 |
| CHARMM22* | 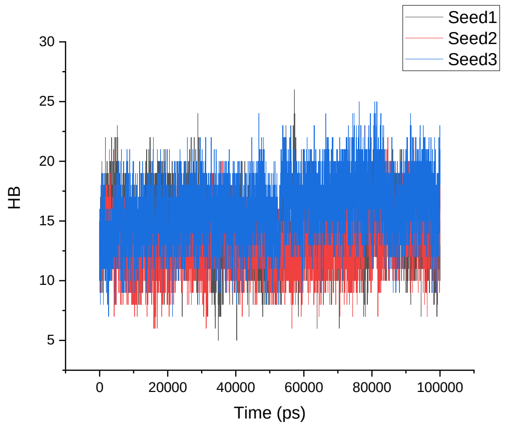 | 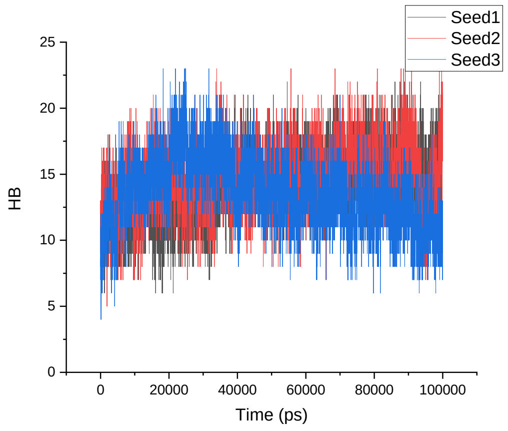 |
| CHARMM36 | 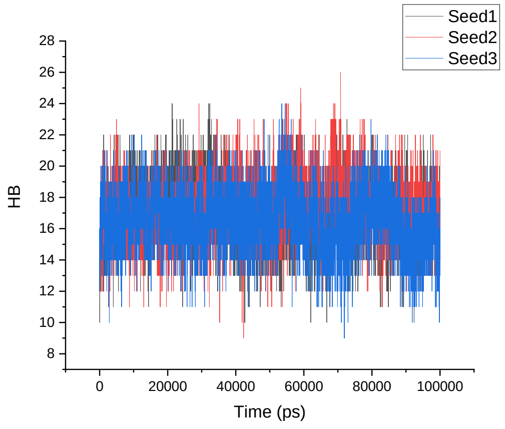 | 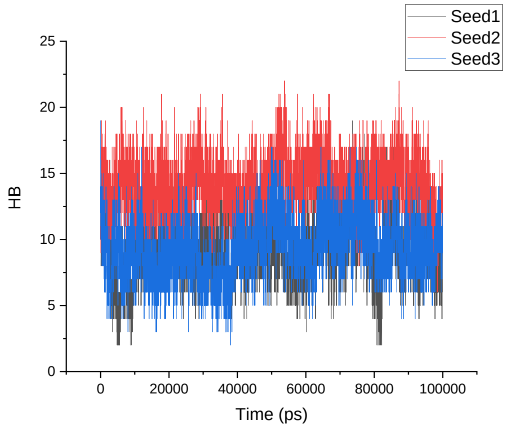 |

<table>
<tr><td>GROMOS 54A8</td><td>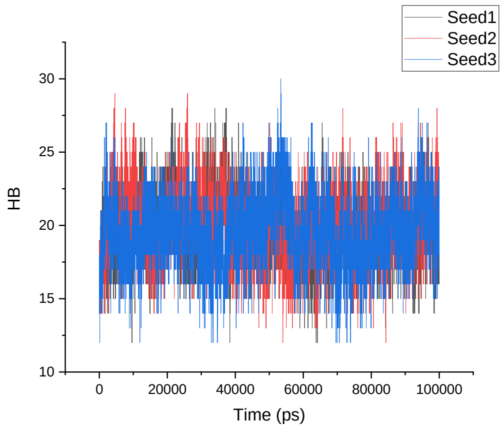
</td><td>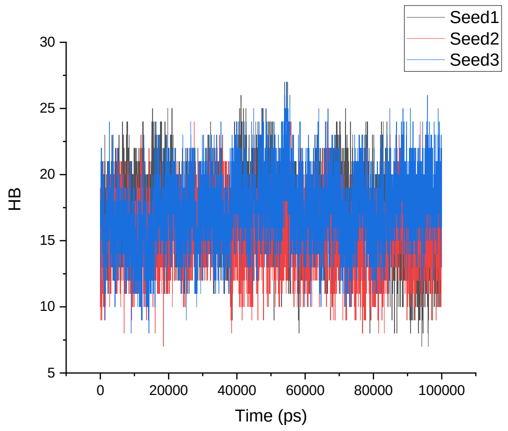
</td></tr>
<tr><td></td><td>1LFC</td><td>α-1LFC</td></tr>
<tr><td>Amber 99SB-ILDN</td><td>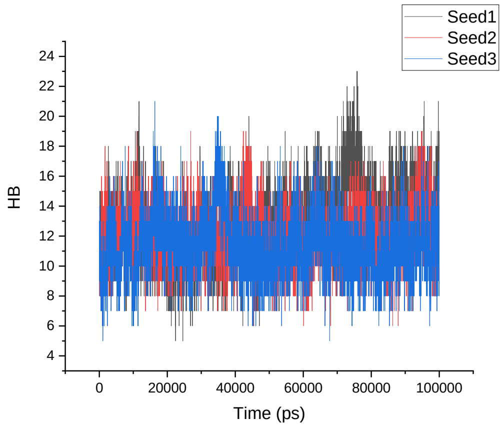
</td><td>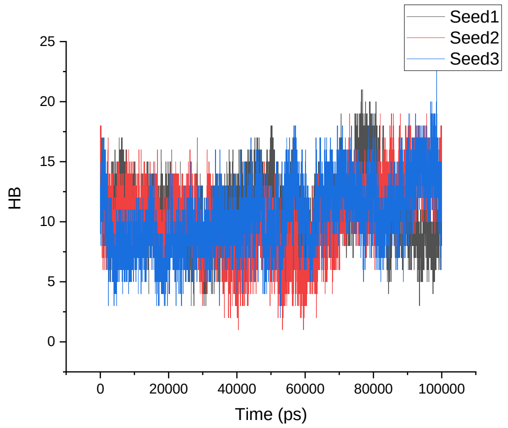
</td></tr>
<tr><td>Amber 99SB*-ILDN</td><td>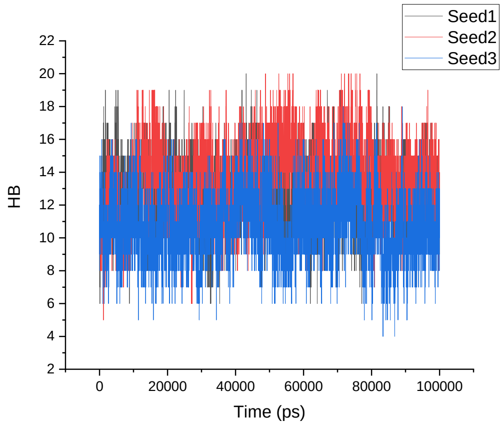
</td><td>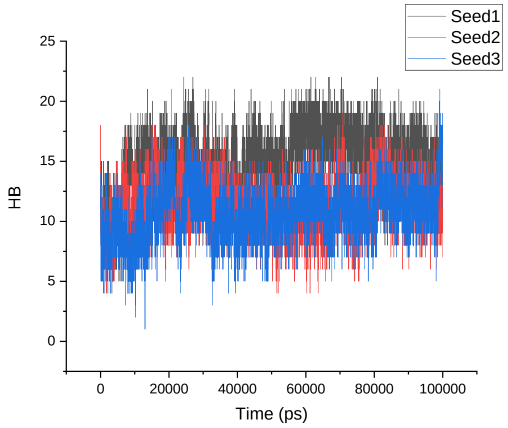
</td></tr>
</table>

<table>
<tr><td>CHARMM22*</td><td>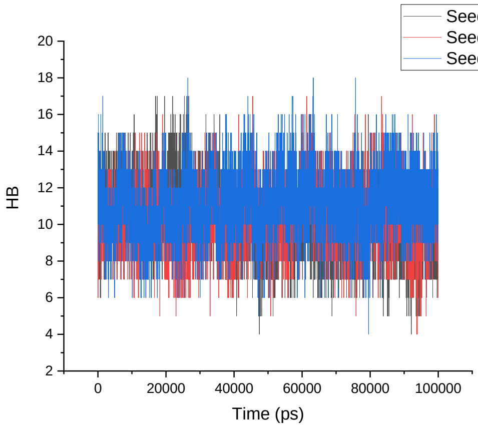
</td><td>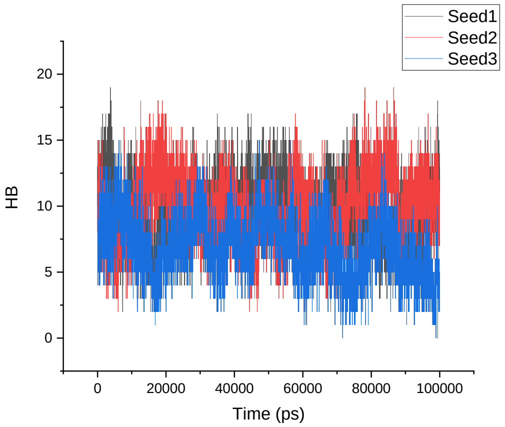
</td></tr>
<tr><td>CHARMM36</td><td>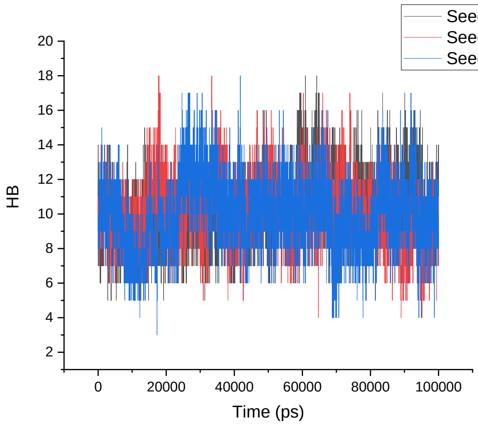
</td><td>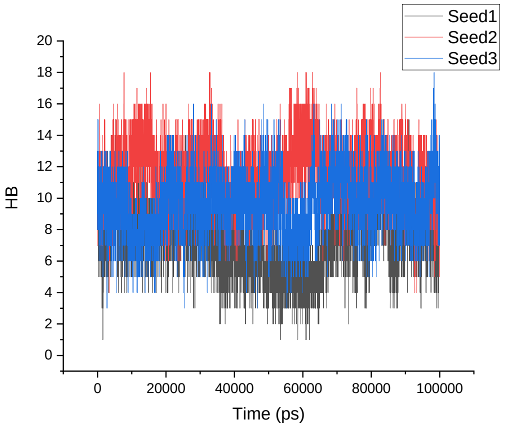
</td></tr>
<tr><td>GROMOS<br>54A8</td><td>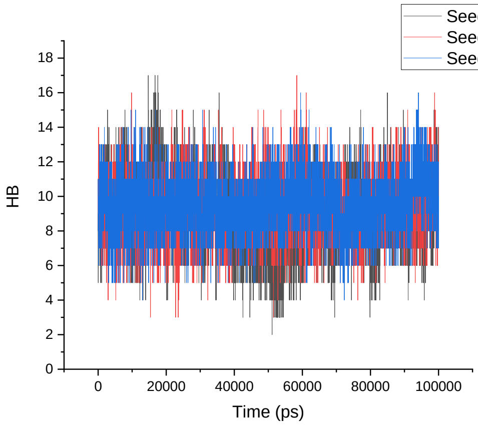
</td><td>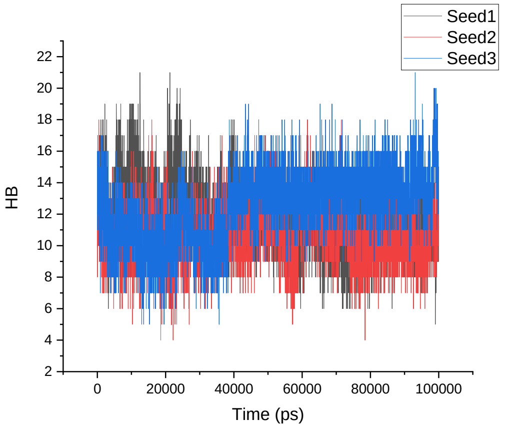
</td></tr>
<tr><td></td><td>8TFV</td><td>α-8TFV</td></tr>
</table>

<table>
<tr>
<td>Amber 99SB-ILDN</td>
<td>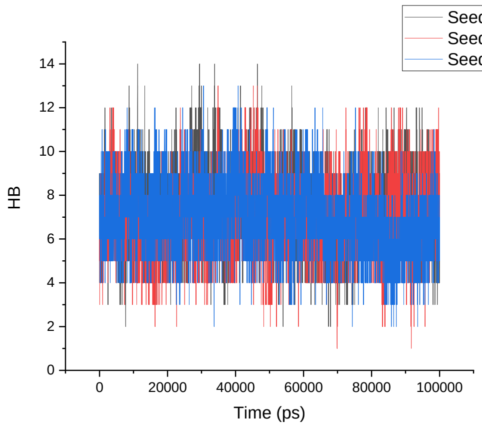
</td>
<td>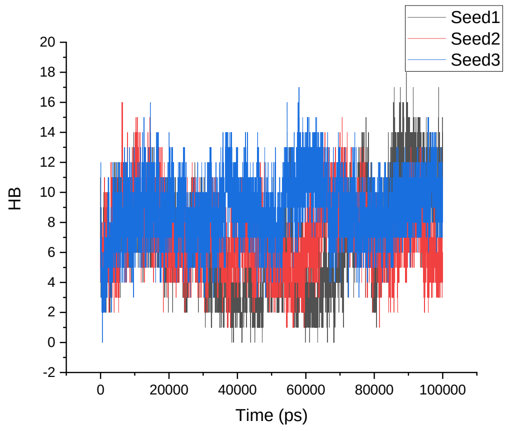
</td>
</tr>
<tr>
<td>Amber 99SB*-ILDN</td>
<td>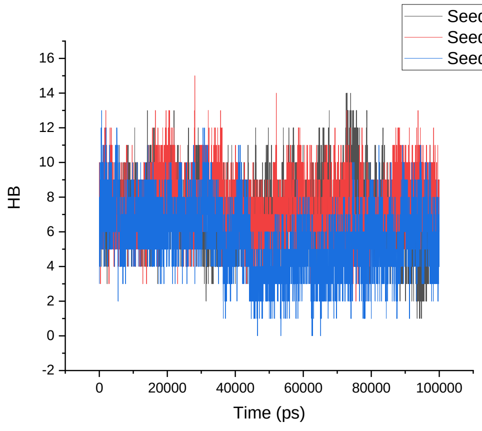
</td>
<td>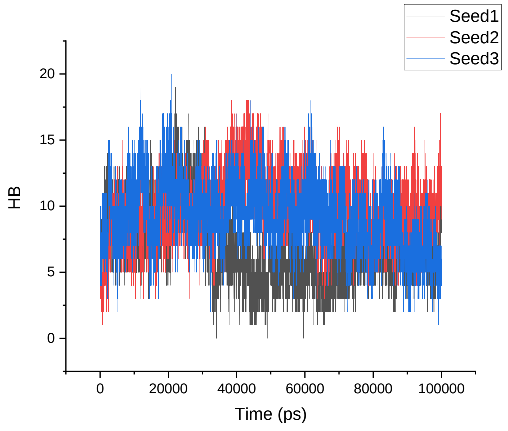
</td>
</tr>
<tr>
<td>CHARMM22*</td>
<td>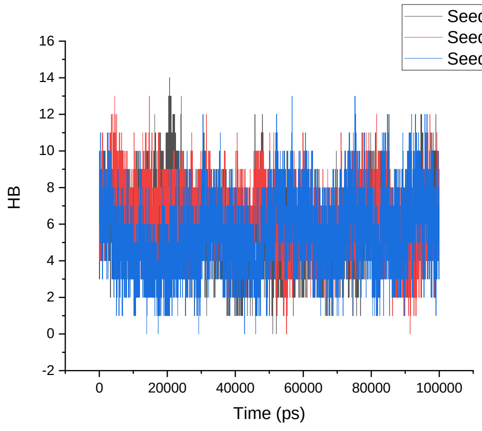
</td>
<td>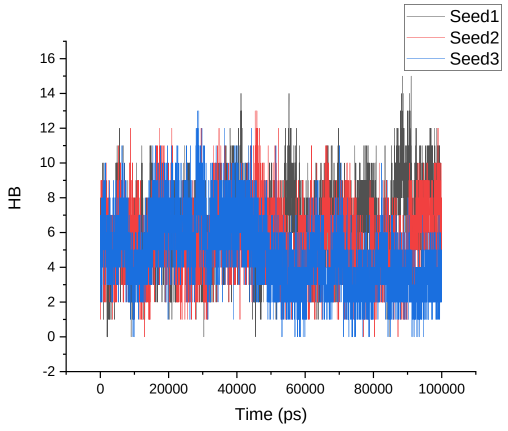
</td>
</tr>
</table>

<table>
<tr><td>CHARMM36</td><td>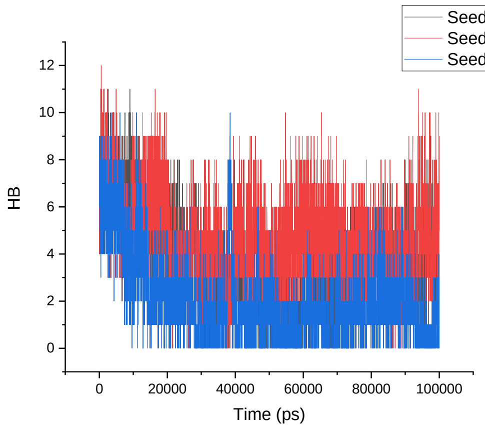
</td><td>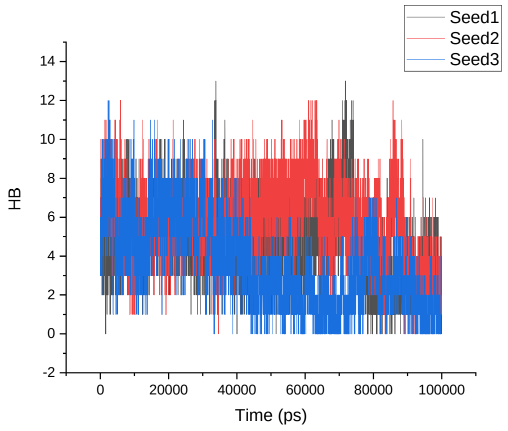
</td></tr>
<tr><td>GROMOS 54A8</td><td>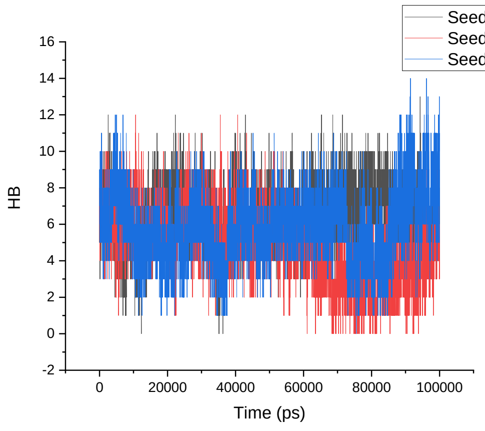
</td><td>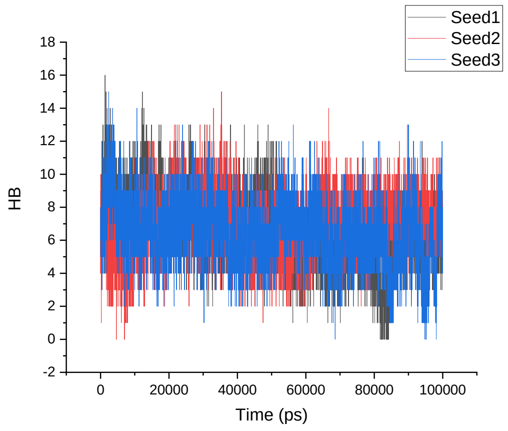
</td></tr>
<tr><td></td><td>1KGM</td><td>α-1KGM</td></tr>
<tr><td>Amber 99SB-ILDN</td><td>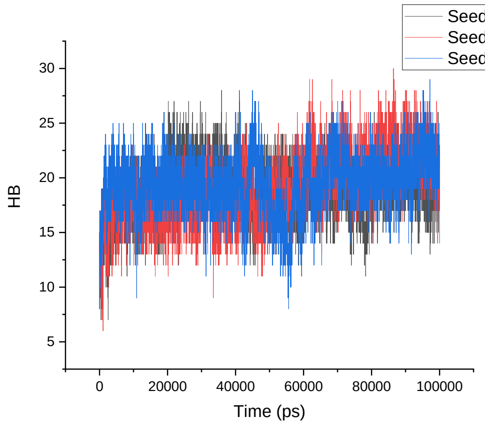
</td><td>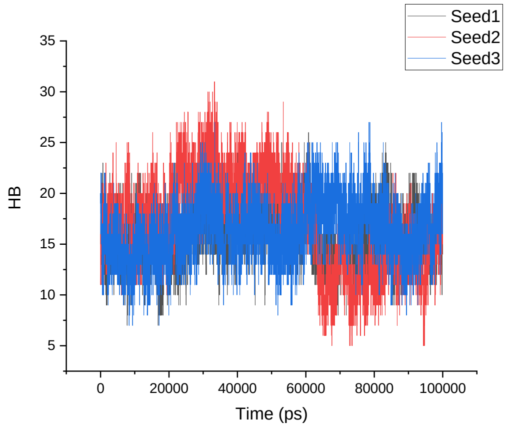
</td></tr>
</table>

| | | |
|---|---|---|
| Amber 99SB*-ILDN | 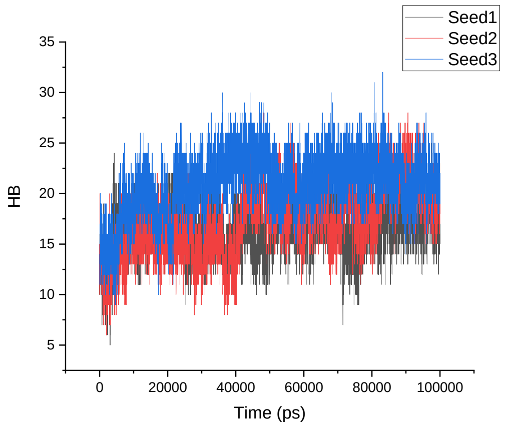 | 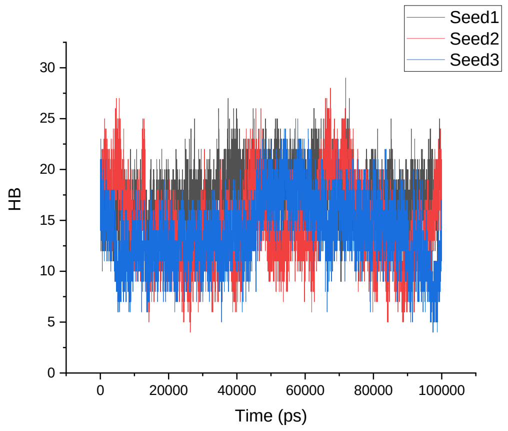 |
| CHARMM22* | 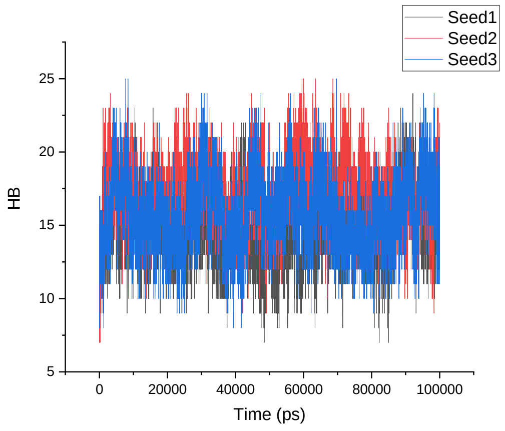 | 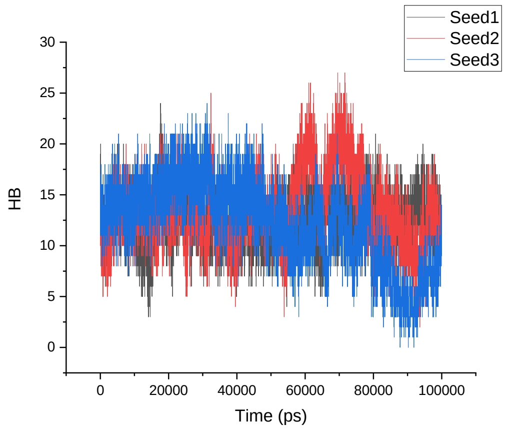 |
| CHARMM36 | 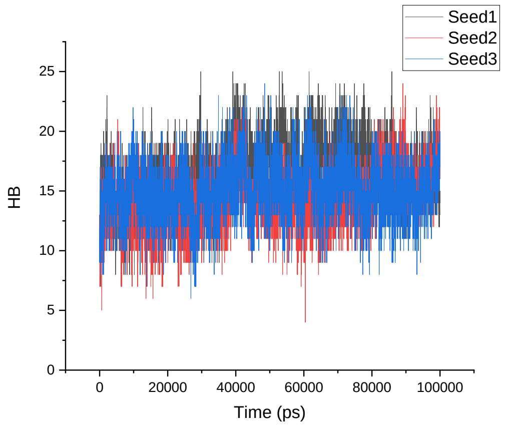 | 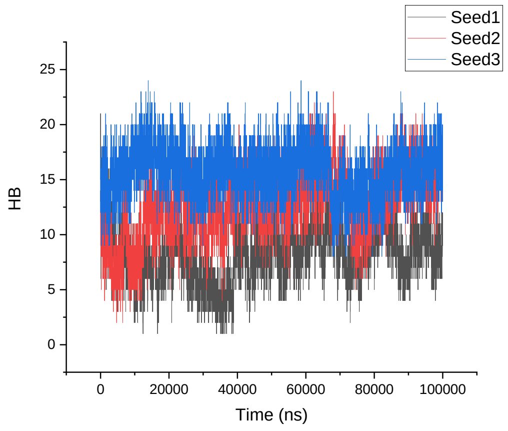 |

<table>
<tr><td>GROMOS 54A8</td><td>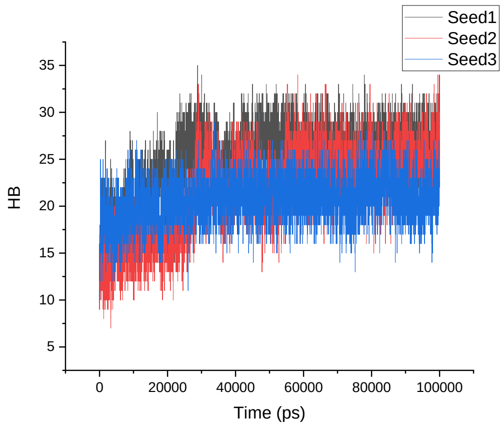</td><td>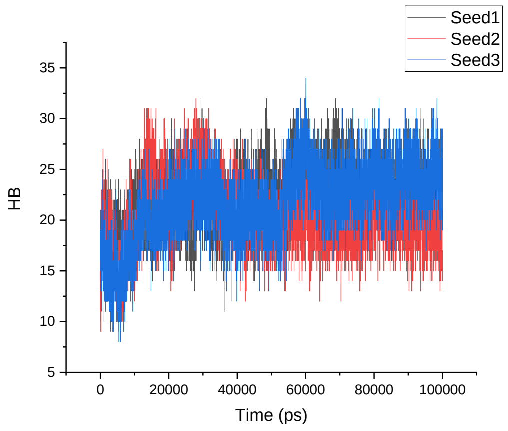</td></tr>
<tr><td></td><td>1M4F</td><td>α-1M4F</td></tr>
<tr><td>Amber 99SB-ILDN</td><td>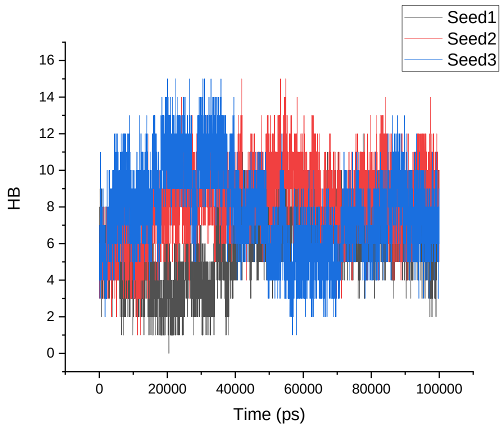</td><td>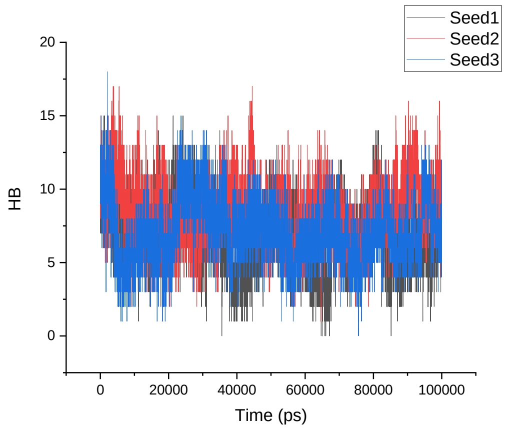</td></tr>
<tr><td>Amber 99SB*-ILDN</td><td>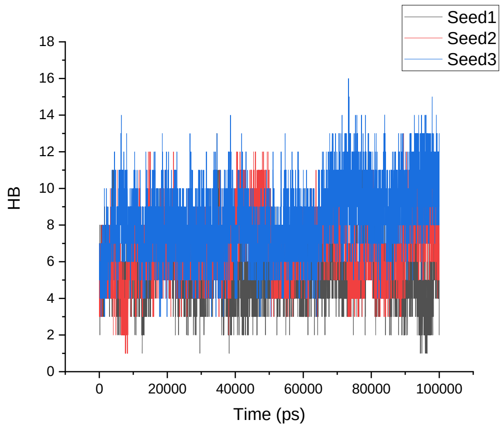</td><td>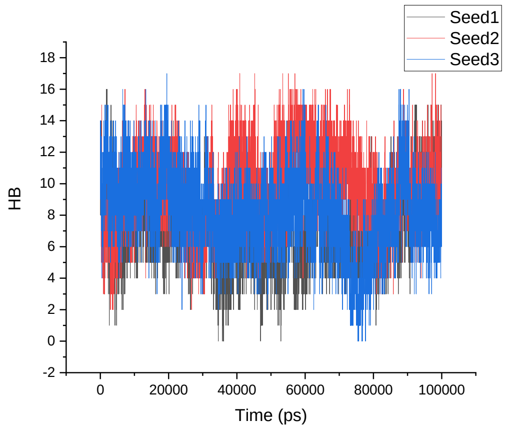</td></tr>
</table>

| | | |
|---|---|---|
| CHARMM22* | 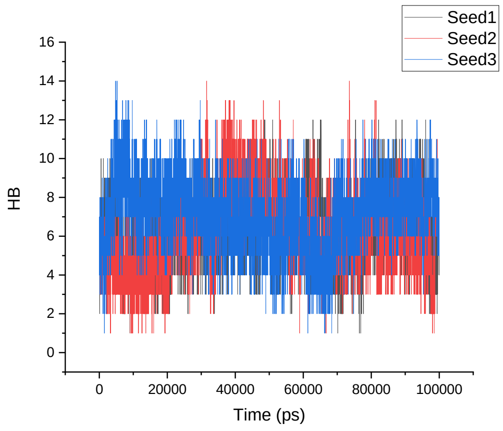 | 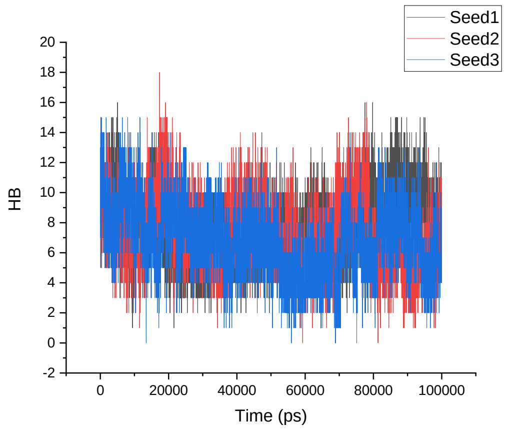 |
| CHARMM36 | 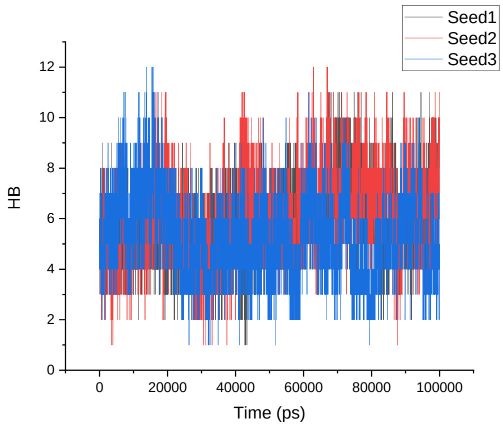 | 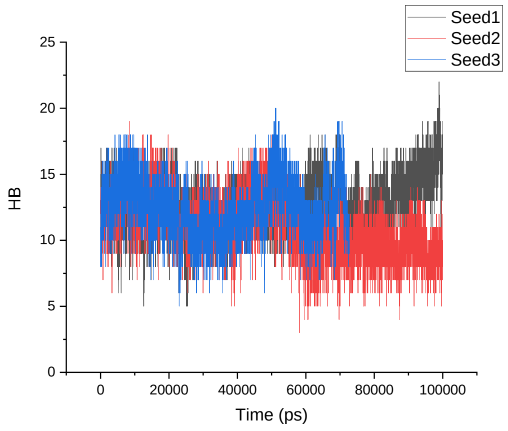 |
| GROMOS 54A8 | 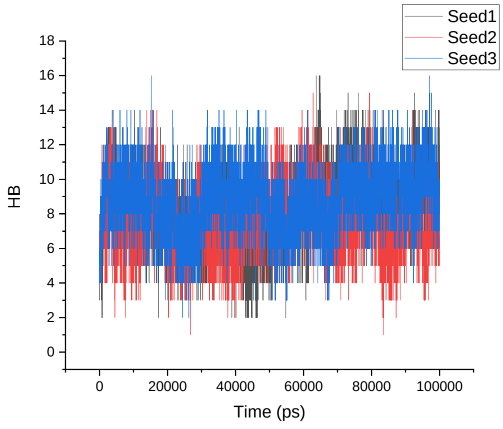 | 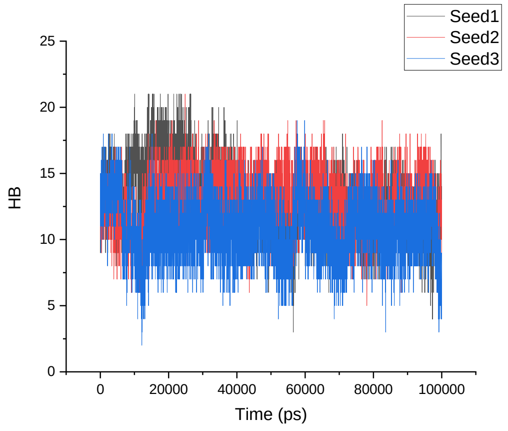 |
| | **1TV0** | **α-1TV0** |

<table>
<tr><td>Amber 99SB-ILDN</td><td>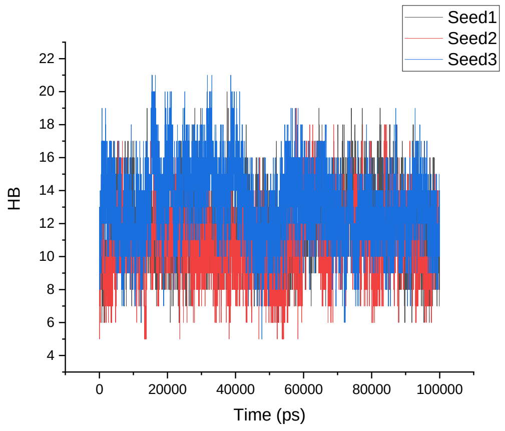
</td><td>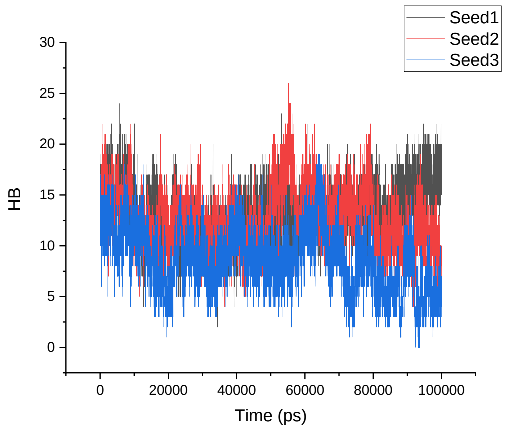
</td></tr>
<tr><td>Amber 99SB*-ILDN</td><td>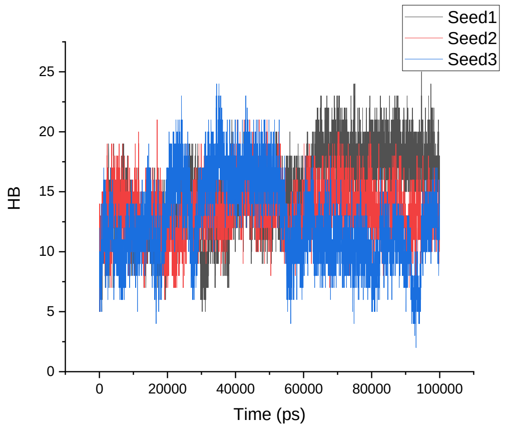
</td><td>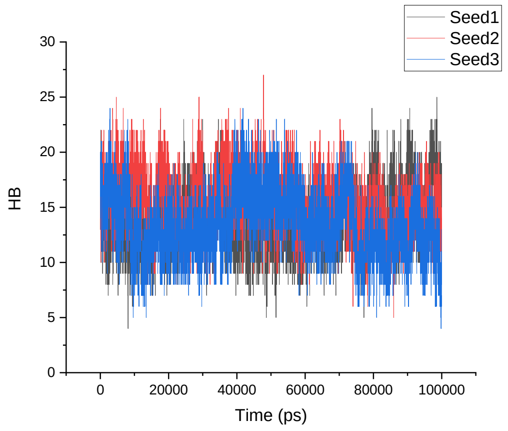
</td></tr>
<tr><td>CHARMM22*</td><td>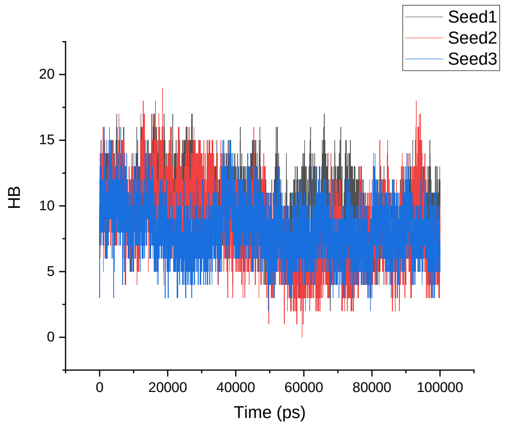
</td><td>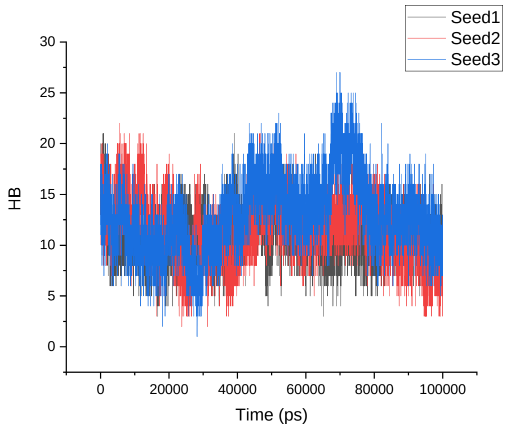
</td></tr>
</table>

| | | |
|---|---|---|
| CHARMM36 | 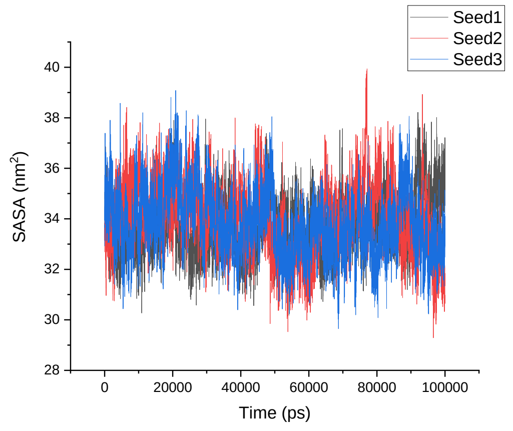 | 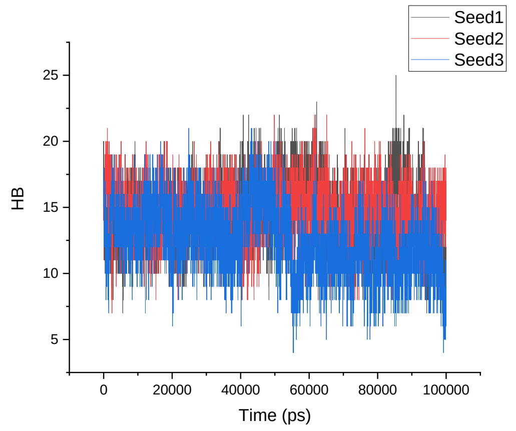 |
| GROMOS 54A8 | 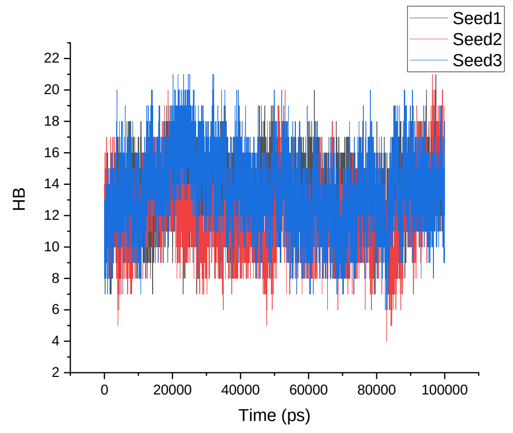 | 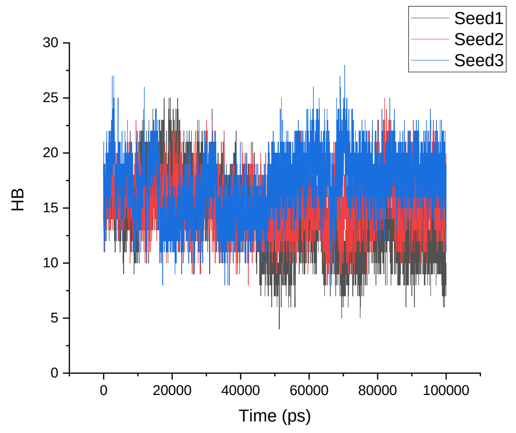 |

**Table S12**. Average number of intra-protein hydrogen bonds (averaged over three simulations for each protein) of the native α-proteins, modeled β-proteins, β-proteins, and modeled α-proteins.

| | HB | SD | HB | SD | HB | SD | HB | SD | HB | SD |
|---|---|---|---|---|---|---|---|---|---|---|
| α-proteins | | | | | | | | | | |
| | **1CQ0** | | **1L2Y** | | **1ROO** | | **1T5Q** | | **1V1D** | |
| Amber 99SB-ILDN | 14 | 3 | 12 | 2 | 19 | 3 | 15 | 3 | 17 | 2 |
| Amber 99SB*-ILDN | 16 | 3 | 13 | 2 | 20 | 3 | 17 | 3 | 17 | 2 |
| CHARMM22* | 13 | 3 | 10 | 2 | 21 | 3 | 16 | 3 | 15 | 2 |
| CHARMM36 | 15 | 2 | 11 | 2 | 17 | 3 | 22 | 3 | 17 | 2 |
| GROMOS 54A8 | 16 | 3 | 10 | 2 | 19 | 3 | 21 | 3 | 20 | 2 |
| Modelled β-proteins | | | | | | | | | | |
| | **β-1CQ0** | | **β-1L2Y** | | **β-1ROO** | | **β-1T5Q** | | **β-1V1D** | |
| Amber 99SB-ILDN | 10 | 2 | 5 | 2 | 14 | 3 | 13 | 3 | 15 | 3 |
| Amber 99SB*-ILDN | 10 | 3 | 5 | 3 | 14 | 4 | 13 | 3 | 16 | 3 |
| CHARMM22* | 8 | 3 | 5 | 2 | 12 | 3 | 10 | 3 | 14 | 3 |
| CHARMM36 | 6 | 3 | 3 | 2 | 8 | 2 | 8 | 3 | 11 | 2 |
| GROMOS 54A8 | 11 | 2 | 6 | 2 | 15 | 3 | 15 | 3 | 17 | 3 |
| β-proteins | | | | | | | | | | |
| | **1LFC** | | **8TFV** | | **1KGM** | | **1M4F** | | **1TV0** | |
| Amber 99SB-ILDN | 12 | 2 | 7 | 2 | 19 | 3 | 7 | 2 | 12 | 2 |
| Amber 99SB*-ILDN | 12 | 2 | 7 | 2 | 18 | 3 | 7 | 2 | 14 | 3 |
| CHARMM22* | 11 | 2 | 6 | 2 | 16 | 2 | 7 | 2 | 9 | 2 |
| CHARMM36 | 10 | 2 | 3 | 2 | 15 | 2 | 6 | 2 | 10 | 2 |
| GROMOS 54A8 | 9 | 2 | 6 | 2 | 22 | 3 | 8 | 2 | 13 | 2 |
| Modelled α-proteins | | | | | | | | | | |
| | **α-1LFC** | | **α-8TFV** | | **α-1KGM** | | **α-1M4F** | | **α-1TV0** | |
| Amber 99SB-ILDN | 11 | 3 | 8 | 2 | 17 | 3 | 7 | 2 | 12 | 3 |
| Amber 99SB*-ILDN | 12 | 2 | 9 | 3 | 15 | 3 | 8 | 2 | 14 | 3 |
| CHARMM22* | 9 | 2 | 6 | 2 | 13 | 3 | 7 | 2 | 12 | 3 |
| CHARMM36 | 10 | 2 | 5 | 2 | 12 | 3 | 12 | 2 | 14 | 2 |
| GROMOS 54A8 | 12 | 2 | 7 | 2 | 22 | 3 | 12 | 2 | 16 | 3 |

**Table S13**. Free energy landscapes of all 300 MD simulations.

| FEL | | | |
|---|---|---|---|
| **1CQ0** | | | |
| | Seed1 | Seed2 | Seed3 |
| **CHARMM22*** | 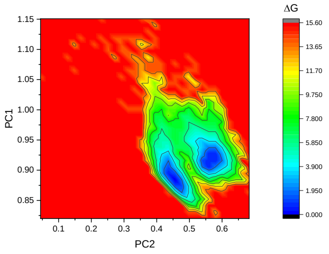 | 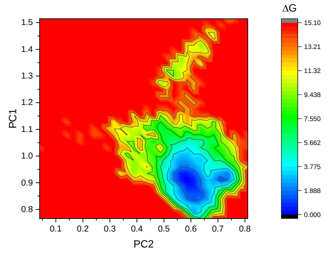 | 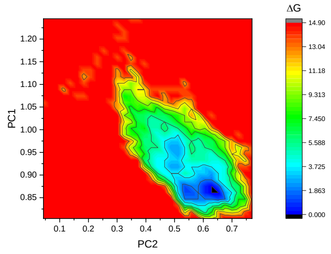 |
| **CHARMM36** | 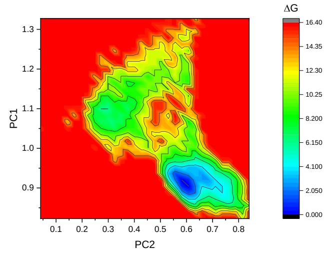 | 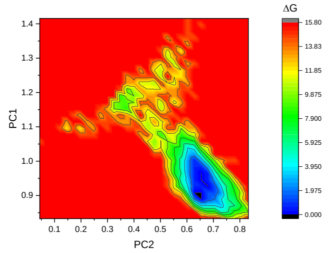 | 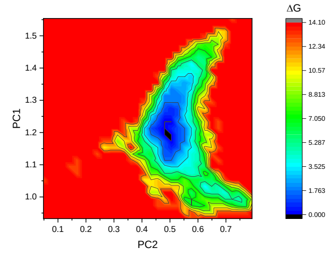 |
| **Amber 99SB-ILDN** | 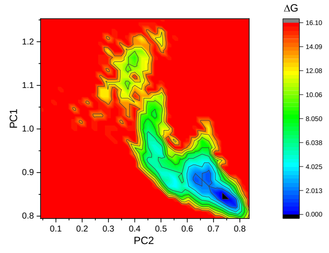 | 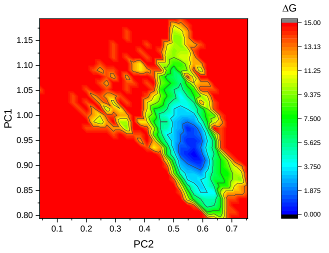 | 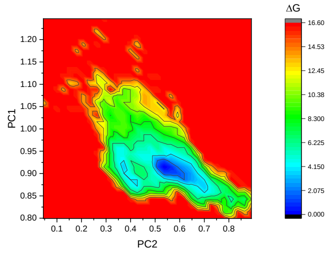 |
| **Amber 99SB-ILDN*** | 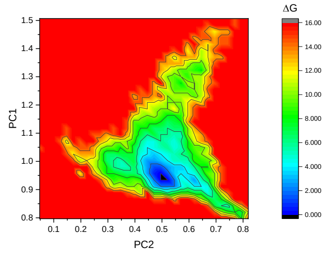 | 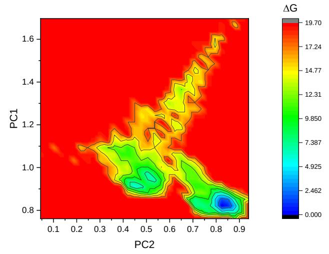 | 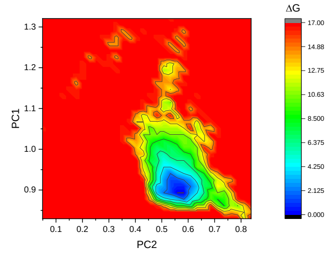 |
| **GROMOS 54A8** | 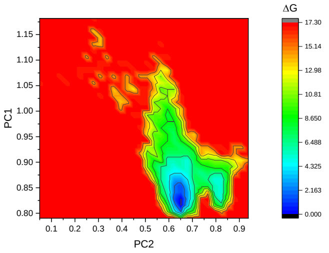 | 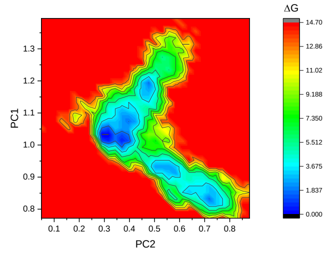 | 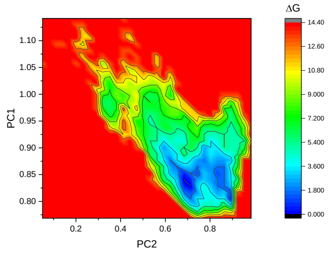 |

# β-1CQ0

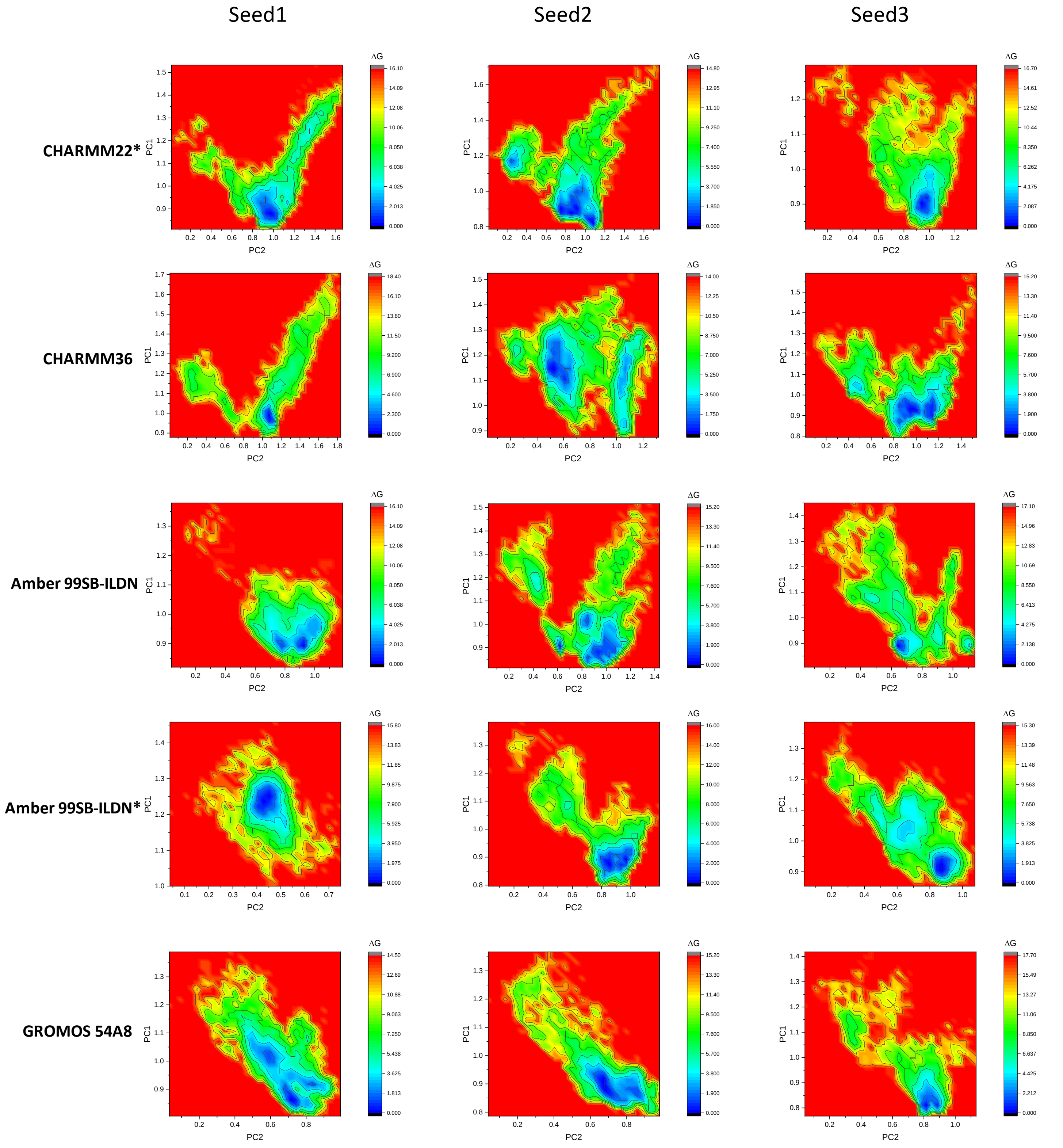
Seed1
Seed2
Seed3
CHARMM22*
CHARMM36
Amber 99SB-ILDN
Amber 99SB-ILDN*
GROMOS 54A8
PC1
PC2
ΔG

# 1L2Y

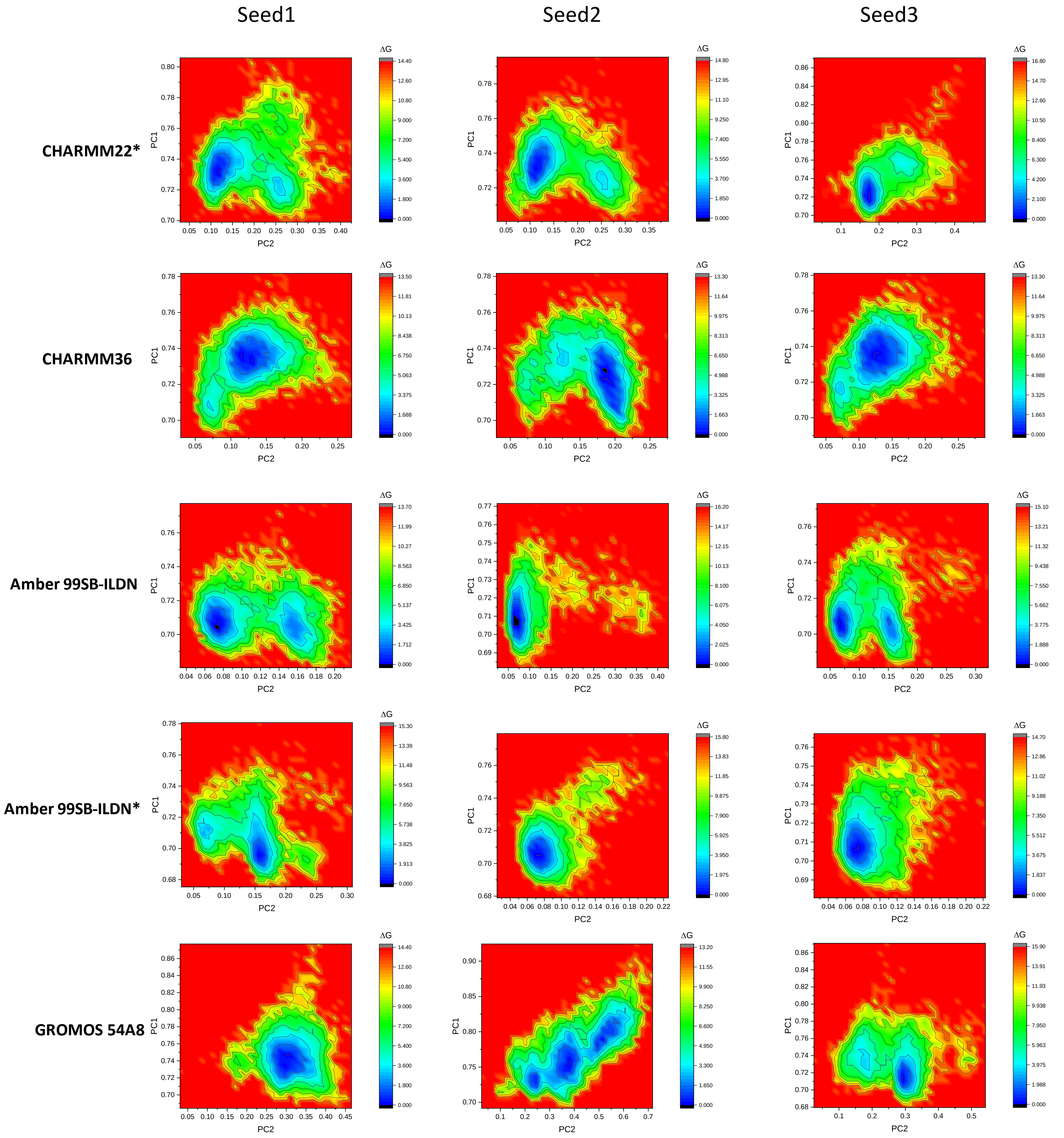
Seed1
Seed2
Seed3
CHARMM22*
CHARMM36
Amber 99SB-ILDN
Amber 99SB-ILDN*
GROMOS 54A8
PC1
PC2
ΔG

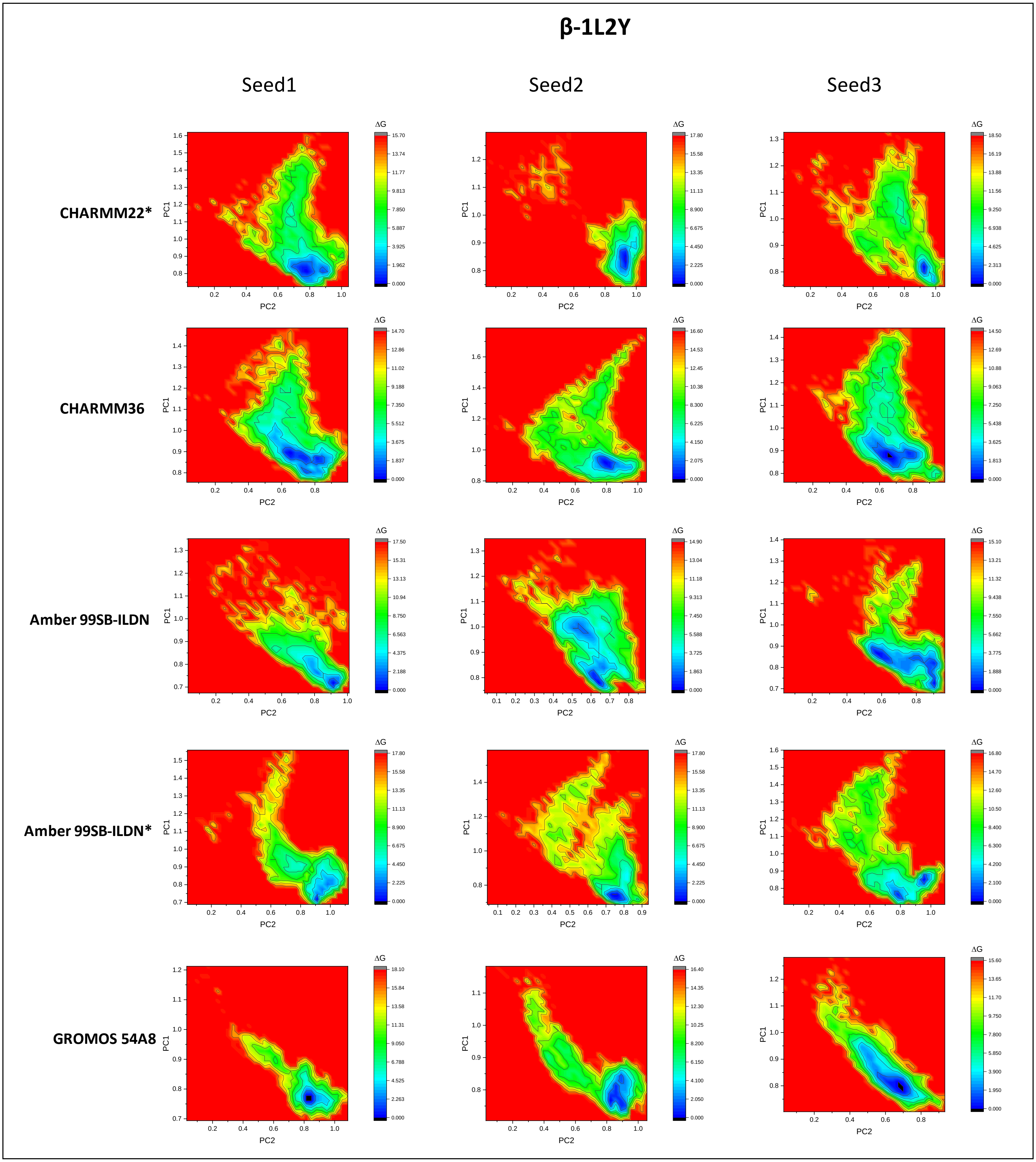
β-1L2Y
Seed1
Seed2
Seed3
CHARMM22*
CHARMM36
Amber 99SB-ILDN
Amber 99SB-ILDN*
GROMOS 54A8
PC1
PC2
ΔG

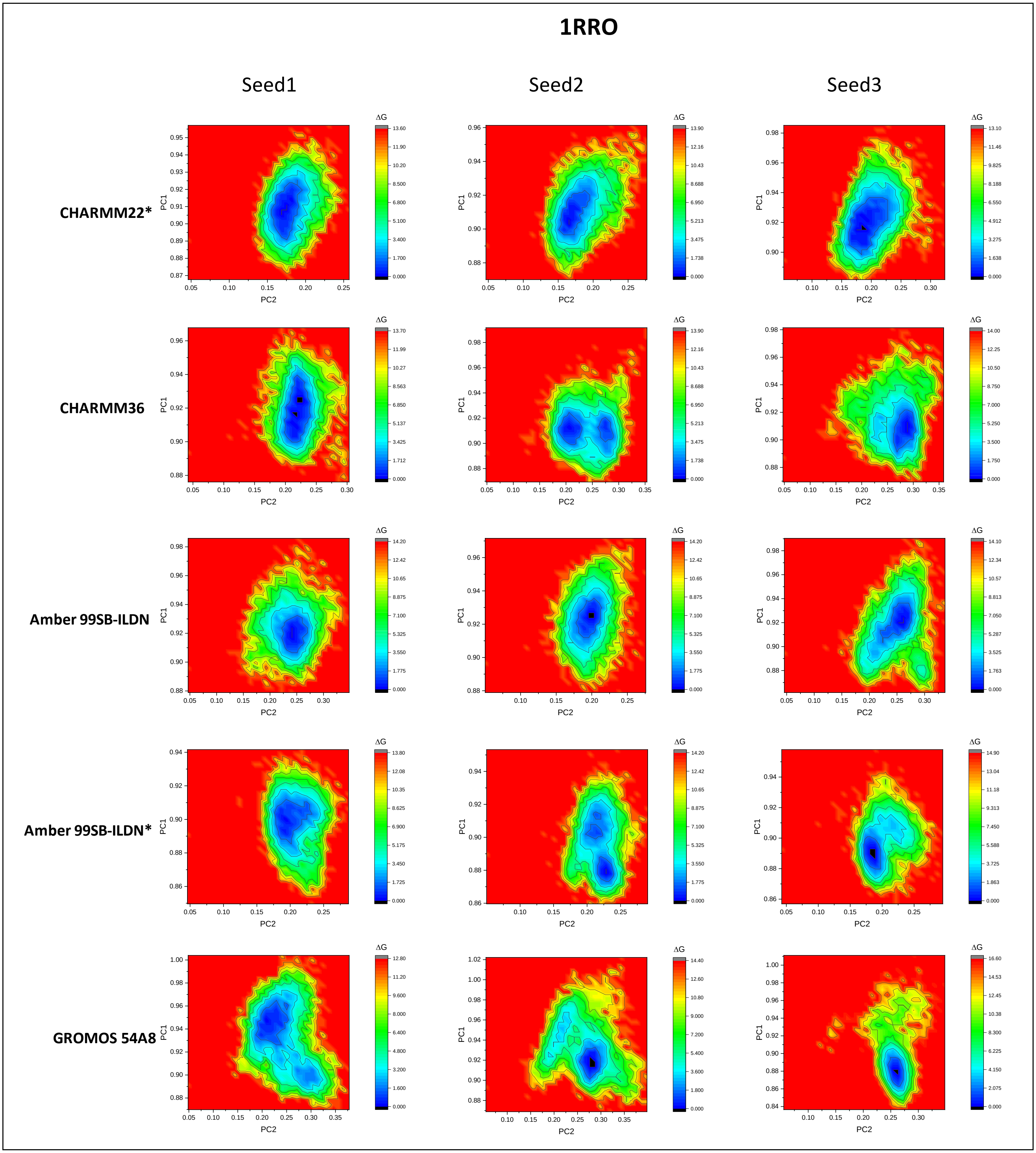
1RRO
Seed1
Seed2
Seed3
CHARMM22*
CHARMM36
Amber 99SB-ILDN
Amber 99SB-ILDN*
GROMOS 54A8
PC1
PC2
ΔG

### β-1RRO

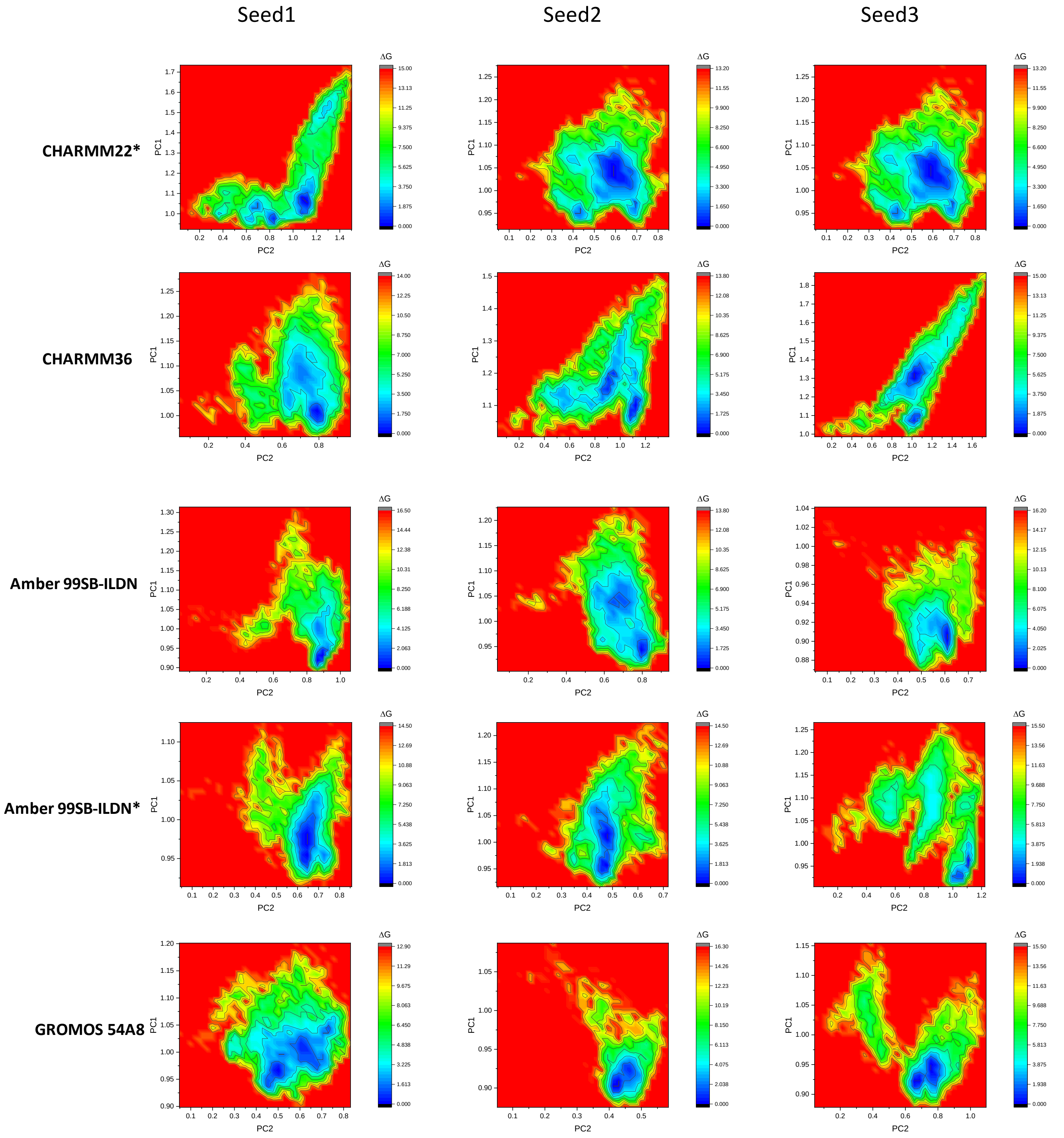
Seed1
Seed2
Seed3
CHARMM22*
CHARMM36
Amber 99SB-ILDN
Amber 99SB-ILDN*
GROMOS 54A8
PC1
PC2
ΔG

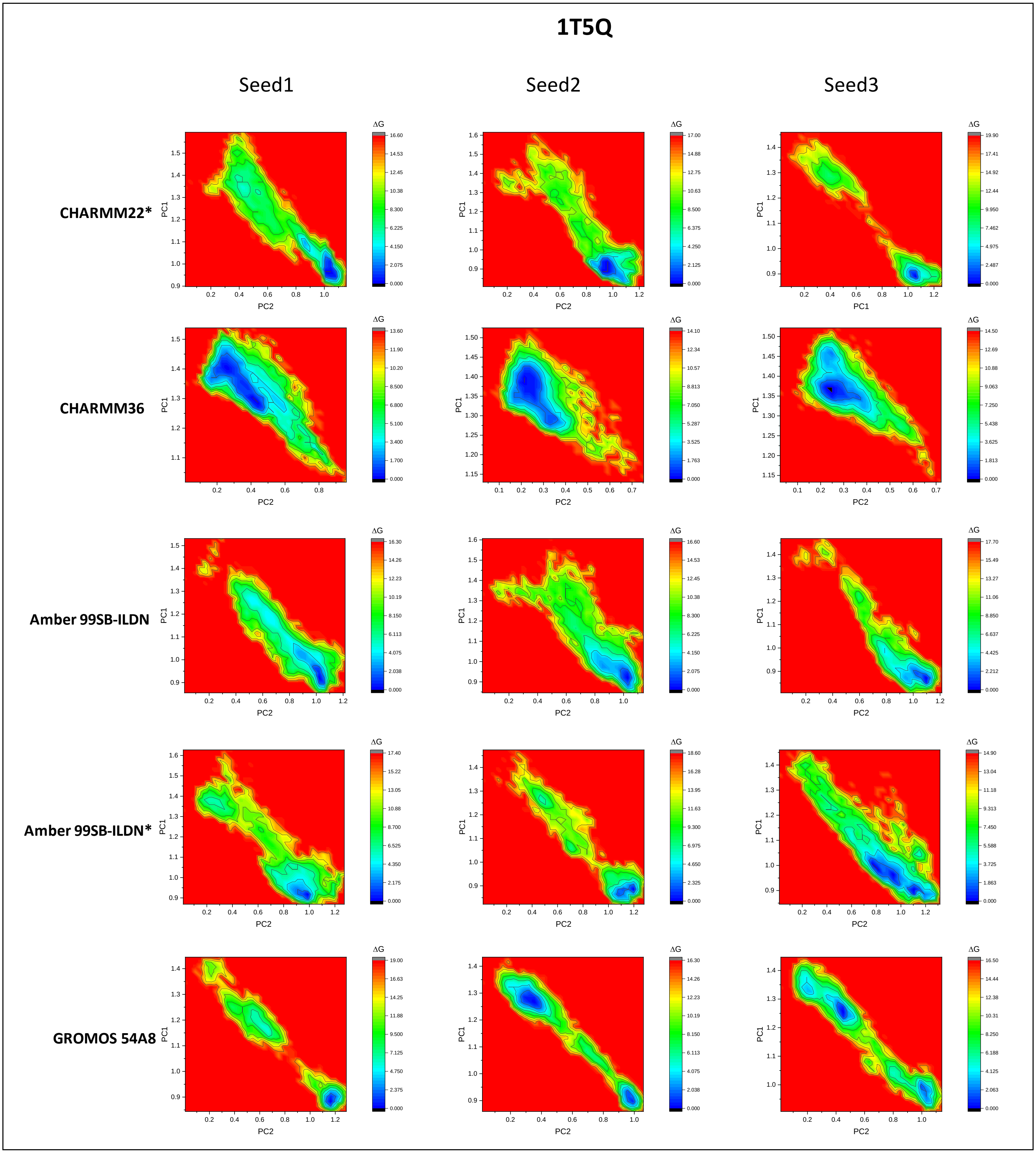
1T5Q
Seed1
Seed2
Seed3
CHARMM22*
CHARMM36
Amber 99SB-ILDN
Amber 99SB-ILDN*
GROMOS 54A8
PC1
PC2
ΔG

# β-1T5Q

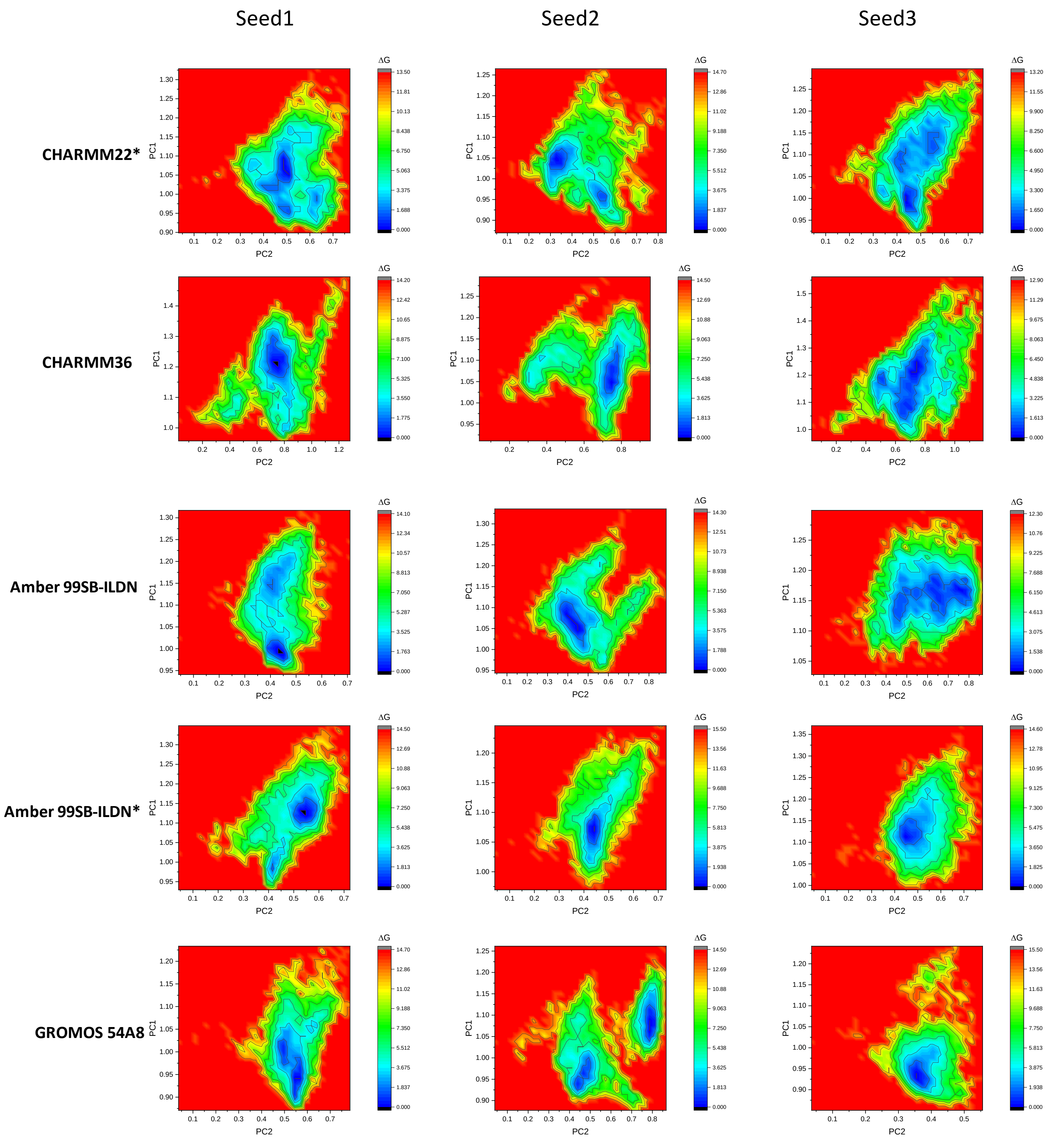

Seed1
Seed2
Seed3
CHARMM22*
CHARMM36
Amber 99SB-ILDN
Amber 99SB-ILDN*
GROMOS 54A8
PC1
PC2
ΔG

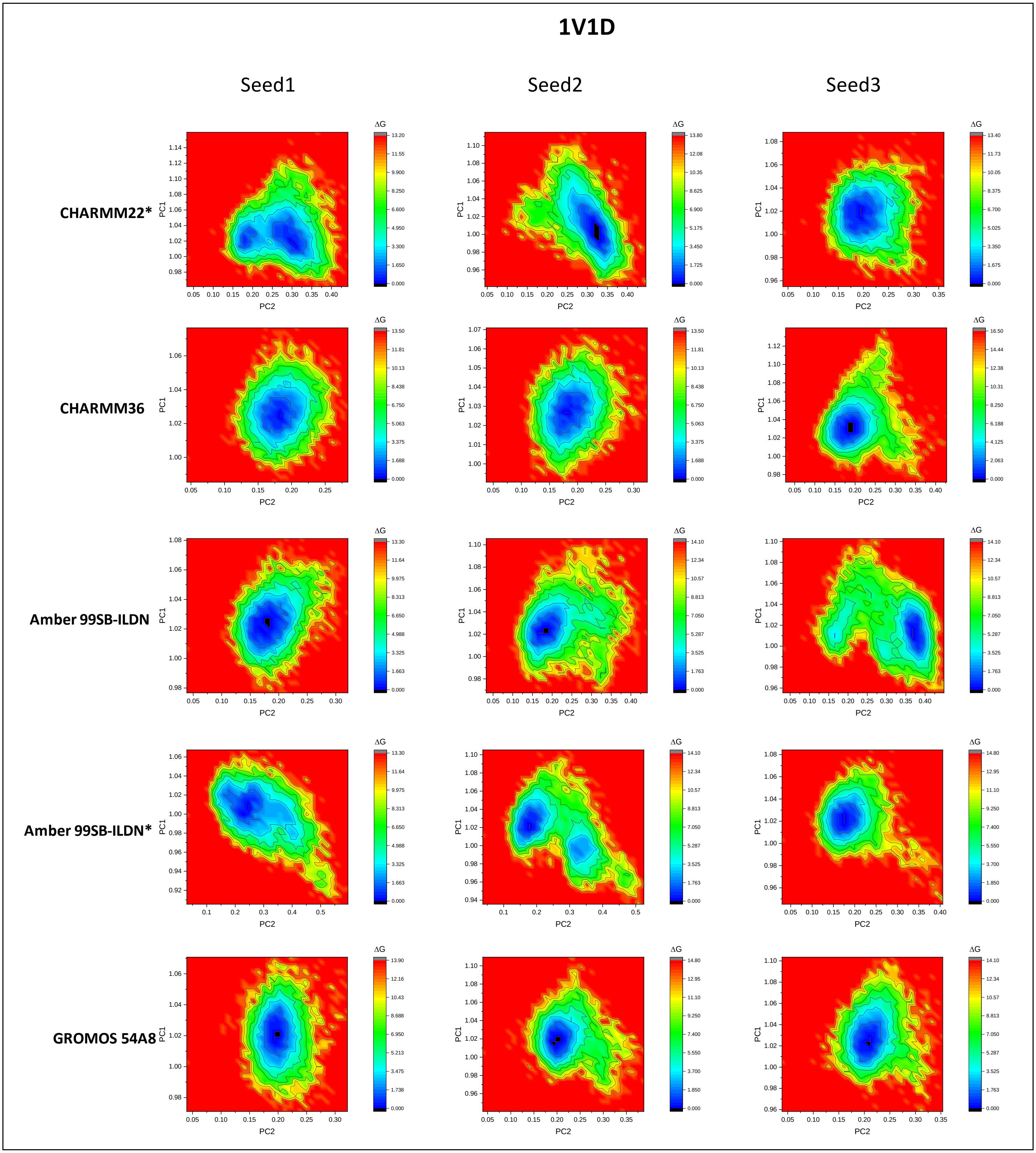

1V1D
Seed1
Seed2
Seed3
CHARMM22*
CHARMM36
Amber 99SB-ILDN
Amber 99SB-ILDN*
GROMOS 54A8
PC1
PC2
ΔG

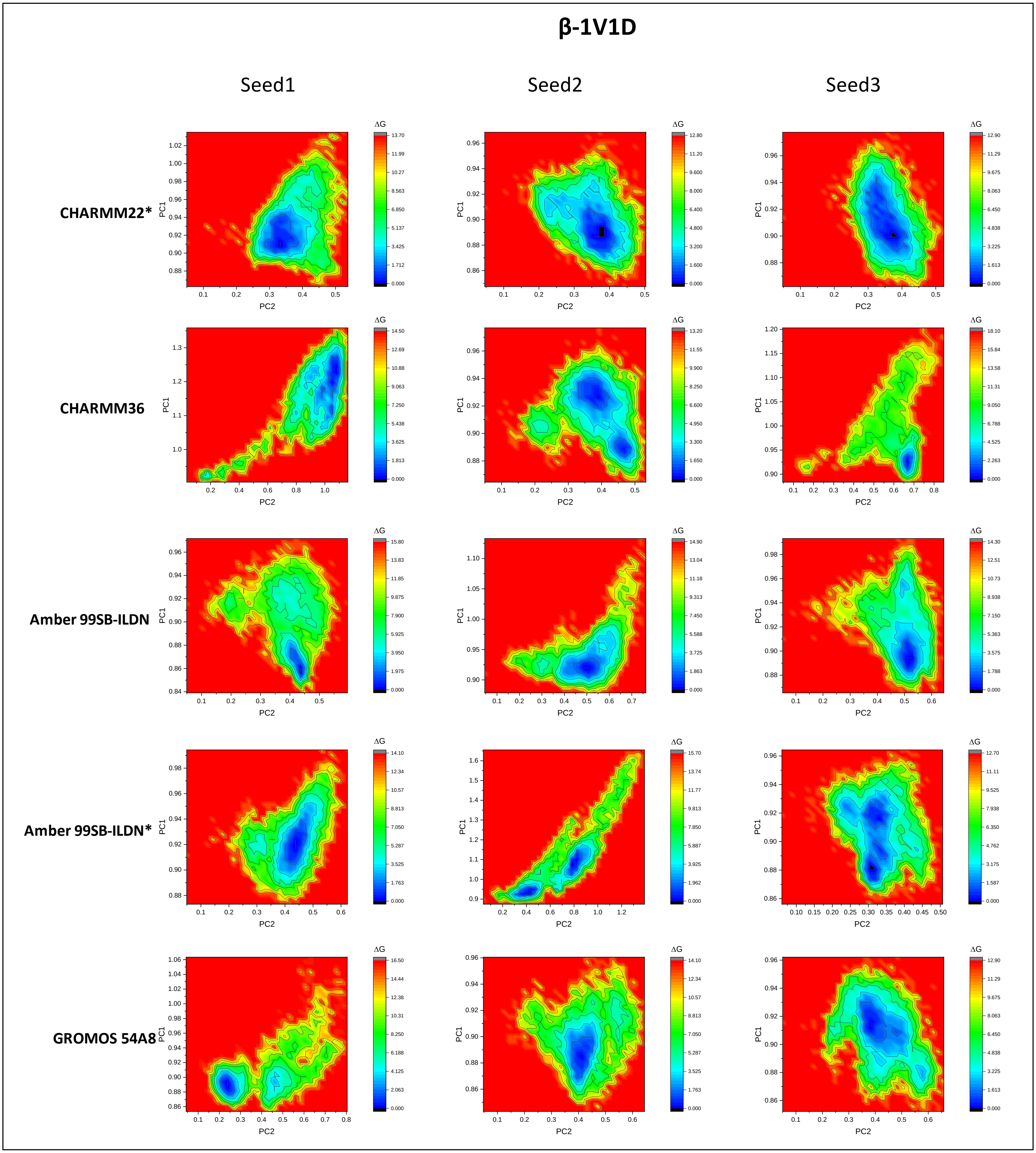
β-1V1D
Seed1
Seed2
Seed3
CHARMM22*
CHARMM36
Amber 99SB-ILDN
Amber 99SB-ILDN*
GROMOS 54A8
PC1
PC2
ΔG

### 1LFC

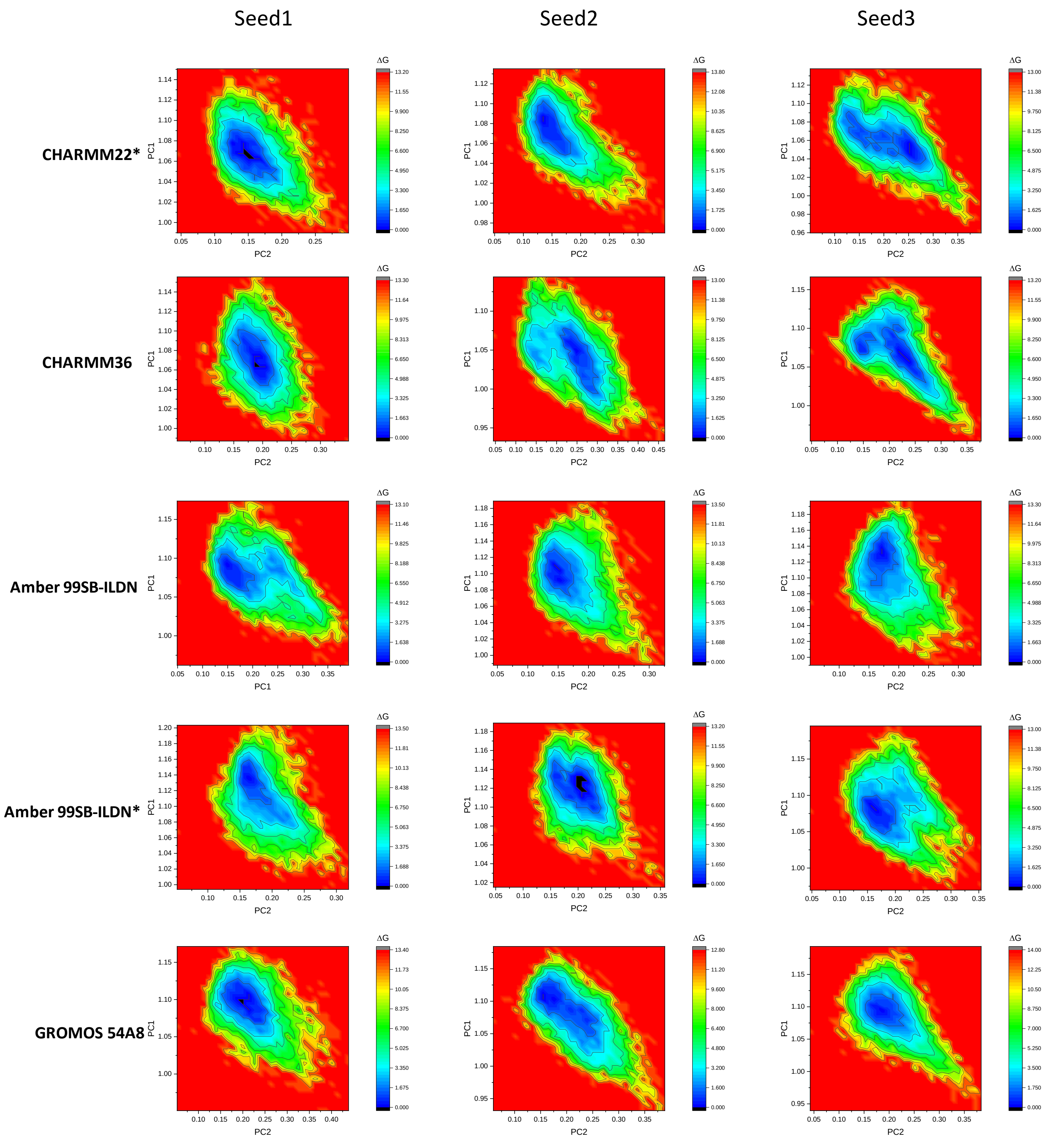

Seed1
Seed2
Seed3
CHARMM22*
CHARMM36
Amber 99SB-ILDN
Amber 99SB-ILDN*
GROMOS 54A8
PC1
PC2
ΔG

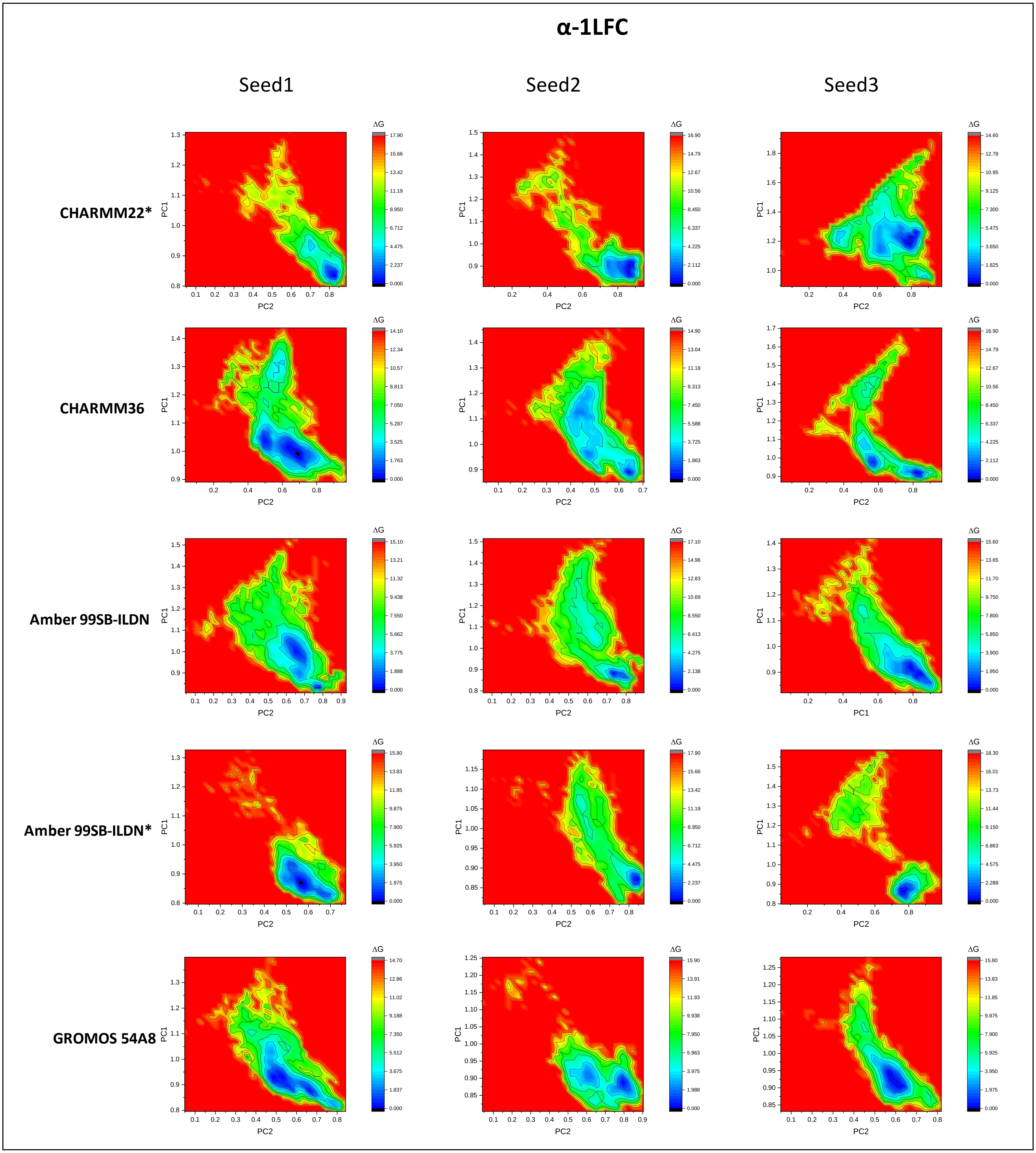
α-1LFC
Seed1
Seed2
Seed3
CHARMM22*
CHARMM36
Amber 99SB-ILDN
Amber 99SB-ILDN*
GROMOS 54A8
PC1
PC2
ΔG

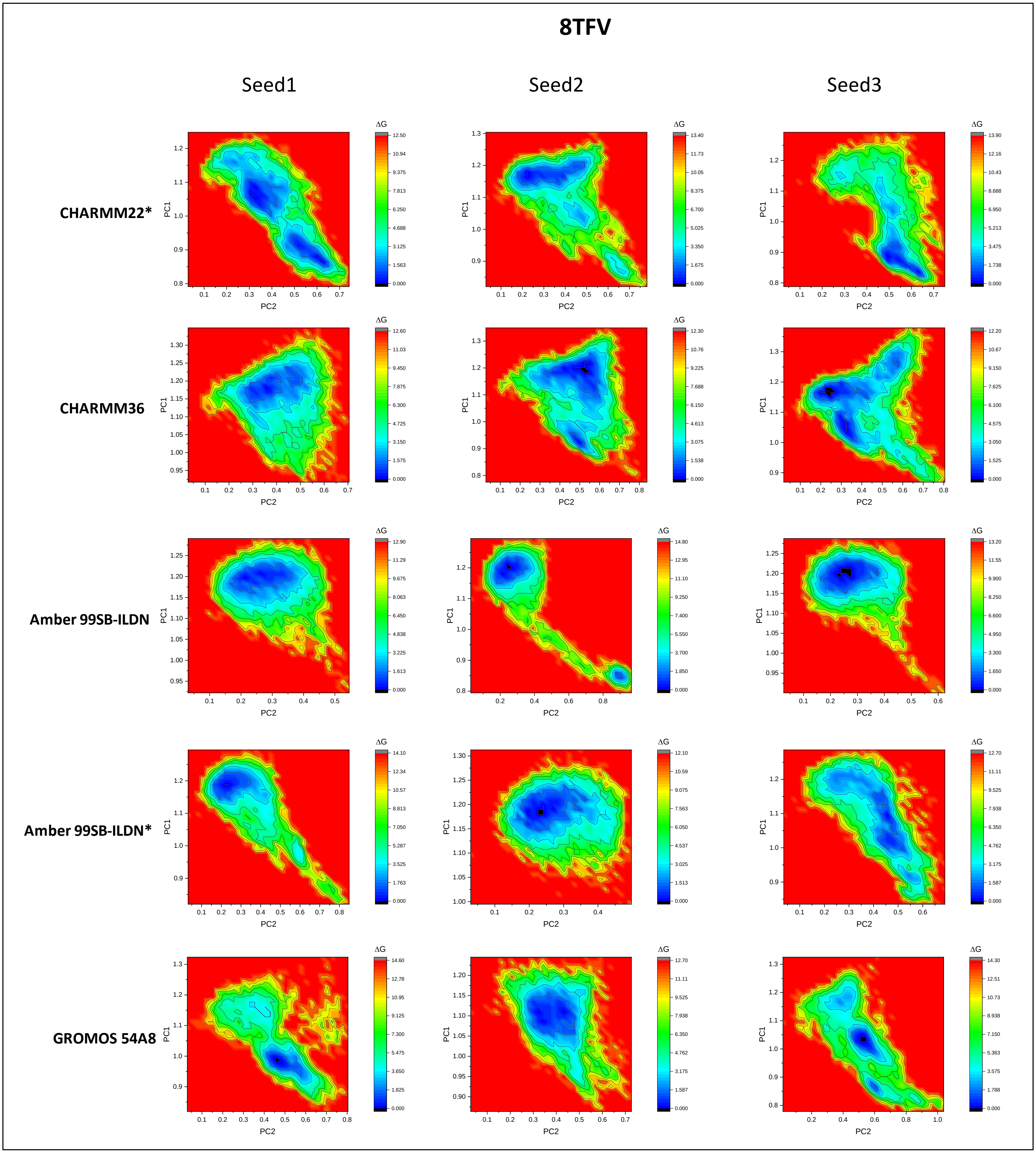
8TFV
Seed1
Seed2
Seed3
CHARMM22*
CHARMM36
Amber 99SB-ILDN
Amber 99SB-ILDN*
GROMOS 54A8
PC1
PC2
ΔG

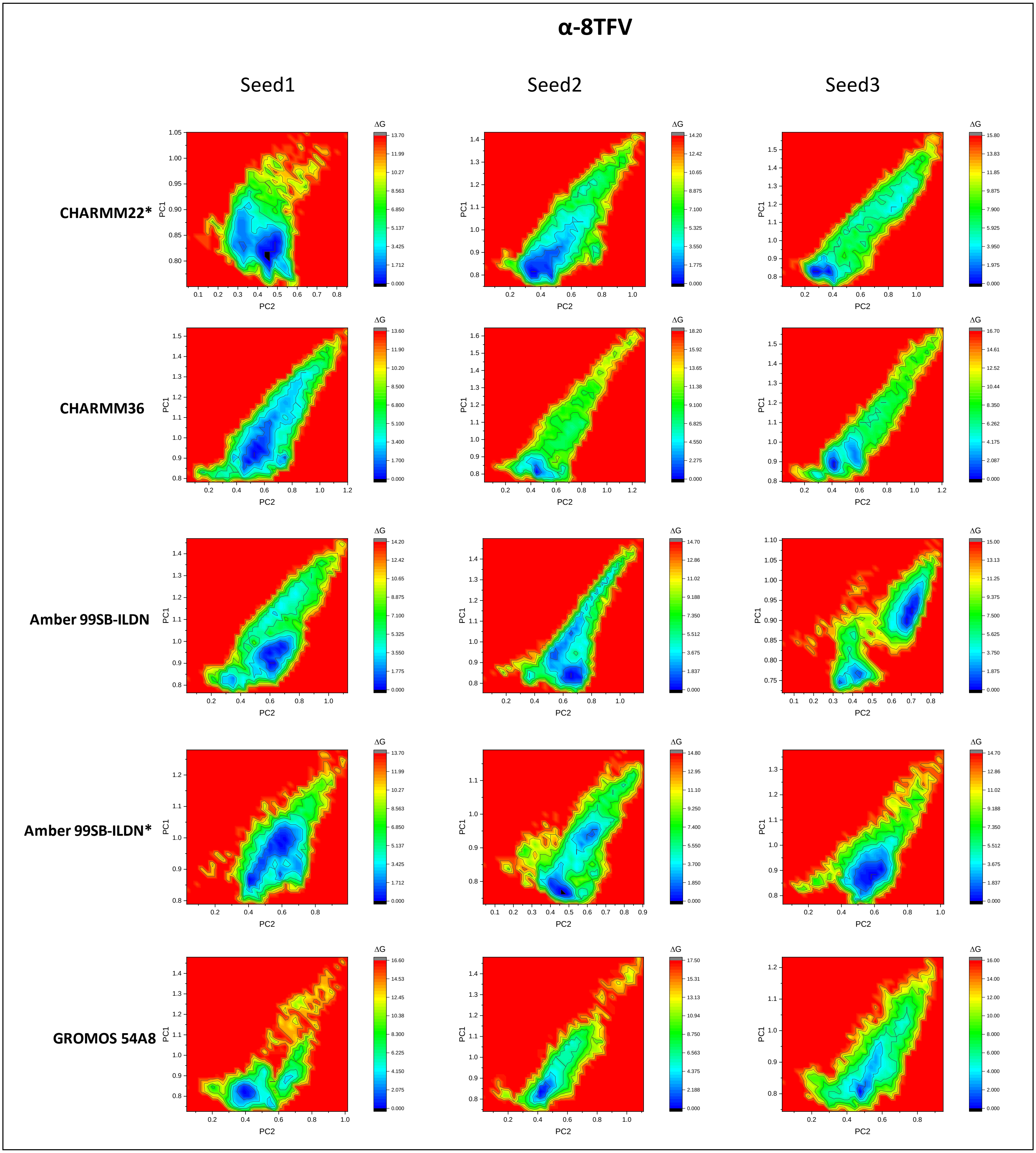
α-8TFV
Seed1
Seed2
Seed3
CHARMM22*
CHARMM36
Amber 99SB-ILDN
Amber 99SB-ILDN*
GROMOS 54A8
PC1
PC2
ΔG

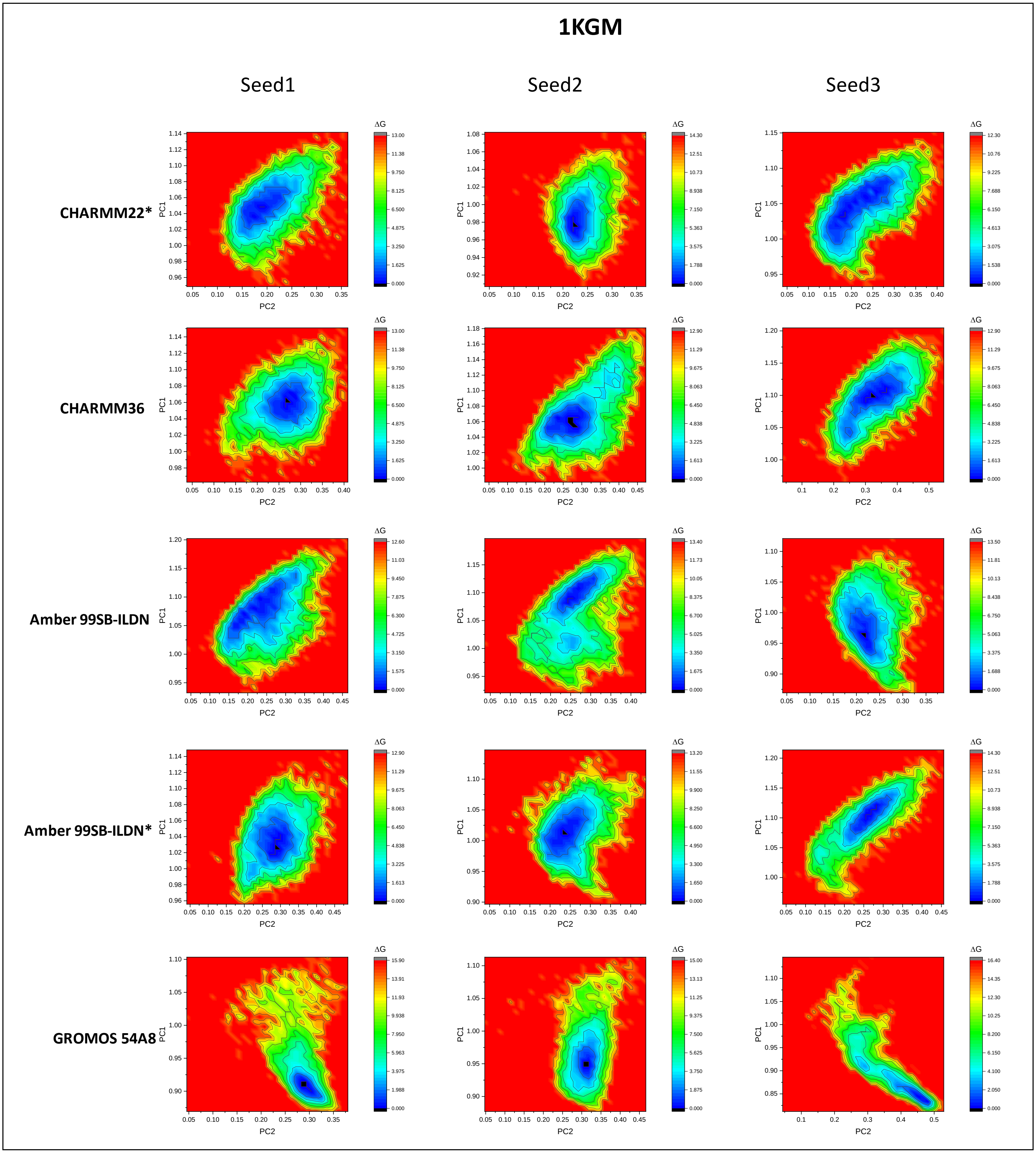
1KGM
Seed1
Seed2
Seed3
CHARMM22*
CHARMM36
Amber 99SB-ILDN
Amber 99SB-ILDN*
GROMOS 54A8
PC1
PC2
ΔG

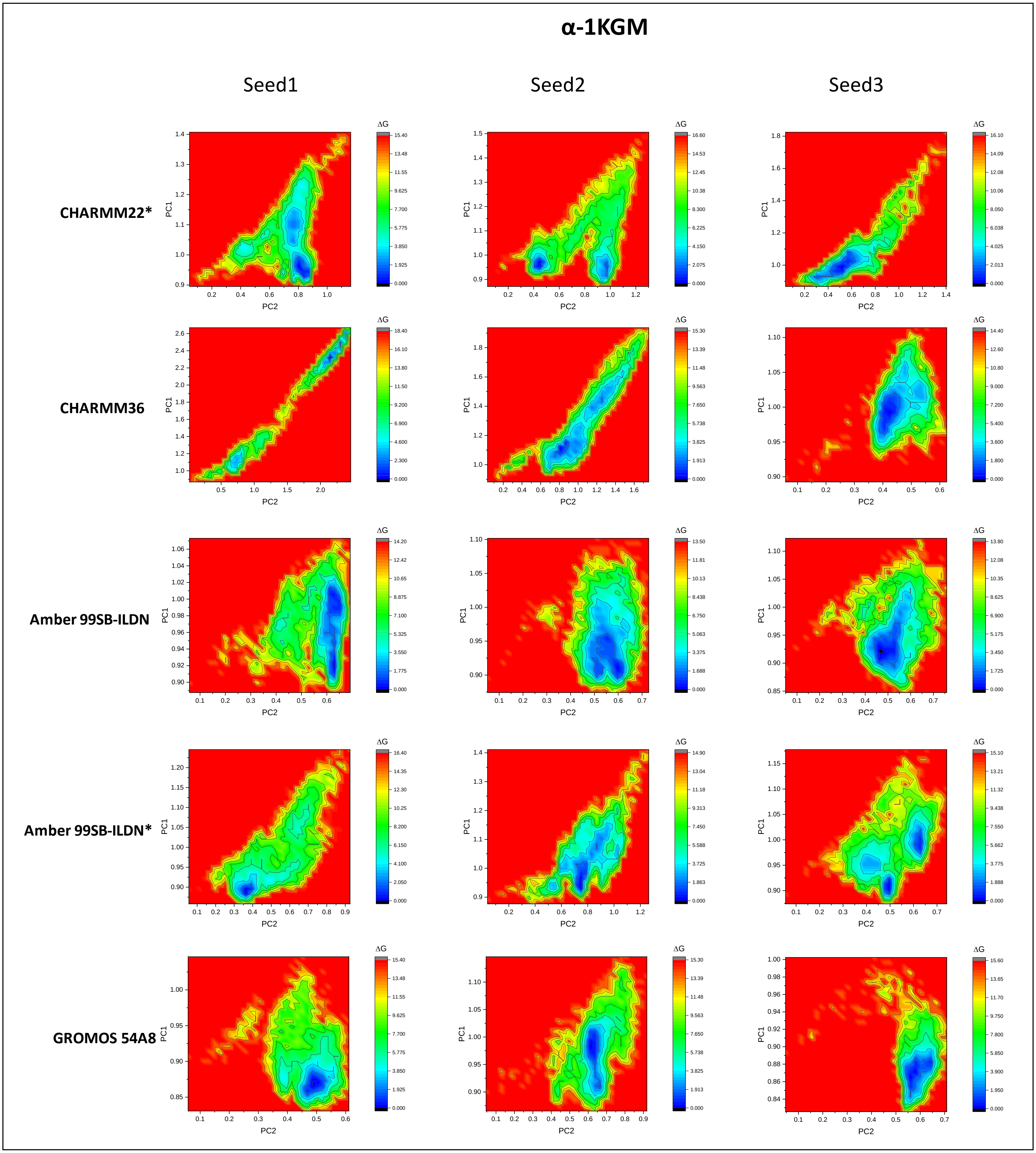
α-1KGM
Seed1
Seed2
Seed3
CHARMM22*
CHARMM36
Amber 99SB-ILDN
Amber 99SB-ILDN*
GROMOS 54A8
PC1
PC2
ΔG

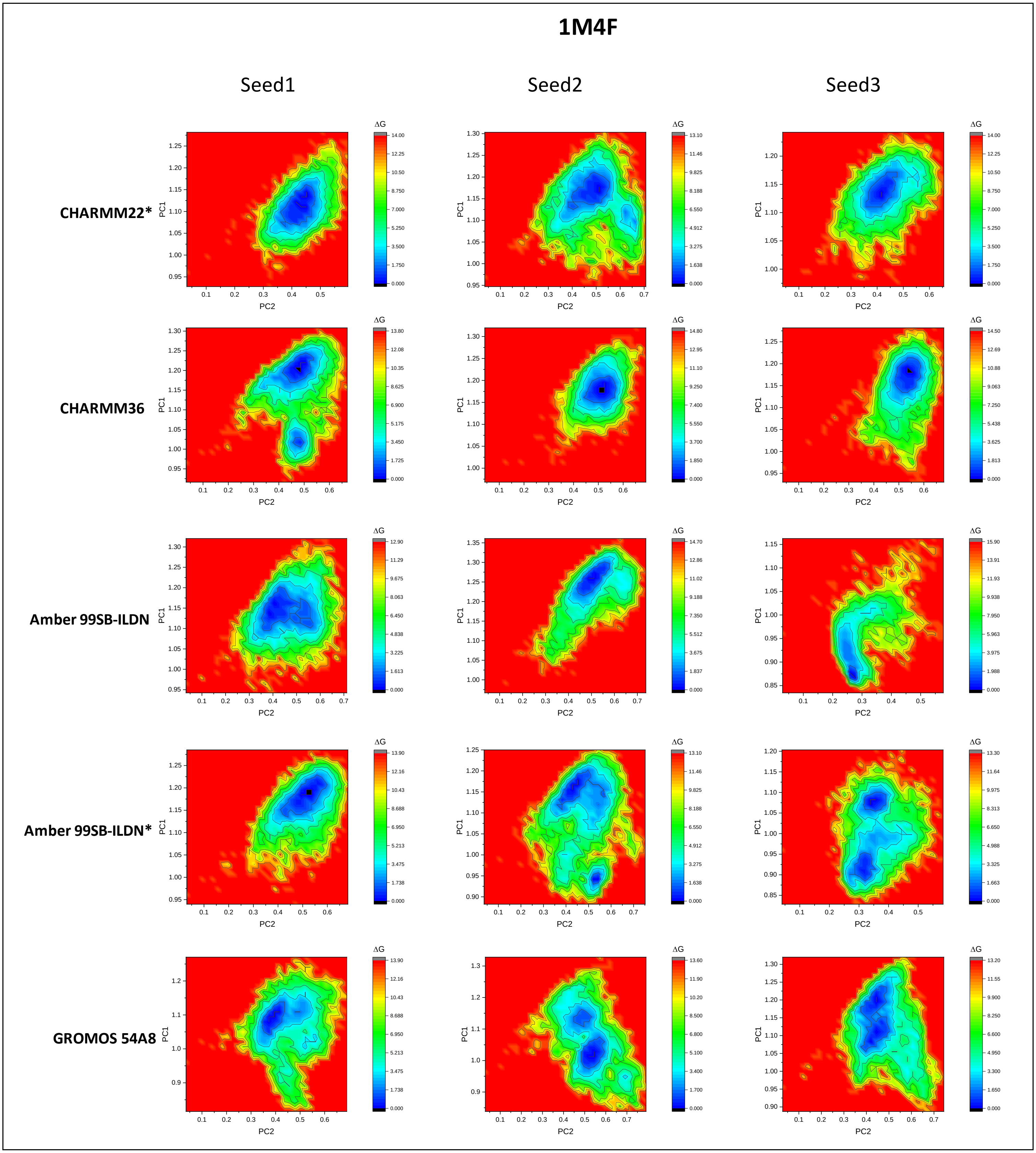

1M4F
Seed1
Seed2
Seed3
CHARMM22*
CHARMM36
Amber 99SB-ILDN
Amber 99SB-ILDN*
GROMOS 54A8
PC1
PC2
ΔG

## α-1M4F

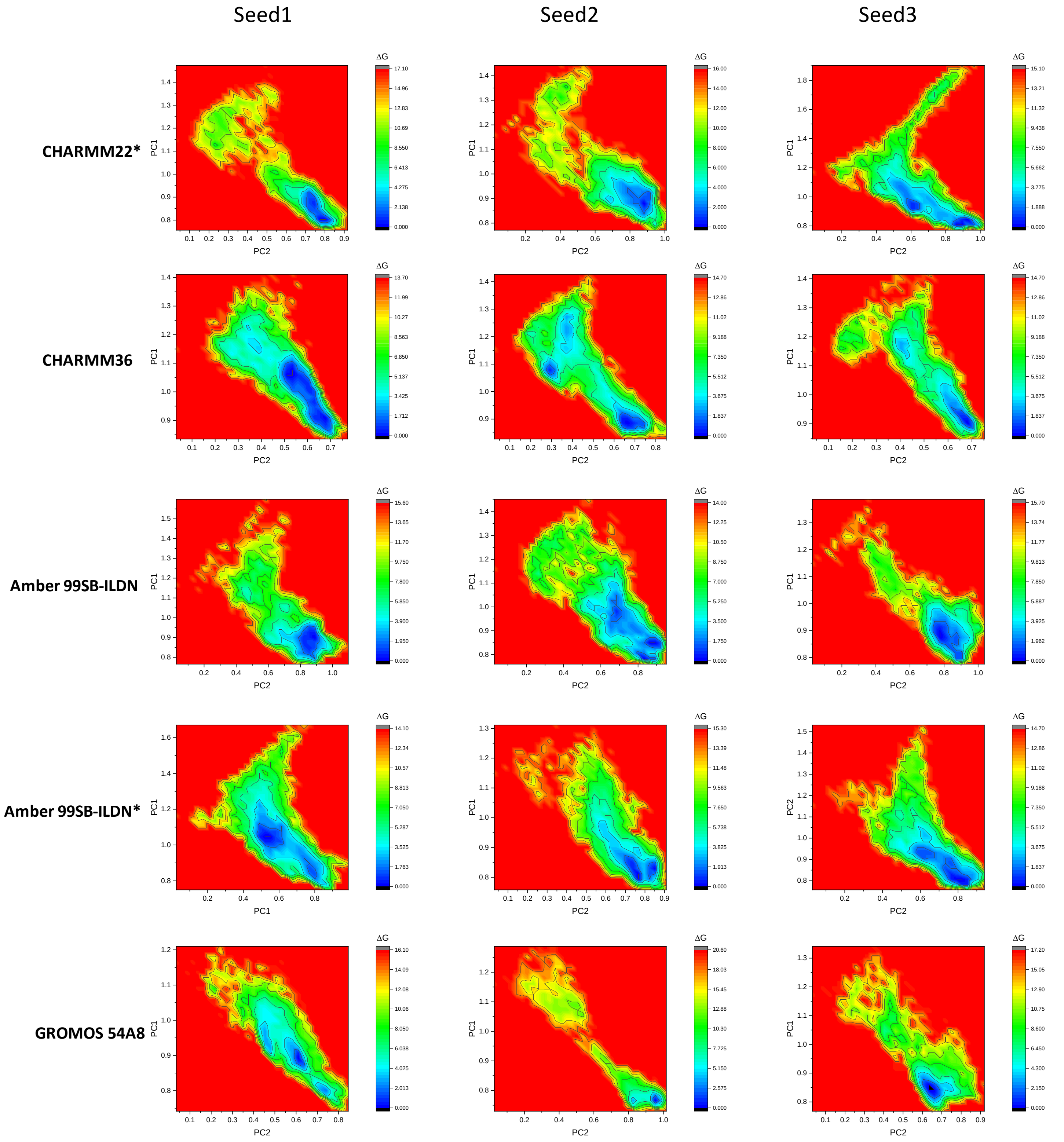

Seed1
Seed2
Seed3
CHARMM22*
CHARMM36
Amber 99SB-ILDN
Amber 99SB-ILDN*
GROMOS 54A8
PC1
PC2
ΔG

# 1TV0

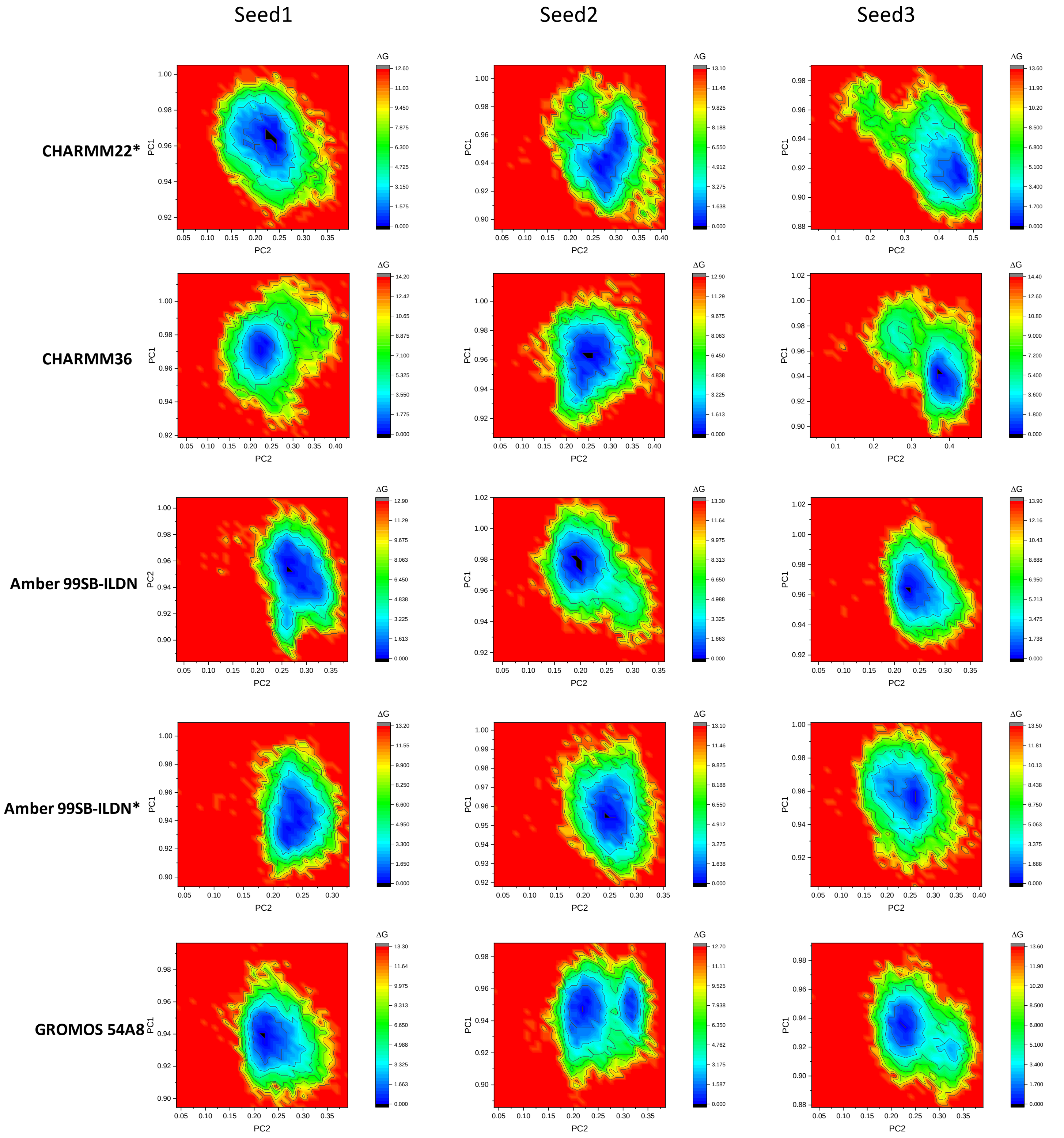

Seed1
Seed2
Seed3
CHARMM22*
CHARMM36
Amber 99SB-ILDN
Amber 99SB-ILDN*
GROMOS 54A8
PC1
PC2
ΔG

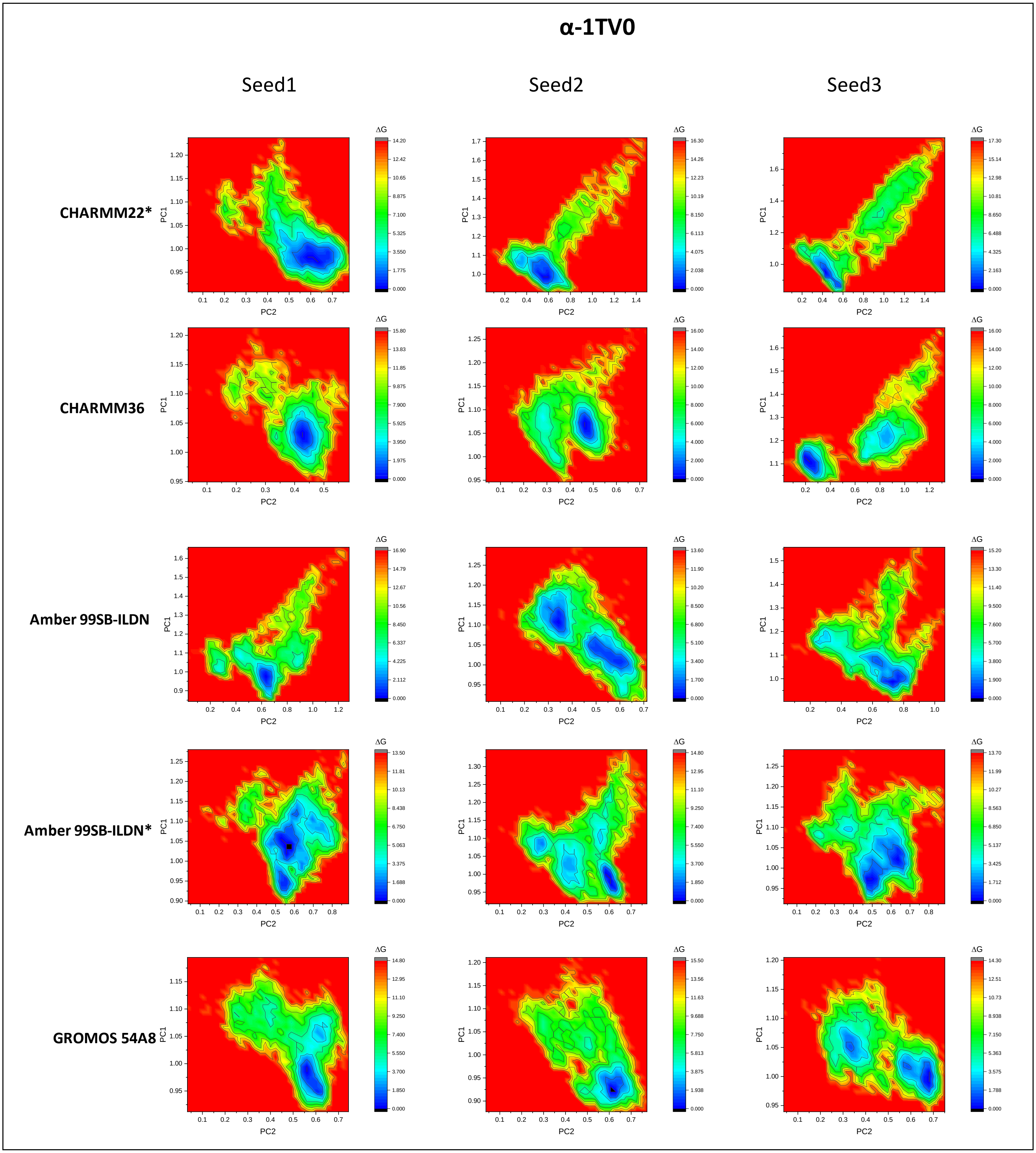
α-1TV0
Seed1
Seed2
Seed3
CHARMM22*
CHARMM36
Amber 99SB-ILDN
Amber 99SB-ILDN*
GROMOS 54A8
PC1
PC2
ΔG

**Table S14**. Average free energy landscape over three simulations of the native α-proteins, modeled β-proteins, β-proteins, and modeled α-proteins.

| α-proteins | | | | | |
|---|---|---|---|---|---|
| | **1CQ0** | **1L2Y** | **1ROO** | **1T5Q** | **1V1D** |
| Amber 99SB-ILDN | 0.008 | 0.001 | 0.002 | 0.008 | 0.003 |
| Amber 99SB*-ILDN | 0.006 | 0.001 | 0.001 | 0.013 | 0.005 |
| CHARMM22* | 0.012 | 0.001 | 0.002 | 0.008 | 0.005 |
| CHARMM36 | 0.015 | 0.002 | 0.002 | 0.022 | 0.002 |
| GROMOS 54A8 | 0.012 | 0.007 | 0.003 | 0.008 | 0.002 |
| Modelled β-proteins | | | | | |
| | **β-1CQ0** | **β-1L2Y** | **β-1ROO** | **β-1T5Q** | **β-1V1D** |
| Amber 99SB-ILDN | 0.014 | 0.016 | 0.008 | 0.020 | 0.003 |
| Amber 99SB*-ILDN | 0.011 | 0.005 | 0.007 | 0.008 | 0.011 |
| CHARMM22* | 0.027 | 0.009 | 0.017 | 0.020 | 0.006 |
| CHARMM36 | 0.028 | 0.021 | 0.029 | 0.037 | 0.009 |
| GROMOS 54A8 | 0.017 | 0.010 | 0.012 | 0.010 | 0.004 |
| β-proteins | | | | | |
| | **1LFC** | **8TFV** | **1KGM** | **1M4F** | **1TV0** |
| Amber 99SB-ILDN | 0.005 | 0.020 | 0.009 | 0.014 | 0.003 |
| Amber 99SB*-ILDN | 0.005 | 0.026 | 0.008 | 0.018 | 0.003 |
| CHARMM22* | 0.005 | 0.028 | 0.007 | 0.014 | 0.004 |
| CHARMM36 | 0.007 | 0.051 | 0.009 | 0.012 | 0.003 |
| GROMOS 54A8 | 0.007 | 0.021 | 0.002 | 0.021 | 0.003 |
| Modelled α-proteins | | | | | |
| | **α-1LFC** | **α-8TFV** | **α-1KGM** | **α-1M4F** | **α-1TV0** |
| Amber 99SB-ILDN | 0.011 | 0.025 | 0.010 | 0.026 | 0.017 |
| Amber 99SB*-ILDN | 0.007 | 0.025 | 0.009 | 0.030 | 0.018 |
| CHARMM22* | 0.016 | 0.021 | 0.017 | 0.020 | 0.018 |
| CHARMM36 | 0.015 | 0.026 | 0.022 | 0.015 | 0.007 |
| GROMOS 54A8 | 0.012 | 0.007 | 0.005 | 0.003 | 0.006 |

**Table S15**. Number of the clusters obtained by clustering analyses of the native α-proteins, modeled β-proteins, β-proteins, and modeled α-proteins.

| α-proteins | | | | | | |
|---|---|---|---|---|---|---|
| | | **1CQ0** | **1L2Y** | **1ROO** | **1T5Q** | **1V1D** |
| Amber 99SB-ILDN | Seed1 | 267 | 8 | 55 | 530 | 69 |
| | Seed2 | 457 | 9 | 34 | 702 | 121 |
| | Seed3 | 287 | 9 | 50 | 300 | 146 |
| Amber 99SB*-ILDN | Seed1 | 620 | 14 | 22 | 621 | 127 |
| | Seed2 | 359 | 3 | 23 | 267 | 125 |
| | Seed3 | 176 | 6 | 16 | 527 | 74 |
| CHARMM22* | Seed1 | 135 | 44 | 13 | 571 | 276 |
| | Seed2 | 627 | 15 | 17 | 571 | 193 |
| | Seed3 | 334 | 28 | 54 | 295 | 143 |
| CHARMM36 | Seed1 | 240 | 6 | 28 | 398 | 19 |
| | Seed2 | 202 | 15 | 30 | 226 | 22 |
| | Seed3 | 774 | 13 | 37 | 209 | 88 |
| GROMOS 54A8 | Seed1 | 300 | 159 | 131 | 229 | 101 |
| | Seed2 | 535 | 190 | 103 | 207 | 164 |
| | Seed3 | 488 | 96 | 38 | 403 | 194 |
| Modelled β-proteins | | | | | | |
| | | **β-1CQ0** | **β-1L2Y** | **β-1ROO** | **β-1T5Q** | **β-1V1D** |
| Amber 99SB-ILDN | Seed1 | 488 | 468 | 433 | 578 | 108 |
| | Seed2 | 765 | 760 | 581 | 547 | 223 |
| | Seed3 | 499 | 525 | 120 | 697 | 190 |
| Amber 99SB*-ILDN | Seed1 | 392 | 488 | 312 | 520 | 103 |
| | Seed2 | 572 | 645 | 254 | 252 | 677 |
| | Seed3 | 707 | 983 | 628 | 733 | 124 |
| CHARMM22* | Seed1 | 1149 | 1135 | 880 | 651 | 150 |
| | Seed2 | 1030 | 344 | 474 | 543 | 139 |
| | Seed3 | 771 | 843 | 196 | 677 | 162 |
| CHARMM36 | Seed1 | 1439 | 1128 | 498 | 1044 | 752 |
| | Seed2 | 1271 | 970 | 176 | 746 | 125 |
| | Seed3 | 784 | 1176 | 330 | 1569 | 269 |
| GROMOS 54A8 | Seed1 | 665 | 263 | 229 | 374 | 193 |
| | Seed2 | 626 | 324 | 176 | 591 | 180 |
| | Seed3 | 487 | 489 | 330 | 419 | 168 |
| β-proteins | | | | | | |
| | | **1LFC** | **8TFV** | **1KGM** | **1M4F** | **1TV0** |
| Amber 99SB-ILDN | Seed1 | 169 | 1115 | 172 | 632 | 145 |
| | Seed2 | 186 | 1038 | 164 | 743 | 197 |
| | Seed3 | 173 | 897 | 64 | 126 | 146 |
| Amber 99SB*-ILDN | Seed1 | 199 | 1086 | 97 | 590 | 92 |
| | Seed2 | 139 | 1064 | 140 | 568 | 119 |
| | Seed3 | 204 | 806 | 97 | 314 | 180 |
| CHARMM22* | Seed1 | 77 | 784 | 65 | 433 | 231 |
| | Seed2 | 94 | 840 | 45 | 571 | 158 |
| | Seed3 | 81 | 894 | 103 | 415 | 132 |
| CHARMM36 | Seed1 | 125 | 853 | 72 | 464 | 119 |
| | Seed2 | 132 | 1138 | 85 | 464 | 180 |

| | | | | | | |
|---|---|---|---|---|---|---|
| | Seed3 | 88 | 1368 | 166 | 518 | 110 |
| GROMOS 54A8 | Seed1 | 531 | 829 | 63 | 571 | 269 |
| | Seed2 | 480 | 1191 | 91 | 698 | 263 |
| | Seed3 | 351 | 1096 | 60 | 609 | 254 |
| Modelled α-proteins | | | | | | |
| | | **α-1LFC** | **α-8TFV** | **α-1KGM** | **α-1M4F** | **α-1TV0** |
| Amber 99SB-ILDN | Seed1 | 1119 | 1026 | 181 | 1003 | 749 |
| | Seed2 | 1018 | 817 | 336 | 991 | 555 |
| | Seed3 | 617 | 318 | 426 | 624 | 1167 |
| Amber 99SB*-ILDN | Seed1 | 274 | 722 | 476 | 1538 | 807 |
| | Seed2 | 517 | 377 | 666 | 632 | 459 |
| | Seed3 | 658 | 727 | 312 | 997 | 688 |
| CHARMM22* | Seed1 | 382 | 320 | 793 | 420 | 584 |
| | Seed2 | 453 | 754 | 542 | 601 | 696 |
| | Seed3 | 1770 | 1294 | 753 | 1130 | 1033 |
| CHARMM36 | Seed1 | 771 | 964 | 1860 | 623 | 207 |
| | Seed2 | 586 | 508 | 1527 | 501 | 283 |
| | Seed3 | 507 | 868 | 205 | 719 | 786 |
| GROMOS 54A8 | Seed1 | 650 | 581 | 152 | 382 | 613 |
| | Seed2 | 638 | 656 | 255 | 235 | 465 |
| | Seed3 | 696 | 654 | 97 | 426 | 583 |

**Table S16**: Average secondary structure averaged over three independent simulations for all proteins.

| | **1CQ0** | | | | | | | | **β-1CQ0** | | | | | | | |
|---|---|---|---|---|---|---|---|---|---|---|---|---|---|---|---|---|
| | Coil | β-Sheet | β-Bridge | Bend | Turn | α-Helix | π-Helix | $3_{10}$-Helix | Coil | β-Sheet | β-Bridge | Bend | Turn | α-Helix | π-Helix | $3_{10}$-Helix |
| PDB | 18 | 0 | 0 | 14 | 11 | 57 | 0 | 0 | | | | | | | | |
| Amber 99SB-ILDN | 29 | 0 | 1 | 17 | 17 | 30 | 0 | 6 | 45 | 12 | 4 | 18 | 15 | 1 | 0 | 4 |
| Amber 99SB*-ILDN | 28 | 0 | 0 | 13 | 22 | 32 | 0 | 5 | 48 | 12 | 4 | 18 | 12 | 2 | 0 | 5 |
| CHARMM22* | 30 | 0 | 0 | 13 | 17 | 36 | 0 | 3 | 45 | 11 | 2 | 21 | 16 | 0 | 1 | 4 |
| CHARM36 | 28 | 0 | 0 | 10 | 9 | 51 | 0 | 2 | 58 | 4 | 3 | 26 | 8 | 0 | 0 | 0 |
| GROMOS 54A8 | 30 | 0 | 1 | 14 | 6 | 46 | 3 | 0 | 47 | 10 | 6 | 26 | 10 | 0 | 0 | 1 |
| | **1L2Y** | | | | | | | | **β-1L2Y** | | | | | | | |
| | Coil | β-Sheet | β-Bridge | Bend | Turn | α-Helix | π-Helix | $3_{10}$-Helix | Coil | β-Sheet | β-Bridge | Bend | Turn | α-Helix | π-Helix | $3_{10}$-Helix |
| PDB | 30 | 0 | 0 | 0 | 15 | 35 | 0 | 20 | | | | | | | | |
| Amber 99SB-ILDN | 27 | 0 | 0 | 13 | 11 | 33 | 0 | 16 | 48 | 0 | 0 | 35 | 12 | 2 | 0 | 3 |
| Amber 99SB*-ILDN | 28 | 0 | 0 | 13 | 11 | 33 | 0 | 16 | 46 | 0 | 3 | 29 | 17 | 4 | 0 | 4 |
| CHARMM22* | 31 | 0 | 0 | 0 | 21 | 44 | 0 | 4 | 49 | 24 | 2 | 25 | 14 | 1 | 0 | 3 |
| CHARM36 | 28 | 0 | 0 | 2 | 20 | 46 | 0 | 4 | 65 | 0 | 0 | 27 | 5 | 2 | 0 | 2 |
| GROMOS 54A8 | 38 | 0 | 0 | 13 | 11 | 35 | 1 | 1 | 51 | 1 | 4 | 34 | 8 | 2 | 0 | 5 |
| | **1ROO** | | | | | | | | **β-1ROO** | | | | | | | |
| | Coil | β-Sheet | β-Bridge | Bend | Turn | α-Helix | π-Helix | $3_{10}$-Helix | Coil | β-Sheet | β-Bridge | Bend | Turn | α-Helix | π-Helix | $3_{10}$-Helix |
| PDB | 31 | 0 | 0 | 6 | 23 | 31 | 0 | 9 | | | | | | | | |
| Amber 99SB-ILDN | 30 | 0 | 0 | 16 | 21 | 23 | 3 | 6 | 43 | 6 | 6 | 30 | 13 | 1 | 0 | 5 |
| Amber 99SB*-ILDN | 32 | 0 | 1 | 10 | 22 | 20 | 10 | 5 | 47 | 7 | 4 | 23 | 17 | 0 | 0 | 2 |
| CHARMM22* | 31 | 0 | 1 | 11 | 21 | 21 | 10 | 4 | 50 | 5 | 2 | 25 | 15 | 8 | 0 | 1 |
| CHARM36 | 35 | 0 | 0 | 10 | 15 | 27 | 9 | 4 | 61 | 3 | 1 | 25 | 10 | 0 | 0 | 2 |
| GROMOS 54A8 | 33 | 0 | 0 | 13 | 22 | 22 | 7 | 2 | 47 | 11 | 4 | 27 | 9 | 2 | 0 | 1 |
| | **1T5Q** | | | | | | | | **β-1T5Q** | | | | | | | |
| | Coil | β-Sheet | β-Bridge | Bend | Turn | α-Helix | π-Helix | $3_{10}$-Helix | Coil | β-Sheet | β-Bridge | Bend | Turn | α-Helix | π-Helix | $3_{10}$-Helix |
| PDB | 13 | 0 | 0 | 7 | 3 | 77 | 0 | 0 | | | | | | | | |
| Amber 99SB-ILDN | 29 | 0 | 1 | 13 | 22 | 21 | 0 | 14 | 33 | 33 | 0 | 15 | 12 | 1 | 0 | 6 |
| Amber 99SB*-ILDN | 22 | 0 | 0 | 9 | 27 | 35 | 0 | 7 | 29 | 36 | 0 | 14 | 12 | 0 | 0 | 8 |
| CHARMM22* | 22 | 0 | 0 | 10 | 20 | 41 | 0 | 7 | 48 | 21 | 1 | 20 | 8 | 2 | 0 | 1 |
| CHARM36 | 15 | 0 | 0 | 2 | 4 | 78 | 0 | 0 | 55 | 15 | 2 | 21 | 6 | 5 | 0 | 1 |
| GROMOS 54A8 | 22 | 0 | 0 | 6 | 7 | 63 | 2 | 1 | 36 | 28 | 3 | 27 | 7 | 0 | 0 | 0 |
| | **1V1D** | | | | | | | | **β-1V1D** | | | | | | | |
| | Coil | β-Sheet | β-Bridge | Bend | Turn | α-Helix | π-Helix | $3_{10}$-Helix | Coil | β-Sheet | β-Bridge | Bend | Turn | α-Helix | π-Helix | $3_{10}$-Helix |
| PDB | 32 | 0 | 0 | 3 | 10 | 55 | 0 | 0 | | | | | | | | |
| Amber 99SB-ILDN | 41 | 0 | 0 | 3 | 10 | 45 | 0 | 1 | 51 | 11 | 4 | 24 | 9 | 0 | 0 | 1 |
| Amber 99SB*-ILDN | 36 | 0 | 0 | 5 | 11 | 46 | 0 | 1 | 46 | 10 | 5 | 26 | 12 | 0 | 0 | 3 |
| CHARMM22* | 41 | 0 | 0 | 4 | 11 | 43 | 0 | 1 | 46 | 13 | 8 | 24 | 9 | 0 | 0 | 2 |
| CHARM36 | 40 | 0 | 0 | 2 | 7 | 51 | 0 | 0 | 53 | 8 | 5 | 23 | 9 | 0 | 0 | 0 |
| GROMOS 54A8 | 41 | 0 | 0 | 5 | 5 | 50 | 0 | 0 | 49 | 17 | 3 | 23 | 7 | 0 | 0 | 1 |

| | 1LFC | | | | | | | | α-1LFC | | | | | | | |
|---|---|---|---|---|---|---|---|---|---|---|---|---|---|---|---|---|
| | Coil | β-Sheet | β-Bridge | Bend | Turn | α-Helix | π-Helix | $3_{10}$-Helix | Coil | β-Sheet | β-Bridge | Bend | Turn | α-Helix | π-Helix | $3_{10}$-Helix |
| PDB | 52 | 8 | 24 | 8 | 8 | 0 | 0 | 0 | | | | | | | | |
| Amber 99SB-ILDN | 22 | 62 | 0 | 12 | 3 | 0 | 0 | 0 | 40 | 0 | 0 | 14 | 19 | 16 | 0 | 10 |
| Amber 99SB*-ILDN | 23 | 62 | 0 | 9 | 5 | 0 | 0 | 2 | 39 | 0 | 3 | 12 | 17 | 22 | 0 | 8 |
| CHARMM22* | 19 | 65 | 0 | 9 | 6 | 0 | 0 | 1 | 48 | 0 | 0 | 12 | 16 | 15 | 0 | 8 |
| CHARM36 | 29 | 54 | 1 | 17 | 1 | 0 | 0 | 0 | 42 | 0 | 0 | 7 | 10 | 38 | 2 | 2 |
| GROMOS 54A8 | 31 | 53 | 1 | 11 | 4 | 0 | 0 | 0 | 33 | 0 | 0 | 15 | 10 | 35 | 5 | 2 |
| | **8TFV** | | | | | | | | **α-8TFV** | | | | | | | |
| | Coil | β-Sheet | β-Bridge | Bend | Turn | α-Helix | π-Helix | $3_{10}$-Helix | Coil | β-Sheet | β-Bridge | Bend | Turn | α-Helix | π-Helix | $3_{10}$-Helix |
| PDB | 43 | 0 | 38 | 0 | 5 | 0 | 0 | 14 | | | | | | | | |
| Amber 99SB-ILDN | 52 | 29 | 0 | 5 | 14 | 0 | 0 | 0 | 45 | 1 | 11 | 21 | 14 | 1 | 0 | 7 |
| Amber 99SB*-ILDN | 56 | 23 | 0 | 8 | 12 | 0 | 0 | 0 | 37 | 0 | 8 | 23 | 16 | 9 | 0 | 6 |
| CHARMM22* | 59 | 10 | 3 | 7 | 19 | 0 | 0 | 3 | 42 | 0 | 20 | 25 | 8 | 0 | 0 | 5 |
| CHARM36 | 71 | 3 | 1 | 13 | 10 | 0 | 0 | 0 | 52 | 0 | 23 | 15 | 7 | 0 | 0 | 2 |
| GROMOS 54A8 | 64 | 6 | 3 | 15 | 10 | 0 | 0 | 1 | 47 | 1 | 2 | 22 | 17 | 8 | 0 | 3 |
| | **1KGM** | | | | | | | | **α-1KGM** | | | | | | | |
| | Coil | β-Sheet | β-Bridge | Bend | Turn | α-Helix | π-Helix | $3_{10}$-Helix | Coil | β-Sheet | β-Bridge | Bend | Turn | α-Helix | π-Helix | $3_{10}$-Helix |
| PDB | 54 | 6 | 11 | 14 | 14 | 0 | 0 | 0 | | | | | | | | |
| Amber 99SB-ILDN | 46 | 27 | 1 | 14 | 13 | 0 | 0 | 0 | 37 | 0 | 8 | 26 | 18 | 2 | 1 | 8 |
| Amber 99SB*-ILDN | 45 | 30 | 1 | 10 | 15 | 0 | 0 | 0 | 38 | 0 | 2 | 23 | 24 | 6 | 1 | 6 |
| CHARMM22* | 44 | 29 | 0 | 14 | 13 | 0 | 0 | 0 | 43 | 0 | 9 | 20 | 16 | 7 | 0 | 5 |
| CHARM36 | 47 | 27 | 2 | 12 | 12 | 0 | 0 | 0 | 53 | 1 | 0 | 26 | 9 | 10 | 0 | 1 |
| GROMOS 54A8 | 50 | 2 | 10 | 27 | 11 | 0 | 0 | 0 | 36 | 1 | 2 | 26 | 19 | 14 | 0 | 2 |
| | **1M4F** | | | | | | | | **α-1M4F** | | | | | | | |
| | Coil | β-Sheet | β-Bridge | Bend | Turn | α-Helix | π-Helix | $3_{10}$-Helix | Coil | β-Sheet | β-Bridge | Bend | Turn | α-Helix | π-Helix | $3_{10}$-Helix |
| PDB | 36 | 0 | 32 | 32 | 0 | 0 | 0 | 0 | | | | | | | | |
| Amber 99SB-ILDN | 42 | 29 | 1 | 25 | 3 | 0 | 0 | 0 | 45 | 1 | 3 | 23 | 21 | 4 | 0 | 4 |
| Amber 99SB*-ILDN | 44 | 28 | 0 | 21 | 6 | 0 | 0 | 2 | 36 | 0 | 0 | 19 | 27 | 6 | 0 | 11 |
| CHARMM22* | 44 | 30 | 0 | 22 | 4 | 0 | 0 | 0 | 40 | 1 | 1 | 17 | 22 | 16 | 0 | 4 |
| CHARM36 | 51 | 27 | 0 | 19 | 3 | 0 | 0 | 0 | 31 | 0 | 0 | 8 | 12 | 47 | 0 | 2 |
| GROMOS 54A8 | 40 | 35 | 1 | 15 | 9 | 0 | 0 | 1 | 33 | 1 | 0 | 23 | 14 | 25 | 2 | 1 |
| | **1TV0** | | | | | | | | **α-1TV0** | | | | | | | |
| | Coil | β-Sheet | β-Bridge | Bend | Turn | α-Helix | π-Helix | $3_{10}$-Helix | Coil | β-Sheet | β-Bridge | Bend | Turn | α-Helix | π-Helix | $3_{10}$-Helix |
| PDB | 41 | 0 | 31 | 28 | 0 | 0 | 0 | 0 | | | | | | | | |
| Amber 99SB-ILDN | 39 | 29 | 2 | 20 | 10 | 0 | 0 | 1 | 44 | 0 | 0 | 18 | 16 | 9 | 0 | 12 |
| Amber 99SB*-ILDN | 41 | 28 | 2 | 21 | 9 | 0 | 0 | 1 | 34 | 0 | 0 | 14 | 19 | 24 | 2 | 7 |
| CHARMM22* | 44 | 16 | 4 | 23 | 13 | 0 | 0 | 1 | 36 | 3 | 1 | 16 | 18 | 20 | 1 | 6 |
| CHARM36 | 49 | 18 | 3 | 26 | 4 | 0 | 0 | 0 | 37 | 0 | 0 | 11 | 10 | 42 | 0 | 1 |
| GROMOS 54A8 | 32 | 39 | 1 | 20 | 8 | 0 | 0 | 0 | 38 | 0 | 2 | 14 | 9 | 35 | 1 | 2 |

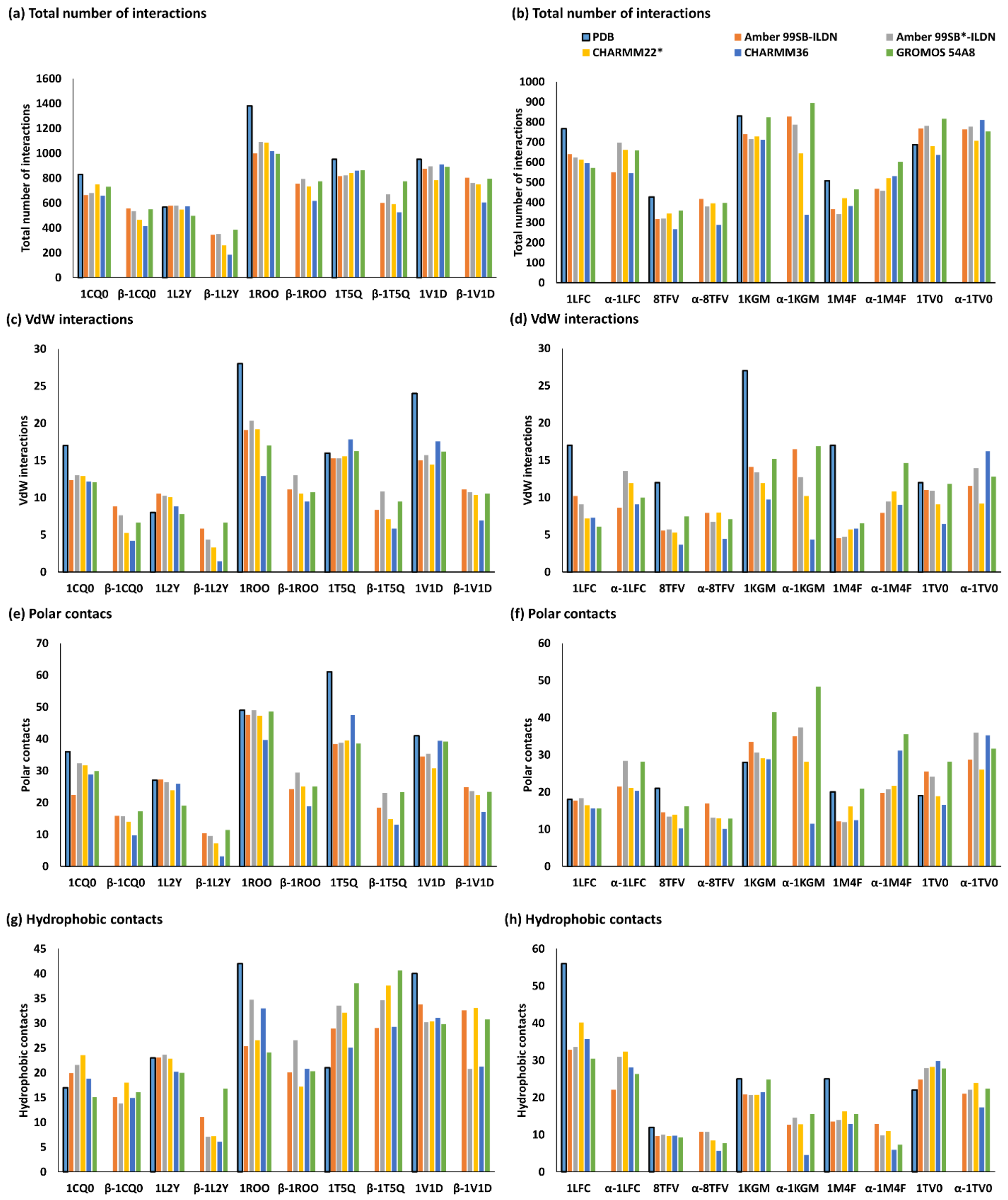


**Figure S1. Averaged number of van der Walls interactions, polar contacts, hydrophobic contacts, ionic interactions, and aromatic contacts.** (a/b) Average total interactions. (c/d) van der Walls interactions. (e/f) Polar contacts. (g/h) Hydrophobic contacts.
